# Correlative three-dimensional super-resolution and block face electron microscopy of whole vitreously frozen cells

**DOI:** 10.1101/773986

**Authors:** David P. Hoffman, Gleb Shtengel, C. Shan Xu, Kirby R. Campbell, Melanie Freeman, Lei Wang, Daniel E. Milkie, H. Amalia Pasolli, Nirmala Iyer, John A. Bogovic, Daniel R. Stabley, Abbas Shirinifard, Song Pang, David Peale, Kathy Schaefer, Wim Pomp, Chi-Lun Chang, Jennifer Lippincott-Schwartz, Tom Kirchhausen, David J. Solecki, Eric Betzig, Harald Hess

## Abstract

Living cells function through the spatial compartmentalization of thousands of distinct proteins serving a multitude of diverse biochemical needs. Correlative super-resolution (SR) fluorescence and electron microscopy (EM) has emerged as a pathway to directly view nanoscale protein relationships to the underlying global ultrastructure, but has traditionally suffered from tradeoffs of structure preservation, fluorescence retention, resolution, and field of view. We developed a platform for three-dimensional correlative cryogenic SR and focused ion beam milled block-face EM across entire vitreously frozen cells that addresses these issues by preserving native ultrastructure and enabling independent SR and EM workflow optimization. Application to a variety of biological systems revealed a number of unexpected protein-ultrastructure relationships and underscored the value of a comprehensive multimodal view of ultrastructural variability across whole cells.

## Main Text

Electron microscopy (EM) has revealed an intricate world inside eukaryotic cells (*1*), spatially organized at all length scales from nanometer-sized molecular assemblies to cell-spanning structures such as actin stress fibers and microtubules. However, even within different regions of the cell, there are notable differences in the structure of individual components, such as nuclear chromatin organization (*2*) or the morphology of the endoplasmic reticulum (ER), which is highly convoluted and compact in the perinuclear region, yet sparsely reticulated in the lamellae (*1*). Thus, a comprehensive picture of cellular organization requires nanometer-level three-dimensional (3D) imaging of whole cells.

While cryogenic (cryo)-EM / tomography offers sub-nanometer 3D resolution (*3*), it is limited to sparse deposits of extracted macromolecules, cellular sections of sub-micron thickness (*4–7*), or thin lamella sculpted with cryo focused ion beam (FIB) milling (*8, 9*). In contrast, serial FIB ablation and imaging of the exposed face of resin-embedded specimens by scanning electron microscopy (FIB-SEM) routinely achieves 8 nm isotropic 3D resolution (*10–12*) not possible with traditional 3D EM by diamond knife serial array (*13, 14*) or block face sectioning (*15*). However, EM produces grayscale images in which the unambiguous identification and 3D segmentation of many subcellular structures can be challenging, and where the distributions of specific proteins can rarely be identified.

In response, correlative light and electron microscopy (CLEM) techniques have been developed that combine the global contrast and high resolution of EM with the molecular specificity of fluorescence microscopy (*16, 17*). With the advent of super-resolution (SR) microscopy (*18*), such techniques now offer a closer match in resolution between the two modalities (supplementary note 1, table S1), allowing specific molecular components to be visualized at nanoscale resolution in the context of the crowded intracellular environment. However, SR/EM correlation often involves tradeoffs in sample preparation between the retention of fluorescent labels, sufficiently dense heavy metal staining for high contrast EM, and faithful preservation of ultrastructure, particularly when chemical fixation is used (*19–22*).

Here we describe a pipeline (fig. S1) for correlative cryo-SR/FIB-SEM imaging of whole cells designed to address these issues. Specifically, cryogenic, as opposed to room temperature, SR performed after high pressure freezing (HPF), allowed us to use a standard EM sample preparation protocol without compromise. We used cryogenic 3D structured illumination (SIM) and single molecule localization (SMLM) microscopy for SR protein specific contrast with 3D FIB-SEM for global contrast of subcellular ultrastructure. The SR modality highlights features not readily apparent from the EM data alone, such as exceptionally long or convoluted endosomes, and permits unique classification of vesicles of like morphology, such as lysosomes, peroxisomes, and mitochondrial-derived vesicles. Cell-wide 3D correlation also reveals unexpected localization patterns of proteins, including intranuclear vesicles positive for an ER marker, intricate web-like structures of adhesion proteins at cell-cell junctions, and heterogeneity in euchromatin or heterochromatin recruitment of transcriptionally-associated histone H3.3 and heterochromatin protein 1α (HP1α) in the nuclei of neural progenitor cells as they transition into differentiated neurons). More generally, whole cell cryo-SR/FIB-SEM can reveal compartmentalized proteins within known subcellular components, help discover new subcellular components, and classify unknown EM morphologies and their roles in cell biology.

### Cryogenic SR below 10K: motivations and photophysical characterization

To avoid artifacts associated with chemical fixation (fig. S2), our pipeline begins with cryo-fixation *via* HPF (*23, 24*) of whole cells cultured on 3 mm diameter, 50 µm thick sapphire disks (supplementary note 2). Unlike plunge freeze methods, HPF reliably freezes specimens hundreds of microns thick (*21, 23, 25, 26*) in their entirety within vitreous ice in milliseconds, providing an exact snapshot of subcellular ultrastructure (fig. S3, movie S1). Each sapphire disk provides an optically flat and transparent back surface for aberration-free SR imaging, along with the high thermal conductivity needed to minimize specimen heating and potential ice re-crystallization under the intense (∼kW/cm^2^), long-lasting illumination used during SMLM. Frozen specimens are inspected, cleaned (movie S2), and loaded onto a solid copper sample holder (fig. S4) in a covered, liquid nitrogen (LN_2_) cooled preparation chamber (fig. S5) before transfer through a load lock to an evacuated optical cryostat modified for SR imaging (fig. S6).

Cryo-SR benefits from a dramatic increase in fluorophore photostability (*27*). This allowed us to achieve the high photon counts required for precise single molecule localization, despite the modest numerical aperture (NA 0.85) we were compelled to use in order to image through the cryostat window, vacuum, and sapphire substrate (fig. S1, supplementary note 3). This along with a remarkably high reactivation efficiency under 405 nm illumination (*28, 29*) allowed us to acquire multicolor SIM/SMLM images of the same cells without substantial photobleaching. Which, in turn, enabled SIM/SMLM correlation in three or more colors (movie S3) as well as the ability to quickly image and assess many cells across the substrate by 3D SIM before concentrating on the best candidates for the much slower, yet even higher resolution, imaging by 3D SMLM.

Most cryo-SR systems to date operate with LN_2_ cooling near 77K (*7, 27, 29–34*). However, we opted for a liquid helium (LHe) cooled microscope, which allowed us to explore photophysics at any temperature down to 8K (supplementary note 4). In particular, we were able to exploit a sharp increase in the lifetime of a dark state D_1_ for many fluorescent molecules with decreasing temperature (fig. S7), allowing them to be efficiently shelved for long periods. Such shelving has important implications for SMLM, as it dictates the dynamic contrast ratio (DCR) defined by the time a given molecule is OFF and shelved in the dark state normalized to the time it is ON and cycling between singlet states S_0_ and S_1_ (fig. S7) to emit light. Molecules with high DCR can be expressed at higher density, creating SMLM images of higher fidelity and resolution, with less chance of spontaneous overlap of the diffraction spots from multiple molecules that would otherwise hinder precise localization.

We measured (Fig. 1A) the DCR of six different fluorophores at both 8K and 77K from the ON/OFF blinking behavior of isolated single molecules (fig. S8, supplementary note 4a). In addition, we compared (Fig. 1B) their static contrast ratios (SCR, defined by the ratio of their signal in the ON state to their local background, fig. S9), which must also be high for precise localization, during SMLM imaging of densely labeled mitochondria (Movie 1, fig. S9, supplementary note 4b). Notably, DCR and SCR tended to increase with shorter emission wavelengths, making such fluorophores better suited to high quality SMLM imaging (Fig. 1C). SCR also often improved at lower temperature (Fig. 1B). These trends are consistent with the photophysical argument that the dark state lifetime should increase with increasing energy from D_1_ to S_0_, normalized to the thermal energy. In particular, we observed substantial gains in the SCR and DCR of JF525 (*35*) when operating with LHe which, in conjunction with mEmerald, enabled high quality two color SMLM of densely labeled structures. However, if only cryo-SIM and/or single color cryo-SMLM is needed, or if further study uncovers fluorophores spectrally distinct from mEmerald that work just as well at 77K, then operation with LN_2_ may prove sufficient.

**Figure 1.**
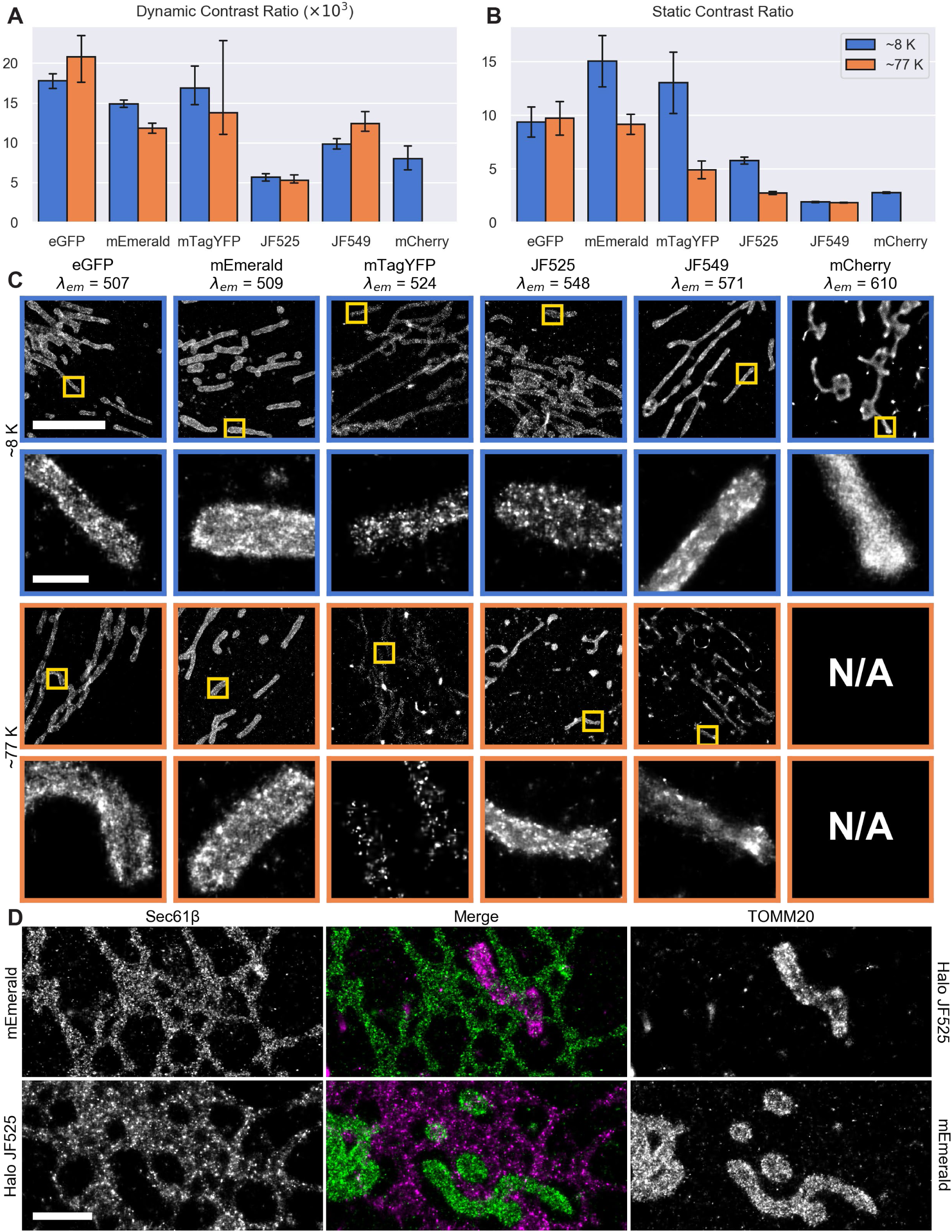
Cryogenic photophysical characterization of fluorophores for single molecule localization microscopy (SMLM) **(A)** Dynamic and **(B)** static contrast ratios for six different fluorophores at ∼8K (blue) and ∼77K (orange) ordered by increasing emission wavelength. **(C)** Corresponding images of mitochondrial outer membrane protein TOMM20 with detailed insets shown below. **(D)** Comparison of mEmerald and JF525 in cryo-SMLM imaging. Top row: U2OS cell transiently expressing ER membrane marker mEmerald-Sec61β and mitochondrial membrane marker Halo-TOMM20 conjugated to JF525. Bottom row: U2OS cell transiently expressing mEmerald-TOMM20 and Halo/JF525-Sec61β. Scale bars: 5 μm and 0.5 μm in (C), first and second rows; 1 μm in (D).

To compare the relative merits of these labels for cryo-SMLM, we imaged two U2OS cells, the first (Fig. 1D, top row) targeting the ER membrane (green) with mEmerald and the mitochondrial outer membrane (magenta) with Halo-JF525 (*36*), and the second having these labels switched between the same two targets (Fig 1D, bottom row). While both labels produced high density, high precision SMLM images of both targets, the Halo-JF525 images exhibited numerous bright puncta in both cases (Fig. 1D, fig. S10B). Although these may result from aggregation of Halo-tagged proteins, the presence of similar puncta in cryo-SMLM images of the ER obtained via SNAP (*37*) or CLIP (*38*) tag targeting of JF525 (fig. S10C, D) suggest that they arise from a subset of extremely long-lived JF525 molecules which undergo numerous switching cycles. Indeed, the long persistence of JF525 and other labels at 8K necessitated a fresh, data-driven approach (supplementary note 5b, fig S11) to the problem of correctly assigning and integrating the multiple photon bursts from each molecule. Even so, in all cases mEmerald yielded images probably more reflective of the true molecular distribution.

### Cell-wide 3D correlation of cryo-SR with FIB-SEM

A key advantage of our pipeline is that inserting the cryo-SR step between cryofixation and freeze substitution / staining for FIB-SEM allowed us to decouple the sample preparation protocols for the two imaging modalities, thereby avoiding the tradeoffs of fluorescence retention, dense heavy metal staining, and ultrastructure preservation of resin-embedding based protocols (*39–42*). Another is that it allowed us to rapidly (∼20 min) survey hundreds of cells across the sapphire disk (fig S12A), identify those of promising morphology and expression levels, inspect these further at higher resolution by 3D cryo-SIM (5 min/cell/color), and select the very best ones of these for the ∼1–2 days/cell/color required for 3D cryo-SMLM and ∼5-10 days/cell needed for EM sample preparation and FIB-SEM imaging.

Thus, after cryo-SR imaging, we removed the frozen, disk-mounted specimens from the cryostat (fig. S12B) and processed them (supplementary note 6) via freeze substitution (*19–21*), heavy metal staining, and embedding in Eponate 12 resin (fig S12C). After coarse trimming of the resin block and removal of the sapphire disk, we re-embedded the remaining tab in Durcupan, imaged it with x-rays (fig. S12D), and correlated this to the original disk-wide view (fig S12E) to identify the region (typically 100 × 100 × 10 μm) containing the cells of interest imaged previously with cryo-SR. Additional microtome trimming isolated this region (fig. S12F-I), which we then imaged at 4-8 nm isotropic voxels in 3D by FIB-SEM (*12*).

To exploit the full potential of correlative microscopy, the different imaging modalities need to be mutually registered to the level of their spatial resolution. Given the high resolution of both cryo-SMLM and FIB-SEM, and our desire to extend their correlation across whole cells in 3D, registration to this level is challenging. For example, slight magnification differences or deviations from ideal flat field and rectilinear imaging coupled with potentially non-uniform FIB milling increase registration errors quickly with increasing field of view. Furthermore, freeze substitution and resin embedding introduce nonlinear and spatially inhomogeneous sample deformations (arrows, Fig. 2A) between the cryo-SR and FIB-SEM imaging steps that have a substantial non-affine component requiring deformable registration to achieve alignment to this level of accuracy.

**Figure 2.**
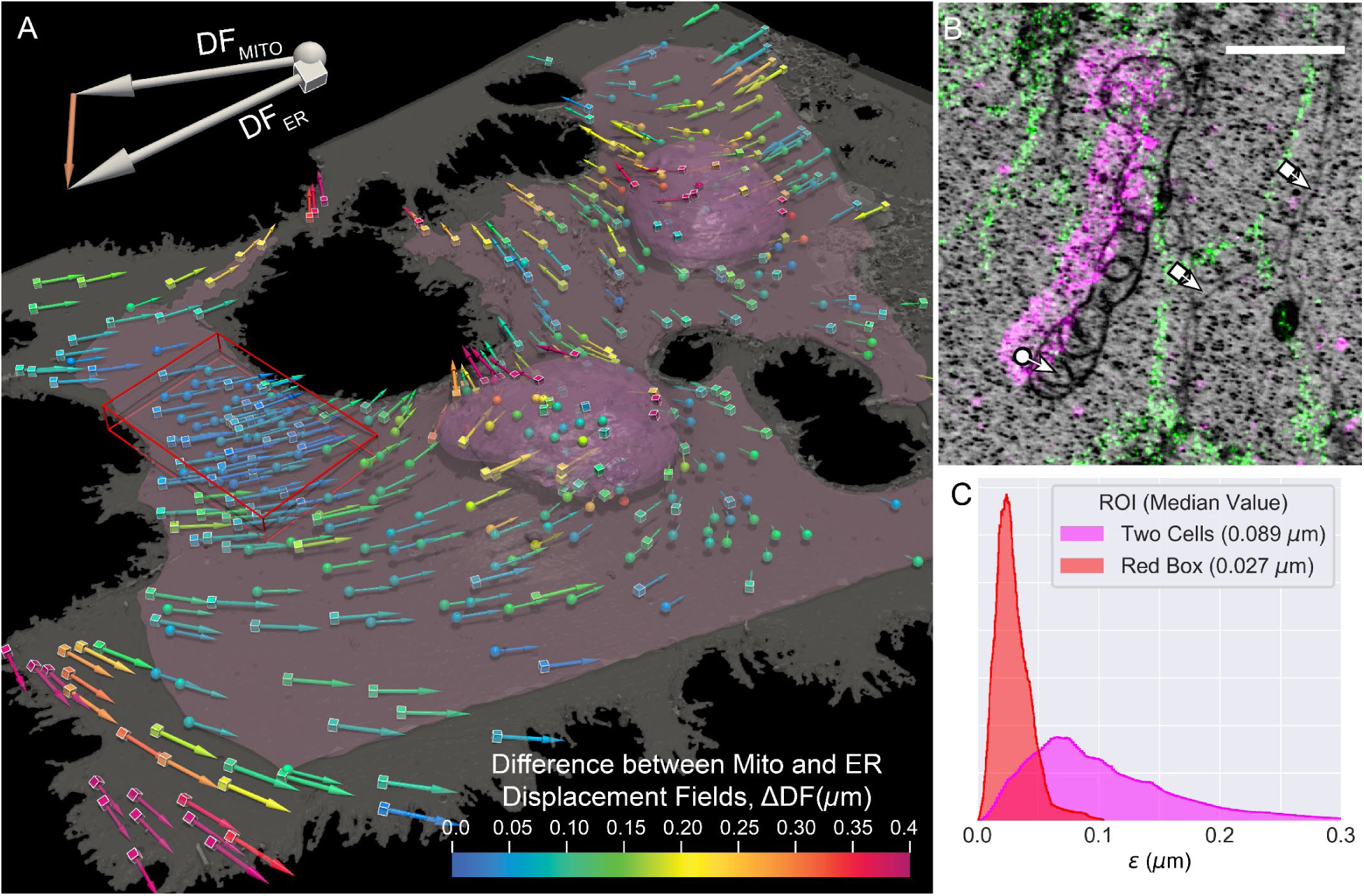
High accuracy correlation of cryo-SMLM and FIB-SEM data using organelle landmarks. **(A)** A perspective view of mitochondrial (spheres) and ER (cubes) landmarks used for registration along with the plasma (grey) and nuclear (pink) membranes as determined by FIB-SEM of two COS-7 cells. Arrows point in the direction of and are sized according to the magnitude of the non-affine component of the final displacement field. Arrows are color coded according to the magnitude of the local difference (ΔDF) between the displacement fields determined by the mitochondrial or the ER landmarks separately. The pink surface indicates the boundaries of the sub-volume containing sufficient landmarks of both types for quantitative comparison of their respective displacement fields. **(B)** XY orthoslice illustrating landmark selection and determination of the displacement vectors (fig. S14). Scale bar: 1 µm. **(C)** Histograms of the correlation accuracy, *ε* (cf. supplementary note 7), for the sub-volume bounded by the pink surface (magenta) and the 61 µm^3^ sub-volume defined by the red box (red), where density of both types of landmarks is higher.

Taking advantage of our protocol, we densely labeled ubiquitous intracellular organelles such as the ER and mitochondria that could be readily identified in both the cryo-SMLM and FIB-SEM data and used them as landmarks (e.g., Fig. 2B, fig. S13) to register the two modalities across the cellular volume (supplementary note 7), using the software package BigWarp (*43*). To quantify the accuracy of this correlation, we independently measured the deformation fields 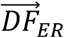 and 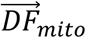 from only ER or mitochondrial landmarks, respectively, after aligning these color channels to one another using fluorescent bead fiducial markers. Since 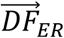 and 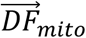 represent independent estimates of the underlying sample deformation, the correlation accuracy ε is given by 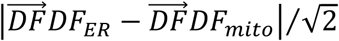 (supplementary note 7). Over a field of view covering the majority of two cells (pink surface, Fig. 2A), we measured a median ε of 89 nm (Fig. 2C), whereas in a small peripheral region (red box, Fig. 2A), the median ε was 27 nm. The difference may be attributable to a higher density of landmarks within the peripheral region, tighter physical constraint on the differential motion between organelles due to the thinness of this region, or greater accuracy in landmark displacement measurement when the sample thickness becomes less than the axial localization precision. Regardless, spatial maps of ε (fig. S14) give a local estimate of the length scale to which spatial relationships between features seen by the two imaging modalities can be reliably inferred.

### Correlative cryo-SR/FIB-SEM reveals vesicle identities and their morphological diversity

Using our correlative pipeline, we imaged two neighboring COS-7 cells (Fig. 3, Movie 2) transiently expressing mEmerald-ER3, a luminal ER marker, and HaloTag-TOMM20 conjugated to JF525 to label the mitochondrial outer membrane. Both the resulting volume rendering (Fig. 3A) and axial or transverse orthoslices (Figs. 3B-M, fig. S15) demonstrate accurate 3D correlation of the cryo-SMLM and FIB-SEM data, high labeling density, and faithful ultrastructure preservation throughout the ∼5000 µm^3^ cellular volume within the field of view, validating the expectation of uncompromised quality in both SR and FIB-SEM data.

**Figure 3.**
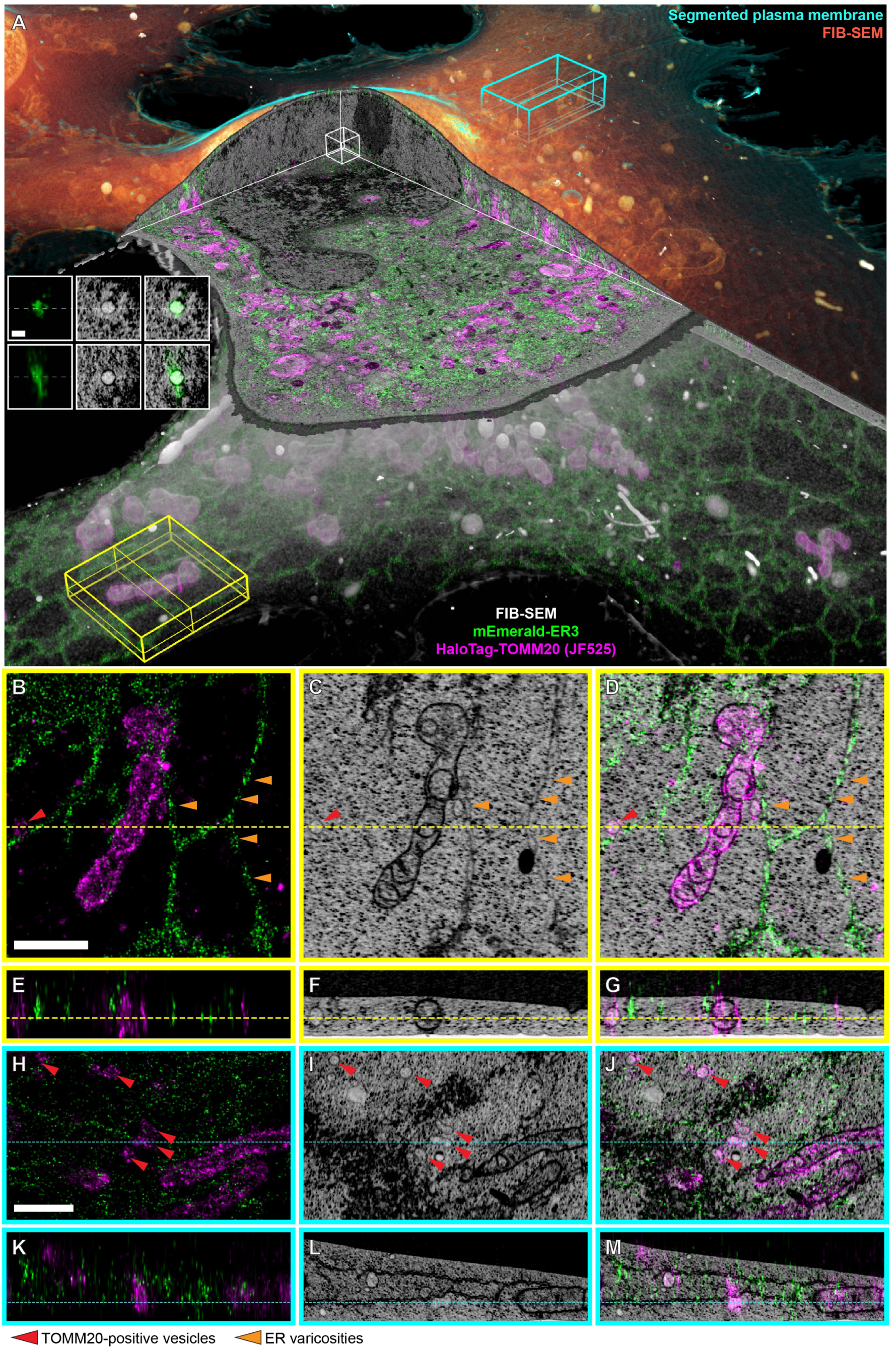
Whole-cell correlative cryogenic single molecule localization and block face electron microscopy. **(A)** Perspective overview of a cryo-SMLM and FIB-SEM (orange and grey) data set of a COS-7 cell transiently expressing mEmerald-ER3 (ER lumen marker, green) and Halo/JF525-TOMM20 (mitochondrial outer membrane marker, magenta) (Movie 2). Cyan, yellow, and white boxes indicate regions with ortho slices shown in panels (B-M) and inset. **Inset:** SMLM (left column), FIB-SEM (middle column), and correlative (right column) orthoslices in XY (top row) and XZ (bottom row) through an intranuclear ER-positive vesicle. Scale bar: 200 nm. **(B, E)** SMLM, **(C, F)** FIB-SEM and **(D, G)** correlated overlay of orthoslices in XY (B-D) and XZ (E-G) in a lamellipodial region. Scale bar: 1 µm. **(H, K)** SMLM, **(I, L)** FIB-SEM and **(J, M)** correlated overlay of orthoslices in XY (H-J) and XZ (K-M) in a thicker region with ER sheets. Scale bar: 1 µm. Red arrows: TOMM20-positive vesicles; orange arrows: varicosities in the ER.

The data also immediately illustrate the power of cryo-SR and FIB-SEM correlation. For example, clusters of ER3 seen by cryo-SMLM to dot ER tubules (orange arrows, Fig. 3B) might easily be dismissed as artifacts of labeling or fixation, but instead are revealed to correlate (Fig. 3D) with varicosities in these tubules as seen by FIB-SEM (Fig. 3C). It is also immediately apparent by FIB-SEM that vesicles of various sizes are ubiquitous throughout the cell. However, these can come in many forms – peroxisomes, lysosomes, endosomes or, as identified by our correlation, TOMM20-positive vesicles (red arrows, Fig. 3H-J, fig. S16). Given their small (∼100-200 nm) size and proximity to mitochondria, these may represent mitochondria derived vesicles (MDVs). MDVs are believed to play a key role in mitochondrial quality control by sequestering unfolded or oxidized mitochondrial proteins and transporting them to lysosomes or peroxisomes for degradation (*44*). There remain, however, many questions about what proteins regulate these processes and where they are distributed – questions which cryo-SR/FIB-SEM may be able to help resolve.

Our data also revealed three instances of intranuclear vesicles, again ∼100-200 nm size, positive for the ER lumen protein ER3 (left inset orthoslices, Fig. 3A; correlation examples 119 and 164, fig. S15). In dividing somatic cells, the nuclear membrane (NM) breaks down at prometaphase and NM proteins are dispersed within the ER, which remains continuous throughout mitosis (*45*). The NM then begins to reassemble in anaphase when ER-like cisternae contact the chromatids of the daughter cells, and NM proteins become immobilized there (*45–48*). Thus, one possibility is that ER lumen-positive intranuclear vesicles in interphase are the remnants of such contacts that do not completely return to the extranuclear ER after the NM is fully re-established. Alternatively, a small fraction of the total ER volume might be disrupted into vesicles during its rearrangement in mitosis and become similarly trapped when the NM reforms.

Another important class of vesicles in cells are peroxisomes, which catabolize long chain fatty acids via β-oxidation and reduce reactive oxygen species such as hydrogen peroxide (*49*). Peroxisomes can adopt a variety of sizes, shapes, and distributions depending on cell type and environment (*49, 50*), but accurately capturing these morphologies and their spatial relationship to other organelles can be difficult with traditional chemical fixation and EM staining protocols against their enzymatic contents (*51, 52*). Furthermore, serial section transmission EM lacks the axial resolution to precisely measure morphological parameters at the sub-100 nm level, whereas cryo-EM tomography lacks the field of view to explore more than a small fraction of the total cellular volume.

To get a more accurate and comprehensive look, we used cryo-SMLM / FIB-SEM to image and semi-automatically segment (supplementary note 8) 466 mature peroxisomes across two vitreously frozen HeLa cells expressing mEmerald tagged to the peroxisomal targeting sequence SKL, and Halo/JF525-TOMM20 to mark mitochondria (Fig. 4, Movie 3, and fig. S17). Independent two-channel SR/EM registration as described above revealed a correlation accuracy (median ε) of 85 nm (fig. S14C, D). Peroxisomes with volumes smaller than 0.01 µm^3^ universally assumed nearly spherical shapes to minimize their surface area under the influence of surface tension (e.g., Fig. 4A, H), but increasingly irregular shapes such as plates (Fig. 4B), cups (Fig. 4C) or hollow spheres (Fig. 4D) form with increasing volume (lower rows, fig. S17). This may serve to regulate reaction kinetics rates within them (*53*). Some irregularly shaped peroxisomes we found were in close proximity to other organelles or as part of multi-organelle assemblies (Fig. 4E-G), consistent with 3D observations in live cells (*54*). These assemblies may facilitate the transfer of cargo between organelles responsible for distinct and possibly incompatible biochemical processes (*54*), such as the sequential breakdown of fatty acids between peroxisomes and mitochondria (*55*).

**Figure 4.**
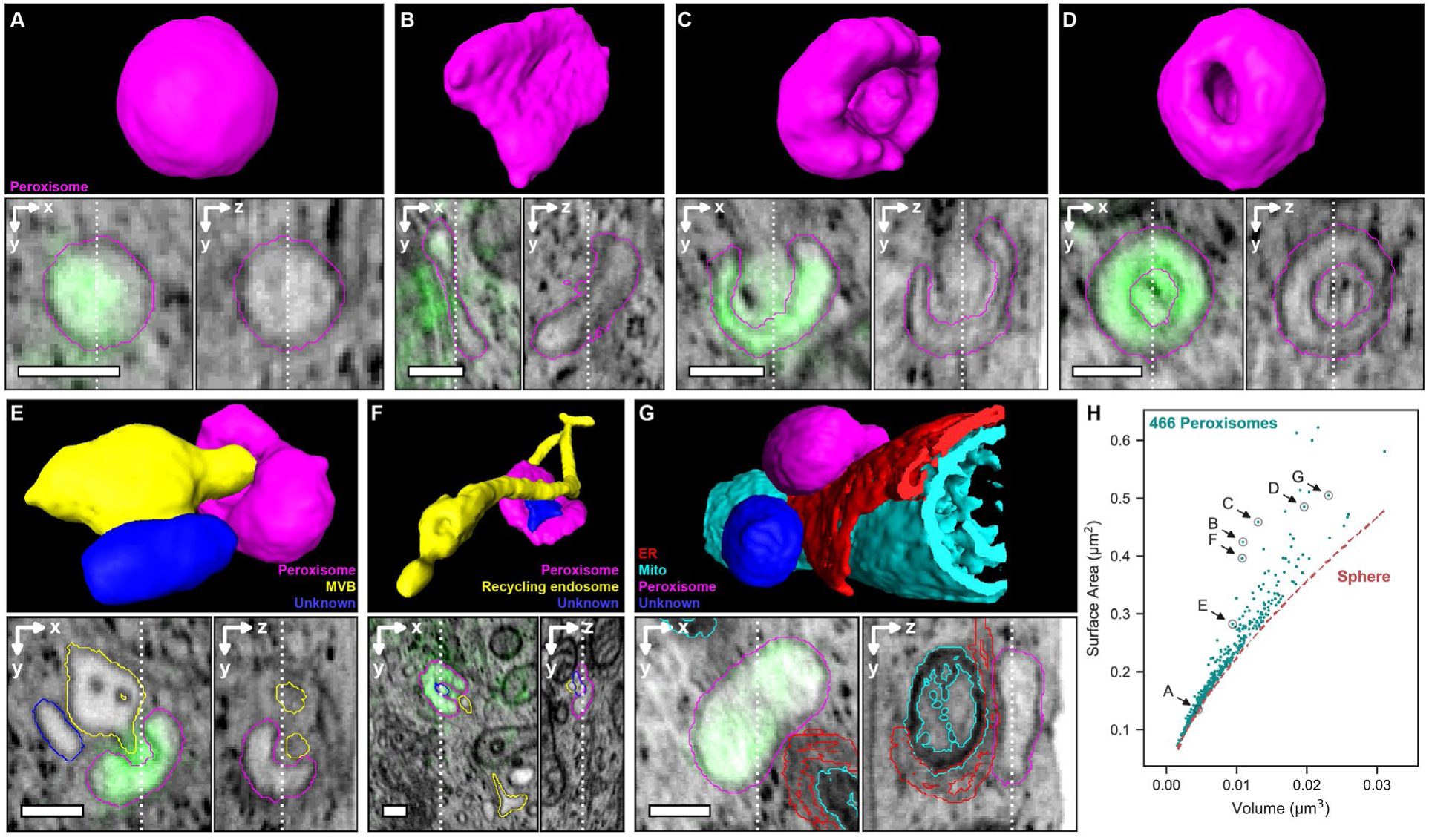
Diversity of peroxisome morphologies and peroxisome-organelle interactions. **(A-D)** FIB-SEM segmentations (top) of four peroxisomal targeting signal (SKL) containing peroxisomes (magenta) and corresponding orthoslices (bottom) with cryo-SMLM overlays of SKL (green) from two HeLa cells expressing mEmerald-SKL. **(E-G)** Three examples of peroxisome/organelle interactions, showing both segmentations (top) and orthoslices (bottom) with overlaid contours of matching colors. Scale bars: 200 nm. **(H)** Surface-to-volume relationship for 466 peroxisomes (fig. S17), with the specific peroxisomes in (A)-(G) indicated, showing the increasing deviation from spherical shape with increasing volume. See also Movie 3.

Finally, we explored the endolysosomal pathway, the compartments of which are notoriously sensitive to artifacts of fixation or protein overexpression (*56–58*). We used correlative cryo-SIM/FIB-SEM to image transferrin (Tfn)-containing endolysosomal compartments in a SUM-159 cell previously incubated for 30 min in media containing Alexa Fluor 647-conjugated Tfn (Fig. 5A, beginning of Movie 4). The density of labeled compartments was low enough to assign each discrete SIM feature (inset, Fig. 5A) to a single structure as seen by FIB-SEM, and then use the latter modality to render each such compartment with 8 nm isotropic voxels. Despite its much lower resolution, SIM was essential not only to identify which compartments in the FIB-SEM data represented endolysosomes, but also to spot the many such structures of extremely convoluted morphology in the crowded intracellular environment that were not readily apparent by FIB-SEM alone. These included elongated tubules (magenta, Figs. 5B-E) that likely represent recycling endosomes, highly corrugated endosomes (Figs. 5B, D, right), and early endosomes with protruding tubules of 50 nm width possibly associated with retromers (*59*) of sub-50 nm width. Notably, given that cryo-SIM is much faster than cryo-SMLM and can use a wide variety of spectrally distinct labels, it can be a broadly useful tool in its own right to guide the 3D segmentation of dense FIB-SEM data and ensure the correct identification of specific subcellular features.

**Figure 5.**
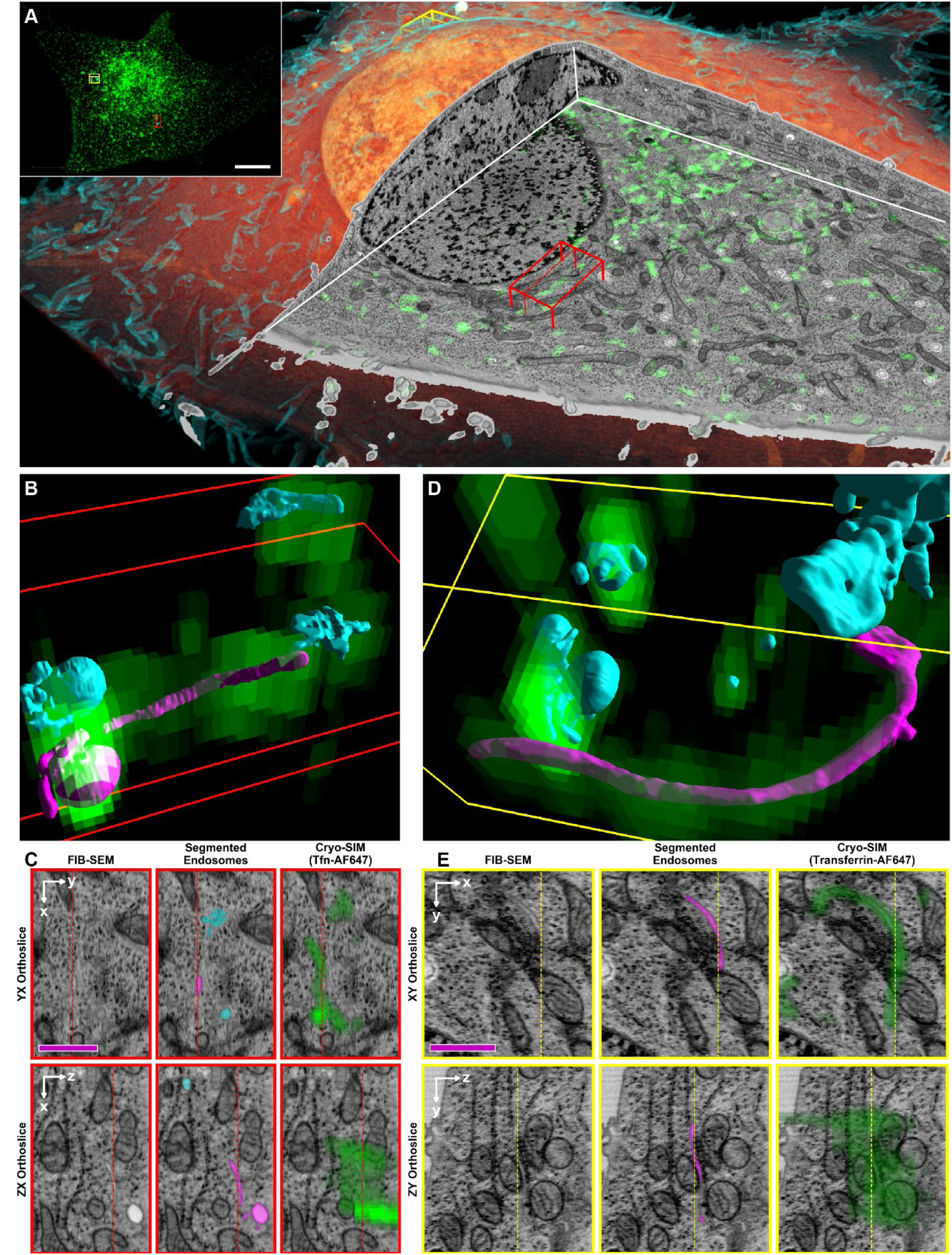
Cryo-SIM/FIB-SEM accurately identifies endosomal compartments and reveals their diverse morphologies at the nanoscale. **(A)** Volume rendered FIB-SEM overview (interior, orange; plasma membrane, cyan) of a SUM159 cell, with cutaway correlative cryo-SIM showing endolysosomal compartments containing AF647-conjugated transferrin (green). **(B)** Segmented Tfn-AF647 containing compartments (colored surfaces) with superimposed 3D cryo-SIM data (green voxels) in the 13 µm^3^ subvolume denoted by the red box in (A). **(C)** XY (top) and ZY (bottom) orthoslices of the same region in (B) showing the FIB-SEM (left) overlaid with segmentations of transferrin labeled compartments (middle) and cryo-SIM of Tfn-AF647 (right). **(D, E)** Same as (B) and (C) for the 19.5 µm^3^ subvolume denoted by the yellow box in (A). Scale bars, (A, inset) 10 µm, (C, E) 1 µm.

### Molecular underpinnings of ultrastructural specialization in neuronal cell-to-cell adhesions

Cell-to-cell adhesions are of central importance in tissue morphogenesis because they mediate cell migration, nucleate cell polarity and spur communication between individual cells in multi-cellular organisms (*60, 61*). While the molecular context and ultrastructure of cellular adhesions to rigid artificial substrates are well characterized (*62, 63*) those between cells in complex 3D environments are not. Neuronal adhesions are particularly interesting because they are crucial for brain development, playing an integral role in sorting neurons based on their maturation status (*64, 65*), forming the laminar structure of the brain (*66, 67*), and ultimately promoting the complex neuronal interactions that drive circuit morphogenesis (*68*). However, they have been difficult to study on account of their disruption by chemical fixation (*19–21*), and because the baroque 3D geometries of neuronal contacts require isotropic 3D-EM and high-resolution LM.

We used cryo-SIM to visualize transiently expressed junctional adhesion molecule (JAM)-C (*64, 66, 69*), a tight-junction component, fused to JF549i-conjugated SNAP-(*35, 70*), and 2x-mVenus-drebrin, a cytoplasmic actin-microtubule crosslinker protein (*71*), in cryofixed mouse cerebellar granule neurons (CGNs) (Fig. 6A, Movie 5). Curiously, we found that JAM-C indicated that the adhesion between two labeled somas was not uniform at their shared membrane contact zone (CZ) (Fig. 6B), but rather formed a web-like structure, and that drebrin preferentially associated with the edges of the JAM-C regions.

**Figure 6.**
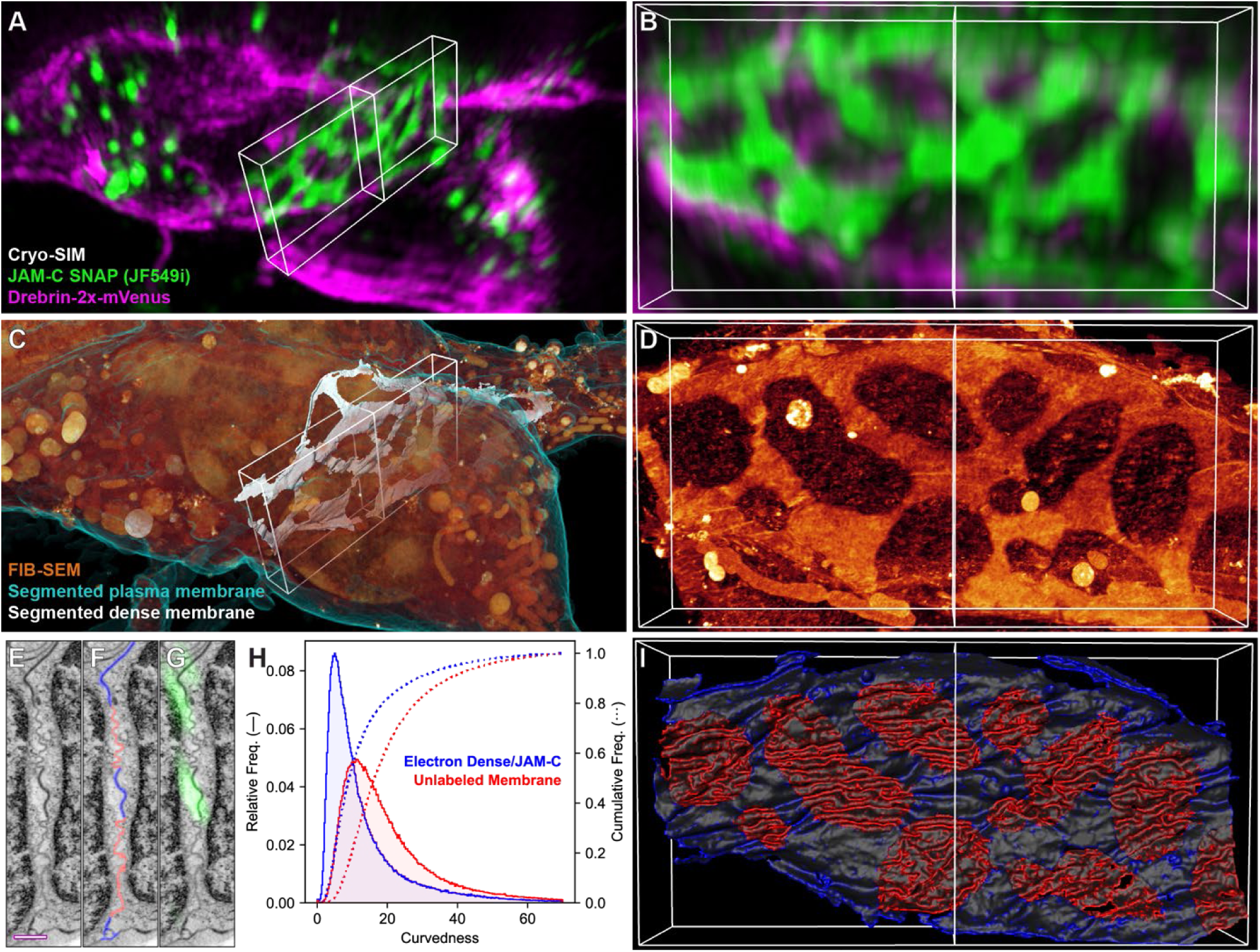
Membrane proteins correlate to membrane ultrastructure at cell-cell adhesions. **(A)** Cryo-SIM volume of cultured mouse cerebellar granule neurons transiently expressing JF549i/SNAP-JAM-C (green) and 2x-mVenus-Drebrin (magenta). **(B)** MIP through an ∼3 μm thick slab (white box in (A)) centered on the contact zone between two cell bodies. **(C)** FIB-SEM volume of the same region in (A), with plasma membrane (cyan), intracellular content (orange), and segmented electron dense regions of the contact zone (white). **(D)** FIB-SEM MIP through the same region in (B), after masking the nuclei. **(E)** Single FIB-SEM slice through the contact zone at the central vertical line in (D). **(F)** same as (E), with more (blue) and less (red) electron dense membranes traced; **(G)** same as (E) overlaid with the JAM-C signal. Scale bar: 500 nm. **(H)** Histograms of the curvedness (supplemental note 9) for the high (blue) and low (red) electron density membrane regions. **(I)** Partial segmentation of the cells’ membranes in the contact zone, color coded according to curvedness, with brighter colors indicating larger values. Note the high correlation between JAM-C (B), electron density (D), and membrane curvedness (I) (Movie 5).

To determine if these protein distributions correlated with membrane ultrastructure at the CZ, we imaged the same cells by FIB-SEM (Fig. 6C) and found that the density of heavy metal staining at the PM was also nonuniform (Fig. 6D), with the densest staining correlating perfectly with JAM-C (compare Figs. 6B, D and G). Moreover, we observed that the densely stained PM was less curved than the electron lucent PM. To quantify this, we segmented the PM within the CZ into high and low electron density regions (Figs. 6F, I), and then calculated the curvedness (supplementary note 9) in each (Figs. 6H and I), finding that the low-density PM was 2.3 times more curved than the high-density, JAM-C rich PM.

While the smooth nature of the adhesion as defined by JAM-C is expected because of the mechanical tension induced by the juxtacrine interaction (*72–74*), the fact that the adhesion does not comprise the whole contact area between these two cells is not. A further surprise is the enrichment of drebrin in the regions adjacent to JAM-C, rather than the laminar stacking of adhesion-associated cytoskeletal adaptor proteins found in focal or cadherin-based adhesions to glass (*62, 63*).

### Chromatin domains and their reorganization during neuronal differentiation

In addition to adhesion, CGNs provide an excellent model system to study the cell biological underpinnings of neural development due to their strongly stereotyped developmental programs as they differentiate from cerebellar granule neuron progenitors (GNPs) (*75*). Intrinsic to this process is the 3D structural reorganization of their nuclear chromatin domains (*76, 77*). To explore this in detail, we first used 3D live cell lattice light sheet microscopy (LLSM) (*78*) and found that flow sorted GNPs expressing the EGFP-Atoh1 marker of the GNP-state (*79, 80*) possess significantly larger nuclei than terminally differentiated CGNs (Figure 7A, B, supplementary note 10). Moreover, longitudinal LLSM live-imaging revealed that GNPs rapidly condense their nuclei to the size of CGNs while Atoh1-EGFP expression fades (Figure 7A, B, movie S6).

**Figure 7.**
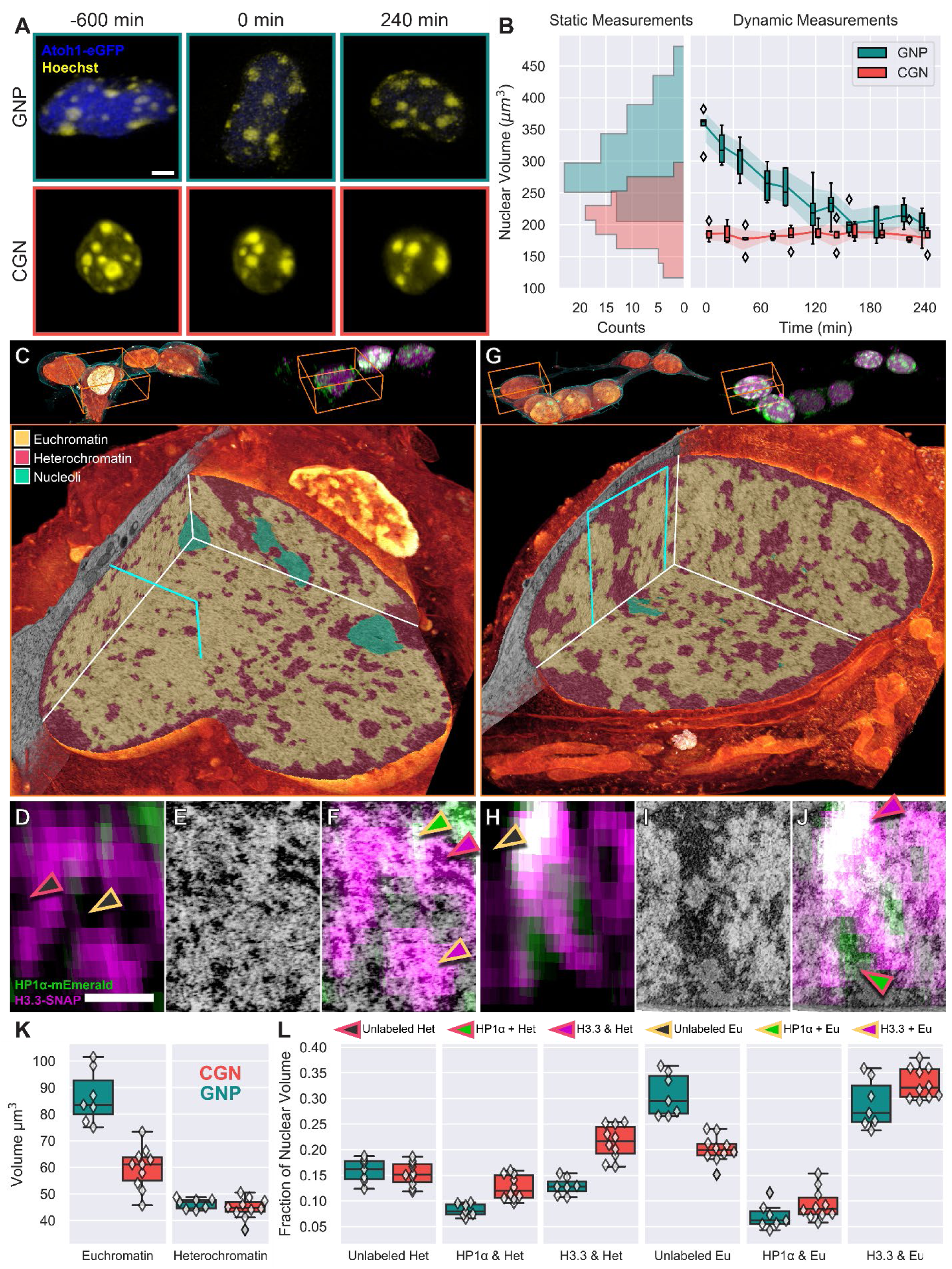
Cryo-SIM/FIB-SEM reveals nuclear rearrangements associated with cerebellar granule neuron progenitor (GNP) differentiation. **(A)** Live-cell lattice light sheet time-lapse images showing an EGFP-Atoh1 positive GNP (top row) condensing its nuclear size to that of a CGN while the size of an EGFP-Atoh1 negative CGN nucleus (bottom row) remains constant. Scale bar: 3 μm. **(B)** Quantification of GNP and CGN nuclear volume for both static (histograms at left, 85 CGNs and 71 GNPs) and time lapse imaging (box plots at right, 5 GNPs and 5 CGNs), showing that, on average, GNPs are 40% larger than CGNs and condense their nuclei to the size of CGNs in approximately 2 hours. **(C)** Top: FIB-SEM (left) and SIM (right) volume renderings of a group of GNPs. Bottom: One such GNP nucleus (orange boxes at top), with cutaway, showing color-coded chromatin territories (heterochromatin, euchromatin or nucleoli) as identified on the basis of the EM data alone. **(D-F)** HP1α (green) or H3.3 (magenta) Cryo-SIM, FIB-SEM and correlative XZ ortho slices of the plane bordered in cyan in (C). Arrowheads indicate different types of labeled chromatin domains, see legend. Scalebar, 1 μm. **(G-J)** Same as (C-F) but for a representative CGN nucleus (Movie 6). **(K)** Quantification of EM segmented and cryo-SIM defined chromatin domains and their correlation for 7 GNP and 9 CGN nuclei.

To uncover the intricate 3D transformations in nuclear architecture that accompany nuclear condensation during GNP differentiation, we then applied cryo-SIM to image a cohort of 7 GNPs and 9 CGNs collectively containing >2000 µm^3^ of the nuclear domain reference proteins mEmerald tagged heterochromatin protein 1 alpha (HP1α), a prototypical heterochromatin marker (*81*), and JF525-conjugated SNAP-Histone 3.3 (H3.3), a replacement histone subunit that is loaded on transcriptionally active nucleosomes (*82*) (Fig. 7C, G, top, fig. S18). We followed this with FIB-SEM imaging and segmentation of the resulting data (*83*) according to the classic EM definitions of compacted heterochromatin, open euchromatin and nucleoli (*2*) (Fig. 7C, G, bottom). Such segmentation revealed that while GNP and CGN nuclei possess similar total nuclear volumes of compacted heterochromatin (GNP = 47±2 µm^3^, CGN = 45±3 µm^3^), GNP nuclei have a significantly higher total nuclear volume of euchromatin (GNP = 84±8 µm^3^, CGN = 61±6 µm^3^) that accounts for a significant fraction of the size differential with CGNs (Fig. 7K).

Registering the cryo-SIM data onto the FIB-SEM results (Fig. 7D-F, H-J, Movie 6, supplementary note 11, figs. S19-S21, movie S4) allowed us to subclassify these classical EM chromatin domains based on their correlation to HP1α or H3.3 (supplementary note 12, figs. S22, S23). Such correlation revealed variations in these chromatin domains linked to neuronal differentiation that are not discernable by ultrastructure alone, including classical compacted heterochromatin domains with alternating layers HP1α and H3.3 (Figure 7J). Indeed, while FIB-SEM showed little difference in the absolute volume of compacted heterochromatin before and after differentiation, correlation with cryo-SIM revealed that CGN nuclei have ∼50% more normalized nuclear volume of HP1α loaded heterochromatin than GNPs (GNP = 8±1%, CGN = 12±2%, Fig. 7L). Moreover, surface area to volume measurements showed HP1α loaded heterochromatin became substantially more compact during nuclear condensation (fig. S23A).

Analysis of H3.3 relationships to heterochromatin and euchromatin also revealed substantial differences between GNP and CGN nuclei. While both GNPs and CGNs had similar amounts of H3.3-loaded euchromatin (GNP = 27±4%, CGN = 32±3%, Fig. 7L) indicative of transcriptionally active regions, GNPs had 50% more normalized nuclear volume of a H3.3-free form of euchromatin than did CGNs (GNP = 29±4%, CGN = 20±2%, Fig. 7L). Live cell LLSM operating in the higher resolution SIM mode revealed that these large H3.3-free voids in GNP nuclei contain mEmerald-cMAP3, a marker of H3K27me3 and H3K4me3-loaded poised chromatin (*84*) suggesting that groups of poised genes are organized in a region-specific fashion in neural progenitors (supplementary note 13, fig. S24, movie S5).

GNP differentiation into CGNs also resulted in the unexpected accumulation of H3.3 in heterochromatin nearly twice as abundant in CGNs as GNPs (GNP = 13±1%, CGN = 22±3%, Fig. 7L). Like classical HP1α-loaded heterochromatin, H3.3-heterochromatin also underwent compaction during CGN differentiation (fig. S23C). The presence of a large fraction of H3.3-loaded heterochromatin in differentiated neurons was surprising given H3.3-loaded heterochromatin species are abundant in pluripotent embryonic stem cells (ESCs) but have not been observed in most of their somatic cell derivatives (*85, 86*). Furthermore, LLS-SIM revealed that H3.3-loaded heterochromatin is likely not due to H3.3 recruitment to telomeres or centromeres as has been reported for ESCs (fig. S24B).

Finally, heterochromatin subdomains exhibited spatially distinct organization patterns depending on whether they were loaded with HP1α or H3.3 (fig. S22A, movie S7). Additional analysis based on the density of heavy metal staining in a correlated 4 nm FIB-SEM data set revealed that H3.3-heterochromatin was less densely packed then HP1α-heterochromatin in CGN nuclei, showing that molecularly defined heterochromatin subdomains are not only spatially distinct at the level of the whole nucleus but are also morphologically distinct at the ultrastructural level (Fig. S22B).

## Discussion

Much of what we know about the structural and functional organization of the cell at the nanoscale comes from a synthesis of the findings of EM and biochemistry. Although this synthesis has proved powerful, fusing the insights from these disparate methods necessarily involves developing models, and therefore possible biases, of how specific proteins are spatially distributed in relation to the EM ultrastructure that bear closer examination. Correlative cryo-SR/FIB-SEM enables such examination by combining two complementary datasets, often revealing unanticipated protein localization patterns or ultrastructural morphologies at variance with such models, while also enabling the potential discovery of new subcategories of functionally distinct subcellular structures that appear morphologically similar by either SR or EM alone. As such, it serves as a prolific generator of observations upon which more refined models can be developed in a way mutually consistent with the findings of SR, EM, live imaging, and biochemistry.

Of course, the value of cryo-SR/FIB-SEM to this enterprise depends on the extent to which it reveals the native ultrastructure of the cells it images, and the extent to which these cells are representative of the normal physiological state of their class. We designed our pipeline with these goals in mind. HPF immediately followed by cryo-SR imaging of cells in vitreous ice without any intervening chemical modification ensures that a faithful, unperturbed snapshot of the cell is captured, and allows SR and EM sample preparation protocols to be decoupled and independently optimized. Widefield cryo-fluorescence imaging to rapidly survey hundreds of cells, followed by higher resolution inspection of likely candidates by multicolor 3D cryo-SIM at a few minutes per cell ensures that only those cells of physiological morphology, or ones in a specific desired physiological state (e.g., (*87*)) are considered for time-intensive cryo-SMLM and FIB-SEM. Lastly, freeze substitution provides excellent preservation of native ultrastructural detail, while subsequent whole-cell 3D FIB-SEM gives a comprehensive picture of subcellular components across all regions of the cell, at 4 or 8 nm isotropic voxels not possible by serial section transmission EM or mechanically sectioned serial block face EM.

That being said, cryo-EM tomography of thin lamellae excavated from whole cells by cryo-FIB (*8, 9*) offers molecular resolution without any risk of ultrastructural perturbation by heavy metal staining and resin embedding. Given, however, that the lamellar volume is typically only a small fraction of the entire cellular volume, many structures of interest will be missed entirely, and those which are seen may not exhibit the same morphology as in other regions of the cell. Thus, FIB-SEM and cryo-EM tomography are complementary and developing a pipeline to do both in conjunction with cryo-SR would be a worthwhile endeavor.

Indeed, the unique ability of FIB-SEM to image whole cells and tissues at 4-8 nm isotropic voxels over volumes as large as 10^7^ µm^3^ makes it an ideal tool to map *in toto* the 3D ultrastructural relationships in living systems. However, to unlock its full potential, robust automated identification and segmentation of specific intracellular features of interest is required, ideally in relationship to neighboring structures with which such features might interact. This remains challenging to accomplish at scale, given the magnitude of the data involved (e.g., 100 GB in Fig. 7 and 19.5 TB in (*12*)), the diversity, spatial density, and conformational complexity of intracellular compartments, and the monochromatic nature of the data. Cryo SR can play an important role in the development of scalable segmentation, both in the validation of training sets for machine learning, and in confirmation of the resulting segmented outputs.

We can also envision a number of possible improvements to our pipeline. First, live cell imaging immediately prior to freezing would allow correlation of dynamics to ultrastructure, refine selection to cells of physiological behavior, and enable pharmacological, optogenetic, or other perturbations to be applied. Second, an extension of cryo-SR/FIB-SEM to more physiologically relevant specimens such as small organisms or organoids should be feasible within the 200 µm thickness limit for HPF by incorporating adaptive optics for aberration-free deep imaging. Third, the axial resolution of both cryo-SIM and cryo-SMLM could be improved ∼5-10× by designing a dual window cryostat using opposed objectives and coherent detection, such as in I^5^S (*88*) and iPALM (*89*). A next generation pipeline combining these improvements could prove an even more powerful discovery platform to link 3D subcellular dynamic processes in cells, small whole organisms, and acute tissue sections to the nanoscale spatial distribution of the proteins driving these processes, all in the context of the global intracellular ultrastructure. However, even in its current form, our cryo-SR/FIB-SEM system can address a broad swath of biological questions and is available to outside users wanting to do so (*90*).

## Materials and methods

### Preparation of vitrified samples

Specimens were cultured on 3 mm diameter, 50 µm thick sapphire disks (Nanjing Co-Energy Optical Crystal Co., Ltd, custom order, supplementary note 2A) before cryofixation with a Wohlwend Compact 2 high pressure freezer. Sample-specific protocols and plasmid maps can be found in supplementary note 14.

### Cryogenic light microscopy

To optically image vitrified samples at diffraction-limited resolution and beyond, they must be maintained below 125K to avoid de-vitrification (*91*), and present a clean, optically flat surface for aberration-free imaging. To achieve these ends, we built our microscope around a modified commercial liquid helium flow cryostat (Janis Research Company, ST-500, supplementary note 3, fig. S6) and imaged cells plated on sapphire coverslips (supplementary note 2A) through the opposite surface, after clearing this surface of residual ice in a custom cryo-preparation chamber (supplementary note 2C, fig. S5 and movie S2). We transferred samples from cold storage to the imaging cryostat using custom tools and procedures adapted to a commercial cryogenic vacuum transfer system (Quorum Technologies, PP3010T, supplementary note 3B, fig. S6). SIM images were processed as previously described (*92*) and SMLM processing is described in supplementary note 15.

### EM sample preparation

Following optical imaging, samples were transferred back to cryo-storage before being freeze-substituted, resin embedded, and re-embedded (supplementary notes 6B, 6C). Desired regions of interest (ROIs) were identified in the plasticized specimens (fig. S12) using an XRadia 510 Versa micro X-Ray system (Carl Zeiss X-ray Microscopy, Inc.) and then trimmed to expose small (∼100 × 100 × 60 µm) stubs (supplementary note 6D).

### FIB-SEM imaging

Standard (8 × 8 × 8 nm^3^ isotropic voxel) FIB-SEM datasets were generated using a customized Zeiss Merlin crossbeam system previously described (*12*) and further modified as specified in supplementary note 16. The SEM image stacks were acquired at 500 kHz/pixel with an 8 nm x-y pixel using a 2 nA electron beam at 1.2 kV landing energy for imaging and a 15 nA gallium ion beam at 30 kV for FIB milling. 4 × 4 × 4 nm^3^ voxel datasets were generated using a similarly customized Zeiss GeminiSEM 500-Capella Crossbeam system. The block face was imaged by a 250 pA electron beam with 0.9 kV landing energy at 200 kHz. The final image stacks were registered using a SIFT (*93*) based algorithm.

## Acknowledgments

We thank L. Lavis, W. Legant, D. Li, L. Shao, C. Ott, N. Alivodej, Niraj Trivedi, Danielle Wong and K. Hayworth. We also thank the Shared Resource teams at Janelia for their skill and dedication in specimen handling and preparation and the Janelia Experimental Technologies team for their manufacturing expertise, in particular Bill Biddle and Bruce Bowers. We gratefully acknowledge the support of the Janelia Visitor Program. mEmerald-N1 and mEmerald-ER3 were a gifts from Michael Davidson (Addgene plasmid #53976; http://n2t.net/addgene:53976; RRID:Addgene_53976 and Addgene plasmid #54082; http://n2t.net/addgene:54082; RRID:Addgene_54082). D.P.H., G.S., C.S.X., M.F., D.E.M., H.A.P, N.I., J.A.B., S.P., D.P., K.S., C-L.C., J.L-S., E.B., and H.F.H were funded by the Howard Hughes Medical Institute (HHMI). L.W., W.P., and T.K. were funded by grants from Biogen, NIH R01 GM075252 and a MIRA NIH award GM130386 (to T.K.). K.R.C., D.R.S., A.S. and D.J.S. were funded by the American Lebanese Syrian Associated Charities (ALSAC) and by grants 1R01NS066936 and R01NS104029-02 from the National Institute of Neurological Disorders (NINDS). LLSM images were acquired at St. Jude Children’s Research Hospital in the Department of Developmental Neurobiology Neuroimaging Laboratory.

## Author contributions

D.P.H., G.S., E.B., and H.F.H. supervised the project and wrote the manuscript with input from all coauthors. D.P.H. and G.S. designed and built the cryogenic optical microscope with input from E.B. and H.F.H. and performed all characterization experiments. D.P.H. and G.S. conducted all optical measurements. C.S.X optimized and adapted the FIB-SEM instruments. C.S.X., S.P., and D.P. conducted all EM experiments. D.E.M. created the instrument control software for the cryo-microscope. J.A.B. customized BigWarp. M.F. and K.S. prepared samples for Fig. 1-3 and the associated supplemental material. C.-L.C and L.W. prepared samples for Fig. 4 and 5, respectively. D.J.S. prepared all granule neuron samples and led related experiments. D.R.S., K.R.C. and D.J.S. performed LLSM and LLSM-SIM experiments. M.F., G.S., and D.P.H. froze specimens. G.S., H.A.P., and N.I. prepared all specimens for EM. D.P.H. and G.S. processed and analyzed all data except for that presented in Fig. 7 and the associated supplemental material which was done by K.R.C., D.R.S., A.S., and D.J.S. D.P.H. and G.S. produced all figures and movies except Fig. S18-S24, Movie S4, and the last part of Movie 6, which were produced by K.R.C. and D.R.S. W.P. assisted in aligning the data in Fig. 5. J.L.-S. and T.K. aided the biological interpretation of the results in Fig. 4 and 5, respectively.

## Data and materials availability

All data needed to evaluate the conclusions of this paper are present here or the supplementary materials. The total size of the raw data used here exceeds tens of terabytes. All data used in this paper is freely available upon reasonable request to those who provide a mechanism for facile data transfer.

**Supplementary Materials:**

Supplementary Notes 1 to 16

Figs. S1 to S49

Tables S1 to S2

Captions for Movies S1 to S7

Movies S1 to S7

---

**Movie 1.**
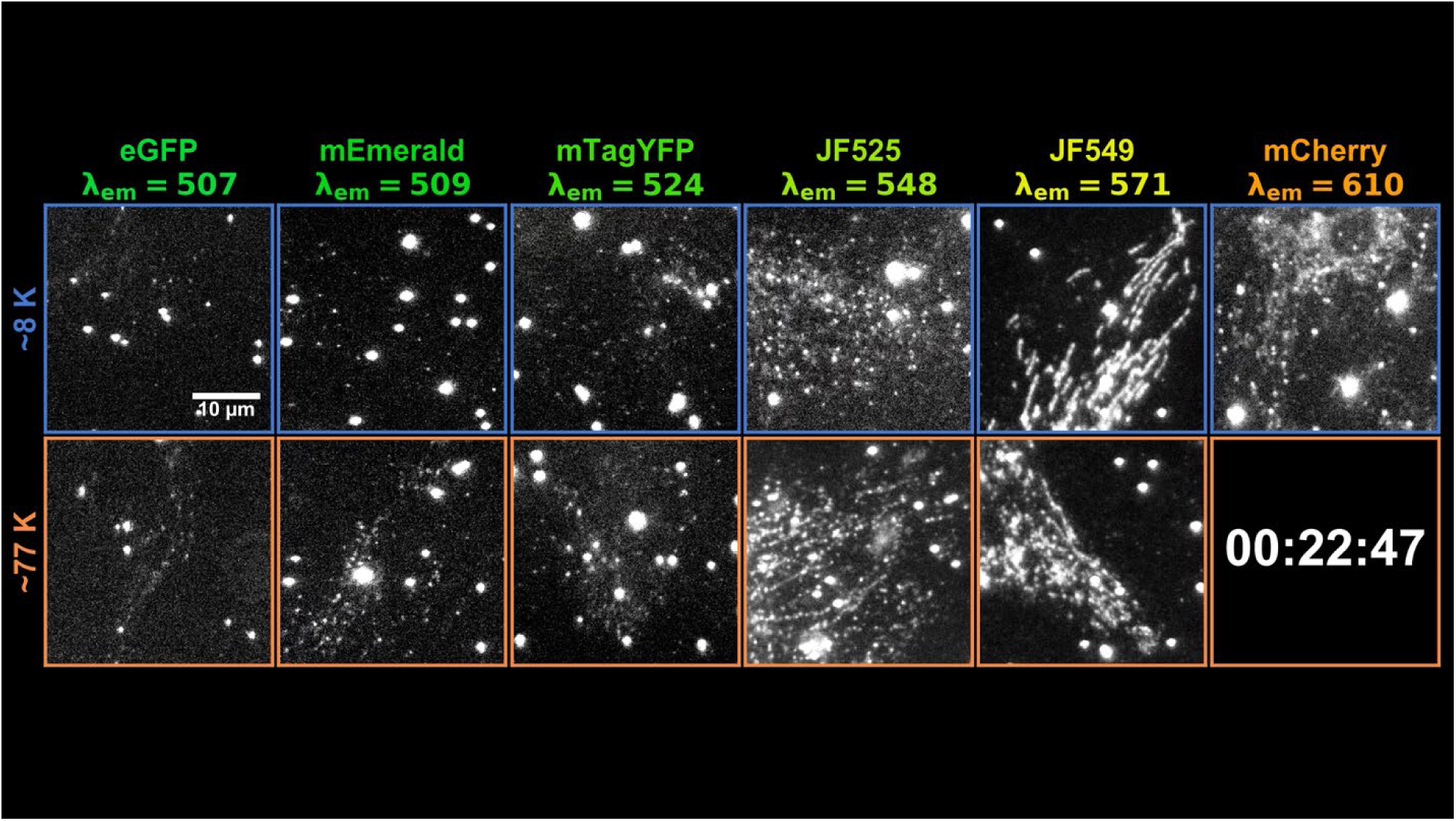
Raw single molecule frames over time since initial illumination illustrating dark state conversion efficiency and background as functions of temperature and emission wavelength. High pressure frozen U2OS cells expressing fluorescent protein or dye labeled TOMM20 to mark the outer mitochondrial membrane, as seen at ten different intervals over 3.5 hours of illumination. Bright continuous emitters are fluorescent bead fiducial markers. As seen, all six emitters asymptote to better single molecule contrast at ∼8K than 77K, yielding more accurate single molecule localization (Fig. 1A-C, fig. S9).

**Movie 2.**
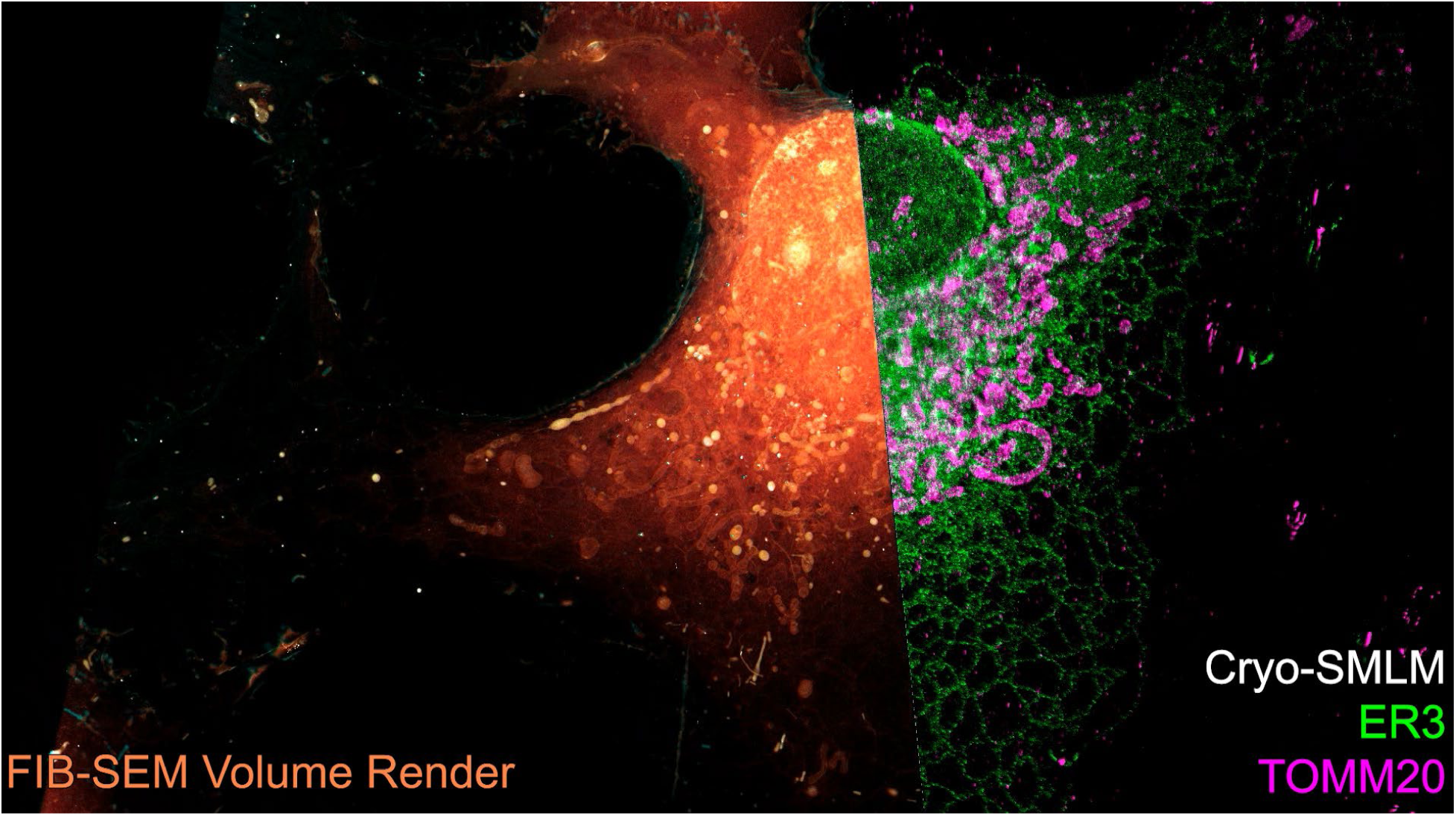
Correlative cryogenic 3D super-resolution and block face electron microscopy of whole vitreously frozen cells. Two COS-7 cells expressing markers for the endoplasmic reticulum (mEmerald-ER3, green) and mitochondria (Halo/JF535-TOMM20, magenta), shown in relation to orthoslice (grayscale) or volume rendered (plasma membrane, cyan; intracellular volume, orange) FIB-SEM data. An ER3-positive intranuclear vesicle and several cytosolic TOMM20-positive vesicles identified by correlation are also highlighted (Fig. 3, fig. S15, S16).

**Movie 3.**
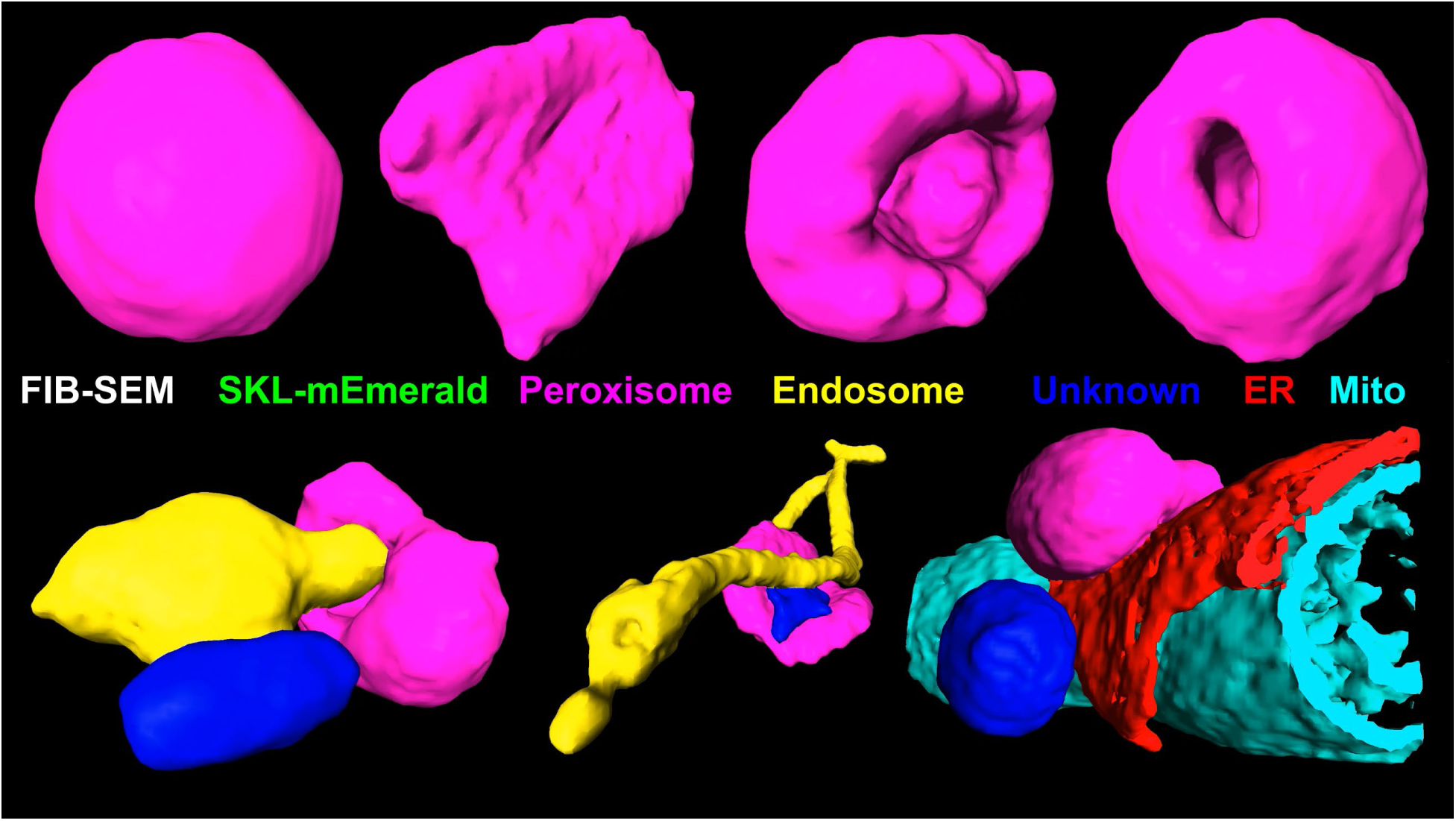
Structural diversity of peroxisomes and their inter-organelle contacts. Peroxisomes from a HeLa cell expressing mEmerald-SKL. Part 1 shows orthoslices of the FIB-SEM and cryo-SMLM data followed by segmentations of SKL labeled peroxisomes. Part 2 shows the same but with segmentations of other organelles in contact with SKL labeled peroxisomes (Fig. 4).

**Movie 4.**
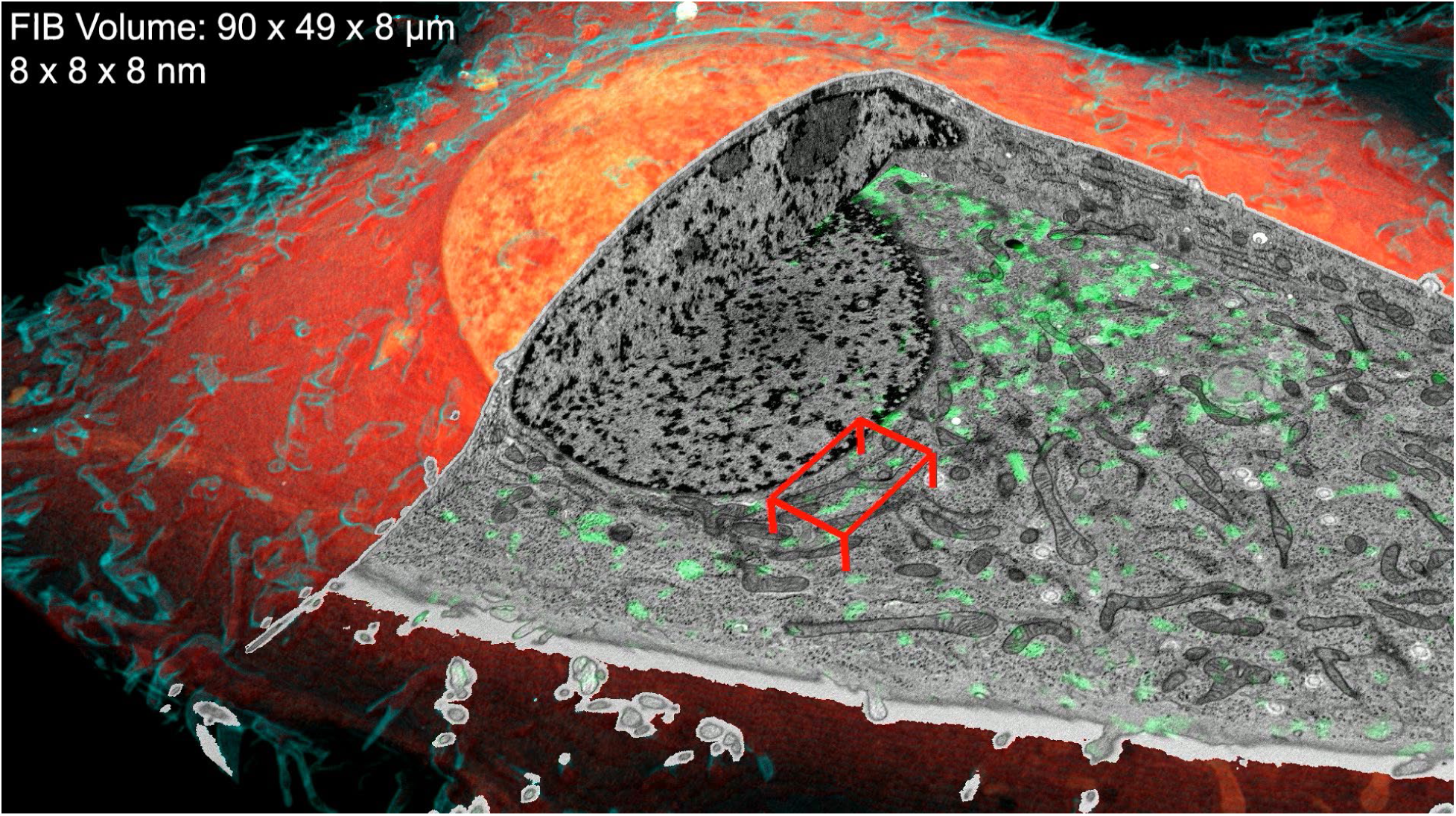
Cryo-SIM/FIB-SEM reveals the morphological heterogeneity of the endolysosomal system. A correlative data set of a SUM-159 cell after endosomal uptake of Alexa Fluor-conjugated transferrin. Part 1: 3D cryo-SIM data (green), correlative orthoslices, and correlative volume render (plasma membrane, cyan; cellular interior, orange). Part 2: ∼13 µm^3^ sub-volume showing segmentations of all transferrin containing compartments. Part 3: same, but for a different ∼20 µm^3^ sub-volume (Fig. 5).

**Movie 5.**
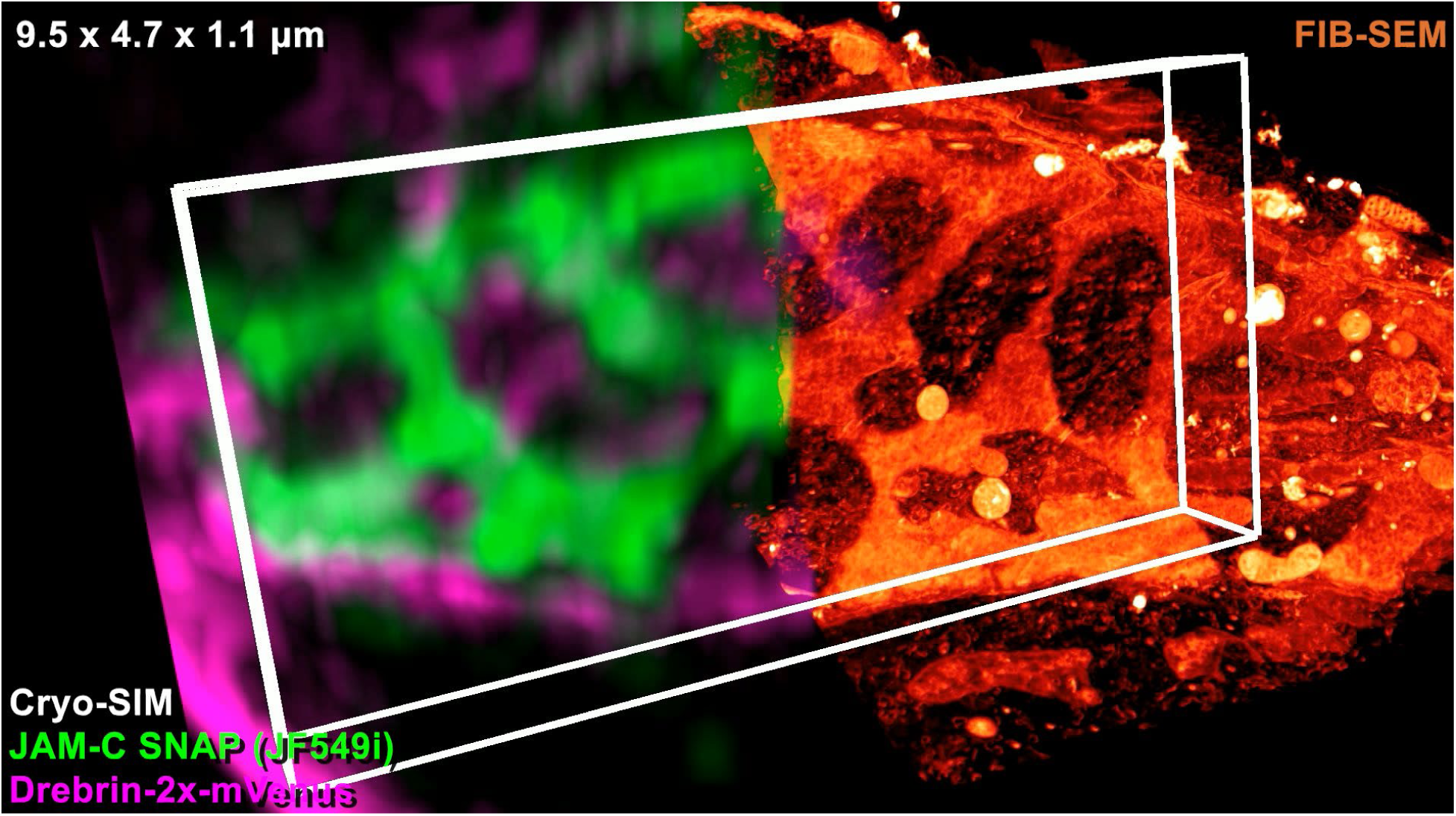
Correlative cryo-SIM/FIB-SEM reveals a web-like adhesion pattern between adjacent cerebellar granule neurons. Part 1: cryo-SIM and FIB-SEM volume renderings of a field of CGNs expressing adhesion proteins JAM-C (green) and drebrin (magenta). Part 2: correlation between electron density at the PM, JAM-C cryo-SIM signal, and PM curvature at the interface between two CGNs (Fig. 6).

**Movie 6.**
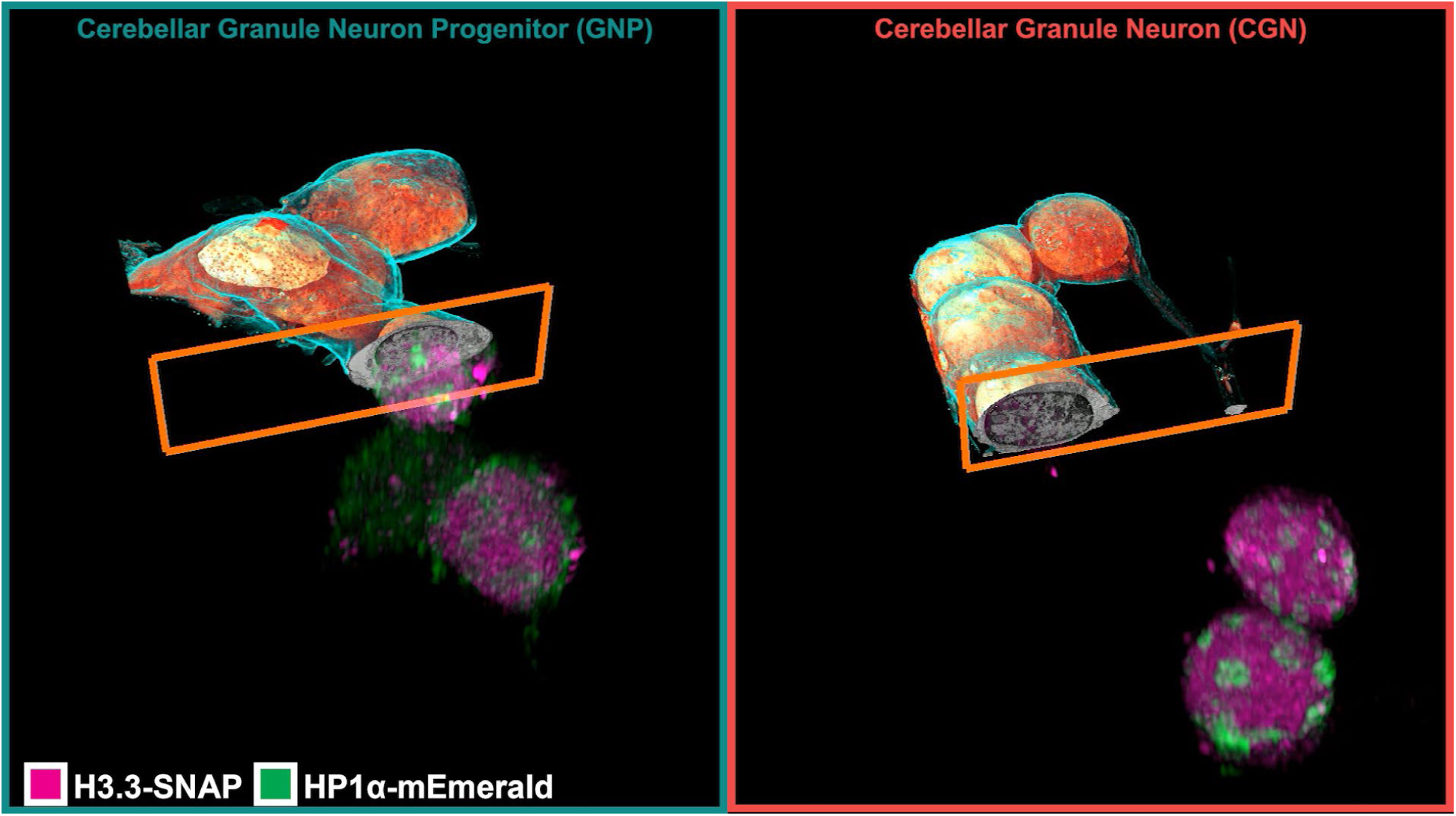
Chromatin compaction during differentiation and identification of novel chromatin subdomains. Correlative data sets of granule progenitor (GNP, left) and cerebellar granule neurons (CGN, right). Part 1: overall correlation between the FIB-SEM (plasma membrane, cyan; cellular interior orange) and cryo-SIM of the nuclear domain reference proteins HP1α (green) and H3.3 (magenta). Part 2: cutaway views of EM-defined chromatin domains for a GNP nucleus (left) and a CGN nucleus (right). Part 3: orthoslices through the CLEM volumes indicating subdomains defined by overlap between EM-defined nuclear domains and nuclear domain reference proteins. Part 4: 3D surface renderings of CLEM defined nuclear chromatin subdomains for the GNP and CGN nuclei (Fig. 7).

