## Supplementary Text for "Correlative three-dimensional super-resolution and block face electron microscopy of whole vitreously frozen cells"

**This PDF file includes:**

Supplementary Notes 1 to 16
Figs. S1 to S49
Tables S1 to S2
Captions for Movies S1 to S7

**Other Supplementary Materials for this manuscript include the following:**

Movies S1 to S7

### Table of Contents


Supplementary Notes ... 4
1. Previous Cryo-SR-CLEM ... 4
2. Frozen Sample Preparation ... 5
a) Coverslip preparation ... 5
b) Freezing ... 6
c) Coverslip preparation for cryogenic optical imaging ... 7
3. Cryostat, Microscopy and Imaging ... 10
a) Cryo-sample holder ... 10
b) Cryostat ... 11
c) Microscope ... 12
d) Objective movement ... 13
e) SIM beam alignment ... 15
f) Chromatic focal shift ... 15
g) Preparation for imaging ... 16
h) SIM ... 17
i) Localization Microscopy ... 19
4. Photophysical Analysis of Single Molecules at Cryogenic Temperatures ... 20
a) Identifying Single Molecules in the Data ... 20
b) Dynamic Contrast Ratio ... 22
c) Static Contrast Ratio ... 22
5. Grouping ... 23
a) Identifying Groups ... 23
b) Data Driven Determination of Grouping Parameters ... 23
c) Aggregating Groups ... 27
6. Preparation for Electron Microscopy ... 29
a) Sample unloading ... 29
b) Freeze-substitution and resin embedding ... 29
c) Re-Embedding ... 30
d) X-Ray correlation and trimming ... 31
7. Correlation and registration ... 31
8. Correlative driven segmentations of peroxisomes ... 33
9. Neuronal Adhesion Curvedness Calculation ... 35
10. LLSM Imaging of GNP Nuclear Condensation ... 35
11. Cryo-SIM/FIB-SEM registration ... 37
12. Cryo-SIM/FIB-SEM segmentation ... 38
13. Biological Insights from Cryo-SR/FIB-SEM of GNP/CGN Nuclei ... 38
14. Cell Culture ... 41
a) Fig. 1, 2, 3, S2, S3, S10, and S11 ... 41
b) Fig. 4 ... 46
c) Fig. 5 ... 47
d) Fig. 6 ... 48
e) Fig. 7 ... 51
15. SMLM Data Processing ... 55
a) Constrained 2D Gaussian Fitting to extract 3D coordinates of the fluorescent events. 55
b) Drift correction ... 56

c) Slab alignment ... 58
d) Multiple Wavelength Channel Registration ... 59
e) Filtering ... 60
16. FIB-SEM imaging ... 61
Fig. S1. ... 63
Fig. S2. ... 64
Fig. S3. ... 66
Fig. S4. ... 67
Fig. S5. ... 68
Fig. S6. ... 69
Fig. S7. ... 70
Fig. S8. ... 71
Fig. S9. ... 74
Fig. S10. ... 75
Fig. S11. ... 77
Fig. S12. ... 79
Fig. S13. ... 80
Fig. S14. ... 82
Fig. S15. ... 83
Fig. S16. ... 86
Fig. S17. ... 87
Fig. S18. ... 88
Fig. S19. ... 89
Fig. S20. ... 90
Fig. S21. ... 91
Fig. S22. ... 92
Fig. S23. ... 94
Fig. S24. ... 95
Fig. S25. ... 97
Fig. S26. ... 98
Fig. S27. ... 99
Fig. S28. ... 100
Fig. S29. ... 102
Fig. S30. ... 104
Fig. S31. ... 105
Fig. S32A. ... 106
Fig. S32B. ... 108
Fig. S32C. ... 109
Fig. S33. ... 111
Fig. S34. ... 112
Fig. S35. ... 114
Fig. S36. ... 116
Fig. S37. ... 117
Fig. S38. ... 118
Fig. S39. ... 119
Fig. S40. ... 121

Fig. S41. 122
Fig. S42. 123
Fig. S43. 124
Fig. S44. 125
Fig. S45. 126
Fig. S46. 127
Fig. S47. 128
Fig. S48. 129
Fig. S49. 130
Table S1. 131
Table S2. 133
Movie S1. 135
Movie S2. 136
Movie S3. 137
Movie S4. 138
Movie S5. 139
Movie S6. 140
Movie S7. 141
References 142

**Supplementary Notes**

#### *1. Previous Cryo-SR-CLEM*

In general SR-CLEM approaches can be split in few groups:

1. Tokuyasu cryo-sectioning (refs (*94–96*)). Those were early attempts, with multiple problems:
    a. Poor quality ultrastructure preservation.
    b. Not applicable to whole cells/thick sections
2. Acrylic resin embedding for in-resin CLEM (refs (*40–42*, *97*, *98*)). In attempt to allow for higher quality EM, resin embedding was introduced. Most fluorophores do not blink without hydration, so hydrophobic resins, such as Durcupan or Epon, cannot be used. Hydrophilic, usually acrylic resins have been used instead. Multiple problems remain:
    a. If fixation is not strong enough, ultrastructure preservation is poor.
    b. With stronger fixation, even with "fixation-resistant fluorophores", the fluorophore survival is low. And even the "stronger" fixation is still not optimal strength fixation, so the ultrastructure quality and staining contrast are compromised.
    c. Acrylic resins are not compatible with large volume 3D EM techniques, such as FIB-SEM
3. Platinum-replica of thin membranous or membrane-bound structures (refs (*99*, *100*)), metal/carbon-coated membranes (refs (*101*, *102*)). These techniques generally provide

good quality data but are only available for studies of membrane-bound structures or cell morphology.

4. Cryo-ET/Cryo-PALM (refs (*7*, *29*, *30*))
    a. CryoEM tomography may be done on small ROI's of thin areas/sections - not on the whole cells.
    b. Plunge-freezing is good for thin areas, but of debatable quality for whole cells (*21*, *103*, *104*).
    c. Imaging on grids presents a problem with heat dissipation, the imaging power has to be limited to avoid de-vitrification (*29*). With low power one can get shimmering, but not blinking (*33*).

#### *2. Frozen Sample Preparation*

##### a) Coverslip preparation

HPF demands high thermal conduction and mechanical durability from its substrates and sapphire has both while also being optically transparent. Fluorescence microscopy additionally requires that the substrate be 1.) optically clear (non-absorbing), 2.) optically flat, 3.) free of distortions and 4.) without any intrinsic fluorescence. Sapphire has excellent optical properties meaning that requirements 1-3 are satisfied. However, sapphire can have brightly fluorescent inclusions, most commonly chromium. Chromium in sapphire is also known as ruby which is a lasing medium and therefore brightly fluorescent. It has two well defined emission peaks at 692 nm (R1) and 693 (R2) (cf. Fig. S26). While these emission peaks can be removed with the appropriate optical filters the vibronic bands surrounding them extend from 665 nm at the short end to 730 nm at the long end (cf. Fig. S26) and are excited by 561 nm light interfering strongly with standard fluorescent imaging. Most common commercial sources of sapphire coverslips for HPF have an un-acceptably high level of impurity. Of all the manufacturers we tested Nanjing Co-Energy Optical Crystal Co., Ltd (COE) offered the lowest fluorescence impurities.

Sapphire coverslips (3 mm diameter, 0.05 mm thickness, Nanjing Co-Energy Optical Crystal Co., Ltd) were cleaned in basic piranha solution (5:1:1 solution of water : ammonia hydroxide : hydrogen peroxide (30%)) for a minimum of 1 hour. Before proceeding all coverslips are inspected for acceptable parallelism; too much of a wedge will degrade image quality, so much so that in some cases SIM reconstruction is impossible. To inspect for parallelism a collimated 2 mm diameter laser beam is directed at the coverslip surface and the reflection is observed on a sheet of paper ~4 m away. If there is more than 1 cm separation between the front and back surface reflections, or if either spot is larger than ~5 mm the coverslip is discarded. Coverslips that pass inspection have a thin (~100 nm) coat of gold was sputtered (sputter coater Desk II, Denton Vacuum) on to a 0.5 mm wide border region to differentiate the surfaces such that the sides with cells plated on them (opposite of the gold coated sides) could be identified at all steps in the process (cf. Fig. S1, top). To prepare bead coated coverslips the cleaned, gold coated sapphire coverslips were transferred to a vacuum chuck and incubated in a 0.2% w/v poly-L-lysine HCl (P2658, Sigma) solution for 15 minutes, rinsed for ~30 seconds, incubated in a solution with 0.8 pM each of green (100 nm, ThermoFischer F8803), orange (100 nm, ThermoFischer F8800), red (100 nm, ThermoFischer F8801) and deep red (40 or 200 nm, ThermoFischer F8789 or F8807) fluorescent beads for another 15 minutes and then rinsed a final time for ~30 seconds.

**b) Freezing**

Immediately prior to freezing, cells are inspected, live, with an inverted wide field microscope (Nikon Instruments, Ti-Eclipse) equipped with a $CO_2$ incubation chamber and a 60X water immersion NA 1.2 objective lens (CFI Plan Apo VC 60XWI). The correction collar on the objective is not designed to image through sapphire and there is still a significant amount of spherical aberration in the images, but they are sufficient to inspect cell morphology and viability.

After quality assurance the cells are transferred to a water jacketed $CO_2$ incubator (ThermoFisher, Midi 40) kept at 37° C, 5% $CO_2$, and 100% humidity while awaiting freezing. Each coverslip is removed from the incubator immediately prior to the freezing procedure. The freezing procedure consists of eight steps:

1. Coat the sides of the aluminum platelets that will contact the sapphire with hexadecene
2. Blot the hexadecene from the platelets with filter paper (Whatman® qualitative filter paper, Grade 1)
3. Prepare the HPF holder by placing the flat platelet (Technotrade International, Alu Platelet grinded 479 (0.3)) into it
4. Remove a coverslip from the incubator

5. Replace the media on the coverslip with a dextran media mixture by dipping the coverslip into three independent 20 µL reservoirs of the dextran mixture
6. Place the coverslip with the replaced media onto the 25 µm well of the other aluminum platelet (Technotrade International, Alu Platelet 389 (0.025/0.275))
7. Blot this "sandwich" with filter paper to remove the excess media and then place it into the holder such that the sapphire is flush with the flat platelet
8. Freezing is completed by following the manufacturer's instructions and the frozen sandwich is stored under liquid nitrogen in a custom-made holder manufactured out of brass or aluminum

Once the sample is frozen it can be stored indefinitely. The dextran mixture is a 25% w/v solution of 40,000 MW dextran (Sigma, 31389-100G) which is used as a cell impermeable, low osmolality cryo-protectant to prevent ice crystal formation in the extra cellular space (*103*). For SMLM samples the dextran mixture has ~4 pM fluorescent beads added. Once frozen in place the beads offer ideal three-dimensional fiducials for calibration, drift correction, and slab alignment (cf. supplemental note 15).

**c) Coverslip preparation for cryogenic optical imaging**

The cryogenic sample preparation chamber (PC, Fig. S5) was designed to facilitate transfer of high-pressure frozen samples from liquid nitrogen storage into the sample holder (SH, cf. supplemental note 3 and Fig. S4) and back as well as for cleaning the sapphire surface in preparation for optical imaging. Two important design goals have to be met: 1) PC must enable samples to be handled while they are submerged in liquid nitrogen so that they never exceed the devitrification temperature (125 K)(*91*) and 2) PC needs to be compatible with the cryogenic transfer device (CTD, part # PP3010T Cryo Transfer Device, Quorum Technologies, Laughton, East Sussex, UK). Vacuum transfer is necessary to ensure that the samples are thermally insulated and kept free of contamination, especially water condensation.

PC consists of three components; a large vacuum chamber (Vacuum Chamber - 11" ID x 6" DP, Abbess Instruments and Systems, Inc.), a liquid nitrogen flask (NF, 8150, Pope Scientific, Inc.) and the preparation stage/mount for SH. Two modifications were made to the main vacuum chamber (cf. Fig. S5C, D): a hole and indent were cut into the acrylic lid to fit the vacuum shuttle and a vacuum compatible electronics port was added to the side.

The most important parts of the preparation stage are an aluminum work platform (WP) and a modified cold finger (MCF, green, Fig. S5A, C, D). MCF is mounted into WP and both are positioned within NF such that when NF is filled with liquid nitrogen both MCF and WP are submerged. MCF here is very similar to that used in the optical cryostat (Fig. S4C). Here, MCF is modified so that the coverslip holding fingers of SH can be actuated with the spring-loaded actuator arms (AA, magenta, Fig. S5A,C,D). Pushing down on a screw head in the direction of a vertical blue arrow translates the motion along the horizontal blue arrow as the arm pivots on the axis indicated by a white dashed line, which in turn moves the spring-loaded finger in SH; actuation of the finger allows samples to be inserted securely into, or removed from, SH (cf. Fig. S4).

WP has a 2 mm rim to prevent samples from falling into the liquid nitrogen bath, an indentation for holding a long-term cryo-storage box (CB), and a preparation area (PA) that has circular indentations and dowel pin posts to facilitate separation of the sapphire coverslips from the aluminum planchettes used during high-pressure freezing and subsequent cleaning of the sapphire surface from ice necessary for optical imaging (cf. Movie S1).

The preparation stage is attached to 10 mm thick Teflon annulus (TA) via 1” thick posts made from G10 Epoxy. WP is mounted on stainless-steel rods (dark blue, Fig. S5A,C, D) so it can rotate 360 degrees while submerged in the liquid nitrogen bath. The WP can be oriented so that the top of SH is facing up (Fig. S5C) for sample preparation, or down (Fig. S5D), for transfer into the CTD (i.e. oriented such that the bayonet on the CTD can be inserted in to the back of the SH). Both the upward and downward orientations are stabilized with spring loaded pins. Figure S5D presents SH in a mid-way position as it is being moved from MCF into CTD. TA has outer and inner diameters of 186 and 85 mm, respectively, so that it sits on the edge of NF with WP, MCF, and SH suspended in the liquid nitrogen bath. An LED mounted on the underside of TA provides oblique illumination which improves the visibility of the surface morphology of the sapphire coverslips.

To prepare the coverslips for imaging they are first removed from CB and the aluminum planchettes are removed using the small circular indentations (cf. S5A and Movie S1) on the stage. The sapphire coverslip is then placed back onto the aluminum platelet with the 25 μm well with the gold coated surface (the bottom) facing upwards. The ice on the gold coated surface is removed (i.e. scraped off) using a small rectangular blade (Fine Science Tools, 10035-05, cf. Movie S1).

Prior to scraping the blade is manually sharpened and polished using successively finer grits of sandpaper and finished with 0.02 μm grit fiber lapping film. Cleaned coverslips are loaded into SH with the bottom (gold plated) side facing up (cf. top of Fig. S1 for sample geometry). A second round of scraping is performed once all coverslips have been loaded into SH. At this point a final pre-load quality control inspection of the coverslips can be performed using a fluorescence stereoscope (Nikon, SMZ18). A final attempt to remove residual contamination is made by spraying liquid nitrogen onto the backside of the coverslips using a liquid nitrogen spray gun (Cry-Ac® B-700, Brymill Cryogenic Systems). Scraping provides an added benefit beyond cleaning, it leaves unique marks on the gold coating allowing individual coverslips to be easily identified with a light microscope aiding the EM preparation steps detailed below (cf. supplemental note 6 and Fig. S12B, C). Note that all tools, such as forceps and scalpels are pre-cooled in liquid nitrogen before being used to handle the coverslips as a precaution against devitrification.

After cleaning the preparation stage is rotated 180 degrees, the working lid (WL) is replaced by vacuum lid (VL) and CTD is docked to VL (Fig. S5D). PC is then evacuated through a vacuum port (VP) using a scroll pump (Edwards nXDS10i, Kurt Lesker). As PC pumps down, the bayonet of CTD is engaged and SH is loosened from MCF. Pumping down PC achieves two goals: 1) the liquid nitrogen is cooled to the freezing temperature of ~63 K providing an extra temperature margin for safe sample transfer, and 2) the pressure in PC is reduced to about 200 Torr of dry nitrogen. Once the liquid nitrogen begins to freeze the pump valve is closed. As soon as the nitrogen begins to thaw SH is retracted into CTD. CTD is then sealed, PC is vented and CTD is undocked from PC and docked to the airlock of the optical cryostat. The airlock is then pumped down to 100 mTorr and the sample is transferred into the cryostat, ensuring that the sample never comes into contact with the atmosphere. Once the sample transfer is completed, the cryostat is pumped down to about $10^{-6}$ Torr. The whole transfer procedure takes less than 30 seconds on average. In our experience, the cryo-sample holder's temperature, as monitored by the on-board temperature sensor, does not exceed 70 K during the transfer process.

#### 3. *Cryostat, Microscopy and Imaging*

##### a) Cryo-sample holder

The cryogenic-sample holder (SH, Fig. S1, inset photograph and Fig. S4A,B) is at the heart of the instrument and has dictated many of the other design decisions that were made. Crucially, it must allow for the loading and unloading of *vitrified* samples, i.e. samples that are at or below 77 K, and it must be compatible with liquid helium cooling which requires the main components be fabricated from highly conductive materials and for all components to be gold coated (to reflect IR blackbody radiation).

The sample holder body (SHB) is machined from Oxygen Free High-Conductivity (OFHC) copper (Super-Conductive 101 Copper with 99.99% copper content, McMaster-Carr) and electro-plated with ~10 μm of gold (Jewel Master Pro kit from Gold Plating Services, Layton, UT). The coverslips are held in dove-tailed pockets by stainless steel fingers, which are spring-loaded by compression springs (SP, part # 70028s, Century Springs Corp. cf. Fig. S4A). One of the coverslips has a Cernox resistance temperature detector (RTD, part # CX-1030-BC-HT-1.4L, Lake Shore Cryotronics, Inc. cf. Fig. S4A) permanently attached to it using thermally conductive epoxy (Stycast 2850, Ellsworth Adhesives). Two contact wires from RTD are pressure connected to SHB and contact ring (CR), respectively (cf. Fig. S4A). The contact ring is electrically insulated from the rest of the sample holder by two Kapton spacers. A bayonet receptacle is mounted at the back of SH and SH is held together by two M2.5 screws.

Once coverslips are mounted into the cryogenic sample holder and cleaned of ice (cf. Movie S2 and supplemental note 2.c)), SH is transferred to the cryostat through an airlock and screwed into the modified cold finger (MCF, cf. Fig. S4C). MCF is similar to the part supplied with the ST-500 Cryostat (Janis Research Company LLC), but is modified to be hollow and have a slightly larger diameter than the original part in order to accommodate SH. A spring-loaded contact pin (CP, part # 0911-0-15-20-86-14-11-0, MILL-MAX MFG. CORP, cf. Fig. S4C) is mounted into the side of MCF by means of an insulating ferrule made from Vespel (part # 87405K45, McMaster-Carr). The contact wires from the contact pin and MCF are routed to a temperature controller (Lakeshore 335, Lake Shore Cryotronics, Inc.), allowing for accurate temperature monitoring of the sample coverslips loaded in SH. Another temperature detector (part # DT-670B-CU-HT, Lake

Shore Cryotronics, Inc.) and a heater element mounted onto MCF in an identical manner to that of the original cold finger inside the ST-500 cryostat.

**b) Cryostat**

The instrument is designed around a commercially available cryostat (CS, Janis Research Company, ST-500, Fig. S1, lower left and Fig. S6). CS was modified in a few ways. Externally, 6 threaded holes (¼-20) were added at even spacing around the rim, with special precaution to not damage the vacuum seal. Internally, the cold finger was modified as described above. A 25 cm access port was added to the rear radiation shield (RS, cf. Fig. S6B) with a small swing door made from 1.6 mm thick gold coated oxygen free copper. The door is held in one of two stable positions (closed or opened) by rare earth magnets embedded in the door and the radiation shield. The door also has a small protrusion (or handle) that can be used to close or open it using the transfer stick. A gold-plated braided copper wire thermally grounds the door to the main radiation shield. Finally, the original optical window has been replaced with 0.75mm thick AR coated Fused Silica custom window (WND, cf. Fig. S6B) made by Mark Optics Inc.The cryostat is bolted, using the six ¼-20 holes, to a custom designed 25 lb solid 316 stainless-steel mount. The mount serves four purposes: 1) to minimize vibrations and effectively anchor the cryostat to the main optical table, 2) orient the cryostat so that the optical axis is parallel to the optical table's surface, 3) to serve as a steady base to build a cage around the cryogen transfer line, the source of most vibrations in the system and 4) to provide a stable platform to which the airlock can be mounted.

The cryostat is continuously under vacuum provided by a turbo pump (P1, HiCube 80 Eco, Pfeiffer Vacuum Inc.). Vacuum is further enhanced by the cryo-pumping action of the MCF. Vacuum isolation is critical to keep the chamber clean and the cold finger thermally isolated. To avoid vibrations from the turbo pump the vacuum line is run through sequential bellows and sections of pipe that are embedded in concrete blocks inside of Styrofoam containers. The final section of vacuum line is bolted to the optical table before it is connected to a valve (V1, part # 310072, MDC Vacuum) attached to the cryostat.

An airlock is bolted onto the back of the stainless-steel rear plate (RP, cf. Fig. S6B) that replaces the original back plate of the cryostat. The airlock consists of a pneumatically controlled gate valve (GV, cf. Fig. S6, part #303001-117, MDC Vacuum), a pneumatically controlled vacuum valve (V2, part #311373, MDC Vacuum) and a mount for the cryogenic transfer device (CTD,

Quorum Technologies, PP3010T). V2 is connected to an oil pump (P2, Varian DS402, Agilent Technologies) that can rapidly evacuate the airlock. A custom electronic switch box operates both pneumatic valves.

**c) Microscope**

Our microscope design is similar to that of the structured illumination microscope reported in ref. (*105*) (cf. Fig. S1, lower right corner). The excitation path is illustrated in light green. The beam from a laser combiner consisting of 488 nm (4 W, Coherent, Genesis CX STM 488nm 4W), 532 nm (10 W, Spectra-Physics, Millennia Xs), 560 nm (5 W, MPB Communications, 2RU-VFL-P-5000-560-B1R) and 642 nm (2 W, MPB Communications, 2RU-VFL-P-2000-642-B1R) is passed through an acousto-optic tunable filter (AOTF, AA Quanta Tech, Optoelectronic AOTF AOTFnC-400.650-TN). Each laser's beam waist is individually adjusted to a $1/e^2$ diameter of 2.5 mm before reaching the AOTF. The AOTF is used to select excitation wavelengths and intensities and is synchronized with the spatial light modulator (SLM) and camera. The system is also equipped with a 405 nm laser (300 mW, Oxxius, LBX-405-300-CIR-PP), which is directly modulated and combined with the other beams after the AOTF using a dichroic filter (Semrock, Di03-R405-t1-25x36). The excitation beam is expanded by 5X and projected onto a binary 2048 x 1536 pixel SLM (Forth Dimension Displays, QXGA-3DM) operated in phase mode using a polarizing beam splitter (PBS, Newport, 10FC16PB.3) and a half-waveplate (HWP1, Bolder Vision Optik, AHWP3). Patterns for structured illumination microscopy (SIM) are displayed on the SLM and the resulting patterned excitation light is Fourier transformed and focused using a 250 mm fl lens (L1, ThorLabs, C254-250-A) onto a specially designed mask (SM, Applied Image) which blocks the unwanted diffraction orders arising from the binary nature of the SLM. In between the SLM and SM are two motorized 0.5" optic rotators (Finger Lakes Instrumentation, High Speed Rotator) the first holding a quarter waveplate (QWP1, Bolder Vision Optik, AQWP3) and the second a half waveplate (HWP2, Bolder Vision Optik, AHWP3). The HWP2 and QWP1 are used to adjust the polarization of the excitation pattern to ensure maximum contrast in the SIM pattern at the sample, cf. supplemental note 3.g). The SIM beamlets at SM are then imaged onto the back pupil of the objective (OL, Nikon, CFI L Plan EPI CRB 100X, NA=0.85) by a 300 mm fl lens (L2, Thorlabs, AC508-300-A) and the 200 mm fl tube lens (TL, Nikon, MXA22018). The total demagnification of the SLM at the focal plane is 83.3 thus the SLM pixel size (8.2 μm) at the sample is 98.4 nm and the resulting illumination pattern covers ~150 μm.

The same optics used to relay the SIM pattern from SM to the back pupil are used to relay the emission light from the back pupil to SM (cf. Fig. S1, light yellow path). Here, SM serves its second purpose: to separate the emission from the excitation light in place of a dichroic mirror. To fulfill this role the mask is manufactured out of optically flat fused silica (0.5” diameter 0.1” thick, λ/10 flatness, 20-10 scratch dig) and the patterned side is coated with protected aluminum forming a mirrored surface. A photograph of SM is shown as an inset in Fig. S1. Hole placement means that at most 9% of information is lost when the mask is used for imaging; even so, the loss is either DC light (i.e. background) or at the very edge of the OTF support of the objective. The main advantage of this strategy is that it allows the use of any combination of excitation and emission wavelengths. The emission light reflected off SM is passed through a filter wheel (FW, Finger Lakes Instrumentation, HS-625) and imaged onto an sCMOS camera (Hamamatsu, C11440-22CUPLUS) with a 150 mm fl lens (L3, ThorLabs, C254-150-A). The total magnification of the sample onto the camera is 50X meaning that the camera’s pixels (6.5 um) are demagnified to 130 nm at the OL focal plane. A 500 mm fl cylindrical lens (CL, ThorLabs, LJ1144RM-A) can be placed in between L3 and the detection camera to introduce astigmatism when performing 3D-localization microscopy.

**d) Objective movement**

In contrast to conventional microscopes this instrument is designed to move the objective, *not* the sample. Moving the sample would either entail moving the entire cryostat, leading to vibration and drift or require the installation of motorized stages inside the cryostat leading to significant engineering effort. Instead, the OL is moved in the lateral plane, perpendicular to the optical path, by two linear stepper motor stages (Physik Instrumente, Miniature High-Resolution Translation Stage, M-112.12S) and axially, along the optical path, by a closed loop piezo stage (Physik Instrumente, LPS65 1/2" PM LS-072, 586092120). Two 2” protected silver mirrors (Newport, 20Z40ER.2) mounted in motorized gimbal mounts (MM1 and MM2, Newport, U200-G mounts with TRA12CC motorized actuators) compensate for OL’s movement by redirecting the optical path to the new back pupil position while making sure that the optical path remains colinear with OL’s optical axis. A benefit of this mechanism is that it minimizes the change in path length between OL and TL. For SIM there is no true infinity space; when the emission is collimated the excitation is not and *vice versa* which means the distances between OL and TL cannot be changed much. Another way to think about it is that when the distance between OL and TL is optimum the excitation light is focused into the back pupil of OL (i.e. for a given 3D-SIM pattern there are three diffraction limited spots, one in the middle and two at the very edge of the back pupil). If the distance changes much in either direction the spots in the back pupil will become defocused, i.e. larger, and because two are at the very edge of the back pupil they will be clipped. Clipping will result in a reduced field of view and less contrast in the illumination pattern.

To derive the equation relating the horizontal objective position, $x$, to the mirrors' (MM1 and MM1, cf. Fig. S1 and Fig. S27) tilt angle, $\alpha$, we note that there are two expressions for $w$ (cf. Fig. S27):

$$\begin{aligned} w &= (x - x_0)\tan(\alpha) \\ w &= \big(L_0 + (x - x_0)\big)\tan(2\alpha - 2\alpha_0) \end{aligned} \tag{1}$$

Because the choice of $x_0$ and $\alpha_0$ is arbitrary we can choose them to be 0 and $\pi/4$, respectively. Combining the two equations leads to

$$x\tan\alpha = (L_0 + x)\frac{\tan^2\alpha - 1}{2\tan\alpha} \tag{2}$$

Solving eq. (2) leads to the desired result

$$\alpha = \tan^{-1}\left(\sqrt{\frac{L_0 + x}{L_0 - x}}\right) \tag{3}$$

For practical purposes the following modified equation is used

$$\alpha = \tan^{-1}\left(\sqrt{\frac{L_0 + (x - x_0)}{L_0 - (x - x_0)}}\right) + \alpha_0 \tag{4}$$

A similar method can be used to determine the relationship between the vertical objective position, $y$, and the tip angle, $\beta$, by noting that there are two equations for $h$:

$$\begin{aligned} h &= y\tan\beta \\ L_0 - h &= y\tan 2\beta \end{aligned} \tag{5}$$

Various algebraic manipulations lead to the desired result

$$\beta = \frac{1}{2}\sin^{-1}\left(\frac{y}{L_0}\right) \tag{6}$$

Again, a modified version is used in practice

$$\beta = \frac{1}{2}\sin^{-1}\left(\frac{y - y_0}{L_0}\right) + \beta_0 \tag{7}$$

In the current design $L_0 = 90$ mm, $x + y \leq 8$ mm and the distance between TL and OL is 200 mm. Based on the above equations the maximum change in path length between TL and OL is estimated to be less than 0.2% using this method. A more conventional approach would have a maximum path length change of approximately 4%.

Initial alignment of the mirrors and objective proceeds as follows. A flat mirror is placed at the location of the back pupil of the objective. MM1's actuators are set at the middle of their travel range and the angles of MM2 are adjusted until the excitation beam is retroreflected onto itself, this sets $\alpha_0$ and $\beta_0$ for both mirrors. Now the objective is put back in place and aligned to the excitation beam, this sets $x_0$ and $y_0$. For widefield imaging the equations maintain acceptable alignment.

**e) SIM beam alignment**

SIM has more stringent alignment requirements. To meet these a motorized inspection system can be inserted into the beam path. Here a 50:50 beam splitter cube (BS, ThorLabs, BS013) picks off a portion of the beam which is imaged by a 200 mm fl lens onto an inspection camera (Basler AG, acA2040-90um). The BS is mounted in a rotation mount so that either the back pupil or the mask can be imaged (the inset photograph of the mask in Fig. S1 was taken with this system). To align the SIM excitation beams a 7-beam pattern, i.e. the average of three 3 beam patterns, is displayed on the SLM. Part of each beam is reflected off the surface of the sapphire coverslip and is imaged by the inspection camera. Fluorescence emitted from the sample allows the back pupil of the objective to be imaged simultaneously with the SIM pattern. As long as 3 of the peripheral SIM beams are detected a circle can be fit to them. Similarly, a circle can be fit to the image of the back pupil. MM1 and MM2 are iteratively adjusted (keeping their relative angles constant) until the centers of the two fitted circles are within some user specified tolerance, usually 0.5 pixels (2.75 µm).

**f) Chromatic focal shift**

Nikon specifies that the CFI L Plan EPI CRB 100x NA = 0.85 air objective used here is a "semi-apochromat", which means that maximum focal plane displacement should be <= 2.5x the depth of field (DOF) of the objective between 486 nm and 656 nm, and <=2x the DOF between either of those wavelengths and 546 nm.

In order to ensure that the registration of different wavelength channels is accurate, we characterized the chromatic aberrations in our system by imaging a field of TetraSpeck beads deposited onto the surface of the sapphire coverslip (cf. supplemental note 2.a)). The same beads can be excited by either 488 or 532 nm laser and imaged using following Semrock fluorescent filters: FF01-513/17, FF01-542/20, FF01-560/25, FF01-588/21, FF02-632/22, and FF02-684/24. We collected volumes scanning the objective axially to obtain 3D PSF's and determined the 3D PSF centers for all detection filters. We used these data to evaluate both axial chromatic aberration (dependence of focal plane position on wavelength) as well as transverse chromatic aberration (dependence of system magnification on wavelength).

The plot in Fig. S25A shows the dependencies of the focal plane position (in the object space, top) and of the relative magnification (bottom) on wavelength. Note: the change magnification is practically zero. Axis of the chromatic focal shift is co-linear with objective optical axis. We define a positive change as being in the direction *away* from the objective. In other words, with all other optical elements fixed the objective's focal plane at 675 nm is about 0.9 μm further from the objective than the focal plane at 500 nm.

##### g) Preparation for imaging

After the samples are loaded but before imaging can proceed the correction collar needs to be adjusted for each coverslip. A modified gear (S1268Z-096A180, Stock Drive Products/Sterling Instrument) is attached to the correction collar of OL using a nylon tipped set screw. The gear interlocks with a small stepper motor (CCM, 0824M012BAESM-4096+10/1 64:1+MG09 - DC Brushless Servomotor with MCBL3002 F AES RS controller, Faulhaber GMBH, MICROMO) such that the correction collar position can be computer controlled. Any slight variations in coverslip thickness can induce serious spherical aberrations because of the high index of sapphire ($n = 1.77$). All coverslips have fiducial markers, either attached to the surface or frozen in the cryo-protectant medium. To find the best setting for the correction collar the user iterates between adjusting the correction collar and acquiring a $z$-stack of a fiducial marker. Each $z$-stack of the fiducial, which is equivalent to the microscope's 3D point spread function (PSF) is inspected and iteration is halted once the width of the PSF in each direction is minimized and the support of the optical transfer function (OTF, the Fourier domain representation of the PSF) is maximized.

At the optimal correction collar position the PSF and corresponding OTF (Fig. S28A and B, respectively) of a bead (ThermoFisher, F8803) demonstrate nearly theoretical performance (white lines, Fig. S28B). Note that because an air objective is being used to image in vitreous ice (n=1.3, (*106*)) the effective axial resolution is reduced by approximately 50% and the focal plane moves 50% more than the objective. To see why the axial resolution is reduced while the lateral resolution is not note that the maximum extent of the OTF in the reciprocal lateral dimension is $2\mathrm{NA}/\lambda$ while the maximal extent in the reciprocal axial dimension is $\left(n-\sqrt{n^2-\mathrm{NA}^2}\right)/\lambda$ (cf. Fig. S28C for geometrical proof). More complex calculations (*107*, *108*), based on a real-space integral formalism agree well with the calculation presented here and show that a single correction collar position is sufficient to image ~15 μm of sample. The OTF based approach we describe offers the advantages of a more intuitive physical picture while also being fast and accurate.

To aid the selection of regions of interest for further super-resolution imaging each entire coverslip is imaged at a single z-plane. To do so, first the user marks the correct focal position at approximately 30 points on the coverslip. Then the program moves the objective in a raster scan acquiring an image at each grid point, interpolating the objective z-position based on the user chosen anchor points. These images are stitched together in a mosaic (cf. Fig. S12A) that is used to choose specimens for further investigation. The mosaic is also correlated to an x-ray image of the embedded sample to identify the cells of interest during the trimming step described in supplemental note 6.d).

**h) SIM**

Structured illumination microscopy is, in a sense, an interferometric microscopy in terms of the excitation pattern. In the case of 3D-SIM (2D-SIM) three (two) plane waves interfere at the sample to produce a sinusoidal standing wave. The quality of the final reconstructed SIM image depends on the quality of the interference produced which in turn depends on the relative polarization of the beams and their relative phase stability. For the later, air currents can have a detrimental effect near the back pupil or near any plane conjugate to it. At these planes the SIM beams are focused to spots that are ~2.5 mm apart from one another and index of refraction fluctuations caused by air currents can affect their relative phase. Fast phase jitter is averaged and reduces the overall contrast of the patterns at the samples while slow phase jitter can result in inconsistent phase stepping of the SIM patterns hampering reconstruction. The entire microscope has a cardboard shield built around it with internal baffles and lens tubes protect the light path where possible.

The contrast of the SIM excitation pattern at the focal plane of the objective is strongly affected by the relative polarization of the SIM beams. Lowest contrast occurs when the beams are p-polarized while highest contrast is when the beams are s-polarized. HWP2 and QWP1 (c.f. Fig. S1, lower right) are used to control the polarization state of the excitation beams at the sample plane. One might naively assume that only a single half waveplate is needed as the SIM beams are linearly polarized at the SIM mask. However, if the beams are not completely p-polarized or s-polarized when they are reflected by a mirror the reflected beam will have a different polarization state compared to the incident beam. This is because the mirror's reflectance for p-polarized light is different from that for s-polarized light and the differential reflectance of the two polarization states causes the reflected beam's polarization state to "twist" from that of the incident beam. Protective coatings exacerbate this effect and dielectrics are particularly bad as p and s-polarized light will have different reflectances *and* phase shifts upon reflection. Furthermore, the amount of "twist" will depend not only on the incident polarization state but also the wavelength. This effect is multiplicative, and our design has five protected silver mirrors (Newport, 20Z40ER.2) between the SIM mask and the back pupil of the objective.

Optimization of HWP2 and QWP1 proceeds as follows. For each wavelength and SIM pattern angle pair, images of beads are taken at linearly spaced phase steps of the SIM pattern. Each bead is fit to a two-dimensional Gaussian, which well approximates the PSF of the microscope, and the fitted amplitude of each bead as a function of phase step is fit to a sinusoid, $I(p) = A\ \cos(2\pi f p + \phi) + O$, where $I$ is the fitted intensity of the bead and $p$ is the phase of the SIM pattern. The contrast ratio is defined as $2\ A/(A + O)$. For a theoretically perfect pattern (where the nodes have zero intensity) and an infinitesimally small bead the contrast ratio will be 1. For the worst possible pattern, i.e. flat, the contrast ratio will be zero. The angles of HWP2 and QWP1 are iteratively adjusted until the median contrast ratio of all the beads in the field of view is maximized. For example, for 642 nm excitation waveplate optimization led to an improvement in contrast ratio from 0.69 to 0.92.

#### i) Localization Microscopy

Samples prepared for single molecule localization microscopy (SMLM) experiments have fiducial beads dispersed in the freezing media prior to freezing (cf. supplemental note 2.a)). These beads are used for three purposes: drift correction, alignment, and auto-focus. Before every SMLM experiment three z-stacks of images are acquired for each combination of excitation and emission wavelengths where the emission wavelength is longer than the excitation wavelength: a scaffold, a calibration and a short calibration. The scaffold is a z-stack of the selected region of interest taken with small z-steps (usually 10 nm) covering the entire axial extent of the sample without the cylindrical lens in place. The calibration is the same as the scaffold except with the cylindrical lens in place. Finally, a short calibration stack is taken which is used to calibrate the autofocus routine described below. Data from the large calibration stack is used to determine the axial position of each localized molecule as described in supplemental note 15.a). The scaffold serves two important functions: 1) it allows the different data sets to be aligned to a common frame of reference and 2) it allows for the aberration induced by the cylindrical lens to be corrected. The aberration is primarily the differential magnification of the sample along the $x$ and $y$ axis and is well characterized by an affine transformation. The second function is subtle but important, without this correction the data is neither a good representation of reality nor can it be correctly correlated to the other light microscopy data or the electron microscopy data.

For 3D-SMLM on specimens thicker than ~0.5 µm multiple focal planes need to be acquired. During the experiment the focal plane is moved in 0.2 µm steps every 250 frames. Each data slab is independently processed (cf. supplemental note 15) and then the slabs are stitched together in one of two ways: 1) the slabs can be registered to their neighbors directly via common fiducial markers, or 2) the slabs can be registered directly to the scaffold data set. The second method opens up the ability to sample non-overlapping planes independently from one another. Once the slabs are registered to each other set using a translation only registration model they can be aligned to the scaffold using an affine transformation model.

SMLM experiments have a long duration (multiple days) resulting in a significant drift. Correcting the drift in the lateral plane can be done as a post processing step (cf. supplemental note 15.b)) as long as the drift is negligible during the camera exposure time and not so large that the sample drifts out of the field of view of the microscope during the experiment. Axial drift presents a more difficult challenge as the axial "field of view" of the microscope is on the order of a single micron. Put another way, if the axial drift during the entire course of the experiment is larger than 1 micron it will be impossible to correct during post processing. To prevent this drift an auto-focus routine runs at a predefined interval during the experiment. To set up the auto-focus the user selects fiducials in the short calibration data set. Each fiducial is fit to an elliptical 2D Gaussian in every frame of the calibration stack and the ellipticity as a function of axial position is calculated. The ellipticity is defined as $(w_x - w_y)/(w_x + w_y)$ where $w_x$ and $w_y$ are the fitted standard deviations of the Gaussian in the $x$ and $y$ directions (not to be confused with the localization precisions in the $x$ and $y$ directions). The first time the auto focus routine is run at the beginning of the experiment the ellipticity of each fiducial is recorded. When the auto focus routine is run subsequently it calculates the median drift of all fiducials and if that drift is greater than a user defined dead band (usually 50 nm) the objective is moved to compensate for the drift. Autofocus is only run on one focal plane in multi-slab experiments.

As described in the main text, cryo-SMLM experiments on standard fluorophores require high power excitation (~1-10 kW/cm$^2$)(*29*). Unfortunately, the objective used in this microscope was originally designed for bright field microscopy and not the high-power laser excitation used for SMLM. In early experiments where standard wide field illumination was used the objective would fail. Specifically, burn marks would form on internal surfaces/interfaces presumably due to local heating at a focal point of the excitation beam. To avoid damaging the objective lens we used SIM-style excitation. The only difference between this excitation and that used for standard 2D-SIM imaging is that the sample was exposed to each orientation and phase pattern for 1 ms and the number of phases per orientation was increased from 5 to 8. Thus, for a 50 ms exposure the sample is exposed to ~2 full rounds of SIM patterns and the effective illumination is indistinguishable to wide field illumination except that the objective is subjected to half the power at each focal spot (the patterns are designed to deposit the majority of the energy into the two diffraction spots) and heat is allowed to dissipate.

#### 4. *Photophysical Analysis of Single Molecules at Cryogenic Temperatures*

##### a) Identifying Single Molecules in the Data

Fig. S8 presents our pipeline for identifying and analyzing single molecules within our cryo-SMLM data sets. First, fiducials (marked as magenta circles in Fig. S8A) are identified as described in supplemental note 15.b)i) and all locations within a 10-pixel radius of the fiducial centroid are removed. Next, high density regions (cf. Fig. S8B, magenta lines), in this case regions containing mitochondria, are removed by following the procedure outlined in Fig. S29D-F and supplemental note 5.b). The remaining localizations are clustered using the HDBSCAN algorithm (hdbscan 0.8.19, (*109*)). Clustered localizations are assigned a p-value (indicated by color in Figure S8B) by comparing clusters to simulated clusters (see note 5.b)) of the same localization precision. Clusters of localization events coming from a single molecule can be thought of as random samples from the underlying molecular distribution, in this case a single molecule. We can compare our experimentally measured cluster (a distribution) to one simulated from the experimental data using the Kolmogorov-Smirnov (KS) test. Note, because the x, y and z coordinates of the localizations are assumed to be independent we calculate the p-value for the KS test for the x, y and z distributions independently and assign the minimum p-value of the three to the cluster. In this case, the null hypothesis assumes that the experimental distribution represents a single molecule and the p-value indicates with what statistical confidence we can reject that hypothesis. Because we want to retain clusters that are from single molecules we want to remove clusters with low p-values. In general, we remove clusters with p-values less than 0.25 before proceeding with further analysis. To make this idea concrete Figure S8C) shows three clusters with the three different p-values assigned by the previous procedure rendered by replacing each localization (indicated by red crosses) with a normalized gaussian with standard deviations in the x and y directions equivalent to the localization's precision in each direction. Note that the far-left example is unlikely to have arisen from a single molecule while we can be much more confident in claiming that the following two examples do.

Once purported single molecules have been identified they can be transformed into blinking traces (cf. Figure S8D). Here each frame in the experiment is assigned either a 1 or a zero depending on whether an emission event was identified and localized in that frame. Individual ON and OFF times are extracted from these traces. Note that an OFF time is only considered if it is bounded by two ON times. All ON or OFF times from all single molecules in a given experiment can be aggregated into histograms (cf. Figure S8E) which show approximately power-law behavior. The same analysis can be applied to different fluorophores and temperatures (cf. Figure S8F-G).

**b) Dynamic Contrast Ratio**

Dynamic contrast (Figure 2A) is defined as the ratio between the lengths of a dark period (an "OFF" time) and the subsequent emissive period (an "ON" time) (cf. Fig. S8D). Figure 2A presents the median dynamic contrast ratio for different fluorophores and temperatures. Dynamic contrast reflects the temporal separation of different emission events, a crucial ingredient for successful SMLM.

**c) Static Contrast Ratio**

Another key factor in SMLM is the brightness of each emissive event relative to the background signal, which can be quantified as the static contrast ratio (Figure 2B). Fig. S9 presents the processing pipeline to calculate the static contrast ratio from a SMLM data set. Data is presented from a U2OS cell expressing mEmerald-TOMM20. The first row shows the final SMLM image, an early frame, and a frame later in the experiment once single molecule blinking is achieved. Fiducial locations are identified following supplemental note 15.b)i) and high density, i.e. biologically labeled, areas are identified following supplemental note 5.b); both areas are indicated in the second row with white (fiducials) and cyan (mitochondria) lines, respectively. Peaks are identified as pixels that have a value greater than $b + 2\sqrt{b + e^2}$ where b is the background counts within the mitochondrial areas and e is the camera read noise. We measured e to be ~2.45 counts. Essentially, we are looking for peaks that have a signal to noise ratio of 2 or higher. Groups of connected pixels are considered to be the same peak. Identified peaks are indicated in (H) and zoom views are shown in (I). The median value of the counts outside the mitochondrial and fiducial masks is the background value ($img_{bg}$), the median value of the counts inside the mitochondrial mask is the peak background value ($peak_{bg}$) and the mean value within the mitochondrial mask and the fit windows shown in (H) is the peak value ($peak$). Static contrast is calculated as $SC = (peak - peak_{bg})/\max(peak_{bg} - img_{bg}, 1)$ for each frame in the data set. For instance, for the frame shown in (C, F, G, H) the contrast ratio is 26. Figure 2B presents the median static contrast value for the last hour of the experiment, error bars are the standard deviation of the static contrast over the last hour of the experiment.

#### *5. Grouping*

In SMLM single molecules frequently appear in more than one camera frame. Sometimes the molecules will remain bright in consecutive frames but more often they will blink, in other words the molecules will change their state between a bright one and a dark one without photochemical bleaching (i.e. destruction of the molecule). Some authors have referred to this blinking behavior as photophysical bleaching. Consequently, the final data set and final rendered image will have spurious clumps of localizations that do not reflect the actual underlying distribution of fluorophores in the sample (cf. Fig. S11). To correct for these phenomena and to generate a data set that more faithfully represents the underlying fluorophore distribution, localizations due to the same molecule need to be identified and combined (grouped) into a single localization event.

##### a) Identifying Groups

To group localizations we use the fact that the experiment is time ordered meaning that for a given localization we only need to look in the subsequent frames for potential matches not the previous frames. All localizations in the first frame are assigned a unique identifier and these data become our "cache." The cache is compared to the next frame using a k-d tree algorithm (as implemented in SciPy (*110*)) to find neighbors within a certain radius (the grouping radius, $r_g$). For all matches the cache values are updated. All localizations in the new frame that do not pair with a cache value are added to the cache with a unique identifier. Cache localizations that have not been updated for a given number of frames (the group gap, $g$) are purged. At the end of the algorithm all localizations in the data set have a been assigned to a group. The grouping algorithm requires two input parameters: the grouping radius ($r_g$) and the group gap ($g$) the determination of both are described below.

##### b) Data Driven Determination of Grouping Parameters

From previous reports (*7*, *27*, *29–34*) and our photophysical data (Fig. 1, Fig. S8 and supplemental note 4) we know that at cryogenic temperatures bleaching events are extremely rare and single molecules can turn off for very long times (on the order of hours) before returning to an emissive state. Therefore, we want to be as aggressive as possible with our grouping parameters while minimizing erroneous grouping between different molecules.

To choose a grouping radius in a principled manner we begin by finding the localization PSF aspect ratio ($\alpha$, cf. Fig. S29A) which is defined as the median aspect ratio for all non-fiducial localizations. An exemplary $\alpha$ distribution and its median value are shown in Fig. S29A. Next, $M$ (where $M$ is usually 4096) synthetic groups of localizations are simulated as follows. For each group $N$ (where $N$ is nominally 512) localization precisions are sampled from the experimental data. The effects of residual drift are modeled by adding, in quadrature, the residual drift to the sampled precisions (eq.(8)). Finally, a set of $N$ three dimensional coordinates is generated by sampling one set of coordinates from a normal distribution described by one set of localization precisions (eq.(9)). The entire process is described by the following equations:

$$\tilde{\sigma}_i = \sqrt{\sigma_i^2 + \Delta^2} \tag{8}$$

$$x_i \sim \mathcal{N}(0, \tilde{\sigma}_i^2) \tag{9}$$

Where boldface indicates vectoral quantities, $\boldsymbol{\sigma}$ are localization precisions, $\boldsymbol{\Delta}$ are residual drifts, $\mathcal{N}(\mu, \sigma^2)$ is a normal distribution with mean $\mu$ and standard deviation $\sigma$, and $\boldsymbol{x}$ are coordinates. For each localization the normalized radius is calculated as $\bar{r}_i = \sqrt{x_i^2 + y_i^2 + (z_i/\bar{\alpha})_i^2}$, an exemplary distribution of $\bar{r}_i$ for a single group is shown in Fig. S29B, along with the 90$^{th}$, 99$^{th}$, and 99.9$^{th}$ percentile cutoffs. Distributions for $M$ simulations are shown in Fig. S29C, along with median values. We chose the median of 99$^{th}$ percentile bootstrap distribution (blue line in the middle plot of Fig. S29C) for the grouping radius to balance the desire to capture all information from each molecule while minimizing the risk of grouping information from spatially adjacent molecules.

Normally, the group gap ($g$) would be determined from the bleaching rate of a given fluorophore. For instance, if we knew a given fluorophore bleached within 1 minute of its first detection 99% of the time then it would be unlikely that two sequential events separated by more than a minute would have come from the same single molecule. However, under cryogenic conditions bleaching is rare therefore we want to be as aggressive in setting $g$ as we can be without compromising image quality by combining events from different molecules. We can develop a heuristic algorithm to determine $g$ as follows. Ignoring the spatial component of grouping for a moment and focusing on the temporal one we can see that the worst case would be if all blinking events within $r_g$ were uniformly distributed in time. If this were true, and we ignored the spatial component, then there would be no way to group these data, to see why consider the following argument. Let $g_0 = \left(\frac{\text{time of experiment}}{\text{\# of events within } r_g}\right)$; if $g < g_0$ then every event would be treated as a separate group, conversely, if $g \geq g_0$ then every event would be treated as the same group. We can use this as the limit for our grouping; if we can determine the characteristic density of our sample, $\rho$, then we can calculate $g_0$, given we know the normalized grouping radius, $r$, using the following equation: $\frac{1}{\rho} = g_0 = g\frac{4}{3}\pi r^3$ ($g\pi r^2$ if the analysis is done in 2D). Rearranging yields:

$$g = \frac{\rho^{-1}}{\frac{4}{3}\pi r^3} \tag{10}$$

To determine $\rho$ we use the following heuristic algorithm. First the localization data is binned into a histogram image with voxels (pixels) of a given volume (area) (cf. Fig. S29D); empirically we have found that using voxels with normalized edges equivalent to 1 camera pixel is best; in our case 130 nm. If the voxels (pixels) are too small the result is dominated by shot noise and if the voxels (pixels) are too large too much background noise will be included. Dividing the histogram image, which is in units of events per unit volume (area), by the total number of frames in the experiment results in data with units of events per unit volume (area) per unit time. The largest 1% of pixels are removed as outliers. Biologically labeled areas of the histogram image are determined by the triangle threshold algorithm (*111*). This step is crucial because the density of fluorophores will be much higher there then in the background areas and including background regions will result in erroneously low $\rho$. Finally, $\rho$ is set as the $99^{th}$ percentile of values in the labeled region of the histogram image. A histogram of histogram image pixel values is shown in Fig. S29E with the triangle threshold (background) shown in blue and $\rho$ shown in red. Fig. S29F shows the same data as (D) with the background in blue and high-density regions with a density greater than $\rho$ shown in red.

In order to improve upon the heuristic algorithm outlined above we ran simulations (cf. Fig. S30) using single molecule data collected for mEmerald and JF525 (see supplemental note 4.a)). Fig. S30A presents the timing of emissive events from 8 single JF525 molecules collected from a high pressure frozen U2OS cell expressing Halo-TOMM20 conjugated to JF525-HT imaged at ~8 K. The numbers on the left axis indicate the total number of events depicted on each line. All 8 molecules were spatially separated from one another in the original data set, but we can use them to model how our grouping procedure would handle them if they were found within a single grouping volume. The left most plot shows emission events color coded according to the molecule from which they originated. The next three plots show the resulting groups as a function of grouping gap. We want to develop a metric that will indicate which grouping gap results in optimal grouping. We chose the difference in molecular probabilities (DMP) between our proposed grouping and that of the ground truth. Fig. S30B presents the 2D DMPs for the three grouping gaps in (A). Empty circles and crosses show ground truth and proposed groupings, respectively. Fig. S30C shows the root mean squared (RMS) of the DMP for a variety of grouping gaps on a logarithmic scale, the gaps from (A) are indicated as colored circles. A gap of 26.2 minutes is optimal for this collection of localizations. We can repeat this procedure for many different random samplings of single molecule traces of different sizes. Fig. S30D presents a scatter plot of optimized grouping gap versus number of events per frame for mEmerald (blue) and JF525 (orange) along with a power-law fit to the data (dashed green line) and the heuristic algorithm (eq. (10), red dotted line). These simulations lead to eqn. (10) to be modified as

$$g = \frac{a\rho^{-b}}{\frac{4}{3}\pi r^3} \tag{11}$$

Where $s = 10.274$ and $b = 0.785$ for mEmerald and JF525. Our simulations allow us to be even more aggressive (i.e., use a larger grouping gap) in our grouping than the heuristic algorithm would have suggested. However, the heuristic algorithm is recommended for fluorophores for which no photophysical data exists.

Fig. S11 presents the aggregated improvements in grouping developed in this section with two exemplary data sets: a U2OS cell expressing mEmerald-Sec61β and a U2OS cell expressing Halo-Sec61β (JF525). The top row of (A) shows a low magnification ROI the two SMLM images. The subsequent rows show zoom images of the indicated ROIs. The last two rows are color-coded according to the frame in which a grouped localization first appeared. The first and third columns show the results of *ad hoc* grouping in which the grouping radius and grouping gap were chosen to be 26 nm and 1.6 seconds, respectively, for mEmerald and 26 nm and 2.6 seconds for JF525. The second and fourth columns show the results of our developments. The results are particularly striking for JF525 where the time-correlated clumps are greatly reduced with our new grouping method. Presented in (B) are the pair correlation functions of the data shown in the last row of (A).

In all cases the uniformity of the sample is increased with our new method. The molecular density of the data shown in the last row of (A) are shown as bars in (C).

##### c) Aggregating Groups

Groups are aggregated with the following equations for localization coordinates (14) and precisions (15).

$$w_{i,j} = \frac{1}{\sigma_{i,j}^2} \tag{12}$$

$$\overline{w_i} = \sum_j^n w_{i,j} \tag{13}$$

$$\overline{c_i} = \frac{1}{\overline{w_i}} \sum_j^n w_{i,j} c_{i,j} \tag{14}$$

$$\overline{\sigma_i}^2 = \frac{n}{(n-1)\overline{w_i}^2} \sum_j^n w_{i,j}^2 \left(c_{i,j} - \overline{c_i}\right)^2 \tag{15}$$

Where $n$ is the number of group members, the subscript $j$ indicates the group member, the subscript $i$ indicates the coordinate (i.e. $x$, $y$, or $z$), $c$ indicates localization coordinate, $\sigma$ indicates localization precision in each coordinate, and bars over variables indicate that they are the aggregated value for the group. Equation (14) states that the grouped coordinates are simply the weighted means of the group coordinates using the inverse square of the localization precision in each direction as the weighting factor. Normalization of the weighting is achieved by dividing by the sum of the weights. The equation for determining the grouped localization precision of the groups, eq. (15), is more complex.

Several models have been proposed to estimate the grouped localization precision ($\bar{\sigma}_i$) of a single blinking molecule from a series of grouped localizations. For instance, Shtengel et al. (*89*) calculated $\bar{\sigma}_i$ as

$$\bar{\sigma}_i^2 = \frac{1}{2}\left(\frac{\sum_j^n w_{i,j}\left(c_{i,j} - \overline{c_i}\right)^2}{n\overline{w_i}} + \frac{1}{\overline{w_i}}\right) \tag{16}$$

and Legant, et al. (*112*) calculated $\bar{\sigma}_i$ as

$$\overline{\sigma_i}^2 = \frac{1}{\sum_j^n 1/\sigma_{i,j}^2} \tag{17}$$

However, what is desired is an estimate of the error on the weighted mean (the aggregated localization coordinate, $\overline{c_i}$). In the unweighted case one can use the standard formula for the standard error of the mean ($\sigma_\mu = \sigma/\sqrt{n}$) but for the weighted case there is no agreed upon formula for the standard error of the weighted mean. A proposed one can be found in (*113*) (eq.(15)).

To test which of these models best fits the true precision, we ran a numerical simulation in several stages (cf. Fig. S31). First a set of localization precisions are chosen from either a theoretical distribution (cf. Fig. S32A) or an experimental distribution (cf. Figs. S32B and S32C) and synthetic localizations are generated as described in supplemental note b). The entire set of localizations is split into *N* groups of *M* localizations (cf. Fig. S31 top row). Each group can be grouped into a single localization (black crosses) using eq.(14). The grouped localization precisions of each group calculated by equations (15), (16) and (17) are shown as the purple, red, and yellow ellipses, respectively. These estimates are calculated for *N* groups and the resulting distributions of estimated grouped localization precisions are shown as the three histograms at right. To estimate the true distribution of grouped localization precisions we generated *N* bootstrap samples of *M* grouped coordinates (black crosses, bottom row). We considered the distribution of the sample standard deviation (green ellipses, magnified 10X) of the bootstrap samples to be the ground truth grouped localization precision distribution (green histogram, at right).

For simulations of synthetic data (cf. Fig. S32A) the square of the localization precisions were sampled from a scaled $\chi^2$ distribution (*114*), i.e. $\sigma_{i,j}^2 \sim \chi_k^2/k$ where *k* is known as the degrees of freedom (DoF) of the distribution. As the *k* increases the distribution becomes more sharply peaked around 1. Because all coordinates are treated independently during SMLM processing we only simulated a single dimension. Histograms of the grouped localization precisions are shown in green, purple, red and yellow for ground truth and eqs. (15), (16) and (17) respectively. In every case *N*=10,000. Each column and row depict a simulation for different values of *M* and *k*, respectively. Only when the distribution of localization precisions is broad (*k* is small), and the group size (*M*) is large does eqn. (15) fail to mirror the ground truth (note these simulations do not include residual drift). Simulations wherein the localization precisions are sampled from experimental distributions of mEmerald and JF525 (cf. Fig. S32B and Fig. S32C) were run similarly.

#### 6. *Preparation for Electron Microscopy*

A critical step of the CLEM process is identifying areas of the sample to image with electron microscopy that have already been imaged with light microscopy. We have followed a protocol similar to the one previously described (*115*). Once all optical experiments are completed the samples are unloaded from the instrument (supplemental note 6.a)), freeze substituted and resin embedded (supplemental note 6.b)), have their coverslips removed and are re-embedded prior to X-ray imaging (supplemental note 6.c)). Finally, the resin embedded samples are trimmed to the identified areas (supplemental note 6.d)) by correlating the X-ray image to a widefield cryo-optical fluorescence map of the entire coverslip (cf. supplemental note 3.g) and Fig. S12A).

##### a) Sample unloading

Sample unloading is done in following steps (cf. Fig. S1 and Fig. S5):

1. CTD is attached to the airlock of CS
2. The airlock is evacuated
3. GV is opened
4. SH is detached from MCF and retracted into CTD, which is then sealed.
5. GV is closed, the airlock is vented and CTD is docked to the sample preparation chamber.
6. SH is transferred into MCF of the preparation chamber.
7. The coverslips are removed from SH and their photos are taken (cf. Fig. S12B)
8. Coverslips are then stored in cryo boxes (CB) for subsequent freeze-substitution and resin embedding.

##### b) Freeze-substitution and resin embedding

After coverslips are removed from the cryogenic microscope, they are prepared for room temperature electron microscopy by freeze substitution and heavy metal staining followed by embedding in resin. During this process coverslips are uniquely identified by scratches on their gold-plated surfaces (cf. Fig. S12B-C, red arrows).

i) Freeze-substitution

Freeze-substitution (FS) was performed with a protocol adapted from (*116*). Briefly, coverslips were transferred to cryotubes containing FS media (2% $OsO_4$, 0.1% Uranyl acetate, and

3% water in acetone) under liquid nitrogen and the following FS schedule was executed using automated FS machine (AFS2, Leica Microsystems):

1. 140°C to -90°C 2h
2. -90°C to -90°C 24h
3. -90°C to 0°C 12 h
4. 0 to 22°C 1h
5. 22°C 1 h

ii) Resin Embedding

Resin embedding was performed immediately after FS. Samples were removed from the AFS2 machine, washed 3 times in anhydrous acetone for a total of 10 min and embedded in Eponate 12 with the following protocol:

1. Acetone/Eponate 12 2:1 1h
2. Acetone/Eponate 12 1:1 1 h
3. Acetone/Eponate 12 1:2 1 h
4. Eponate 12 2 h
5. Eponate 12 2 h
6. Eponate 12 2 h

Coverslips were placed in the slots of a flat embedding silicone mold, cells side up, and immersed in Eponate 12 which was polymerized for 48 hours at 60°C.

**c) Re-Embedding**

Following EPON embedding, the epoxy is removed from all coverslip surfaces not containing cells using a razor blade. Then the coverslip is separated from the resin block containing the cells by sequential immersion in liquid nitrogen and hot water. In a process similar to pothole formation, hot water gets into the cracks between the resin and the coverslip and then expands when it freezes in the liquid nitrogen. Moreover, the sapphire coverslip and polymer resin have very different thermal expansion coefficients. Both mechanisms combine to ensure easy coverslip separation.

Once the coverslip is removed, the exposed surface is immediately re-embedded in Durcupan resin which helps minimize streaks during FIB-SEM imaging (*12*).

##### d) X-Ray correlation and trimming

Following Durcupan re-embedding, an X-Ray of the entire block is taken using an XRadia 510 Versa micro X-Ray system (Carl Zeiss X-ray Microscopy, Inc.). By correlating the X-Ray image (cf. Fig. S12D) and the fluorescent map (cf. Fig. S12A), we can identify the areas in the resin block (cf. Fig. S12E) that we wish to prepare for FIB-SEM imaging. Correlation is straightforward as many landmarks are easily identifiable in both images, exemplary landmarks are indicated with yellow arrows in Fig. S12A, D, E. Once the desired area is identified the sample block is re-mounted on a copper stud using Durcupan (cf. Fig. S12F) and trimmed using an ultramicrotome (EM UC7, Leica Microsystems). Trimming is done in few iterations with X-Ray images taken between the steps to ensure accuracy (cf. Fig. S12G, H). A cross-section of an X-Ray tomogram overlaid with the MIP of 3-channel cryo-SIM image is presented in Fig. S12I.

As a final preparation step, the sample is sputter coated with 10 nm of gold and 100 nm of carbon in a sputter-coating system (PECS 682, Gatan).

#### *7. Correlation and registration*

Correlation lies at the heart of any correlative imaging technique and determines to what spatial error a fluorescently labeled protein of interest can be located within an electron micrograph; therefore, correct registration and quantification of correlation accuracy are critical. In general, data recorded in different modalities are registered using fiducial markers visible in all modalities. Gold nanoparticles have been demonstrated to work well for thin, nearly two-dimensional, samples (*115*) but are unsuitable for the method described here because it is difficult to place them uniformly throughout a 3D sample while preserving that placement after all sample processing steps. Therefore, a different volumetric registration method is needed, especially for protocols using freeze-substitution and resin embedding where deformations, such as swelling or shrinkage, between vitrified and embedded samples occur.

To describe our process, we will use the data presented in Fig. 2 and Fig. 3 as an example. First, the different fluorescence data volumes (Halo-TOMM20 (JF525) and mEmerald-ER3) are registered to each other using the fluorescent spheres suspended in the vitreous ice as fiducials. Second, the FIB-SEM is registered to the two-color LM volume by using organelle, specifically the mitochondria and ER, landmarks (cf. Fig. 2B). Used separately the mitochondrial or ER landmarks form two local displacement vector fields, $\overrightarrow{DF}DF_{mito}$ and $\overrightarrow{DF}DF_{ER}$, mapping the 3D-EM volume to the LM volume. These mappings represent two estimates of the same underlying deformation that the sample has undergone between the two imaging modalities. The mean absolute error of correlation $\langle\varepsilon\rangle$ between the estimated deformation and ground truth deformation $\overrightarrow{DF}DF_{true}$ should be:

$$\langle\varepsilon\rangle = \frac{1}{N}\sum_{i=1}^{N}\left|\overrightarrow{DF}DF_{mito_i} - \overrightarrow{DF}DF_{true_i}\right|$$

or (18),

$$\langle\varepsilon\rangle = \frac{1}{N}\sum_{i=1}^{N}\left|\overrightarrow{DF}DF_{ER_i} - \overrightarrow{DF}DF_{true_i}\right|$$

where summation is done over all image voxels. Assuming that $\overrightarrow{DF}DF_{mito}$ and $\overrightarrow{DF}DF_{ER}$ are statistically independent, we can write:

$$\langle\varepsilon\rangle = \frac{1}{\sqrt{2}}\frac{1}{N}\sum_{i=1}^{N}\left|\overrightarrow{DF}DF_{mito_i} - \overrightarrow{DF}DF_{ER_i}\right| = \frac{1}{\sqrt{2}}\frac{1}{N}\sum_{i=1}^{N}\left|\overrightarrow{\Delta DF}\Delta DF_i\right| \quad (19).$$

The difference between the two deformation fields, $\Delta DF = DF_{mito} - DF_{ER}\overrightarrow{\Delta DF}\Delta DF = \overrightarrow{DF}DF_{mito}\overrightarrow{DF}DF_{ER}$, gives an estimate of the registration error (times $\sqrt{2}$). To calculate the deformation fields and to warp the image volumes, we used BigWarp, a Fiji plugin (*43*). A total of 448 landmarks (217 mitochondrial and 231 ER) were identified within the EM and LM volumes (Fig. 2A and Fig. S15A). The landmark selection process is illustrated in Fig. 2B.

Fig. 2A presents the non-affine component of the displacement field generated using the combined set of both the mitochondrial and ER landmarks as vectors originating from each landmark position. The length of vector is proportional to the magnitude of the non-affine component of the displacement field at that point, and the color represents the magnitude of $\Delta DF$. Since $DF_{mito}$ and $DF_{ER}$ correspond to two separate wavelength channels, the difference between the two displacement fields gives the upper bound estimate of the overall registration error as it includes the registration error between the two optical channels and the errors of registering each optical data set to the EM data set. Of course, this estimation is only accurate in the areas where both mitochondrial and ER landmarks are present, this region is indicated by the dark pink surface in Fig. 2A and the grey contour in Fig. S14A. A histogram of the magnitude of $\Delta DF$ in that area is shown as grey in Fig. 2C and peaks at 92 nm with a mean value of 145 nm. However, a higher density of landmarks can improve correlation accuracy. To demonstrate this, we selected 60 landmarks within the 61 $\mu m^3$ sub-volume indicated by the red box in Fig. 2A. A histogram of the magnitude of $\Delta DF$ within this sub-volume is presented in Fig. 2C (red) and has a peak of 32 nm and mean of 42 nm.

Fig. S14A presents all the ER (circles) and mitochondrial (squares) landmarks used for CLEM overlaid on the "mid-cell" FIB-SEM slice (cf. Fig. S33). Landmarks are color-coded according to distance from the coverslip. Fig. S15 presents cross-sections through exemplary CLEM sub-volumes centered on the ten landmarks labeled in Fig. S14A. Fig. S14B presents the "mid-cell" slice of the magnitude of $\Delta DF$ which illustrates that the registration accuracy is higher in the thinner areas of the cells where landmarks in the ER channel can be more easily identified. Note that the areas around the perimeter of the cells, which have high values of $\Delta DF$, do not indicate high registration errors. The purportedly poor registration in these areas is actually due to the fact that there are no mitochondrial landmarks present, and thus our cross-registration error evaluation procedure cannot be applied. The magenta contour in Fig. S14A is a projection of the mask within which $\Delta DF$ is estimated and excludes the above-mentioned areas.

#### 8. *Correlative driven segmentations of peroxisomes*

Generalized methods for the segmentation of EM images is an open and difficult problem. However, SR-CLEM data offers a path to specialized segmentation algorithms that leverage the sparsity and specificity of SR-LM. To demonstrate this possibility, we used our correlative data of SKL-mEmerald expressing HeLa cells (cf. Fig. 4 and Fig. S14C-D) to automatically identify and segment peroxisomes in the EM data. Our algorithm consists of 5 steps:

1) Identification of peroxisomes in the LM data
2) Extraction of small EM ROIs
3) Segmentation of membranes in the EM data
4) Identification of peroxisomes in the segmented EM data
5) Filtering failed segmentations.

Peroxisome locations were identified in two ways, either thresholding or difference of gaussians (DoG) blob detection (*117*). The threshold was determined using Li's iterative Minimum

Cross Entropy method (*118*) as implemented in SciKit-Image (*119*) applied to the axial MIP of the SKL-mEmerald SIM image. Using this threshold objects in the 3D-SIM volume were identified and objects smaller than the PSF were removed. An attempt was made to split each object into smaller objects using a watershed algorithm in order to separate spatially adjacent objects. In the DoG approach the lateral centroids of peroxisomes were first found in the axial MIP of the SIM data and then the axial position was found by determining the location of the brightest signal in a 0.312 μm square *xy* patch surrounding centroid. Objects found using both methods were combined into a single set with object locations found by thresholding taking precedence.

Object bounding boxes were refined using the SMLM data as follows. A subset of the SMLM data corresponding to the initially found SIM bounding box is extracted and the new bounding box is calculated as a rectangle where each side is centered on the median SMLM location and has a length that is twice the max of the $1^{st}$ to $50^{th}$ percentile and the $50^{th}$ to $99^{th}$ percentile. Using these bounding boxes subsets of the EM data are extracted for further processing.

Membranes and ribosomes in each EM subset are segmented with a trained random forest classifier using Ilastik (*83*) using all possible features. The background, i.e. the parts of the image that are classified as neither membrane nor ribosome, is split into objects using a watershed algorithm. Any objects smaller than 262,144 $nm^3$ or touching the border are removed. Of the remaining objects the one whose surface is closest to the center of the bounding box, and therefore the centroid of the LM signal, is chosen as the peroxisome.

The algorithm can and does fail. An unsupervised clustering algorithm, Ward's method (as implemented in scikit-learn (*120*)), was used to split the set of segmentations into successful and failed segmentations using the surface area to volume ratio, roundness (defined below), and two different measures of surface roughness (defined below) as features. Roundness is calculated as $SA_{sphere}/SA_{object}$ where $SA_{object}$ is the surface are of the object and $SA_{sphere}$ is the surface area of a sphere with the same volume as the object. Thus, roundness is bounded between 1, for a sphere, and 0, for a plane. The two measures of surface roughness are the fractional changes in either the surface area or the surface area to volume ratio between an object and a smoothed version of itself. The resulting successful segmentations are shown in Fig. S17 ordered in terms of increasing volume and color coded according to roundness (a scatter plot of surface area and volume is shown in Fig. 4).

#### *9. Neuronal Adhesion Curvedness Calculation*

First, the membrane between the two cell bodies that was collocated with the JAM-C SIM signal (white box, Fig. 6) was manually segmented using Amira 6.7 (Thermo Fisher Scientific). Next, the electron dense and lucent regions of the segmented membrane were separated using the magic wand tool in Amira (Fig. 6F). The resulting segmentation was converted into a triangular mesh surface using the 'Generate Surface' module in Amira with constrained smoothing and a smoothing extent of 3 (see Amira documentation and ref. (*121*)). Further smoothing of the surface was performed using the 'Smooth Surface' module in Amira with 50 iterations and lambda equal to 0.6 (see Amira documentation and ref. (*122*)). The final smoothed surface was split into two surfaces, one for the electron dense membrane and one for the electron lucent membrane (Fig. 6H-I, blue and red, respectively). Curvedness for each membrane component was calculated using the 'Curvature' module in Amira using the 'on triangles' method with the parameters nLayers = 1 and nAverage = 5 (see Amira documentation). The curvedness for each surface triangle is defined as $\frac{1}{2}\sqrt{C_1^2 + C_2^2}$ where $C_1$ and $C_2$ are the two principal curvatures for the parabolic approximation of the surface in the neighborhood of the triangle.

#### *10. LLSM Imaging of GNP Nuclear Condensation*

The LLSM optical path was previously described in detail (*78*). Single timepoints (3 biological replicates totaling N=71 and N=85 for each GNP and CGN cohort, respectively) and time-lapses (3 biological replicates totaling N=5 for each GNP and CGN cohort) of all LLSM samples were carried out in z-stage + objective scan mode (where the detection objective and the light-sheet moved together in discrete steps). Here we used two pairs of cylindrical lenses to illuminate a thin stripe on a spatial light modulator (Forth Dimension QXGA with 2048 by 1536 pixels) to generate a lattice light sheet ~160 µm wide along the y-axis. A two-camera solution was used with dichroics and emission filters (Semrock Di03-R561-t1-25x36; emission filter on the camera 1 is a Semrock FF01-530/43-25; emission filter on the camera 2 was a Semrock LP02-568RU-25), which allowed readout of one camera during exposure of the other for rapid acquisition. All LLSM images were acquired over a 512 x 512-pixel field of view resulting in 0.104 µm x-y pixels while dithering the light sheet with a dithering amplitude of 10 µm. All volumes were acquired with 151 z-axis planes spaced by 0.2 µm to satisfy Nyquist sampling. For time-lapse imaging, images were acquired every

10 minutes for 12 hours. Fields of view were selected to contain Atoh1-EGFP positive cells representing GNPs and Atoh1-EGFP negative cells representing CGNs. Sample chamber conditions were optimized for >12-hour neuron imaging. This includes the use of thermoelectric cooling (TEC) control for maintaining 37°C media temperature. pH and $O_2$ homeostasis were maintained by perfusing $CO_2$ and $O_2$ into the top of the chamber directly over the coverslip. $DiH_2O$ was slowly dripped into the chamber using a syringe pump to compensate for evaporation in the media. By doing so, media levels remained at equilibrium, and the focus of the samples was maintained through the entirety of acquisition.

Subsequent imaging processing was necessary to obtain LLSM volume calculations. First, all LLSM datasets were deconvolved after background subtraction using a Richardson-Lucy maximum likelihood algorithm adapted to run on FIJI, using experimentally measured PSFs for each color channel for 10 iterations (*123*). Then channel alignment corrections for the LLSM data were performed in the Data Transformation Gallery in Arivis 4D to interactively apply channel shift transformations to optimize channel overlap. For time-lapse data, LLSM image photobleach correction were performed using the Histogram Matching method contained in the Bleach Correction plugin in ImageJ (*124*). Next, deconvolved LLSM volumes were imported into Amira (FEI) where threshold-based segmentation and label analysis were used to calculate 3D volumes of each nuclei. Criteria for measuring nuclear volume for CGNs was simply defined as nuclei that contained no Atoh1-EGFP signal at the start of time lapse imaging. For differentiating GNPs, a cell must have been Atoh1-EGFP positive at the beginning of an imaging sequence and subsequently lost Atoh1-EGFP expression without an apparent cell division that would reduce nuclear size by the completion of mitosis (Atoh1 protein half-life is ~30 min). Many GNPs maintained Atoh1-EGFP expression during 12-hour imaging runs showing the timing of GNP differentiation was not homogeneous across the progenitor-cell population and loss of EGFP signal was not photobleaching related (data not shown). As displayed in Figure 7A, B, GNP nuclei rapidly condense in an approximately two-hour period coinciding with the loss of Atoh1-EGFP signal from the nucleus of these cells. The optics used for the experiments described herein included the following: 560-, 640- and 488-nm laser lines, with maximum power of 500 mW and 300 mW, respectively, and a quad-band emission filter to resolve spectrally the imaged channels.

All the cell samples were imaged in Fluorobrite low-fluorescence medium supplemented with 10% heat-inactivated horse serum. The correction collar on the detection objective was adjusted to compensate for the index of refraction of the media and any spherical aberration. Cells were seeded on 5-mm coverslips and mounted in custom-fabricated sample holders for imaging. Images were acquired with the Slidebook software package, using a custom-designed 15 µm–square light-sheet pattern in the 5-phase structured illumination z-galvo and objective scan mode, with all colors being captured at each z position. The acquired images were background subtracted and SIM images were reconstructed using an open-source, GPU-accelerated SIM reconstruction software tailored to LLSM SIM data with freshly acquired optical transfer functions (OTFs) generated on each day of imaging (*125*).

#### *11. Cryo-SIM/FIB-SEM registration*

For cryo SR / FIB-SEM chromatin domain quantitation, a pipeline was developed that consisted of multiple image processing, registration, segmentation, and chromatin domain analysis steps (Fig. S19). First, raw SIM images were reconstructed using the following parameters: 0.007 Wiener Filter, 0.7 gamma apodization, 15 pixel radii of the singularity suppression at the OTF origins (*125*). FIB-SEM volumes were aligned using the SIFT-algorithm FIJI plugin (*93*) and resliced so X, Y, and Z coordinates match SIM images. Before light microscopy to electron microscopy registration, normalized mutual information was used to align SIM channel, and both SIM and FIB-SEM volumes were cropped down to single cells, as registering across large ROIs proved less useful than single-cell crops in the 3D Slicer. The merged SIM channels were then imported into the 3D Slicer software package along with its corresponding FIB-SEM stack. Next, 8-10 landmarks were selected between the SIM composite (moving) to FIB-SEM (fixed) prioritizing unique in-focus center of volume features. It is important to note, over-reliance of landmarks on SIM z-volume top or bottom were suboptimal for registering light microscopy information to FIB-SEM data. Once landmarks were placed, an affine transformation was applied to merged HP1α and H3.3 SIM volumes to the FIB-SEM data. If additional fine-scale registration was needed after landmark registration, a normalized mutual information-based algorithm was used to achieve optimal alignments (reserved for <25% of cell analyzed in this study). Final registered SIM volumes were resampled with nearest-neighbor interpolation to match the voxel number and dimensions of the FIB-SEM volume (Fig. S18).

#### *12. Cryo-SIM/FIB-SEM segmentation*

For cryo SR / FIB-SEM segmentation (see Fig. S19 for the pipeline), post-registration images were converted to HDF5 and loaded into Ilastik (*83*) in which pixel classification algorithm was used to generate a probability map. FIB-SEM segmentation masks for euchromatin, heterochromatin, and nucleoli were obtained using a two-stage classifier. SIM segmentation masks for each SIM channels, HP1α and H3.3, were obtained by single-stage classification. Finally, images were converted back to TIFFs for further processing and importation into MATLAB for quantification of volume overlaps, surface area to volume ratios, and heterochromatin volume fractions. All SIM volumes were interpolated to the FIB-SEM coordinate space. For visualization, TIFFs were Gaussian blurred (kernel sigma = 32 nm), and isosurfaces were rendered to reveal the precise subnuclear organization of each chromatin subdomain in a representative GNP and CGN (Fig. S22 and S23).

For cryoCLEM analyses quality assurance testing, we performed three tests. First, a SIM reconstruction parameter sweep was performed to generate a spectrum of final SIM reconstructed images (Fig. S18). These images were then passed through the cryoCLEM image analysis pipeline while all other parameters from the pipeline remained constant. It is important to note that a sweep of these parameters resulted in minor differences in final quantifications (Fig. S18A, boxplots). Second, the same strategy was implemented to explore the type of interpolation performed on the SIM dataset when resampled to FIB-SEM voxel dimensions, or vice versa (Fig. S18B, boxplots). Third, we explored how a shift in registration affected the final domain volumes (Fig. S20). This was done by zero-padding the SIM masks in positive and negative x, y, and z directions while keeping the FIB-SEM masks unaltered (Fig. S20, panel J).

#### *13. Biological Insights from Cryo-SR/FIB-SEM of GNP/CGN Nuclei*

Additional analyses revealed further 3D chromatin structural alterations that accompany neuronal differentiation in addition to the apparent differences in cryoCLEM-defined chromatin subdomain organization in GNP and CGN nuclei (Fig. 7). Surface area to volume ratio analysis of domains like HP1α-labeled heterochromatin, or unlabeled euchromatin (e.g., H3.3 free euchromatin) displayed differences linked to differentiation status. For example, both the cryoCLEM-defined HP1α-labeled heterochromatin and H3.3-labeled heterochromatin chromatin domains were not only more abundant nuclear fractions in the smaller CGN nuclei, but also the relatively lower surface area to volume ratios of these domains showed that HP1α- or H3.3-labeled material was composed of relatively larger objects in CGNs and therefore had undergone compaction during differentiation (Fig. S20A,B). While both GNPs and CGNs had a similar amounts of H3.3-loaded euchromatin indicative of transcriptionally active chromatin (Fig. 7), the surface area to volume ratio of this domain is smaller in CGNs indicating that it also compacts during GNP differentiation to CGNs (despite the fact that FIB-SEM ground-truth still shows that it is open euchromatin, Fig. S23B). In contrast, H3.3 free-euchromatin was more compacted in larger GNP nuclei showing that domain compaction was not merely a factor of the nuclear size difference between GNPs and CGNs (Fig. S20D).

We also performed a preliminary characterization of unlabeled euchromatin and H3.3-labeled heterochromatin using live-cell LLSM-SIM to test follow-on hypotheses related to initial conclusions drawn from nuclear cryoCLEM. First, we were intrigued by the H3.3-free euchromatin fraction, as this euchromatin domain makes up the most substantial fraction of GNP nuclei that differs compared to post-nuclear condensation CGNs. The absence of the H3.3 in this euchromatin fraction suggested that it was less likely to be transcriptionally active, and we therefore hypothesized that it might represent poised chromatin, a euchromatin domain that is epigenetically silenced awaiting developmental or other conditional signals to activate transcriptional activity (*126*). To test this hypothesis, we implemented the cMAP3 imaging probe developed by Delachat et al. (*84*) that harbors the chromobox domain of drosophila polycomb protein, which imparts H3K27me3 binding, and PHD domain of human TAF3, which imparts H3K4me3 binding, fused to fluorescent proteins like EGFP or Emerald (*84*). Given the relative binding affinities of the chromobox and PHD domains, cMAP3 preferentially binds to poised chromatin sites in the genome which harbor a high density of both the H3K27me3 transcriptionally inactive epigenetic mark and the H3K4me3 promoter/enhancer epigenetic mark (*84*). LLSM-SIM (Fig. S24A, movie S5) revealed 1) cMAP3 Emerald staining was more abundant in GNP nuclei, 2) aggregations of cMAP3 stain were located in large H3.3-free voids and 3) GNPs possessed larger cMAP3 aggregates than CGNs much like the higher compaction state of H3.3-free euchromatin from our cryoCLEM surface area to volume ratio analysis. This suggests that GNPs organize their poised chromatin in defined locations within their nuclei. Second, we used LLSM-SIM to image Emerald-TERF1, a telomere marker, and CenpA-Halo, a centromere marker, to further characterize the H3.3-Heterochromatin cyroCLEM defined domain (Fig. S24B). Telomeres and centromere were mostly located at the GNP and CGN nuclear periphery, as expected from a variety of earlier studies (*127*, *128*). However, telomere and centromere puncta were seldom embedded within H3.3 labeled regions suggesting that H3.3 recruitment to telomeres or centromeres is not a significant fraction of H3.3-labeled heterochromatin we detected via cryoCLEM. Also, H3.3-heterochromatin comprises roughly 20% of CGN nuclear volume, while telomeres and centromeres combined are known to occupy roughly of the 4% of genome in mouse cells (*129*).

Finally, we analyzed whether the presence or absence of HP1α or H3.3 in heterochromatin defined by cryoCLEM conferred gross nuclear morphology or ultrastructural differences to heterochromatin. Isosurface renders revealed that HP1α-heterochromatin (with or without co-incident H3.3) was generally located in large chromodomains that adorn the nuclear periphery, while heterochromatin labeled with only H3.3 was located in heterochromatin tendrils in the CGN nuclear interior (Fig. S22A). Intrigued by the apparent morphological and positional differences our isosurface renders revealed, we expanded on our heterochromatin analysis pipeline to include an analysis of grayscale values of the heterochromatin FIB-SEM domains with respect to localization of functional labels determined by super-resolution fluorescence data. First, heterochromatin-specific voxels (c.f. supplemental note 12) were ranked, and the median grayscale value was used as a threshold. Next, combinations of domains; such as HP1α alone-, H3.3 alone-, HP1α plus H3.3- or unlabeled-heterochromatin, were assigned by the coincidence or absence of the SIM channels (c.f. supplemental note 12) and the heterochromatin voxels. For every domain, we calculated the following ratio:

$$Volume\ Fraction = \frac{Total\ voxels\ of\ domain\ above\ median\ threshold}{Total\ voxels\ of\ specific\ domain}$$

Volume analysis was performed on the full 3D volumes of four individual CGNs totaling thousands of FIB-SEM image planes at 4-nm isotropic voxel size using custom scripts in MATLAB. The left panel in Figure S22B shows a FIB-SEM slice highlighting representative regions containing HP1α alone-, H3.3 alone-, HP1α plus H3.3- and unlabeled-heterochromatin that were defined by cryoCLEM. The HP1α alone-heterochromatin was the heterochromatin population with the most voxels harboring dense, heavy metal staining based on our volume fraction analysis (Fig. S22B, right panel). Interestingly, the presence of H3.3 in heterochromatin coincided with a significant increase in the number of voxels with low heavy metal staining, indicating a less densely packed heterochromatin configuration. Taken together, our isosurface renders and volume fraction analysis exposed morphological and ultrastructural variations in heterochromatin subpopulations not discernable by ultrastructure alone: 1) cryoSIM signals define heterochromatin populations displaying differential morphological and positional differences in CGN nuclei and 2) these morphological differences also extend to the ultrastructural level by delineating heterochromatin populations with different degrees of heavy metal staining and the number of open voxels without compacted material. The cryoCLEM method will be instrumental in future work to assay the role of nuclear proteins in functionally generating the chromatin configurations observed in this study. For example, an attractive downstream hypothesis from our the current cryoCLEM work is that H3.3 may not only be a marker for a more "open" form of heterochromatin, but also possibly play a functional role in the formation of this type heterochromatin.

#### *14. Cell Culture*

##### a) Fig. 1, 2, 3, S2, S3, S10, and S11

COS-7 (ATCC CRL-1651) and U2OS cells (ATCC HTB-96) were grown in phenol red-free Dulbecco's modified eagle Medium (DMEM, Corning 17-205-CV) supplemented with 10% (v/v) FBS (Gibco, 26140-079), 2 mM L-glutamine (Gibco 25030-081), 100 U/ml penicillin and 100 U/ml streptomycin (Gibco 15140-122) and incubated at 37° C with 5% $CO_2$ Transfections of approximately $7x10^5$ cells were performed using the SE cell line 96-well nucleofector kit (Lonza V4SC-1096) with programs CM 104 for U2OS and FF 104 for Cos-7 on the Amaxa Nucleofector II system (Lonza). All plasmid constructs were used at a concentration of 100 ng DNA/$7x10^5$ cells in U2OS and 300 ng DNA/$7x10^5$ cells in COS-7.

Janelia Fluor staining was done using 100 nM solutions for 30 minutes at 37° C. CellMask deep red plasma membrane Stain (Invitrogen C10046) was used according to manufacturer's instructions.

Sapphire coverslips (COE Optics, custom order) were coated with poly-l-lysine hydrobromide (PLL, Sigma P7890) dissolved in $H_2O$ at 200mg/mL and incubated for 15 minutes at room temperature (RT), followed by a 15 minute incubation with $H_2O$ containing 0.8 pM of 0.2 µm FluoSpheres (Invitrogen T7280) at RT. Coverslips were then sterilized with 70% ethanol before coating with 10 µg/mL human fibronectin (HFN, EMD Millipore Corp. FC010) for 30 minutes at 37° C. Cells were then seeded to reach 30-50% confluency at the time of HPF or fixation.

Glyoxal fixation for Fig. S2 was performed as previously described (*130*). Chemical fixation for Fig. S3 was done by treating cells with 4% (w/v) paraformaldehyde (PFA, Electron Microscopy Sciences 19208) and 0.1% glutaraldehyde (from 8% Glutaraldehyde, Electron Microscopy Sciences 111-30-8) in PHEM buffer (see below) for 10 minutes at RT, followed by quenching in 100 mM glycine (Sigma G7403) in phosphate buffered saline (PBS, Gibco 10010-023) for 5 minutes at RT.

**For 1 L PHEM Buffer add the following to 973 mL of Milliq water:**

- 25 ml HEPES Buffer (25 mM from a 1 M solution, Corning 25-060-CI)
- 2 mL $MgCl_2$ (2 mM, Sigma M1028)
- 3.8035 g EGTA (10 mM, Sigma E3885)
- 18.141 g PIPES (60 mM, Sigma P6757)

TOMM20 10aa linker Halo

```
ATGGTGGGTCGGAACAGCGCCATCGCCGCCGGTGTATGCGGGGCCCTTTTCATTGGGTACTGCA
TCTACTTCGACCGCAAAAGACGAAGTGACCCCAACTTCAAGAACAGGCTTCGAGAACGAAGAAA
GAAACAGAAGCTTGCCAAGGAGAGAGCTGGGCTTTCCAAGTTACCTGACCTTAAAGATGCTGAA
GCTGTTCAGAAGTTCTTCCTTGAAGAAATACAGCTTGGTGAAGAGTTACTAGCTCAAGGTGAAT
ATGAGAAGGGCGTAGACCATCTGACAAATGCAATTGCTGTGTGTGGACAGCCACAGCAGTTACT
GCAGGTCTTACAGCAAACTCTTCCACCACCAGTGTTCCAGATGCTTCTGACTAAGCTCCCAACA
ATTAGTCAGAGAATTGTAAGTGCTCAGAGCTTGGCTGAAGATGATGTGGAAGGCGGTAGCGGGG
ATCCACCGGTCGCCACCATGGCAGAAATCGGTACTGGCTTTCCATTCGACCCCCATTATGTGGA
AGTCCTGGGCGAGCGCATGCACTACGTCGATGTTGGTCCGCGCGATGGCACCCCTGTGCTGTTC
CTGCACGGTAACCCGACCTCCTCCTACGTGTGGCGCAACATCATCCCGCATGTTGCACCGACCC
ATCGCTGCATTGCTCCAGACCTGATCGGTATGGGCAAATCCGACAAACCAGACCTGGGTTATTT
CTTCGACGACCACGTCCGCTTCATGGATGCCTTCATCGAAGCCCTGGGTCTGGAAGAGGTCGTC
CTGGTCATTCACGACTGGGGCTCCGCTCTGGGTTTCCACTGGGCCAAGCGCAATCCAGAGCGCG
TCAAAGGTATTGCATTTATGGAGTTCATCCGCCCTATCCCGACCTGGGACGAATGGCCAGAATT
TGCCCGCGAGACCTTCCAGGCCTTCCGCACCACCGACGTCGGCCGCAAGCTGATCATCGATCAG
AACGTTTTTATCGAGGGTACGCTGCCGATGGGTGTCGTCCGCCCGCTGACTGAAGTCGAGATGG
ACCATTACCGCGAGCCGTTCCTGAATCCTGTTGACCGCGAGCCACTGTGGCGCTTCCCAAACGA
GCTGCCAATCGCCGGTGAGCCAGCGAACATCGTCGCGCTGGTCGAAGAATACATGGACTGGCTG
CACCAGTCCCCTGTCCCGAAGCTGCTGTTCTGGGGCACCCCAGGCGTTCTGATCCCACCGGCCG
AAGCCGCTCGCCTGGCCAAAAGCCTGCCTAACTGCAAGGCTGTGGACATCGGCCCGGGTCTGAA
TCTGCTGCAAGAAGACAACCCGGACCTGATCGGCAGCGAGATCGCGCGCTGGCTGTCGACGCTC
GAGATTTCCGGCTAA
```

TOMM20 10aa linker eGFP

ATGGTGGGTCGGAACAGCGCCATCGCCGCCGGTGTATGCGGGGCCCTTTTCATTGGGTACTGCA
TCTACTTCGACCGCAAAAGACGAAGTGACCCCAACTTCAAGAACAGGCTTCGAGAACGAAGAAA
GAAACAGAAGCTTGCCAAGGAGAGAGCTGGGCTTTCCAAGTTACCTGACCTTAAAGATGCTGAA
GCTGTTCAGAAGTTCTTCCTTGAAGAAATACAGCTTGGTGAAGAGTTACTAGCTCAAGGTGAAT
ATGAGAAGGGCGTAGACCATCTGACAAATGCAATTGCTGTGTGTGGACAGCCACAGCAGTTACT
GCAGGTCTTACAGCAAACTCTTCCACCACCAGTGTTCCAGATGCTTCTGACTAAGCTCCCAACA
ATTAGTCAGAGAATTGTAAGTGCTCAGAGCTTGGCTGAAGATGATGTGGAAGGCGGTAGCGGGG
ATCCACCGGTCGCCACCATGGTGAGCAAGGGCGAGGAGCTGTTCACCGGGGTGGTGCCCATCCT
GGTCGAGCTGGACGGCGACGTAAACGGCCACAAGTTCAGCGTGTCCGGCGAGGGCGAGGGCGAT
GCCACCTACGGCAAGCTGACCCTGAAGTTCATCTGCACCACCGGCAAGCTGCCCGTGCCCTGGC
CCACCCTCGTGACCACCCTGACCTACGGCGTGCAGTGCTTCAGCCGCTACCCCGACCACATGAA
GCAGCACGACTTCTTCAAGTCCGCCATGCCCGAAGGCTACGTCCAGGAGCGCACCATCTTCTTC
AAGGACGACGGCAACTACAAGACCCGCGCCGAGGTGAAGTTCGAGGGCGACACCCTGGTGAACC
GCATCGAGCTGAAGGGCATCGACTTCAAGGAGGACGGCAACATCCTGGGGCACAAGCTGGAGTA
CAACTACAACAGCCACAACGTCTATATCATGGCCGACAAGCAGAAGAACGGCATCAAGGTGAAC
TTCAAGATCCGCCACAACATCGAGGACGGCAGCGTGCAGCTCGCCGACCACTACCAGCAGAACA
CCCCCATCGGCGACGGCCCCGTGCTGCTGCCCGACAACCACTACCTGAGCACCCAGTCCGCCCT
GAGCAAAGACCCCAACGAGAAGCGCGATCACATGGTCCTGCTGGAGTTCGTGACCGCCGCCGGG
ATCACTCTCGGCATGGACGAGCTGTACAAGTAA

TOMM20 10aa linker mEmerald

ATGGTGGGTCGGAACAGCGCCATCGCCGCCGGTGTATGCGGGGCCCTTTTCATTGGGTACTGCA
TCTACTTCGACCGCAAAAGACGAAGTGACCCCAACTTCAAGAACAGGCTTCGAGAACGAAGAAA
GAAACAGAAGCTTGCCAAGGAGAGAGCTGGGCTTTCCAAGTTACCTGACCTTAAAGATGCTGAA
GCTGTTCAGAAGTTCTTCCTTGAAGAAATACAGCTTGGTGAAGAGTTACTAGCTCAAGGTGAAT
ATGAGAAGGGCGTAGACCATCTGACAAATGCAATTGCTGTGTGTGGACAGCCACAGCAGTTACT
GCAGGTCTTACAGCAAACTCTTCCACCACCAGTGTTCCAGATGCTTCTGACTAAGCTCCCAACA
ATTAGTCAGAGAATTGTAAGTGCTCAGAGCTTGGCTGAAGATGATGTGGAAGGCGGTAGCGGGG
ATCCACCGGTCGCCACCATGGTGAGCAAGGGCGAGGAGCTGTTCACCGGGGTGGTGCCCATCCT
GGTCGAGCTGGACGGCGACGTAAACGGCCACAAGTTCAGCGTGTCCGGCGAGGGCGAGGGCGAT
GCCACCTACGGCAAGCTGACCCTGAAGTTCATCTGCACCACCGGCAAGCTGCCCGTGCCCTGGC
CCACCCTCGTGACCACCTTGACCTACGGCGTGCAGTGCTTCGCCCGCTACCCCGACCACATGAA
GCAGCACGACTTCTTCAAGTCCGCCATGCCCGAAGGCTACGTCCAGGAGCGCACCATCTTCTTC
AAGGACGACGGCAACTACAAGACCCGCGCCGAGGTGAAGTTCGAGGGCGACACCCTGGTGAACC
GCATCGAGCTGAAGGGCATCGACTTCAAGGAGGACGGCAACATCCTGGGGCACAAGCTGGAGTA
CAACTACAACAGCCACAAGGTCTATATCACCGCCGACAAGCAGAAGAACGGCATCAAGGTGAAC
TTCAAGACCCGCCACAACATCGAGGACGGCAGCGTGCAGCTCGCCGACCACTACCAGCAGAACA
CCCCCATCGGCGACGGCCCCGTGCTGCTGCCCGACAACCACTACCTGAGCACCCAGTCCAAGCT
GAGCAAAGACCCCAACGAGAAGCGCGATCACATGGTCCTGCTGGAGTTCGTGACCGCCGCCGGG
ATCACTCTCGGCATGGACGAGCTGTACAAGTAA

TOMM20 10aa linker mTagYFP

ATGGTGGGTCGGAACAGCGCCATCGCCGCCGGTGTATGCGGGGCCCTTTTCATTGGGTACTGCA
TCTACTTCGACCGCAAAAGACGAAGTGACCCCAACTTCAAGAACAGGCTTCGAGAACGAAGAAA
GAAACAGAAGCTTGCCAAGGAGAGAGCTGGGCTTTCCAAGTTACCTGACCTTAAAGATGCTGAA
GCTGTTCAGAAGTTCTTCCTTGAAGAAATACAGCTTGGTGAAGAGTTACTAGCTCAAGGTGAAT
ATGAGAAGGGCGTAGACCATCTGACAAATGCAATTGCTGTGTGTGGACAGCCACAGCAGTTACT
GCAGGTCTTACAGCAAACTCTTCCACCACCAGTGTTCCAGATGCTTCTGACTAAGCTCCCAACA
ATTAGTCAGAGAATTGTAAGTGCTCAGAGCTTGGCTGAAGATGATGTGGAAGGCGGTAGCGGGG
ATCCACCGGTCGCCACCATGGTTAGCAAAGGCGAGGAGCTGTTCGCCGGCATCGTGCCCGTGCT
GATCGAGCTGGACGGCGACGTGCACGGCCACAAGTTCAGCGTGCGCGGCGAGGGCGAGGGCGAC
GCCGACTACGGCAAGCTGGAGATCAAGTTCATCTGCACCACCGGCAAGCTGCCCGTGCCCTGGC
CCACCCTGGTGACCACCCTCACCTACGGCGTACAGTGCTTCGCCCGCTACCCCAAGCACATGAA
GATGAACGACTTCTTCAAGAGCGCCATGCCCGAGGGCTACATCCAGGAGCGCACCATCCTCTTC
CAAGACGACGGCAAGTACAAGACCCGCGGCGAGGTGAAGTTCGAGGGCGACACCCTGGTGAACC
GCATCGAGCTGAAGGGCAAGGACTTCAAGGAGGACGGCAACATCCTGGGCCACAAGCTGGAGTA
CAGCTTCAACAGCCACAACGTCTACATCACCCCCGACAAGGCCAACAACGGCCTGGAGGTGAAC
TTCAAGACCCGCCACAACATCGAGGGCGGCGGCGTGCAGCTGGCCGACCACTACCAGACCAACG
TGCCCCTGGGCGACGGCCCCGTGCTGATCCCCATCAACCACTACCTGAGCTACCAGACCGACAT
CAGCAAGGACCGCAACGAGGCCCGCGACCACATGGTGCTCCTGGAGTCCGTCAGCGCCTGCAGC
CACACCCACGGCATGGACGAGCTGTACCGCTAA

TOMM20 10aa linker mCherry2

ATGGTGGGTCGGAACAGCGCCATCGCCGCCGGTGTATGCGGGGCCCTTTTCATTGGGTACTGCA
TCTACTTCGACCGCAAAAGACGAAGTGACCCCAACTTCAAGAACAGGCTTCGAGAACGAAGAAA
GAAACAGAAGCTTGCCAAGGAGAGAGCTGGGCTTTCCAAGTTACCTGACCTTAAAGATGCTGAA
GCTGTTCAGAAGTTCTTCCTTGAAGAAATACAGCTTGGTGAAGAGTTACTAGCTCAAGGTGAAT
ATGAGAAGGGCGTAGACCATCTGACAAATGCAATTGCTGTGTGTGGACAGCCACAGCAGTTACT
GCAGGTCTTACAGCAAACTCTTCCACCACCAGTGTTCCAGATGCTTCTGACTAAGCTCCCAACA
ATTAGTCAGAGAATTGTAAGTGCTCAGAGCTTGGCTGAAGATGATGTGGAAGGCGGTAGCGGGG
ATCCACCGGTCGCCACCATGGTGAGCAAGGGCGAGGAGGATAACATGGCCATCATCAAGGAGTT
CATGCGCTTCAAGGTGCACATGGAGGGCTCCGTGAACGGCCACGAGTTCGAGATCGAGGGCGAG
GGCGAGGGCCGCCCCTACGAGGGCACCCAGACCGCCAAGCTGAAGGTGACCAAGGGTGGCCCCC
TGCCCTTCGCCTGGGACATCCTGTCCCCTCAGTTCATGTACGGCTCCAAGGCCTACGTGAAGCA
CCCCGCCGACATCCCCGACTACTTGAAGCTGTCCTTCCCCGAGGGCTTCAATTGGGAGCGCGTG
ATGAACTTCGAGGACGGCGGCGTGGTGACCGTGACCCAGGACTCCTCCCTGCAGGACGGCGAGT
TCATCTACAAGGTGAAGCTGCGCGGCACCAACTTCCCCTCCGACGGCCCCGTAATGCAGTGTCG
TACCATGGGCTGGGAGGCCTCCACTGAGCGGATGTACCCCGAGGACGGCGCCCTGAAGGGCGAG
ATCAAGCAGAGGCTGAAGCTGAAGGACGGCGGCCACTACGACGCTGAGGTCAAGACCACCTACA
AGGCCAAGAAGCCCGTGCAGCTGCCCGGCGCCTACAACGTCGACATCAAGTTGGACATCCTTTC
CCACAACGAGGACTACACCATCGTGGAACAGTACGAACGCGCCGAGGGCCGCCACTCCACCGGC
GGCATGGACGAGCTGTACAAGTAA

mEmerald 21aa linker Sec61b

ATGGTGAGCAAGGGCGAGGAGCTGTTCACCGGGGTGGTGCCCATCCTGGTCGAGCTGGACGGCG
ACGTAAACGGCCACAAGTTCAGCGTGTCCGGCGAGGGCGAGGGCGATGCCACCTACGGCAAGCT
GACCCTGAAGTTCATCTGCACCACCGGCAAGCTGCCCGTGCCCTGGCCCACCCTCGTGACCACC

TTGACCTACGGCGTGCAGTGCTTCGCCCGCTACCCCGACCACATGAAGCAGCACGACTTCTTCA
AGTCCGCCATGCCCGAAGGCTACGTCCAGGAGCGCACCATCTTCTTCAAGGACGACGGCAACTA
CAAGACCCGCGCCGAGGTGAAGTTCGAGGGCGACACCCTGGTGAACCGCATCGAGCTGAAGGGC
ATCGACTTCAAGGAGGACGGCAACATCCTGGGGCACAAGCTGGAGTACAACTACAACAGCCACA
AGGTCTATATCACCGCCGACAAGCAGAAGAACGGCATCAAGGTGAACTTCAAGACCCGCCACAA
CATCGAGGACGGCAGCGTGCAGCTCGCCGACCACTACCAGCAGAACACCCCCATCGGCGACGGC
CCCGTGCTGCTGCCCGACAACCACTACCTGAGCACCCAGTCCAAGCTGAGCAAAGACCCCAACG
AGAAGCGCGATCACATGGTCCTGCTGGAGTTCGTGACCGCCGCCGGGATCACTCTCGGCATGGA
CGAGCTGTACAAGTAAATGTACAAGTCCGGACTCAGATCTGGCTCCAGCGCAGGCAGCGCATCC
GGCGGAAGCGGAAGCCCTGGTCCGACCCCCAGTGGCACTAACGTGGGATCCTCAGGGCGCTCTC
CCAGCAAAGCAGTGGCCGCCCGGGCGGCGGGATCCACTGTCCGGCAGAGGAAAAATGCCAGCTG
TGGGACAAGGAGTGCAGGCCGCACAACCTCGGCAGGCACCGGGGGGATGTGGCGATTCTACACA
GAAGATTCACCTGGGCTCAAAGTTGGCCCTGTTCCAGTATTGGTTATGAGTCTTCTGTTCATCG
CTTCTGTATTTATGTTGCACATTTGGGGCAAGTACACTCGTTCGTAG

Halo 21aa Linker Sec61b

ATGGCAGAAATCGGTACTGGCTTTCCATTCGACCCCCATTATGTGGAAGTCCTGGGCGAGCGCA
TGCACTACGTCGATGTTGGTCCGCGCGATGGCACCCCTGTGCTGTTCCTGCACGGTAACCCGAC
CTCCTCCTACGTGTGGCGCAACATCATCCCGCATGTTGCACCGACCCATCGCTGCATTGCTCCA
GACCTGATCGGTATGGGCAAATCCGACAAACCAGACCTGGGTTATTTCTTCGACGACCACGTCC
GCTTCATGGATGCCTTCATCGAAGCCCTGGGTCTGGAAGAGGTCGTCCTGGTCATTCACGACTG
GGGCTCCGCTCTGGGTTTCCACTGGGCCAAGCGCAATCCAGAGCGCGTCAAAGGTATTGCATTT
ATGGAGTTCATCCGCCCTATCCCGACCTGGGACGAATGGCCAGAATTTGCCCGCGAGACCTTCC
AGGCCTTCCGCACCACCGACGTCGGCCGCAAGCTGATCATCGATCAGAACGTTTTTATCGAGGG
TACGCTGCCGATGGGTGTCGTCCGCCCGCTGACTGAAGTCGAGATGGACCATTACCGCGAGCCG
TTCCTGAATCCTGTTGACCGCGAGCCACTGTGGCGCTTCCCAAACGAGCTGCCAATCGCCGGTG
AGCCAGCGAACATCGTCGCGCTGGTCGAAGAATACATGGACTGGCTGCACCAGTCCCCTGTCCC
GAAGCTGCTGTTCTGGGGCACCCCAGGCGTTCTGATCCCACCGGCCGAAGCCGCTCGCCTGGCC
AAAAGCCTGCCTAACTGCAAGGCTGTGGACATCGGCCCGGGTCTGAATCTGCTGCAAGAAGACA
ACCCGGACCTGATCGGCAGCGAGATCGCGCGCTGGCTGTCGACGCTCGAGATTTCCGGCATGTA
CAAGTCCGGACTCAGATCTGGCTCCAGCGCAGGCAGCGCATCCGGCGGAAGCGGAAGCCCTGGT
CCGACCCCCAGTGGCACTAACGTGGGATCCTCAGGGCGCTCTCCCAGCAAAGCAGTGGCCGCCC
GGGCGGCGGGATCCACTGTCCGGCAGAGGAAAAATGCCAGCTGTGGGACAAGGAGTGCAGGCCG
CACAACCTCGGCAGGCACCGGGGGGATGTGGCGATTCTACACAGAAGATTCACCTGGGCTCAAA
GTTGGCCCTGTTCCAGTATTGGTTATGAGTCTTCTGTTCATCGCTTCTGTATTTATGTTGCACA
TTTGGGGCAAGTACACTCGTTCGTAG

CLIP10f 18aa Linker Sec61b

ATGGACAAAGACTGCGAAATGAAGCGCACCACCCTGGATAGCCCTCTGGGCAAGCTGGAACTGT
CTGGGTGCGAACAGGGCCTGCACCGTATCATCTTCCTGGGCAAAGGAACATCTGCCGCCGACGC
CGTGGAAGTGCCTGCCCCAGCCGCCGTGCTGGGCGGACCAGAGCCACTGATCCAGGCCACCGCC
TGGCTCAACGCCTACTTTCACCAGCCTGAGGCCATCGAGGAGTTCCCTGTGCCAGCCCTGCACC
ACCCAGTGTTCCAGCAGGAGAGCTTTACCCGCCAGGTGCTGTGGAAACTGCTGAAAGTGGTGAA
GTTCGGAGAGGTCATCAGCGAGAGCCACCTGGCCGCCCTGGTGGGCAATCCCGCCGCCACCGCC
GCCGTGAACACCGCCCTGGACGGAAATCCCGTGCCCATTCTGATCCCCTGCCACCGGGTGGTGC

AGGGCGACAGCGACGTGGGGCCCTACCTGGGCGGGCTCGCCGTGAAAGAGTGGCTGCTGGCCCA
CGAGGGCCACAGACTGGGCAAGCCTGGGCTGGGTTCCGGACTCAGATCTGGCTCCAGCGCAGGC
AGCGCATCCGGCGGAAGCGGAAGCCCTGGTCCGACCCCCAGTGGCACTAACGTGGGATCCTCAG
GGCGCTCTCCCAGCAAAGCAGTGGCCGCCCGGGCGGCGGATCCACTGTCCGGCAGAGGAAAAA
TGCCAGCTGTGGGACAAGGAGTGCAGGCCGCACAACCTCGGCAGGCACCGGGGGGATGTGGCGA
TTCTACACAGAAGATTCACCTGGGCTCAAAGTTGGCCCTGTTCCAGTATTGGTTATGAGTCTTC
TGTTCATCGCTTCTGTATTTATGTTGCACATTTGGGGCAAGTACACTCGTTCG

SNAPf 18aa Linker Sec61b

ATGGACAAAGACTGCGAAATGAAGCGCACCACCCTGGATAGCCCTCTGGGCAAGCTGGAACTGT
CTGGGTGCGAACAGGGCCTGCACCGTATCATCTTCCTGGGCAAAGGAACATCTGCCGCCGACGC
CGTGGAAGTGCCTGCCCCAGCCGCCGTGCTGGGCGGACCAGAGCCACTGATGCAGGCCACCGCC
TGGCTCAACGCCTACTTTCACCAGCCTGAGGCCATCGAGGAGTTCCCTGTGCCAGCCCTGCACC
ACCCAGTGTTCCAGCAGGAGAGCTTTACCCGCCAGGTGCTGTGGAAACTGCTGAAAGTGGTGAA
GTTCGGAGAGGTCATCAGCTACAGCCACCTGGCCGCCCTGGCCGGCAATCCCGCCGCCACCGCC
GCCGTGAAAACCGCCCTGAGCGGAAATCCCGTGCCCATTCTGATCCCCTGCCACCGGGTGGTGC
AGGGCGACCTGGACGTGGGGGGCTACGAGGGCGGGCTCGCCGTGAAAGAGTGGCTGCTGGCCCA
CGAGGGCCACAGACTGGGCAAGCCTGGGCTGGGTTCCGGACTCAGATCTGGCTCCAGCGCAGGC
AGCGCATCCGGCGGAAGCGGAAGCCCTGGTCCGACCCCCAGTGGCACTAACGTGGGATCCTCAG
GGCGCTCTCCCAGCAAAGCAGTGGCCGCCCGGGCGGCGGATCCACTGTCCGGCAGAGGAAAAA
TGCCAGCTGTGGGACAAGGAGTGCAGGCCGCACAACCTCGGCAGGCACCGGGGGGATGTGGCGA
TTCTACACAGAAGATTCACCTGGGCTCAAAGTTGGCCCTGTTCCAGTATTGGTTATGAGTCTTC
TGTTCATCGCTTCTGTATTTATGTTGCACATTTGGGGCAAGTACACTCGTTCGTAGG

ER3 3aa linker mEmerald

ATGCTGCTATCCGTGCCGTTGCTGCTCGGCCTCCTCGGCCTGGCCGTCGCCGACCGGTCGATGG
TGAGCAAGGGCGAGGAGCTGTTCACCGGGGTGGTGCCCATCCTGGTCGAGCTGGACGGCGACGT
AAACGGCCACAAGTTCAGCGTGTCCGGCGAGGGCGAGGGCGATGCCACCTACGGCAAGCTGACC
CTGAAGTTCATCTGCACCACCGGCAAGCTGCCCGTGCCCTGGCCCACCCTCGTGACCACCTTGA
CCTACGGCGTGCAGTGCTTCGCCCGCTACCCCGACCACATGAAGCAGCACGACTTCTTCAAGTC
CGCCATGCCCGAAGGCTACGTCCAGGAGCGCACCATCTTCTTCAAGGACGACGGCAACTACAAG
ACCCGCGCCGAGGTGAAGTTCGAGGGCGACACCCTGGTGAACCGCATCGAGCTGAAGGGCATCG
ACTTCAAGGAGGACGGCAACATCCTGGGGCACAAGCTGGAGTACAACTACAACAGCCACAAGGT
CTATATCACCGCCGACAAGCAGAAGAACGGCATCAAGGTGAACTTCAAGACCCGCCACAACATC
GAGGACGGCAGCGTGCAGCTCGCCGACCACTACCAGCAGAACACCCCCATCGGCGACGGCCCCG
TGCTGCTGCCCGACAACCACTACCTGAGCACCCAGTCCAAGCTGAGCAAAGACCCCAACGAGAA
GCGCGATCACATGGTCCTGCTGGAGTTCGTGACCGCCGCCGGGATCACTCTCGGCATGGACGAG
CTGTACAAGTAA

**b) Fig. 4**

HeLa cells (ATCC CCL-2) cultured in complete EMEM (ATCC 30-2003; 10% FBS, Gibco 26140-079; 1% Penicillin/Streptomycin, Gibco 15140-122) were plated onto fibronectin coated sapphire coverslips in a 35 mm glass bottomed dish (MatTek Corporation P35G-1.5-20-C) 24 hours prior to transfection. Cells were transfected with 50 ng TOMM20-Halo-N10 (see above) and 50 ng mEmerald-SKL (see below) plasmids using Transit-LT1 (Mirus MIR 2304) for 18 h. The transfected cells were incubated with 100 nM JF525-HT for 30 minutes at 37° C.

mEmerald 2aa linker SKL

```
ATGGTGAGCAAGGGCGAGGAGCTGTTCACCGGGGTGGTGCCCATCCTGGTCGAGCTGGACGGCG
ACGTAAACGGCCACAAGTTCAGCGTGTCCGGCGAGGGCGAGGGCGATGCCACCTACGGCAAGCT
GACCCTGAAGTTCATCTGCACCACCGGCAAGCTGCCCGTGCCCTGGCCCACCCTCGTGACCACC
TTGACCTACGGCGTGCAGTGCTTCGCCCGCTACCCCGACCACATGAAGCAGCACGACTTCTTCA
AGTCCGCCATGCCCGAAGGCTACGTCCAGGAGCGCACCATCTTCTTCAAGGACGACGGCAACTA
CAAGACCCGCGCCGAGGTGAAGTTCGAGGGCGACACCCTGGTGAACCGCATCGAGCTGAAGGGC
ATCGACTTCAAGGAGGACGGCAACATCCTGGGGCACAAGCTGGAGTACAACTACAACAGCCACA
AGGTCTATATCACCGCCGACAAGCAGAAGAACGGCATCAAGGTGAACTTCAAGACCCGCCACAA
CATCGAGGACGGCAGCGTGCAGCTCGCCGACCACTACCAGCAGAACACCCCCATCGGCGACGGC
CCCGTGCTGCTGCCCGACAACCACTACCTGAGCACCCAGTCCAAGCTGAGCAAAGACCCCAACG
AGAAGCGCGATCACATGGTCCTGCTGGAGTTCGTGACCGCCGCCGGGATCACTCTCGGCATGGA
CGAGCTGTACAAGTCC GGA TCC AAG CTG TAG
```

**c) Fig. 5**

SUM159 human breast carcinoma cells were provided by J. Brugge; cells were grown in DMEM/F-12/GlutaMAX (Life Technologies, 10565-042), supplemented with 5% fetal bovine serum (FBS, Atlanta Biologicals, S11050), 100 U/ml penicillin and streptomycin (VWR International, 97063-708), 1 µg/ml hydrocortisone (Sigma-Aldrich, H4001), 5 µg/ml insulin (Sigma-Aldrich, I9278-5ML), and 10 mM HEPES (Corning, 25-060-CI), pH 7.4. Cells were maintained at 37 °C and 5% $CO_2$. All cells were routinely verified to be mycoplasma free using a PCR-based assay.

Coverslips were sterilized in 70% ethanol and placed under UV for at least 30 minutes before placing in 6-well TC plates. Matrigel (Corning, 354230) was thawed on ice and resuspended in chilled DPBS (Thermo Fisher, 14190144) with 2 mg Matrigel in 25 mL DPBS. 1 ml of diluted Matrigel was added to each well of 6-well TC plates.

Cells were seeded on coverslips pretreated with Matrigel the night before imaging. 10 µg/mL AF646 labeled transferrin (Thermo Fisher, T23366) were diluted in phenol red free culture medium (Thermo Fisher A1896702) and incubated with cells for 30 minutes at 37 °C followed by three washes with complete medium.

**d) Fig. 6**

CGNs were prepared according to established protocol (*131*). Cerebella were dissected from the brains of P7 C57BL/6 mice, the pia was stripped away, and cerebellar tissue treated with papain and triturated using a fine-bore fire-polished Pasteur pipettes to dissociate dissected cerebella. The resulting single-cell suspension in CMF-PBS was layered onto a 60% to 35% Percoll gradient and separated by centrifugation. The small cell fraction at the 35% to 65% Percoll interface was isolated and plated on poly-L-ornithine coated plastic dishes to pan away contaminating glia or fibroblasts. Pure cerebellar cells were nucleofected with up to 50 µg of pCIG2 expression vector combinations encoding fluorescence protein- or SNAP or Halo tag-labeled imaging probes using the Amaxa Mouse Neuron kit with the O-005 program on the Amaxa Nucleofector II system (Lonza). Neuronal adhesions were labeled by nucleofection of plasmids encoding Junctional Adhesion Molecule-C SNAP, Drebrin 2x-Venus and Clathrin light chain (LC) Halo, while neuronal nuclei were labeled by nucleofection of plasmids encoding Heterochromatin Protein 1 alpha and Histone 3.3. Undifferentiated GNPs were maintained via nucleofection of pCIG2 SmoM2 receptor that restrains GNP differentiation into CGNs. Nucleofected cells were plated in MatTek glass-bottomed movie dishes with 3 mm sapphire disks coated with poly-L-ornithine and laminin.

JAM-C pHluorin

```
ATGGCGCTGAGCCGGCGGCTGCGACTTCGACTGTACGCGCGGCTGCCTGACTTCTTCCTGCTGC
TGCTCTTCAGGGGCTGCATGATAGAGGCAGTGAATCTCAAATCCAGCAACCGAAACCCAGTGGT
ACATGAATTTGAAAGTGTGGAATTGTCTTGCATCATTACGGACTCACAGACAAGTGACCCTAGG
ATTGAATGGAAGAAAATCCAAGATGGCCAAACCACATATGTGTATTTTGACAACAAGATTCAAG
GAGACCTGGCAGGTCGCACAGATGTGTTTGGAAAAACTTCCCTGAGGATCTGGAATGTGACACG
ATCGGATTCAGCCATCTATCGCTGTGAGGTCGTTGCTCTAAATGACCGAAAAGAAGTTGATGAG
ATTACCATTGAGTTAATTGTGCAAGTGAAGCCAGTGACCCCTGTCTGCAGAATTCCAGCCGCTG
TACCTGTAGGCAAGACGGCAACACTGCAGTGCCAAGAGAGCGAGGGCTATCCCCGGCCTCACTA
CAGCTGGTACCGCAATGATGTGCCACTGCCTACAGATTCCAGAGCCAATCCCAGGTTCCAGAAT
TCCTCTTTCCATGTGAACTCGGAGACAGGCACTCTGGTTTTCAATGCTGTCCACAAGGATGACT
CTGGGCAGTACTACTGCATTGCTTCCAATGACGCAGGTGCAGCCAGGTGTGAGGGGCAGGACAT
GGAAGTCTATGATAGTAAAGGAGAAGAACTTTTCACTGGAGTTGTCCCAATTCTTGTTGAATTA
GATGGTGATGTTAATGGGCACAAATTTTCTGTCAGTGGAGAGGGTGAAGGTGATGCAACATACG
GAAAACTTACCCTTAAATTTATTTGCACTACTGGAAAACTACCTGTTCCATGGCCAACACTTGT
CACTACTTTAACTTATGGTGTTCAATGCTTTTCAAGATACCCAGATCATATGAAACGGCATGAC
TTTTTCAAGAGTGCCATGCCCGAAGGTTATGTACAGGAAAGAACTATATTTTTCAAAGATGACG
GGAACTACAAGACACGTGCTGAAGTCAAGTTTGAAGGTGATACCCTTGTTAATAGAATCGAGTT
```

AAAAGGTATTGATTTTAAAGAAGATGGAAACATTCTTGGACACAAATTGGAATACAACTATAAC
GATCACCAGGTGTACATCATGGCAGACAAACAAAAGAATGGAATCAAAGCTAACTTCAAAATTA
GACACAACATTGAAGATGGAGGCGTTCAACTAGCAGACCATTATCAACAAAATACTCCAATTGG
CGATGGGCCCGTCCTTTTACCAGACAACCATTACCTGTTTACAACTTCTACTCTTTCGAAAGAT
CCCAACGAAAAGAGAGACCACATGGTCCTTCTTGAGTTTGTAACAGCTGCTGGGATTACACATG
GCATGGATGAACTATACAAAGAGCTCTTGAACATTGCTGGGATTATTGGGGGAGTCCTTGTTGT
CCTTATTGTTCTTGCTGTGATTACGATGGGCATCTGCTGTGCGTACAGACGAGGCTGCTTCATC
AGCAGTAAACAAGATGGAGAAAGCTATAAGAGCCCAGGGAAGCATGACGGTGTTAACTACATCC
GGACGAGTGAGGAGGGTGACTTCAGACACAAATCGTCCTTTGTTATCTGA

Drebrin Glycine Linker 2x Venus (dimer of Venus yfp)

ATGGCCGGCGTCAGCTTCAGCGGCCACCGCCTGGAGCTGCTGGCGGCGTACGAGGAGGTGATCC
GGGAGGAGAGCGCAGCCGACTGGGCTCTGTACACATACGAGGATGGCTCAGATGACCTCAAGCT
TGCAGCGTCAGGAGAAGGGGGCTTGCAGGAGCTTTCCGGCCACTTCGAGAACCAGAAAGTGATG
TATGGTTTCTGCAGCGTCAAGGACTCCCAAGCTGCCCTGCCAAAATATGTGCTCATCAACTGGG
TTGGTGAGGATGTGCCTGATGCCCGAAAATGTGCTTGCGCCAGTCATGTGGCCAAGGTGGCTGA
GTTCTTCCAGGGTGTTGATGTCATTGTGAATGCCAGCAGTGTGGAAGACATCGATGCTGGTGCC
ATTGGGCAGCGGCTCTCCAATGGACTGGCACGGCTCTCCAGCCCAGTATTGCACCGCCTGCGCC
TTCGGGAGGATGAAAATGCTGAACCGGTGGGTACCACCTACCAGAAGACGGATGCAGCAGTGGA
GATGAAGCGGATTAACCGTGAGCAGTTTTGGGAGCAGGCCAAGAAGGAGGAAGAGCTGCGGAAG
GAGGAGGAGCGGAAGAAGGCTCTGGACGCCAGGCTCAGGTTTGAACAGGAACGGATGGAGCAGG
AGCGGCAGGAGCAGGAAGAACGTGAGCGGCGCTACCGGGAGCGGGAGCAGCAGATTGAGGAGCA
CAGGAGGAAACAGCAGAGTCTGGAAGCTGAAGAAGCCAAGAGGAGGTTAAAGGAGCAGTCTATC
TTTGGTGACCAGCGGGATGAAGAGGAAGAGTCCCAGATGAAGAAGTCGGAGTCAGAGGTGGAGG
AGGCGGCTGCCATCATTGCCCAGCGGCCTGATAACCCACGGGAGTTCTTCAGACAGCAGGAACG
AGTGGCATCGGCCTCTGGTGGCAGCTGTGACGCGCCTGCGCCTGCACCCTTCAACCACCGACCA
GGTCGTCCGTACTGCCCTTTCATAAAGGCATCGGACAGTGGGCCTTCCTCCTCCTCCTCTTCCT
CCTCTTCCCCTCCACGGACTCCCTTTCCCTATATCACCTGCCACCGCACCCCAAACCTCTCTTC
CTCCCTCCCATGCAGCCACCTGGACAGCCACCGGAGGATGGCACCCACTCCTATTCCCACCCGG
AGCCCATCTGATTCCAGCACAGCCTCTACCCCCATCGCTGAGCAGATCGAGAGGGCCCTGGATG
AGGTCACATCCTCGCAGCCTCCACCTCCACCTCCACCACCTCCACCAACTCAAGAGGCCCAGGA
GACTACCCCAAGCCTGGATGAAGAGCTCAGCAAGGAGGCCAAAGTAACAGCAGCTCCTGAGGTC
TGGGCTGGCTGTGCGGCAGAGCCCCCTCAGGCACAGGAACCTCCCCTGTTGCAAAGCAGCCCCC
TGGAGGACTCGATGTGCACAGAATCTCCAGAGCAGGCTGCCCTGGCTGCCCCTGCGGAGCCTGC
TGCCTCTGTCACCTCAGTAGCTGATGTCCATGCAGCTGACACCATTGAGACCACCACTGCCACT
ACTGACACCACTATTGCCAACAACGTCACCCCTGCCGCTGCCAGCCTCATTGATCTATGGCCTG
GCAACGGGGAAGAGGCCTCAACACTTCAGGCTGAACCCAGGGTGCCCACACCACCCTCAGGTGC
TGAGGCCTCCCTGGCAGAGGTGCCCCTGCTGAATGAGGCCGCTCAGGAGCCGCTGCCGCCGGTA
GGCGAAGGCTGTGCTAACCTTCTTAATTTTGATGAGCTGCCAGAACCTCCAGCCACCTTCTGTG
ACCCAGAGGAGGAAGTAGGAGAAACGCTGGCTGCCTCCCAGGTCCTAACTATGCCCTCAGCTCT
AGAGGAGGTAGATCAGGTGCTGGAGCAGGAGCTGGAGCCAGAACCTCACCTGCTGACCAATGGA
GAGACCACTCAAAAGGAGGGGACCCAGGCCAGCGAAGGATACTTCAGTCAGTCACAGGAGGAAG
AGTTCGCCCAATCAGAAGAGCCATGTGCAAAGGTTCCGCCTCCTGTATTTTACAACAAGCCTCC
AGAAATCGACATCACCTGCTGGGATGCAGACCCAGTTCCTGAAGAGGAAGAGGGCTTCGAGGGT
GGTGATAGCGGCGGCGGGAGCATGGTGAGCAAGGGCGAGGAGCTGTTCACCGGGGTGGTGCCCA

TCCTGGTCGAGCTGGACGGCGACGTAAACGGCCACAAGTTCAGCGTGTCCGGCGAGGGCGAGGG
CGATGCCACCTACGGCAAGCTGACCCTGAAGCTGATCTGCACCACCGGCAAGCTGCCCGTGCCC
TGGCCCACCCTCGTGACCACCCTGGGCTACGGCCTGCAGTGCTTCGCCCGCTACCCCGACCACA
TGAAGCAGCACGACTTCTTCAAGTCCGCCATGCCCGAAGGCTACGTCCAGGAGCGCACCATCTT
CTTCAAGGACGACGGCAACTACAAGACCCGCGCCGAGGTGAAGTTCGAGGGCGACACCCTGGTG
AACCGCATCGAGCTGAAGGGCATCGACTTCAAGGAGGACGGCAACATCCTGGGGCACAAGCTGG
AGTACAACTACAACAGCCACAACGTCTATATCACCGCCGACAAGCAGAAGAACGGCATCAAGGC
CAACTTCAAGATCCGCCACAACATCGAGGACGGCGGCGTGCAGCTCGCCGACCACTACCAGCAG
AACACCCCCATCGGCGACGGCCCCGTGCTGCTGCCCGACAACCACTACCTGAGCTACCAGTCCG
CCCTGAGCAAAGACCCCAACGAGAAGCGCGATCACATGGTCCTGCTGGAGTTCGTGACCGCCGC
CGGGATCACTCTCGGCATGGACGAGCTGTACAAGATGGTGAGCAAGGGCGAGGAGCTGTTCACC
GGGGTGGTGCCCATCCTGGTCGAGCTGGACGGCGACGTAAACGGCCACAAGTTCAGCGTGTCCG
GCGAGGGCGAGGGCGATGCCACCTACGGCAAGCTGACCCTGAAGCTGATCTGCACCACCGGCAA
GCTGCCCGTGCCCTGGCCCACCCTCGTGACCACCCTGGGCTACGGCCTGCAGTGCTTCGCCCGC
TACCCCGACCACATGAAGCAGCACGACTTCTTCAAGTCCGCCATGCCCGAAGGCTACGTCCAGG
AGCGCACCATCTTCTTCAAGGACGACGGCAACTACAAGACCCGCGCCGAGGTGAAGTTCGAGGG
CGACACCCTGGTGAACCGCATCGAGCTGAAGGGCATCGACTTCAAGGAGGACGGCAACATCCTG
GGGCACAAGCTGGAGTACAACTACAACAGCCACAACGTCTATATCACCGCCGACAAGCAGAAGA
ACGGCATCAAGGCCAACTTCAAGATCCGCCACAACATCGAGGACGGCGGCGTGCAGCTCGCCGA
CCACTACCAGCAGAACACCCCCATCGGCGACGGCCCCGTGCTGCTGCCCGACAACCACTACCTG
AGCTACCAGTCCGCCCTGAGCAAAGACCCCAACGAGAAGCGCGATCACATGGTCCTGCTGGAGT
TCGTGACCGCCGCCGGGATCACTCTCGGCATGGACGAGCTGTACAAGTAA

Halo 15aa linker Clathrin LC

ATGCCTACCCTCTATAAAAAGGCTGGAAGTACTATGGCCGAAATTGGTACTGGTTTTCCCTTTG
ATCCTCACTATGTCGAGGTACTTGGCGAGCGAATGCACTATGTCGACGTTGGTCCACGCGACGG
AACACCAGTGCTCTTTCTTCATGGAAACCCTACTTCTTCTTATGTATGGCGGAATATCATACCT
CATGTAGCACCAACCCACCGCTGCATAGCACCAGACCTGATTGGGATGGGCAAGAGCGACAAAC
CTGATTTGGGGTACTTCTTCGACGACCATGTTCGCTTCATGGATGCCTTTATTGAAGCACTCGG
CCTGGAAGAGGTCGTCCTCGTCATTCATGACTGGGGGTCTGCACTCGGCTTTCATTGGGCAAAG
AGAAACCCCGAACGGGTTAAAGGAATAGCCTTCATGGAATTTATACGGCCTATACCAACTTGGG
ACGAATGGCCCGAATTTGCTCGGGAAACTTTCCAGGCATTCCGCACAACAGATGTAGGAAGAAA
ACTTATCATCGATCAGAATGTTTTTATAGAAGGCACATTGCCAATGGGAGTAGTAAGGCCACTT
ACCGAGGTAGAGATGGACCATTATAGAGAACCCTTTTTGAATCCAGTCGACCGAGAACCACTGT
GGAGGTTCCCTAACGAACTTCCTATCGCCGGCGAGCCAGCTAACATTGTCGCTTTGGTAGAGGA
ATACATGGACTGGCTGCATCAGTCACCAGTCCCAAAACTGTTGTTCTGGGGCACCCCAGGGGTA
CTTATCCCACCCGCCGAAGCAGCTAGGCTCGCCAAGTCCCTCCCAAATTGCAAAGCCGTGGACA
TCGGTCCCGGCCTCAACCTTCTCCAGGAAGATAACCCAGACTTGATAGGGTCAGAGATCGCCCG
CTGGTTGAGTACTCTCGAAATCTCAGGAGAACCCACTACTGAGGACCTCTATTTCCAATCCGAC
AATGCCAATTCTGTGGACTGGATACGATACCGAATCCGGGGTCGCACAAGGGACATCTCCGGAC
TCAGATCTCGGGCTCAAGCTTCGAACTCTGCAGTCGACATGGCTGATGACTTTGGCTTCTTCTC
GTCGTCGGAGAGTGGTGCCCCGGAGGCGGCGGAGGAGGACCCGGCAGCCGCCTTCCTGGCCCAG
CAGGAGAGCGAGATTGCAGGCATAGAGAACGACGAGGGCTTCGGGGCACCTGCCGGCAGCCATG
CGGCCCCCGCACAGCCGGGCCCCACGAGTGGGGCTGGTTCTGAGGACATGGGGACCACAGTCAA
TGGAGATGTGTTTCAGGAGGCCAACGGTCCTGCTGATGGCTACGCAGCCATTGCCCAGGCTGAC
AGGCTGACCCAGGAGCCTGAGAGCATCCGCAAGTGGCGAGAGGAGCAGAGGAAACGGCTGCAAG

```
AGCTGGATGCTGCATCTAAGGTCACGGAACAGGAATGGCGGGAGAAGGCCAAGAAGGACCTGGA
GGAGTGGAACCAGCGCCAGAGTGAACAAGTAGAGAAGAACAAGATCAACAACCGGGCATCCGAG
GAGGCTTTCGTGAAGGAATCCAAGGAGGAGACCCCAGGCACAGAGTGGGAGAAGGTGGCCCAGC
TATGTGACTTCAACCCCAAGAGCAGCAAGCAGTGCAAAGATGTGTCCCGCCTGCGCTCGGTGCT
CATGTCCCTGAAGCAGACGCCACTGTCCCGCTAA
```

**e) Fig. 7**

We established the developmental transition of GNPs into CGNs as a novel model system to study rearrangements of nuclear chromatin domains linked to differentiation of neuronal-lineage cells. CGNs were prepared as described previously (*132*). Briefly, cerebella were dissected from the brains of postnatal day 7 (P7) Atoh1-EGFP transgenic B6 mice and the pial layer removed. The tissue was treated with a Neural Tissue Dissociation Kit (Miltenyi Biotec) and triturated into a single-cell suspension by using fine-bore Pasteur pipettes. The suspension was layered onto a discontinuous (35/60%) Percoll gradient and separated by centrifugation. The small-cell fraction was then isolated (95% GNPs and CGNs) and then sorted based on Atoh1-EGFP expression, a transcription factor for GNPs, on a BD FACS Aria Fusion cell sorter (BD Biosciences). Post-sorted cultures routinely contained roughly >95% GNPs from the Atoh1-EGFP high population and >95% differentiated CGNs from the Atoh1-EGFP low population. Expression vectors encoding fluorescently labeled nuclear proteins of interest were introduced into sorted cells via Amaxa nucleofection using an Amaxa Mouse Neuron Nucleofector Kit in accordance with the manufacturer's instructions and program (O-005). After cells had been allowed to recover from the nucleofection for 10 min, each cohort, Atoh1-EGFP high and low, were plated on 5 mm, NaOH-etched coverslips (No. 1) in 6-cm MatTek dishes precoated with low concentrations of poly-L-ornithine and then laminin (3 μg/cm$^3$) to facilitate neuronal attachment. Cells were incubated for 24 h before coverslips were mounted for lattice light sheet microscopy (LLSM).

For CryoCLEM, CGNs were prepared as described above. Briefly, cerebella were dissected from the brains of P7 C57Bl6 mice and the pial layer removed, and then the tissue was treated with a Neural Tissue Dissociation Kit (Miltenyi Biotec) and triturated into a single-cell suspension by using fine-bore Pasteur pipettes. The suspension was layered onto a discontinuous (35/60%) Percoll gradient and separated by centrifugation. The small-cell fraction was then isolated. Expression vectors encoding fluorescently labeled nuclear proteins and pCIG2 expressing proteins of interest were introduced into granule neurons via Amaxa nucleofection, using an Amaxa Mouse Neuron Nucleofector Kit following the manufacturer's instructions and program O-005. After cells had been allowed to recover from the nucleofection for 10 min, cells were plated on 3 mm, NaOH-etched sapphire coverslips in 6-cm MatTek dishes precoated with low concentrations of poly-L-ornithine and laminin to facilitate neuronal attachment. Cells were incubated for 24 h before coverslips were high-pressure frozen.

HP1α 10aa linker Emerald

```
ATGGGAAAGAAAACCAAGCGGACAGCTGACAGTTCTTCTTCAGAGGATGAGGAGGAGTATGTTG
TGGAGAAGGTGCTAGACAGGCGCGTGGTTAAGGGACAAGTGGAATATCTACTGAAGTGGAAAGG
CTTTTCTGAGGAGCACAATACTTGGGAACCTGAGAAAAACTTGGATTGCCCTGAGCTAATTTCT
GAATTTATGAAAAAGTATAAGAAGATGAAGGAGGGTGAAAATAATAAACCCAGGGAGAAGTCAG
AAAGTAACAAGAGGAAATCCAATTTCTCAAACAGTGCCGATGACATCAAATCTAAAAAAAAGAG
AGAGCAGAGCAATGATATCGCTCGGGGCTTTGAGAGAGGACTGGAACCAGAAAAGATCATTGGG
GCAACAGATTCCTGTGGTGATTTAATGTTCCTAATGAAATGGAAAGACACAGATGAAGCTGACC
TGGTTCTTGCAAAAGAAGCTAATGTGAAATGTCCACAAATTGTGATAGCATTTTATGAAGAGAG
ACTGACATGGCATGCATATCCTGAGGATGCGGAAAACAAAGAGAAAGAAACAGCAAAGAGCTCG
GGAAGCACGGATCCACCGGTCGCCACCATGGTGAGCAAGGGCGAGGAGCTGTTCACCGGGGTGG
TGCCCATCCTGGTCGAGCTGGACGGCGACGTAAACGGCCACAAGTTCAGCGTGTCCGGCGAGGG
CGAGGGCGATGCCACCTACGGCAAGCTGACCCTGAAGTTCATCTGCACCACCGGCAAGCTGCCC
GTGCCCTGGCCCACCCTCGTGACCACCTTGACCTACGGCGTGCAGTGCTTCGCCCGCTACCCCG
ACCACATGAAGCAGCACGACTTCTTCAAGTCCGCCATGCCCGAAGGCTACGTCCAGGAGCGCAC
CATCTTCTTCAAGGACGACGGCAACTACAAGACCCGCGCCGAGGTGAAGTTCGAGGGCGACACC
CTGGTGAACCGCATCGAGCTGAAGGGCATCGACTTCAAGGAGGACGGCAACATCCTGGGGCACA
AGCTGGAGTACAACTACAACAGCCACAAGGTCTATATCACCGCCGACAAGCAGAAGAACGGCAT
CAAGGTGAACTTCAAGACCCGCCACAACATCGAGGACGGCAGCGTGCAGCTCGCCGACCACTAC
CAGCAGAACACCCCCATCGGCGACGGCCCCGTGCTGCTGCCCGACAACCACTACCTGAGCACCC
AGTCCAAGCTGAGCAAAGACCCCAACGAGAAGCGCGATCACATGGTCCTGCTGGAGTTCGTGAC
CGCCGCCGGGATCACTCTCGGCATGGACGAGCTGTACAAGTGA
```

H3.3 Glycine linker SNAPf

```
ATGGCTCGTACAAAGCAGACTGCCCGCAAATCCACCGGTGGTAAAGCACCCAGGAAACAACTGG
CTACAAAAGCCGCTCGCAAGAGTGCGCCCTCTACTGGAGGGGTGAAGAAACCTCATCGTTACAG
GCCTGGTACTGTGGCCCTCCGTGAAATCAGACGCTATCAGAAGTCCACTGAACTTCTGATCCGC
AAGCTCCCCTTTCAGCGTCTGGTGCGAGAAATTGCTCAGGACTTCAAAACAGATCTGCGCTTCC
AGAGTGCAGCTATTGGTGCTTTGCAGGAGGCAAGTGAGGCCTATCTGGTTGGCCTTTTTGAAGA
TACCAATCTGTGTGCTATCCATGCCAAACGTGTAACAATTATGCCAAAAGATATCCAGCTTGCA
CGCCGCATACGCGGAGAACGTGCCAGCGGCGGCGGGAGCATGGATAAGGATTGTGAAATGAAAC
GCACAACACTTGACAGCCCCCTTGGGAAACTTGAACTTTCTGGTTGTGAGCAAGGGCTCCACCG
AATCATATTTCTGGGCAAAGGAACATCTGCTGCCGACGCTGTAGAAGTACCTGCACCTGCCGCA
GTTTTGGGCGGCCCAGAACCCTTGATGCAAGCAACTGCATGGCTCAACGCCTATTTTCACCAGC
CCGAGGCCATTGAAGAATTTCCCGTTCCAGCCCTGCATCATCCCGTATTCCAGCAAGAATCATT
CACTCGGCAAGTTTTGTGGAAACTGCTTAAAGTTGTTAAGTTCGGGGAGGTAATCTCATACTCT
CATTTGGCTGCATTGGCCGGAAACCCCGCTGCAACAGCAGCTGTAAAGACCGCCCTCAGTGGGA
```

ACCCCGTCCCCATCCTGATCCCCTGTCATCGAGTAGTACAAGGTGATTTGGATGTCGGAGGCTA
CGAGGGTGGCCTCGCTGTAAAGGAATGGTTGCTCGCACATGAGGGCCACCGGCTTGGCAAGCCT
GGGCTTGGGTGA

cMAP3 Emerald (epitope tag, nls, Pc CBX domain, 5aa linker, Taf3 Phd domain)

ATGGACTACAAGGACGACGACAAGGACCCCAAGAAGAAGAGGAAGGTGATGAAGAAGCACCACC
ACCACCACCACGACAACGCCACCGACGACCCCGTGGACCTGGTGTACGCCGCCGAGAAGATCAT
CCAGAAGAGGGTGAAGAAGGGCGTGGTGGAGTACAGGGTGAAGTGGAAGGGCTGGAACCAGAGG
TACAACACCTGGGAGCCCGAGGTGAACATCCTGGACAGGAGGCTGATCGACATCTACGAGCAGA
CCAACAAGGGCGGCGGCGGCAGCATGGTGAGCAAGGGCGAGGAGCTGTTCACCGGCGTGGTGCC
CATCCTGGTGGAGCTGGACGGCGACGTGAACGGCCACAAGTTCAGCGTGAGCGGCGAGGGCGAG
GGCGACGCCACCTACGGCAAGCTGACCCTGAAGTTCATCTGCACCACCGGCAAGCTGCCCGTGC
CCTGGCCCACCCTGGTGACCACCCTGACCTACGGCGTGCAGTGCTTCGCCAGGTACCCCGACCA
CATGAAGCAGCACGACTTCTTCAAGAGCGCCATGCCCGAGGGCTACGTGCAGGAGAGGACCATC
TTCTTCAAGGACGACGGCAACTACAAGACCAGGGCCGAGGTGAAGTTCGAGGGCGACACCCTGG
TGAACAGGATCGAGCTGAAGGGCATCGACTTCAAGGAGGACGGCAACATCCTGGGCCACAAGCT
GGAGTACAACTACAACAGCCACAAGGTGTACATCACCGCCGACAAGCAGAAGAACGGCATCAAG
GTGAACTTCAAGACCAGGCACAACATCGAGGACGGCAGCGTGCAGCTGGCCGACCACTACCAGC
AGAACACCCCCATCGGCGACGGCCCCGTGCTGCTGCCCGACAACCACTACCTGAGCACCCAGAG
CAAGCTGAGCAAGGACCCCAACGAGAAGAGGGACCACATGGTGCTGCTGGAGTTCGTGACCGCC
GCCGGCATCACCCTGGGCATGGACGAGCTGTACAAGGGCGGCTACCCCGGCGGCAGCATGTACG
TGATCAGGGACGAGTGGGGCAACCAGATCTGGATCTGCCCCGGCTGCAACAAGCCCGACGACGG
CAGCCCCATGATCGGCTGCGACGACTGCGACGACTGGTACCACTGGCCCTGCGTGGGCATCATG
ACCGCCCCCCCCGAGGAGATGCAGTGGTTCTGCCCCAAGTGCGCCAACAAGGACCCCAAGAAGA
AGAGGAAGGTGTACCCCTACGACGTGCCCGACTACGCCTGA

Emerald 11aa linker TERF1

ATGGTATCCAAGGGGGAGGAACTCTTCACAGGTGTTGTGCCTATTTTGGTAGAGCTTGATGGAG
ACGTAAATGGACATAAATTCTCAGTGTCAGGTGAGGGTGAGGGCGATGCCACCTACGGAAAGTT
GACCCTCAAGTTCATTTGCACCACAGGGAAATTGCCAGTTCCTTGGCCAACTCTCGTAACAACT
TTGACCTACGGAGTACAGTGTTTTGCCCGCTACCCAGACCATATGAAGCAGCACGATTTTTTTA
AGTCCGCAATGCCAGAGGGCTACGTCCAAGAGCGAACTATTTTCTTCAAGGATGACGGCAATTA
CAAAACCCGCGCCGAGGTAAAATTCGAGGGTGATACATTGGTAAACAGGATAGAGTTGAAAGGT
ATCGATTTCAAAGAGGACGGTAACATTCTTGGTCATAAACTCGAATACAATTATAACTCTCATA
AGGTTTACATTACTGCCGATAAACAAAAAAACGGAATTAAAGTCAATTTCAAGACACGACACAA
TATAGAAGATGGTTCTGTACAGCTGGCTGACCACTATCAACAGAATACACCCATCGGCGATGGT
CCTGTCCTCCTGCCAGACAATCACTATTTGTCTACTCAAAGTAAATTGTCAAAAGATCCCAACG
AGAAAAGGGATCATATGGTGCTTTTGGAATTTGTCACCGCAGCCGGAATAACTTTGGGGATGGA
TGAATTGTACAAGGGGGGGGGGGGGTACCGGTCGCCGAGCTCACCATGGCGGAGACGGTCTCC
TCAGCGGCCCGGGACGCGCCGAGCCGTGAGGGCTGGACAGATTCGGATTCTCCAGAGCAGGAGG
AGGTGGGAGACGACGCGGAGCTGCTCCAGTGCCAGCTTCAGCTGGGGACCCCGAGAGAGATGGA
GAACGCGGAGCTTGTGGCTGAGGTGGAGGCCGTGGCTGCGGGCTGGATGCTCGACTTCCTCTGC
CTGTCTCTGTGCCGAGCCTTCCGTGACGGCCGCTCCGAGGACTTTCGTCGTACTCGTGACAGCG
CCGAGGCTATTATTCATGGACTACACAGACTTACAGCTTACCAATTGAAAACTGTGTATATATG
TCAGTTTTTGACAAGAGTTGCATCTGGAAAGGCCCTTGATGCACAGTTTGAAGTTGATGAGCGT

ATTACACCCTTGGAATCAGCCCTGATGATTTGGAACTCAATTGAAAAGGAACATGACAAACTGC
ATGACGAAATAAAGAATTTAATTAAAATTCAGGCTGTAGCTGTTTGTATGGAAATTGGCAGCTT
TAAGGAAGCAGAAGAAGTATTTGAAAGAATATTTGGTGATCCAGAATTTTACACGCCTTTAGAA
AGGAAGTTACTTAAGATAATCTCTCAGAAGGATGTGTTCCACTCCCTTTTCCAACACTTCAGCT
ATAGCTGCATGATGGAGAAAATTCAGAGTTATGTGGGTGATGTGTTAAGTGAAAAATCATCAAC
TTTTCTAATGAAGGCAGCAACAAAAGTAGTGGAAAATGAGAAAGCGAGGACACAAGCGTCTAAG
GATAGGCCAGATGCCACCAACACTGGAATGGACACTGAAGTTGGTTTGAATAAAGAGAAAAGTG
TTAATGGCCAGCAGTCTACAGAAACTGAACCCTTAGTGGATACAGTATCCTCAATAAGGTCTCA
CAAGAACGCCTTATCGCAGTTAAAACACAGACGTGCTCCATCAGATTTCAGTAGGAACGAAGCA
AGAACAGGAACTCTTCAGTGTGAAACAACGATGGAAAGGAACCGAAGAACCAGTGGAAGGAATA
GATTGTGTGTCTCAGAGAATCAGCCAGACACTGATGACAAAAGTGGACGCAGGAAAAGACAGAC
ATGGCTTTGGGAAGAAGACAGAATTTTGAAGTGTGGTGTAAAGAAATATGGAGAGGGAAATTGG
GCTAAAATACTATCCCATTATAAGTTCAACAACCGAACAAGTGTCATGTTAAAAGATAGATGGA
GAACAATGAAGAGACTGAAACTGATTAGCTGA

Halo 12aa Linker CenpA

ATGCCTACCCTCTATAAAAAGGCTGGAAGTACTATGGCCGAAATTGGTACTGGTTTTCCCTTTG
ATCCTCACTATGTCGAGGTACTTGGCGAGCGAATGCACTATGTCGACGTTGGTCCACGCGACGG
AACACCAGTGCTCTTTCTTCATGGAAACCCTACTTCTTCTTATGTATGGCGGAATATCATACCT
CATGTAGCACCAACCCACCGCTGCATAGCACCAGACCTGATTGGGATGGGCAAGAGCGACAAAC
CTGATTTGGGGTACTTCTTCGACGACCATGTTCGCTTCATGGATGCCTTTATTGAAGCACTCGG
CCTGGAAGAGGTCGTCCTCGTCATTCATGACTGGGGGTCTGCACTCGGCTTTCATTGGGCAAAG
AGAAACCCCGAACGGGTTAAAGGAATAGCCTTCATGGAATTTATACGGCCTATACCAACTTGGG
ACGAATGGCCCGAATTTGCTCGGGAAACTTTCCAGGCATTCCGCACAACAGATGTAGGAAGAAA
ACTTATCATCGATCAGAATGTTTTTATAGAAGGCACATTGCCAATGGGAGTAGTAAGGCCACTT
ACCGAGGTAGAGATGGACCATTATAGAGAACCCTTTTTGAATCCAGTCGACCGAGAACCACTGT
GGAGGTTCCCTAACGAACTTCCTATCGCCGGCGAGCCAGCTAACATTGTCGCTTTGGTAGAGGA
ATACATGGACTGGCTGCATCAGTCACCAGTCCCAAAACTGTTGTTCTGGGGCACCCCAGGGGTA
CTTATCCCACCCGCCGAAGCAGCTAGGCTCGCCAAGTCCCTCCCAAATTGCAAAGCCGTGGACA
TCGGTCCCGGCCTCAACCTTCTCCAGGAAGATAACCCAGACTTGATAGGGTCAGAGATCGCCCG
CTGGTTGAGTACTCTCGAAATCTCAGGAGAACCCACTACTGAGGACCTCTATTTCCAATCCGAC
AATGCCAATTCTGTGGACTGGATACGATATCGAATCCGGGGTCGCACAAGGGACATCTCGGGAA
GCACGGATGTACCGGTCGCCGAGCTCACCATGGGCCCGCGTCGCAAACCGCAGACCCCAAGGAG
GAGACCCTCCAGCCCGGCGCCTGGACCCTCGCGACAGAGCTCCAGTGTAGGCTCTCAGACACTG
CGCAGAAGACAGAAATTCATGTGGCTTAAGGAAATCAAGACCCTGCAGAAGAGCACAGACCTCT
TGTTCAGGAAGAAGCCTTTCAGCATGGTTGTTAGAGAAATATGTGAGAAGTTCAGCCGTGGTGT
GGATTTTTGGTGGCAAGCCCAGGCCTTGTTGGCCCTTCAGGAGGCAGCAGAAGCTTTCCTCATC
CACCTCTTTGAGGACGCCTACCTCCTCTCCTTACATGCTGGTCGGGTCACGCTTTTCCCCAAAG
ACATTCAGTTGACCAGGAGAATCCGAGGCTTCGAGGGCGGACTCCCCTAA

#### *15.SMLM Data Processing*

##### a) Constrained 2D Gaussian Fitting to extract 3D coordinates of the fluorescent events.

Single molecule x-y localization was performed using PeakSelector software written in IDL, as described in (*89*, *94*). It consisted of two steps: in the first step the approximate x-y positions of fluorescent events are determined using difference of gaussians method. In the next step constrained 2D Gaussian fitting, which allows for the determination of the z-coordinate concurrently with x-y localization (described below), is performed in 7x7 pixel windows centered around the x-y positions found in the first step.

To calibrate the constrained 2D Gaussian fitting the calibration data sets collected (cf. supplemental note 3.i)) are first analyzed using unconstrained 2D Gaussian fitting:

$$I(x,y) = a_0 + a_1 \cdot e^{-\frac{(x-x_0)^2}{2 \cdot w_x{}^2} - \frac{(y-y_0)^2}{2 \cdot w_y{}^2}} \tag{20}$$

where $a_0$, $a_1$, $x_0$, $y_0$, $w_x$, and $w_y$ are independent fitting parameters. The fitting was performed within the PeakSelector software using the MPFIT routine written for IDL [(*133*)]. We call this iteration 0 of the calibration. The values of $w_x$ and $w_y$ as functions of Z-position for eight typical fluorescent beads are shown in Fig. S34A. Fifth order polynomial fits of the combined data set, after correcting for different axial positions, are shown as solid lines. Constrained 2D Gaussian fitting is performed using these polynomial fits for the same fluorescent events:

$$I(x,y) = a_0 + a_1 \cdot e^{-\frac{(x-x_0)^2}{2 \cdot w_x(z_0)^2} - \frac{(y-y_0)^2}{2 \cdot w_y(z_0)^2}}$$

$$w_x(z_0) = \sum_{i=0}^{5} b_i \cdot z_0{}^i \tag{21}$$

$$w_y(z_0) = \sum_{i=0}^{5} c_i \cdot z_0{}^i$$

where $a_0$, $a_1$, $x_0$, $y_0$, and $z_0$ are independent fitting parameters with coefficients $\{b_i\}$ and $\{c_i\}$ determined at the previous step (iteration 0). We call this iteration 1 of the calibration. Because the convergence paths are different for the unconstrained and constrained fitting procedures, the values of $w_x$ and $w_y$ extracted in iteration 0 deviate from the values $w_x = \sum_{i=0}^{5} b_i \cdot z_0{}^i$ and $w_y = \sum_{i=0}^{5} c_i \cdot z_0{}^i$ determined from the same images in iteration 1 (cf. Fig. S34C and D). Therefore, using coefficients $\{b_i\}$ and $\{c_i\}$ from iteration 0 results in incorrect values of the Z coordinate. To overcome this problem, we perform another round of polynomial fitting, this time on the values of $w_x$ and $w_y$ determined at iteration 1 (cf. Fig. S34E). Even though the sets of coefficients $\{b_i\}$ and $\{c_i\}$ determined after iteration 0 and iteration 1 are slightly different, the convergence paths of constrained 2D Gaussian fitting using these two sets are close enough that the resulting values of $w_x$ and $w_y$ determined via constrained 2D Gaussian fittings are the same for coefficients $\{b_i\}$ and $\{c_i\}$ extracted after iteration 0 and iteration 1. However, the values of Z-coordinate are not the same, as shown in Fig. S34E, where the value of extracted Z-coordinate vs actual Z-position is plotted for one fiducial. Constrained 2D Gaussian fitting with coefficients $\{b_i\}$ and $\{c_i\}$ from iteration 0 (blue squares) results in systematic errors in the Z coordinate, while using the coefficients from iteration 1 (red squares) provides more accurate results (cf. Fig. S34E). To confirm that iteration 1 is sufficient, we performed two more iterations without noticeable improvement in the data quality for either.

**b) Drift correction**

i) Fiducial finding

Finding fiducials in the SMLM data is a crucial step in drift correction, slab alignment and alignment to the scaffold. Fiducials were either manually identified in PeakSelector or automatically identified using the following algorithm implemented in Python. First all localizations with an axial localization precision greater than 15 nm are removed. The cutoff of 15 nm was determined empirically and works well with most data sets though in some cases it was enlarged to 30 nm. A two-dimensional histogram of the filtered data in the lateral plane is calculated with bin sizes equivalent to the acquisition pixel size (0.13 μm), i.e. the histogram is generated with a magnification of unity. Peaks are found within the histogram using a Difference of Gaussians algorithm (as implemented in scikit-image). The peaks located in the 2D histogram give the location of the fiducials to within a single pixel (0.13 μm).

Once the initial fiducial locations have been found the fiducials need to be extracted from the SMLM data. First all fiducials within a given radius (capture radius) of the estimated location are collected. This initial data set is filtered so that only the highest amplitude localization in each frame is kept; afterwards the fiducial data set has at most a single point per frame. This process is repeated for all fiducial positions found in the previous step. Spurious fiducials can be filtered by looking at the total number of points per fiducial. One would expect fiducials to be present in every frame of the experiment; only data sets with points in at least 80% of the frames are kept.

ii) Drift calculation

Drift is estimated in an iterative procedure. First, fiducials are found in the data set as described above. All fiducials are immobilized in ice thus it is a good assumption that all fiducials will have the same relative position throughout the experiment. An estimate of the drift is calculated using the following equations:

$$\hat{x}_i = x_i - \langle x_i \rangle_f \tag{22}$$

where $\boldsymbol{x}_i$ are the coordinates of the $i^{th}$ fiducial and the $\langle\,\rangle_f$ denotes an average across all frames, and

$$\delta_f = \left. \sum_i^n \frac{\hat{x}_{i,f}}{\sigma_{i,f}^2} \middle/ \sum_i^n \frac{1}{\sigma_{i,f}^2} \right. \tag{23}$$

where $\boldsymbol{\delta}_f$ is the estimated drift in frame *f*, $\hat{\boldsymbol{x}}_{i,f}$ are the displacements from the mean position of the $i^{th}$ fiducial in the $f^{th}$ frame and $\boldsymbol{\sigma}_{i,f}^2$ are the estimated variances in the fiducial's position. In other words, the estimated drift is calculated by taking a weighted average of the fiducials' deviation from their mean position (eq. (22)) on a per frame basis (eq. (23)) where the weights of each coordinate are the square of the localization precision in each direction (eq.(23)). The drift is removed from data set and the procedure repeated on the drift corrected data. In subsequent iterations $\boldsymbol{\delta}_f$ is not the total drift but the residual drift which is added to the initial estimate. Iteration is halted when the root mean squared deviation of $\boldsymbol{\delta}_f$ is less than some tolerance, usually $10^{-6}$.

There are a few differences between the first iteration and the subsequent ones. In the first iteration all found fiducials are used to estimate the drift. In subsequent iterations all fiducials with less than 0.25 pixels (32 nm) of residual drift are kept. If there are less then 5 such fiducials, fiducials with less than 0.5 pixels (65 nm) of residual drift are retained such that there are at least 5 fiducials used for drift correction. Secondly, for the first round either the user defines a large capture radius or one is estimated from the data as the standard deviation of the brightest fiducial, while in subsequent iterations the capture radius is calculated as the larger of either three times the largest residual drift of the previously used fiducials or 0.5 pixels (62 nm). Finally, for data sets consisting of multiple slabs the drift of one slab can be used to initialize the procedure for the subsequent slab. Residual drifts for data taken with a 1000 mm and 500 mm cylindrical lens are show in Fig. S35; these represent the best possible resolution for our SMLM images.

**c) Slab alignment**

Thicker specimens require data to be recorded at multiple focal planes. These data are then drift corrected independently from one another as described above. However, the slabs are not aligned to one another or to the scaffold (cf. supplemental note 3.i)). Slab alignment can proceed in two ways: the slabs can be registered to each other and the resulting stack can be aligned to the scaffold or each slab can be aligned directly to the scaffold. The second method is attractive because it removes the need for the slabs to be adjacent to one another, thus if the experimenter is only interested in a few focal planes then only those planes need to be recorded. The first method is easier because it is more amenable to automation and the data collected in this manuscript all had overlapping slabs.

In some cases, slabs were aligned manually using PeakSelector by following a general procedure for registering two color channels (*134*), except in this case it was each slab as the "first color channel" and the scaffold as "second color channel".

In other cases, the slabs were aligned using an automatic procedure implemented in Python. First, fiducials are extracted from each slab as described above and their mean positions are determined using eq. (14). For each pair of adjacent slabs an initial guess of matching fiducials is made by finding all neighboring fiducials within a given radius (usually 10 pixels or 1.3 μm) using a k-dimensional tree algorithm (scipy). The resulting point clouds are input into a coherent point drift algorithm (CPD, (*135*)) that estimates the transform that registers the two point clouds while taking into account the potential for outliers. The collected transforms are propagated from the first slab onwards such that all slabs are aligned to a common frame of reference.

To align to the scaffold, fiducials are extracted from the aligned stack in the usual manner. Scaffold fiducials are found using a difference of gaussians algorithm on the axial MIP of the scaffold data set. Fiducials are fit throughout the stack with a gaussian PSF. The axial position ($z_0$) is found by fitting a one-dimensional gaussian to the fitted amplitude and the lateral position is estimated by linearly interpolating the lateral coordinates ($x_0$, $y_0$), which are functions of the axial position, to $z_0$. A few corresponding fiducials between the aligned SMLM stack and the scaffold are manually found by comparing the MIP of the scaffold and a histogram of the aligned SMLM stack. These manually correspondences are used to estimate an initial transform from which more correspondences are identified in a nearest neighbors fashion. The enlarged set of correspondences are input into the same CPD algorithm used earlier except the estimated transformation is affine instead of translation.

**d) Multiple Wavelength Channel Registration**

The reference scaffold used for slab alignment was acquired with 488 nm excitation and an emission filter set consisting of a 532 nm long pass (Semrock, 532RU) and a 633 nm short pass (Semrock, 633SP) filter. Under these conditions the emission of green and orange beads was recorded simultaneously into a single reference scaffold and thus multiple color channels were registered to the same reference. However, because the emission spectra of beads and fluorophores are not the same, we need to take into account the chromatic focal shift discussed in section 3.f).

In order to account for the chromatic shift in our system when registering the processed SMLM and SIM datasets for different fluorescent channels, we first recorded the spectra of fluorophores and beads under all excitation and emission conditions, as shown in Fig. S25A. We then calculated the weight-averaged values of the chromatic shift for each excitation/emission condition as shown in the table above plot in Fig S25B.

The registration of PALM data sets was done in the following way:

1. We registered each slab of mEmerald PALM data set (collected with $\lambda_{exc}$=488nm and 488RU+520/35 emission filters) to the reference data set (collected with $\lambda_{exc}$=488nm and 532RU+633SP emission filters). For this we used green fluorescent beads that are detectable under $\lambda_{exc}$=488nm with both 488RU+520/35 and 532RU+633SP emission filter sets.
2. We registered each slab of JF525 SMLM data set (collected with $\lambda_{exc}$=532nm and 532RU+633SP emission filters) to the reference data set (collected with $\lambda_{exc}$=488nm and 532RU+633SP emission filters). For this we used orange fluorescent beads that are detectable with 532RU+633SP emission filter set under both $\lambda_{exc}$=488nm and $\lambda_{exc}$=532nm.
3. We shifted the registered mEmerald data set by 82nm “down”. The amount of shift was calculated as shown below:

$$\Delta Z_{JF525-mEmerald} = \left( \Delta Z_{mEmerald,exc488}^{488RU+520/35} - \left( \Delta Z_{GB,exc488}^{488RU+520/35} - \Delta Z_{GB,exc488}^{532RU+633SP} \right) \right) - \left( \Delta Z_{JF525\ exc532}^{532RU+633SP} - \left( \Delta Z_{GB,exc488}^{532RU+633SP} - \Delta Z_{GB,exc532}^{532RU+633SP} \right) \right) \tag{24}$$

$$\Delta Z_{JF525-mEmerald} = \left( (-20.2) - ((-3.8) - 103.1) \right) - \left( 176.3 - (193.3 - 189.8) \right) = -82.1nm \tag{25}$$

#### e) Filtering

One issue with cryo-SMLM is interfering fluorescence from out-of-focus objects. A common source is emission from fiducials at a different focal plane. Another is detritus either embedded in the ice or stuck to the ice. In any case we would like to filter the localizations arising from these sources from our data and the final rendered images. Unfortunately, it is difficult to filter these contaminating signals during the peak extraction step, especially at the edges of the contamination. One could instead filter based on the extracted peak parameters such as localization precision or number of photons. However, the parameter thresholds are chosen arbitrarily, and the results are frequently unsatisfactory. To solve this problem, we have used a modern supervised classification algorithm: gradient boosted trees (as implemented in XGBoost 0.81). For each data set, we choose regions that are visually clean (“good” ROIs) and regions that have only contamination (“bad” ROIs) and use these as training data for the classification algorithm. The algorithm is trained based on the following features: # of photons, $\sigma_x$, $\sigma_y$, $\sigma_z$, offset, amplitude, $\chi^2$ and polynomial features of degree 2 of the previous list, i.e. $\sigma_x^2$, $\sigma_y\sigma_z$, offset × amplitude, etc.

Fig. S36 presents the results of three filtering algorithms on an exemplary data set. The first column of (A) shows the unfiltered data set and the next three columns show the data set after filtering with the following algorithms: classical thresholding, thresholding on a manually engineered data feature and gradient boosted trees. All data is presented after grouping and is shown on the same intensity scale. The second and third rows of (A) show the "good" and "detail" insets indicated in the top-left image. On the one hand, the classical thresholding algorithm is too aggressive and removes good and bad localizations. On the other hand, thresholding on an engineered feature leaves enough bad localizations behind to create artifacts in the final image. Using gradient boosted trees strikes a balance between these two extremes and offers optimal filtering. Fig. S36B compares distributions of various peak parameters between the "good" (top row) and "bad" (bottom row) regions shown in the upper left corner of (A) along with the manually chosen thresholds shown as orange lines. Fig. S36C is the same as (B) but for the manually engineered feature: $\chi^2 \times \mathtt{offset/amp}$. Fig. S36D shows the number of grouped localizations in the data set for each filtering method.

#### *16. FIB-SEM imaging*

A custom Zeiss Merlin crossbeam system (*12*) was further modified for this work in the following ways: the FEI Magnum FIB column was replaced with a Zeiss Capella FIB column; the Zeiss Capella FIB column was repositioned at 90 degrees to the SEM column; and the custom NI LabVIEW control software was re-written to switch from the Zeiss RemCon communication protocol to the Zeiss API which reduced the overhead by several seconds per imaging/milling cycle.

Imaging conditions were further optimized for cellular imaging from (*12*). Standard (8 x 8 x 8 $nm^3$ isotropic voxel) image stacks were acquired at 500 kHz/pixel with a x-y pixel resolution of 8 nm using a 2 nA electron beam at 1.2 kV landing energy for imaging and a 15 nA gallium ion beam at 30 kV for FIB milling. No stage bias was used. Both backscattered and secondary electron signals were collected by InLens detector to provide better signal-to-noise ratio.

The milling was done in 4 nm steps to form raw image volumes. Milling in 4 nm steps (even for 8 x 8 x 8 $nm^3$ isotropic voxels) has few advantages:

1. When milling is done at 4 nm, two images are collected for each 8 nm voxel, which are later binned. Therefore, each image can be collected at reduced SNR – at roughly 2x imaging speed. Faster imaging results in lower electron dose and lower sample damage ("cooking") by the electron beam. Along with faster FIB milling this results in fewer milling artifacts.

2. In a rare case when a single frame is lost or data is corrupted, less information is lost.

There is no significant temporal overhead for milling at 4 nm steps and faster SEM imaging relative to milling at 8 nm steps and slower SEM imaging.

Compared to sample-biased conditions used previously (*12*), more FIB milling artifacts such as streaks were present in raw images due contamination from secondary electron signal. A Fourier mask was applied to remove these artifacts.

4 x 4 x 4 $nm^3$ voxel datasets were generated using a similarly customized Zeiss Gemini 500-Capella Crossbeam system. The block face was imaged by a 250 pA electron beam with 0.9 kV landing energy at 200 kHz. The x-y pixel resolution and z milling step were both set at 4 nm.

The final image stacks were registered using a SIFT (*93*) based algorithm. 8 nm pixel datasets were then binned by a factor of 2 along z to achieve isotropic 8 nm voxels.

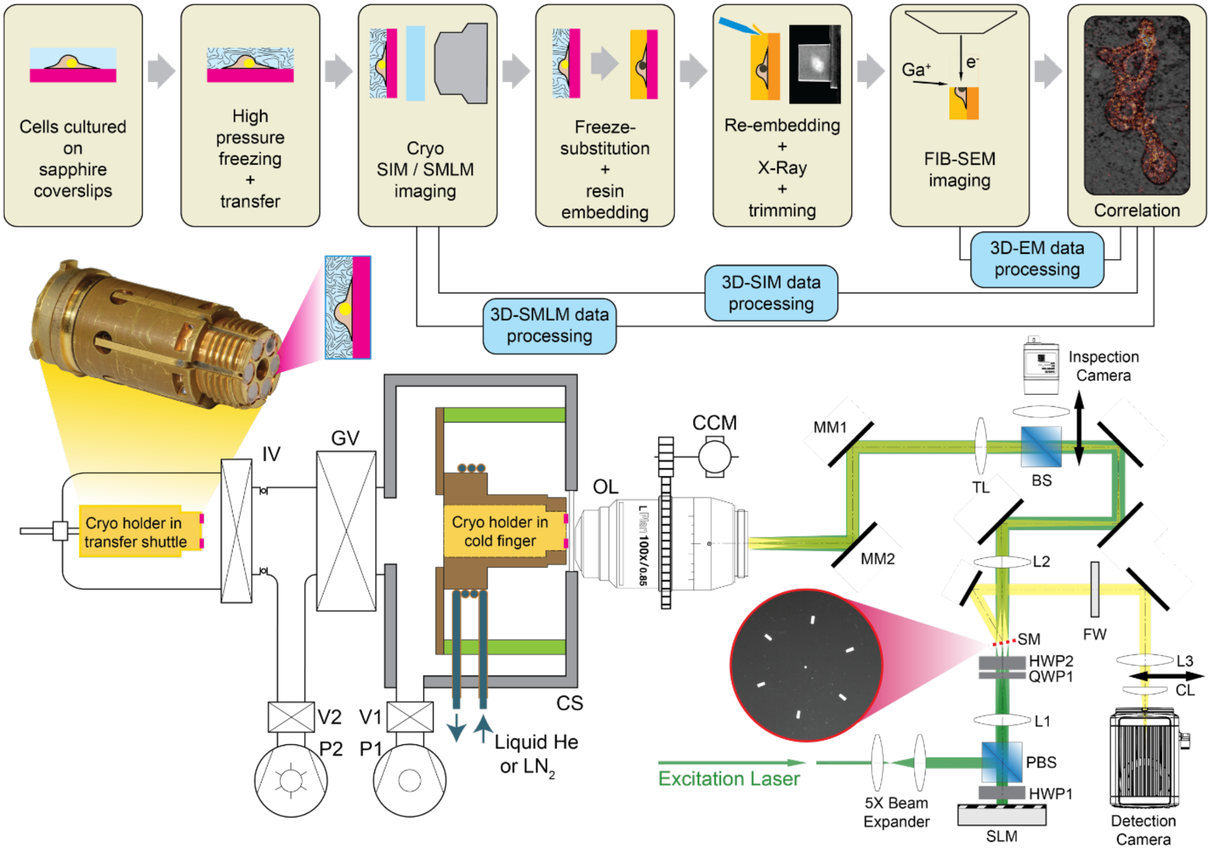


**Fig. S1.**

**Pipeline, experimental workflow and instrument schematic.** Top row: Sequence of sample preparation and imaging steps. Cells cultured on sapphire coverslips are high pressure frozen and transferred into the cryogenic fluorescence microscope for imaging and then removed, freeze substituted, resin embedded, trimmed and imaged by FIB-SEM. Cryo-SR and FIB-SEM data is then processed, registered and correlated. Bottom left: Schematic of the optical cryostat (see Fig. S6 for details) and sample transfer airlock. Inset shows a photograph of the custom designed cryo-sample holder (see Fig. S4 for details). Bottom right: Schematic of the SIM/SMLM microscope and optical paths. Inset shows the mirrored SIM mask (SM) which separates excitation and detection light in a wavelength-independent manner. See supplementary note 3 for details and abbreviations.

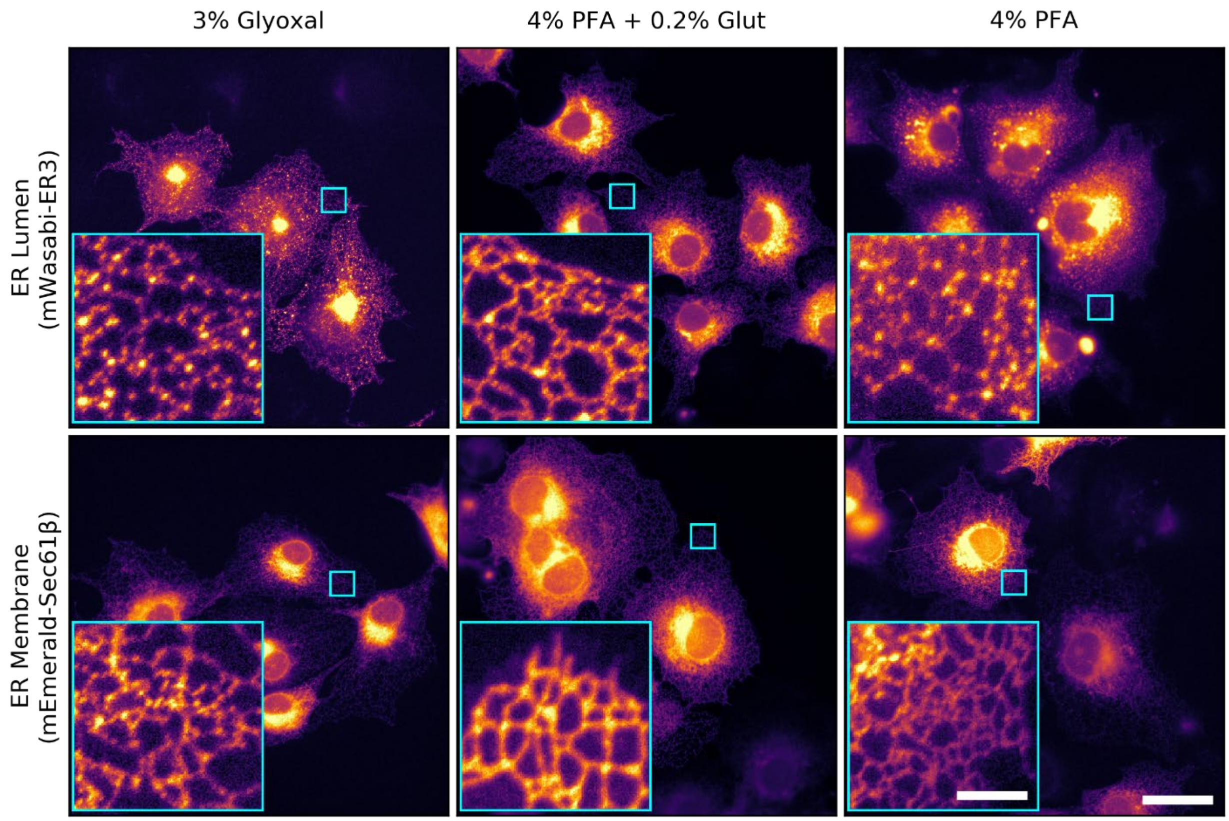


**Fig. S2.**

**Comparison of chemical fixation methods.** Widefield fluorescence images of COS7 cells transiently expressing either ER lumen marker mWasabi-ER3 (top row) or membrane marker mEmerald-Sec61β (bottom row), chemically fixed by either 3% glyoxal ((*130*) left column), 4% paraformaldehyde (PFA) with 0.2% glutaraldehyde (glut, center column), and 4% PFA (right column). Note that in all cases the membrane protein (bottom row) is better preserved then the lumen protein. 4% PFA + 0.2% glut offers the best preservation of both the ER lumen and membrane.

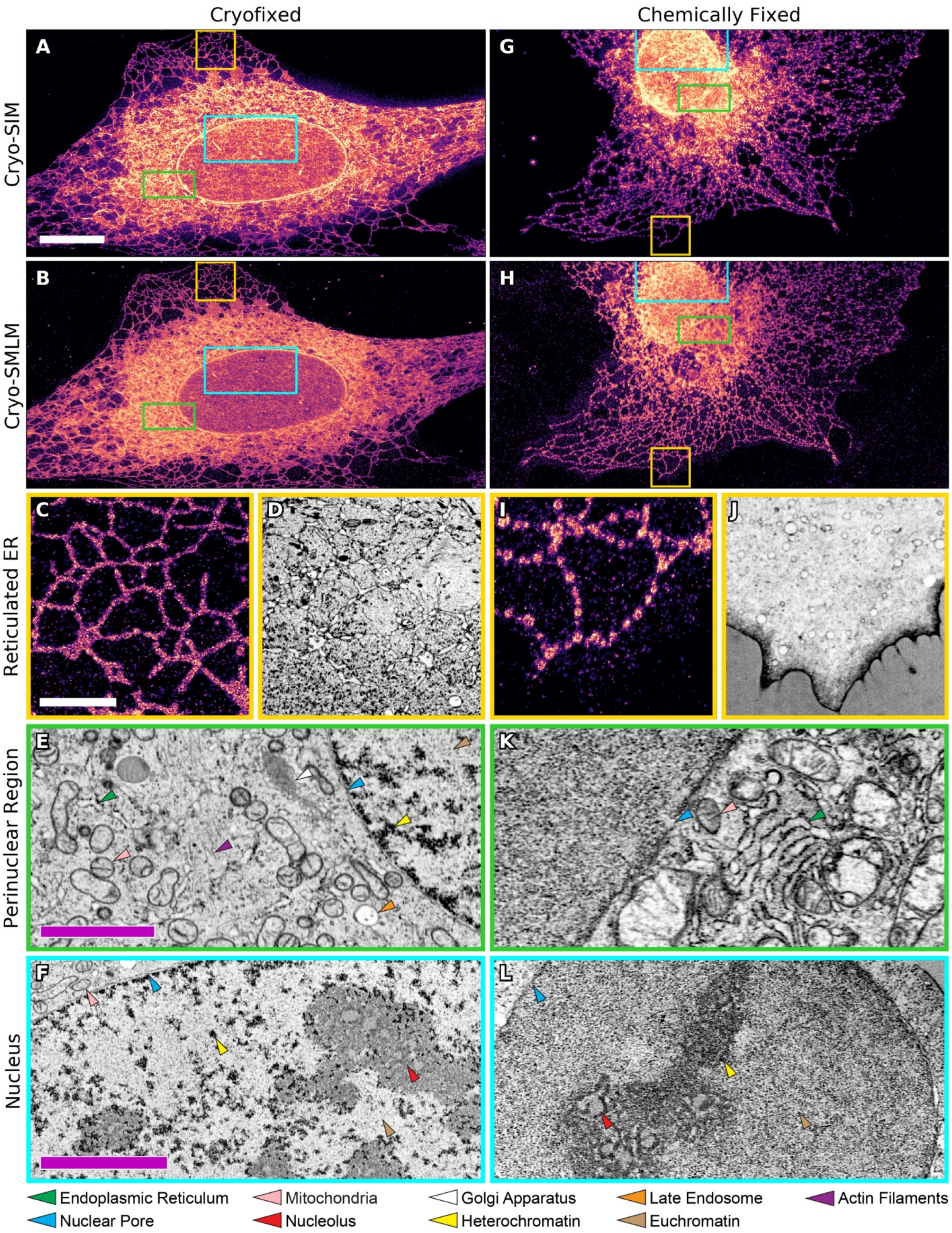

Cryofixed
Chemically Fixed
Cryo-SIM
Cryo-SMLM
Reticulated ER
Perinuclear Region
Nucleus
A
B
C
D
E
F
G
H
I
J
K
L
Endoplasmic Reticulum
Mitochondria
Golgi Apparatus
Late Endosome
Actin Filaments
Nuclear Pore
Nucleolus
Heterochromatin
Euchromatin

**Fig. S3.**

**Cryofixation offers ultrastructure preservation superior to chemical fixation.** Cryo-correlative data from two U2OS cells transiently expressing mEmerald-Sec61β. **(A-F)** High pressure frozen cell, **(G-L)** cell chemically fixed at room temperature with 4% paraformaldehyde (PFA) with 0.1% glutaraldehyde (*112*). Low magnification cryo-SIM (A, G) and cryo-SMLM (B, H) MIPs presented with color coded boxes representing the corresponding zoom regions in subsequent panels. (C, I) Correlated cryo-SMLM and (D, J) FIB-SEM (MIP through 32 nm thick slabs) of reticulated ER in the periphery of the cells. (E, K) FIB-SEM orthoslices in the perinuclear region and (F, L) the nucleus (Movie S1). Colored arrow heads indicate common cellular structures according to legend at bottom. In all cases, structure preservation via high pressure freezing is superior. Scale bars: (A, B, G, H) 10 μm; (C, D, I, J) 2 μm; (E, K) 2 μm; (F, L) 4 μm.

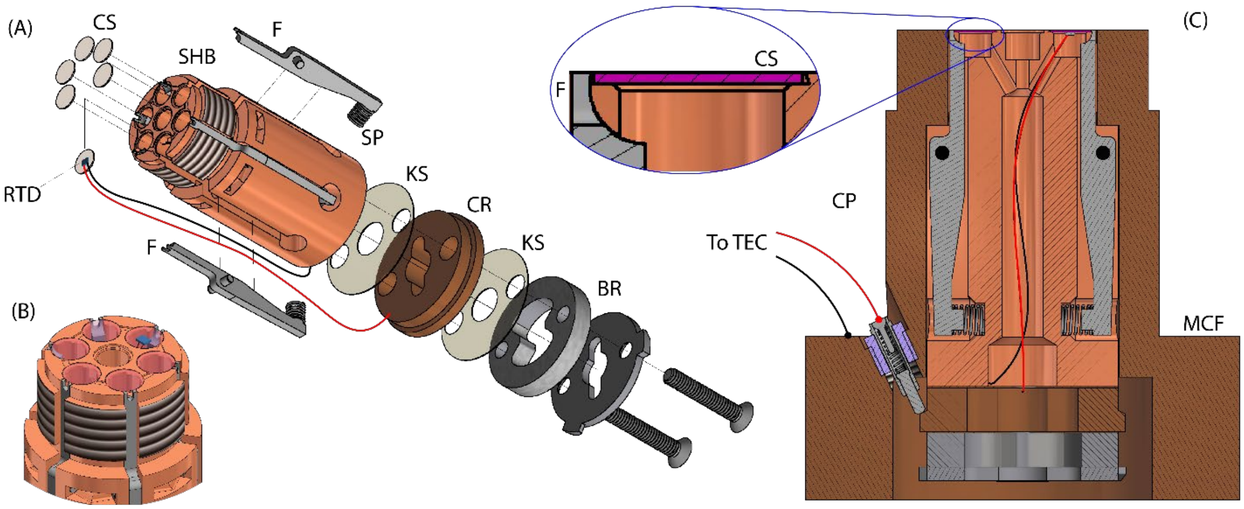


**Fig. S4.**

**Custom designed cryogenic sample holder**. **(A)** Exploded view of the sample holder. Six sapphire coverslips (CS) with HPF cultured cells are held in dovetailed pockets in sample holder body (SHB) by spring-loaded (SP) stainless steel fingers (F). A resistance temperature detector (RTD) epoxied to one CS is connected by two contact wires, one to SHB and the other to contact ring (CR), the latter electrically insulated by Kapton spacers (KS). Bayonet receptacle (BR) permits transfer of the assembly from the sample preparation chamber to the imaging cryostat. **(B)** Top oblique view of the assembled cryogenic sample holder. **(C)** Sample assembly as inserted within modified cold finger (MCF), similar to the standard part supplied with Janis ST-500 Cryostat. Spring-loaded contact pin (CP) transfer sample temperature signal from RTD to an external temperature controller (TEC). Another temperature detector and a heater element (neither shown here) regulate temperature within the cryostat itself.

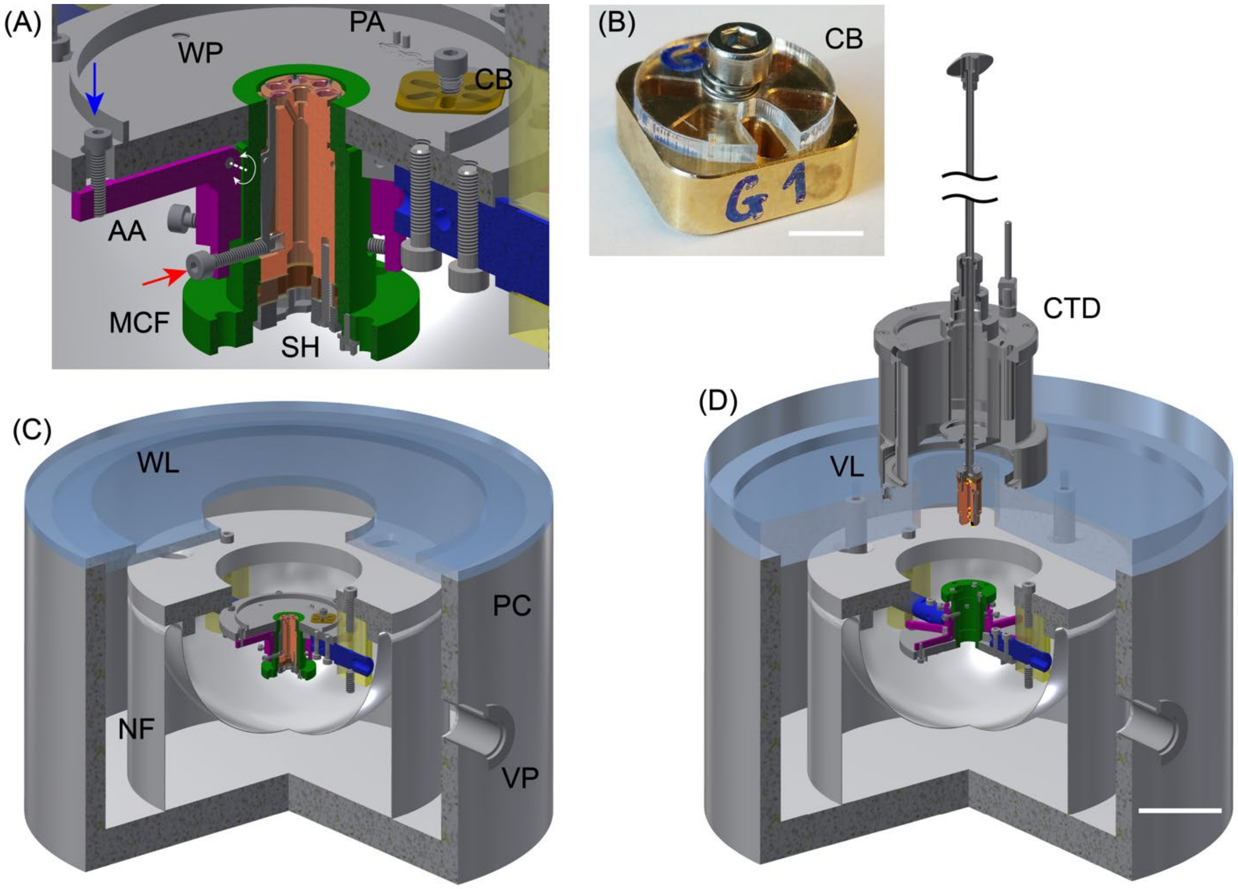


**Fig. S5.**
**Schematics and operation of the cryogenic sample preparation chamber. (A)** The work platform (WP) with sample holder (SH), held by modified cold finger (MCF). Actuator arm (AA) pivots on the axis indicated by a white dotted line. Pushing one arm of AA in the direction indicated by the blue arrow makes the other arm of AA move in the direction indicated by the red arrow, which permits coverslips to be loaded and unloaded from SH. **(B)** Photograph of the cryo box (CB) used for sample storage. Scale bar: 5 mm. **(C)** Preparation chamber (PC) with the nitrogen flask (NF) and WP suspended in NF facing upward for sample loading. The work lid (WL) is placed over PC to reduce water condensation. **(D)** PC with WP turned upside down for sample transfer. WL is replaced with vacuum lid (VL) for vacuum docking of cryo transfer device (CTD) used to transfer SH between PC and optical cryostat. SH is shown in a mid-way position as it is being retracted from MCF into CTD. Scale bar: 50 mm.

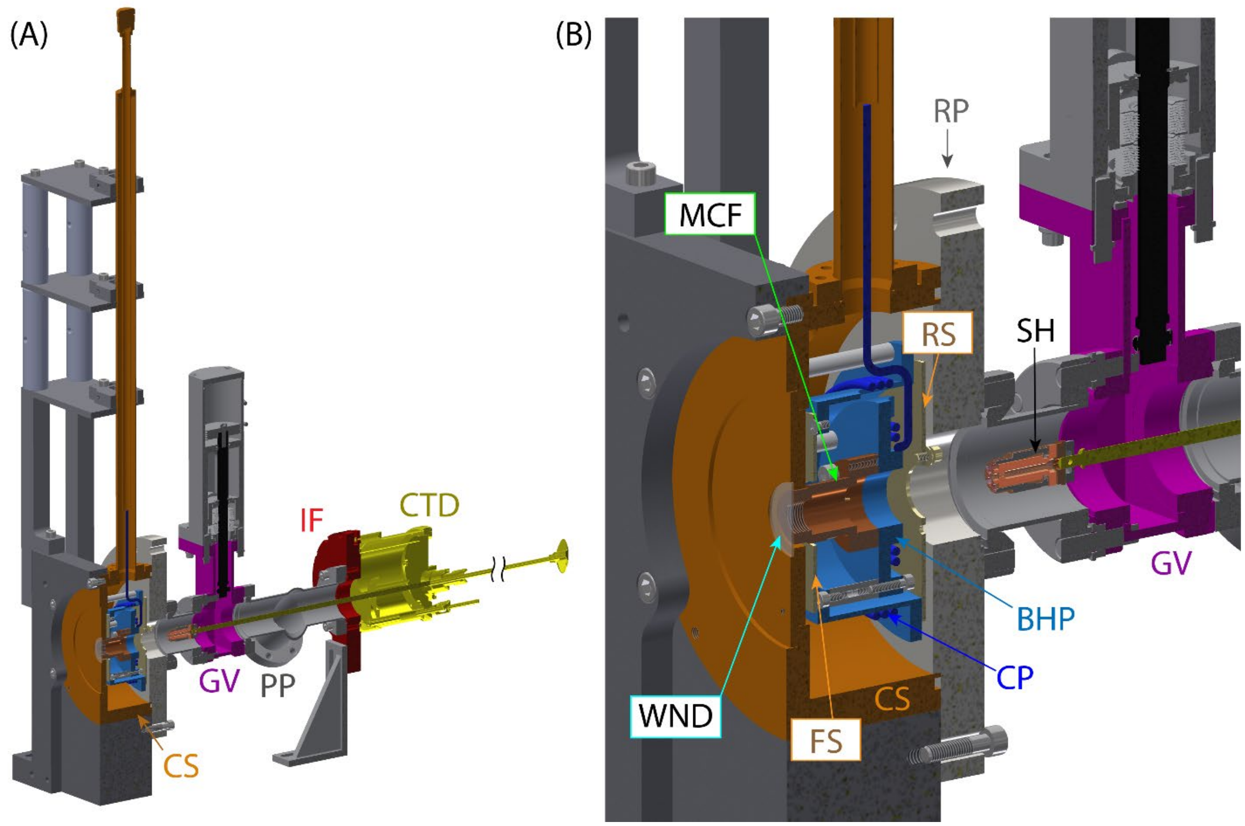


**Fig. S6.**

**Optical cryostat and transfer interlock**. **(A)** Half-section view of the optical cryostat and transfer interlock. Modified Janis ST-500 optical cryostat (CS, dark orange) is mounted in the metal cage so that cryogen transfer line is oriented vertically. Modified rear plate (RP) of CS is connected to Gate Valve (GV). On the opposite side, GV is connected to Interlock Flange (IF) that can accept cryogenic transfer device (CTD). The volume between GV and IF is the body of the interlock that can be evacuated via pumping port (PP). **(B)** Zoomed view of CS and GV. Modified cold finger (MCF) is mounted on the bottom heat exchanger plate (BHP), which is cooled by a constant flow of a cryogen through cooling pipes (CP). Sample holder (SH) is shown in the retracted position (attached to the front end of the CTD transfer rod) before it is docked into MCF. The heat exchanger is protected from black-body radiation by front and rear radiation shields (FS and RS, respectively). The rear shield has a swing door that can be opened and closed for sample loading. The front of the sample holder in the docked position is separated from the cryostat optical window (WND) by ~300 μm vacuum gap.

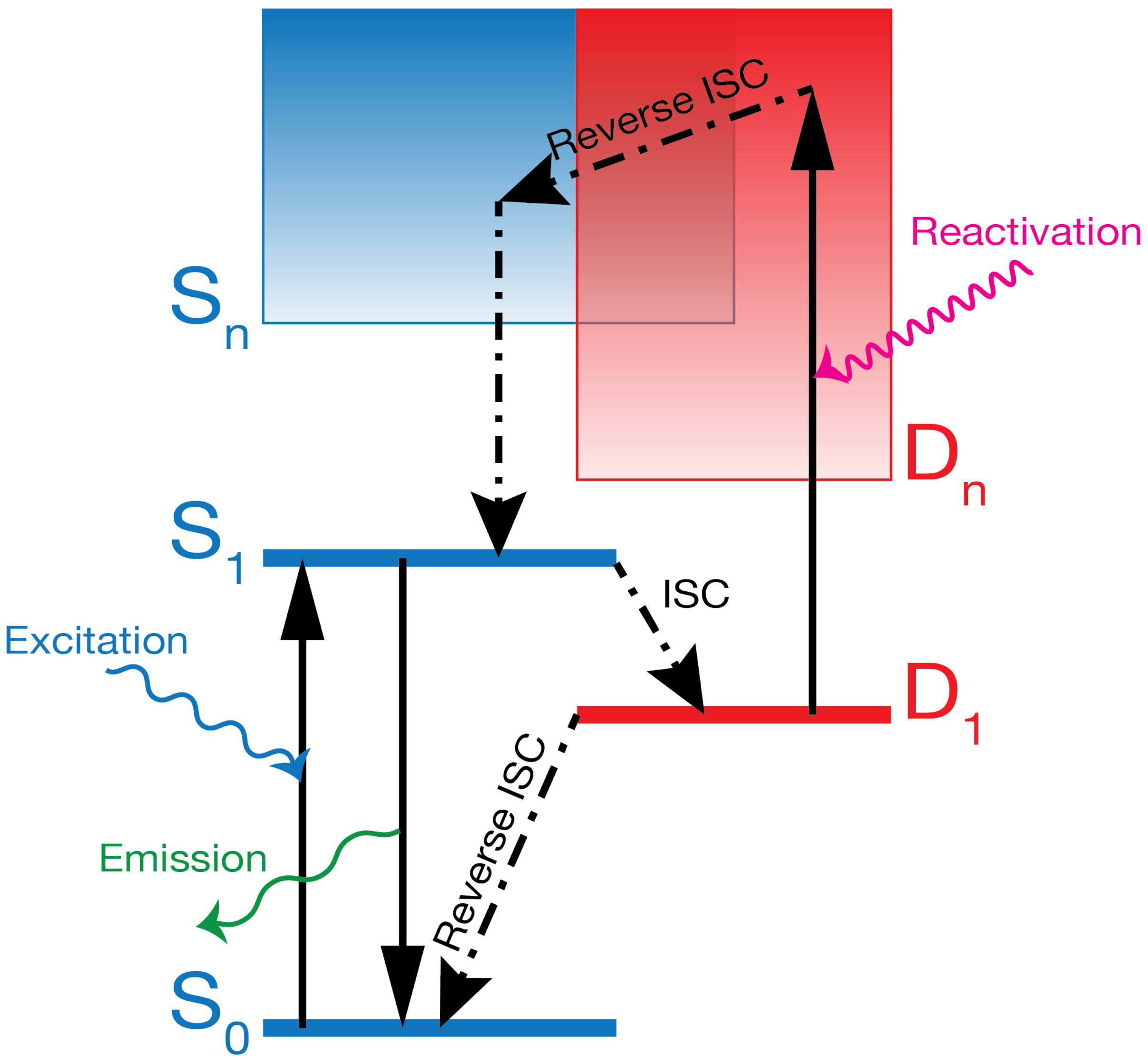


**Fig. S7.**

**Hypothetical energy level diagram for dark state shelving and photoactivation.** Jablonski diagram hypothesized for the energy landscape of a fluorophore at cryogenic temperatures. Horizontal bars represent stable energy levels. $S_0$, $S_1$ are singlet states, the transitions between which correspond to single photon excitation or emission. $D_1$ and $D_n$ are dark states of unknown character. Solid and dot-dashed arrows represent radiative and non-radiative transitions, respectively. Wavy arrows represent absorption or emission of a photon. The lifetime for the spontaneous transition $D_1 \rightarrow S_0$ must be long to achieve a high dynamic contrast ratio in SMLM. For SMLM there also should exist a radiative transition $D_1 \rightarrow D_n$ which can be used to return the molecule to emissive S0↔S1 cycling.

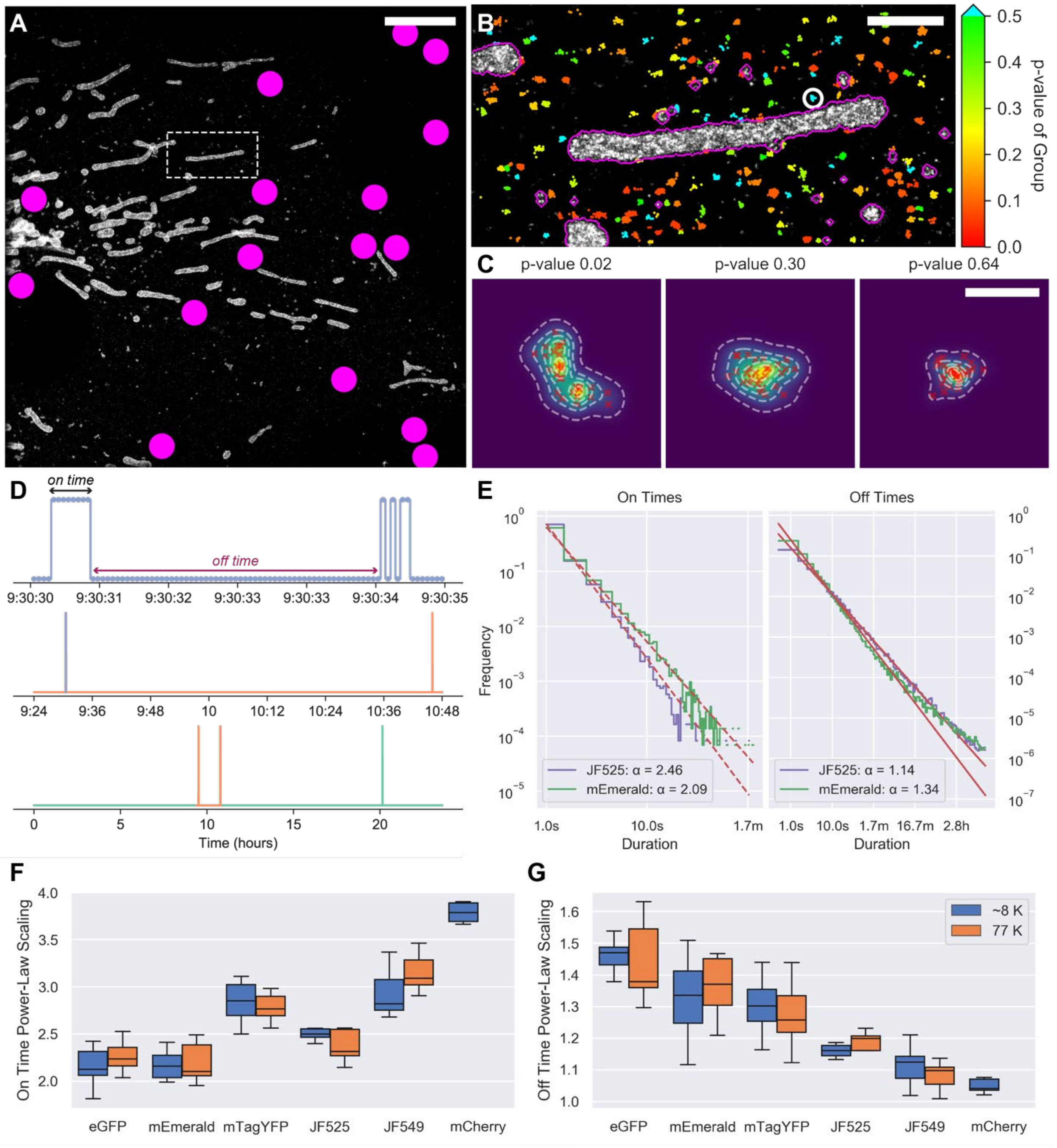


**Fig. S8.**

**Characterizing the on/off switching behavior of single molecules from cryo-SMLM data. (A)** Cryo-SMLM image of a U2OS cell transiently expressing mEmerald-TOMM20; fiducial locations are marked with magenta circles. Scale bar, 5 µm. **(B)** Single mitochondrion and surrounding localization clusters from the boxed region in (A). Scale bar, 1 µm. Large clusters (with magenta borders) are removed, and remaining ones are color coded according to the likelihood that each arises from a single molecule (supplementary note 4a). **(C)** Three examples of the remaining localization clusters, rendered by overlapping Gaussian images, with localized centers shown as red crosses, and p-values as shown. Scale bar, 100 nm. **(D)** Time traces from the cluster circled in white in (B), at three temporal scales. Top trace at single frame resolution shows individual on and off events. **(E)** Histograms of on and off times for mEmerald (top) and Halo-JF525 (bottom), each fused to TOMM20 at the mitochondrial outer membrane, indicating the power law dependence of each. **(F, G)** Box plots of on and off time power law scaling for different fluorophores and temperatures.

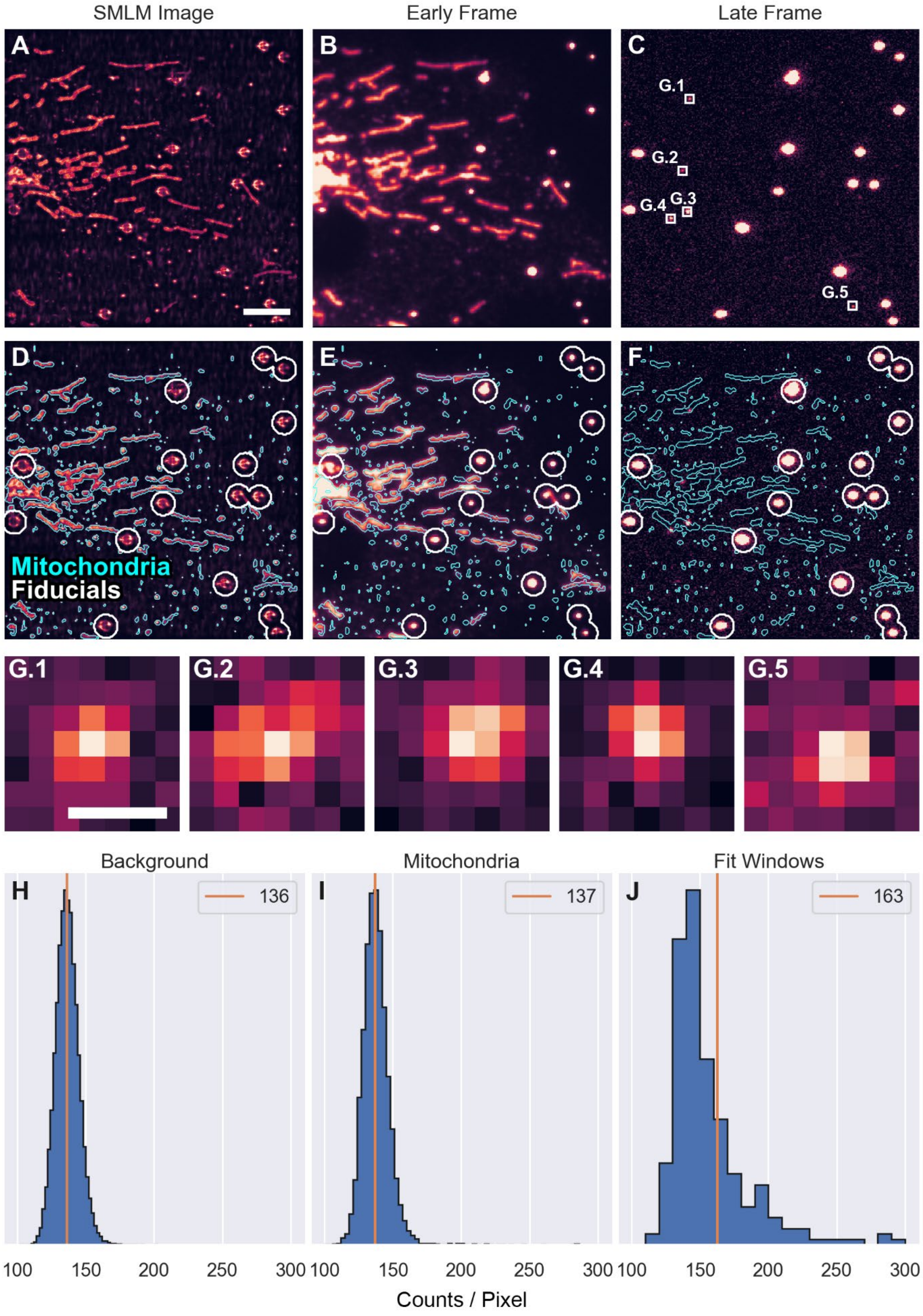
SMLM Image
Early Frame
Late Frame
A
B
C
G.1
G.2
G.4
G.3
G.5
D
E
F
Mitochondria
Fiducials
G.1
G.2
G.3
G.4
G.5
Background
Mitochondria
Fit Windows
H
136
I
137
J
163
100
150
200
250
300
Counts / Pixel

**Fig. S9.**

**Method to calculate the static contrast ratio in SMLM data. (A)** SMLM reconstruction, **(B)** an early raw frame in the acquisition, and **(C)** a frame late in the acquisition with boxes around detected peaks. Scale bar: 5 µm. See also Movie 1. **(D-F)** Same frames as shown above overlaid with mitochondrial and fiducial masks (supplementary note 4b). **(G)** Zoomed views of detected peaks in (C). Scale bar: 0.5 µm. Histograms of counts per pixel in: **(H)** the late frame outside of the fiducial and mitochondrial masks; **(I)** the late frame within the mitochondrial mask but outside the fit windows; and **(J)** the late frame within the fit windows. Orange lines indicate the median, median, and mean values, respectively.

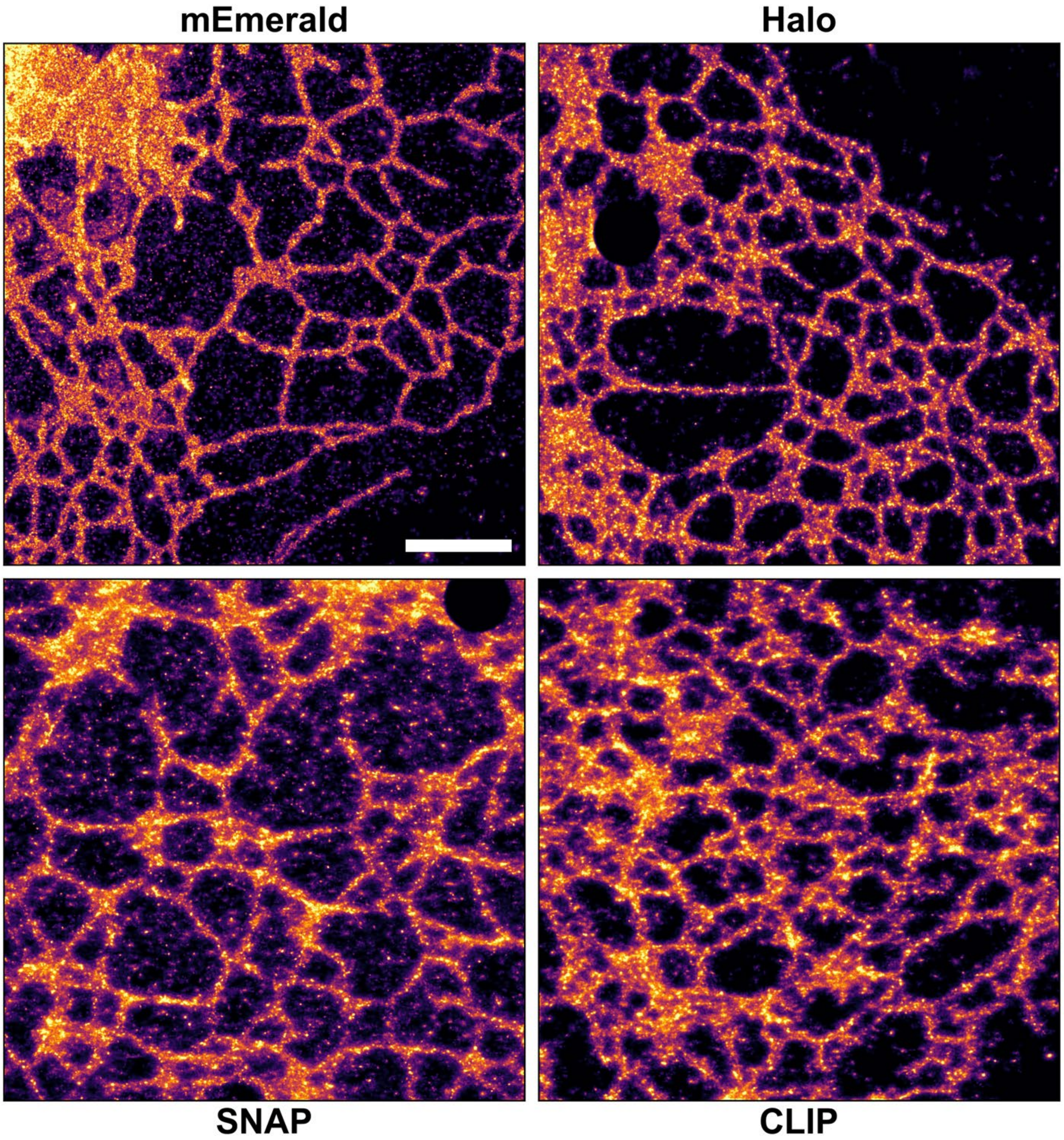


**Fig. S10.**

**Comparison of different labeling strategies.** Cryo-SMLM data sets from U2OS cells transiently expressing Sec61β fused to mEmerald, HaloTag, SNAP-tag, or CLIP-tag. The latter three are conjugated to JF525. Fiducial beads have been removed where applicable. Scale bar: 2 µm.

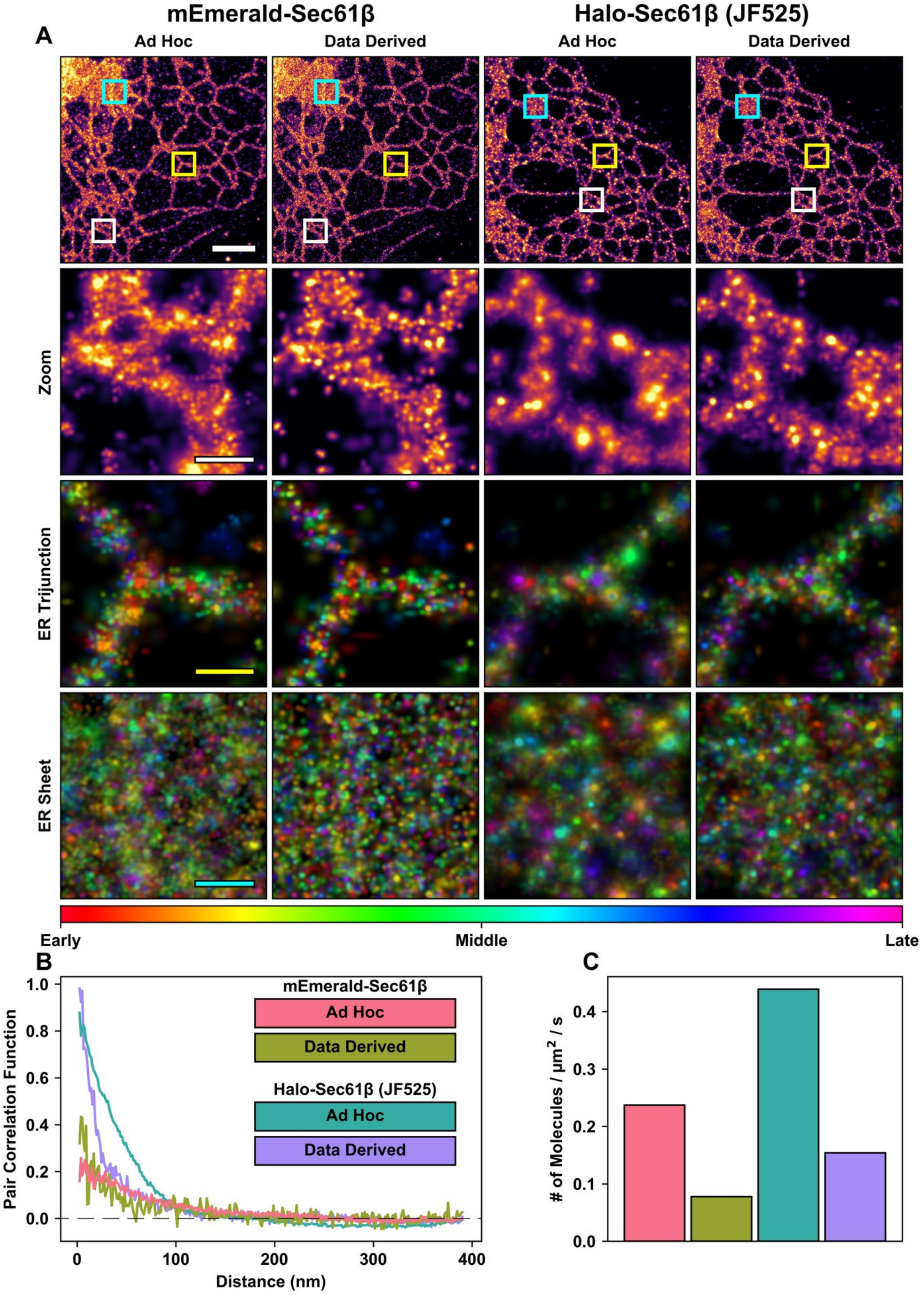
mEmerald-Sec61β
Halo-Sec61β (JF525)
A
Ad Hoc
Data Derived
Ad Hoc
Data Derived
Zoom
ER Trijunction
ER Sheet
Early
Middle
Late
B
Pair Correlation Function
Distance (nm)
mEmerald-Sec61β
Ad Hoc
Data Derived
Halo-Sec61β (JF525)
Ad Hoc
Data Derived
C
### of Molecules / µm² / s

**Fig. S11.**

**Data derived grouping of molecular switching events improves data quality and accuracy.** **(A)** SMLM images of U2OS cells transiently transfected with ER membrane protein Sec61β, fused to either mEmerald (columns 1 and 2) or HaloTag conjugated to JF525 (columns 3 and 4), with multiple switching events grouped using either parameters chosen *ad hoc* (columns 1 and 3) or derived from our cryo-photophysical measurements (columns 2 and 4, supplementary note 5). Scale bar, 2 µm. Top row: large field of view comparisons, with colored boxes denoting regions of differing ER morphology shown in subsequent rows. Second row: zoomed views from white boxes in the top row. Rows 3 and 4: zoomed views from yellow and cyan boxes in the top row, respectively, with localization events color-coded by time of acquisition as shown. Scale bars, 300 nm. **(B)** Pair correlation functions and **(C)** estimated molecular densities for the regions in the bottom row of (A), demonstrating the ability of our data driven approach to more accurately assign multiple switching events to the correct molecular source.

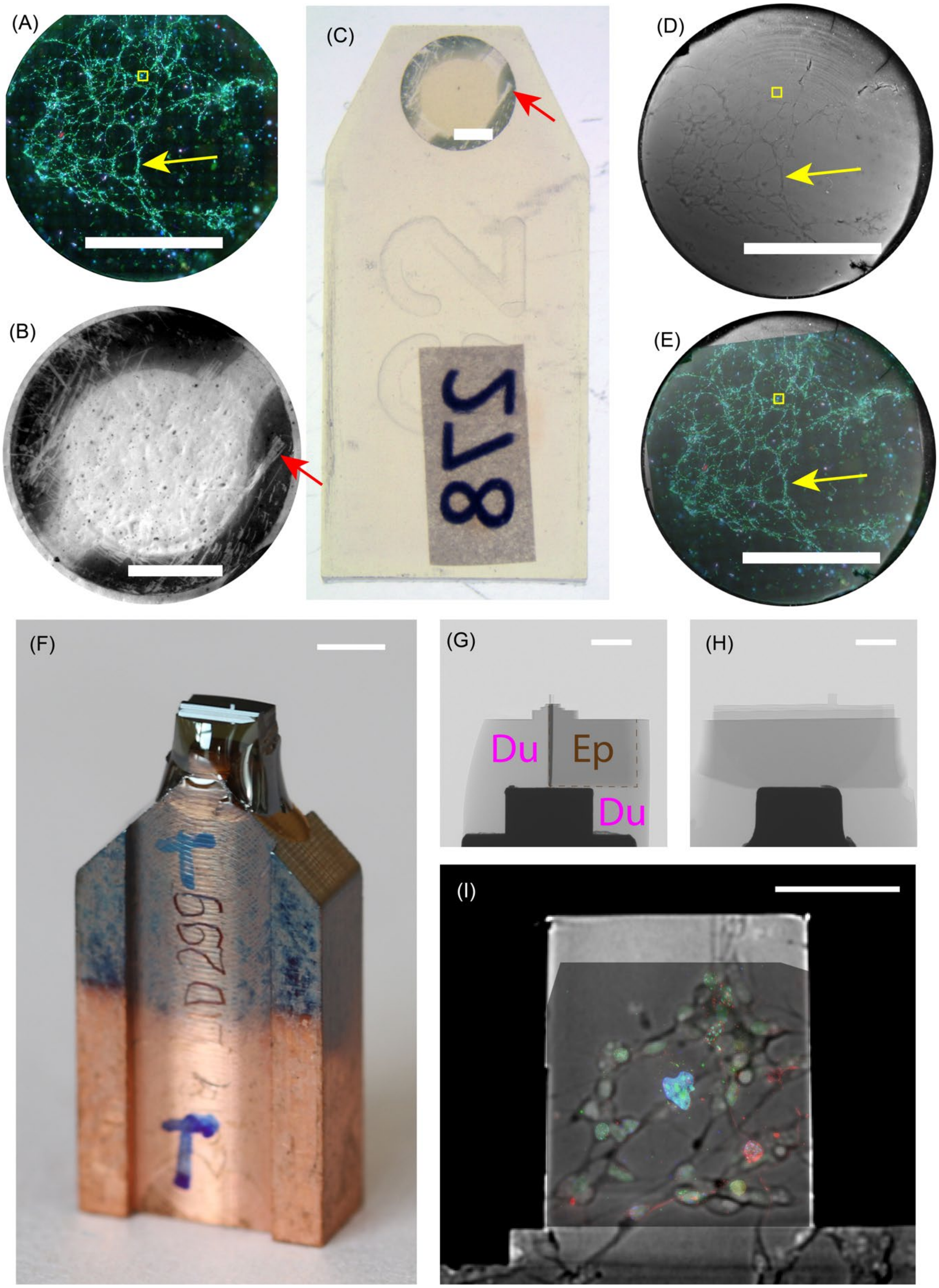
(A)
(B)
(C)
(D)
(E)
(F)
(G)
Du
Ep
Du
(H)
(I)

**Fig. S12.**
**Identifying the cryo-SR field of view after resin embedding in preparation for correlative FIB-SEM imaging. (A)** Widefield fluorescence image of the entire sapphire disk acquired under cryogenic conditions prior to SR imaging. Yellow box indicates the cryo-imaged field of view. **(B)** Brightfield photo of the same disk in the preparation chamber (fig. S5) under liquid nitrogen after removal from the cryostat. **(C)** Brightfield photo of the same disk after freeze substitution and embedding in Eponate 12 resin. Scratches (red arrows in (B) and (C)) uniquely identify each disk. **(D)** X-ray image of the specimen within the resin block. **(E)** Overlay of the images in (A) and (D), yellow arrows indicate landmarks common to both. **(F)** Resin block re-embedded in Durcupan resin and mounted on a copper stud for trimming. **(G, H)** Orthogonal X-ray projections of the sample after trimming. The lighter volume (marked by Du in magenta in (G)) is Durcupan epoxy. A slightly darker volume marked by brown Ep and outlined by a dashed brown line in (G) is original Epon-embedded volume. Still darker line on the left boundary of Epon volume in (G) is a layer that contained media+cryoprotectant during high-pressure freezing. **(I)** Overlay of the cryogenic 3-channel SIM image with the X-ray projection of the sample stub prepared for FIB-SEM imaging. Scale bars: (A-E) 1 mm; (F) 2 mm; (G, H) 500 µm; (I) 50 µm.

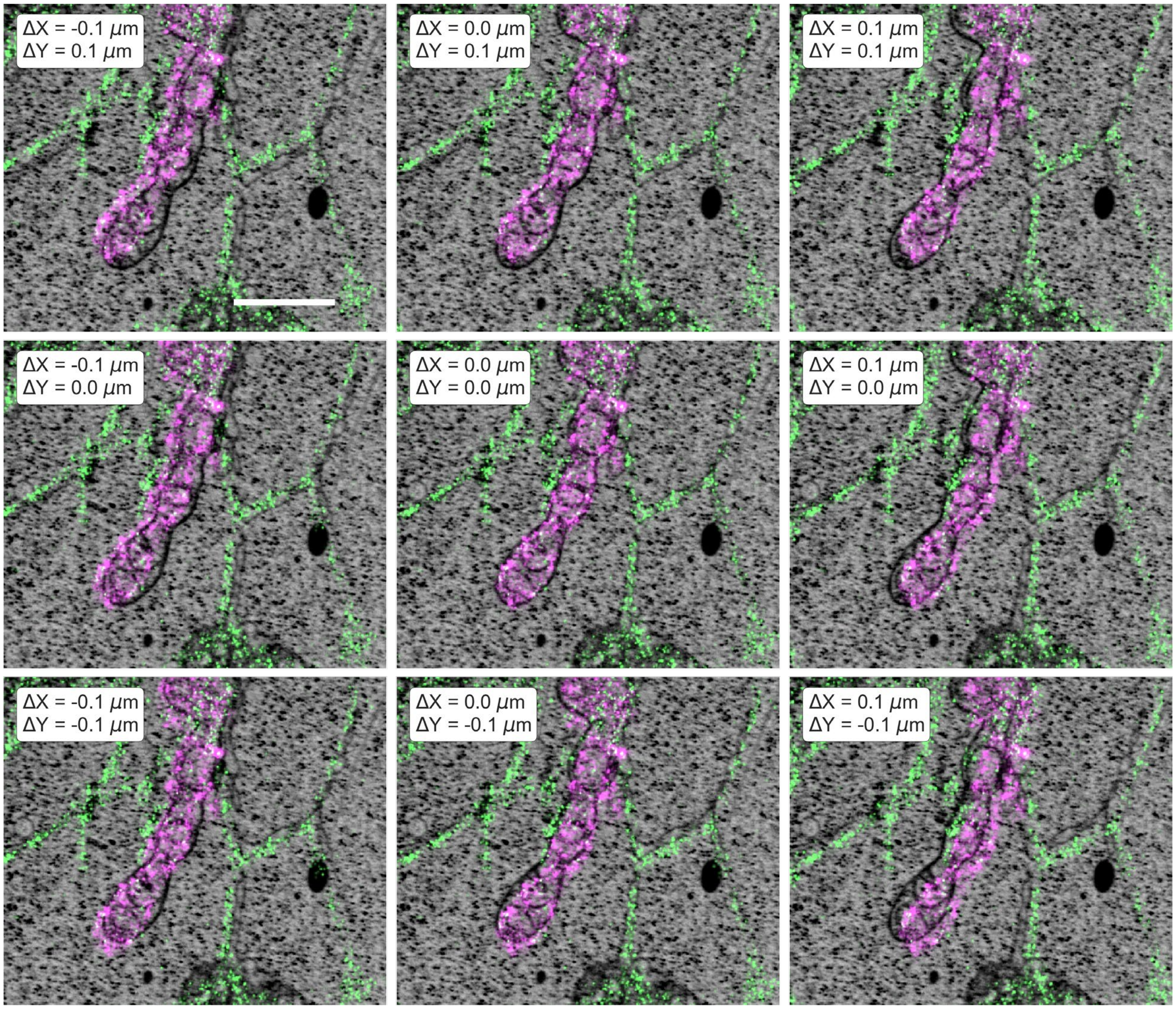


**Fig. S13.**

**Sensitivity of Cryo SMLM and FIB-SEM image overlap to different offsets.** Center panel: cryo SMLM and FIB-SEM images registered by aligning mitochondria and ER organelles in two data sets. Perimeter panels: cryo SMLM image shifted by the displacements $\Delta X$= -0.1, 0.0, 0.1 μm and $\Delta Y$= -0.1, 0.0, 0.1 μm relative to FIB-SEM image to demonstrate precision of manual landmark registration. Scale bar: 1.0 μm.

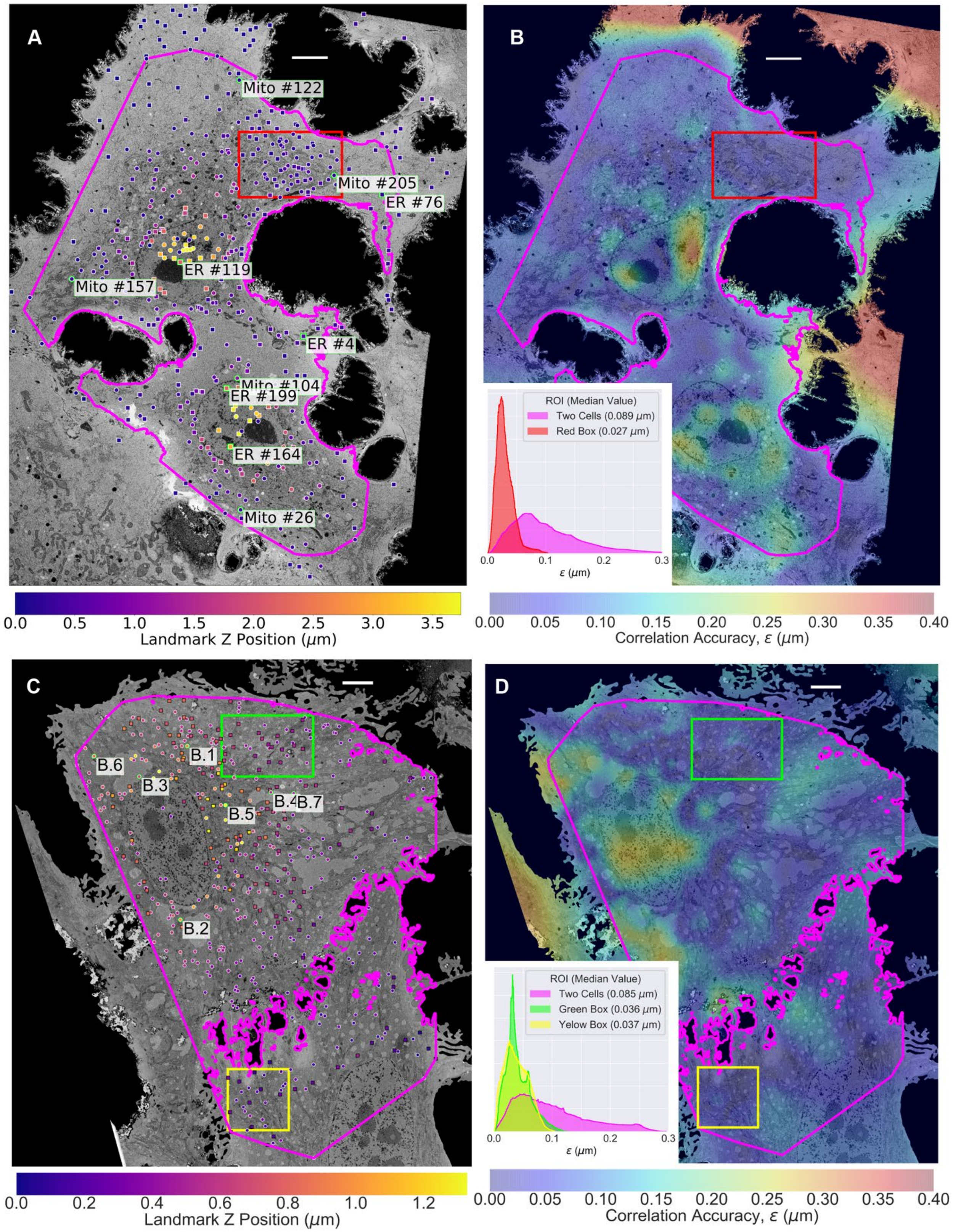
A
B
C
D
Mito #122
Mito #205
ER #76
ER #119
Mito #157
ER #4
Mito #104
ER #199
ER #164
Mito #26
ROI (Median Value)
Two Cells (0.089 μm)
Red Box (0.027 μm)
ε (μm)
Landmark Z Position (μm)
Correlation Accuracy, ε (μm)
B.1
B.6
B.3
B.4
B.7
B.5
B.2
ROI (Median Value)
Two Cells (0.085 μm)
Green Box (0.036 μm)
Yelow Box (0.037 μm)
ε (μm)
Landmark Z Position (μm)
Correlation Accuracy, ε (μm)

**Fig. S14.**

**Figure S14. Organelle landmark positions for correlation of light and electron microscopy, and maps of differences in the resulting two independent measurements of the displacement field. (A)** Map of all mitochondrial and ER landmarks (circles and squares, respectively) used for LM and EM image registration in Fig. 3 projected onto the "mid-cell" EM surface (fig. S33). Landmarks are color-coded by their Z position within the specimen. Numbered landmarks with green borders are shown in orthoslices in fig. S15. **(B)** Map of the correlation accuracy, ε (cf. supplementary note 7), at the "mid-cell" height for the data in Fig. 3. Inset: histograms of ε, color coded by the boundaries defining their regions of measurement in the "mid-cell" image. **(C)** Map similar to (A) of all peroxisomal and mitochondrial landmarks (circles and squares, respectively) used for LM and EM image registration in Fig. 4 projected onto the "mid-cell" EM surface. Landmarks with green borders represent the seven peroxisomes shown in Fig. 4. **(D)** Map similar to (B) of ε at the "mid-cell" height for the data in Fig. 4. Inset: distribution of ε values inside the color-coded boundaries in the "mid-cell" image. Scale bars, 5 μm.

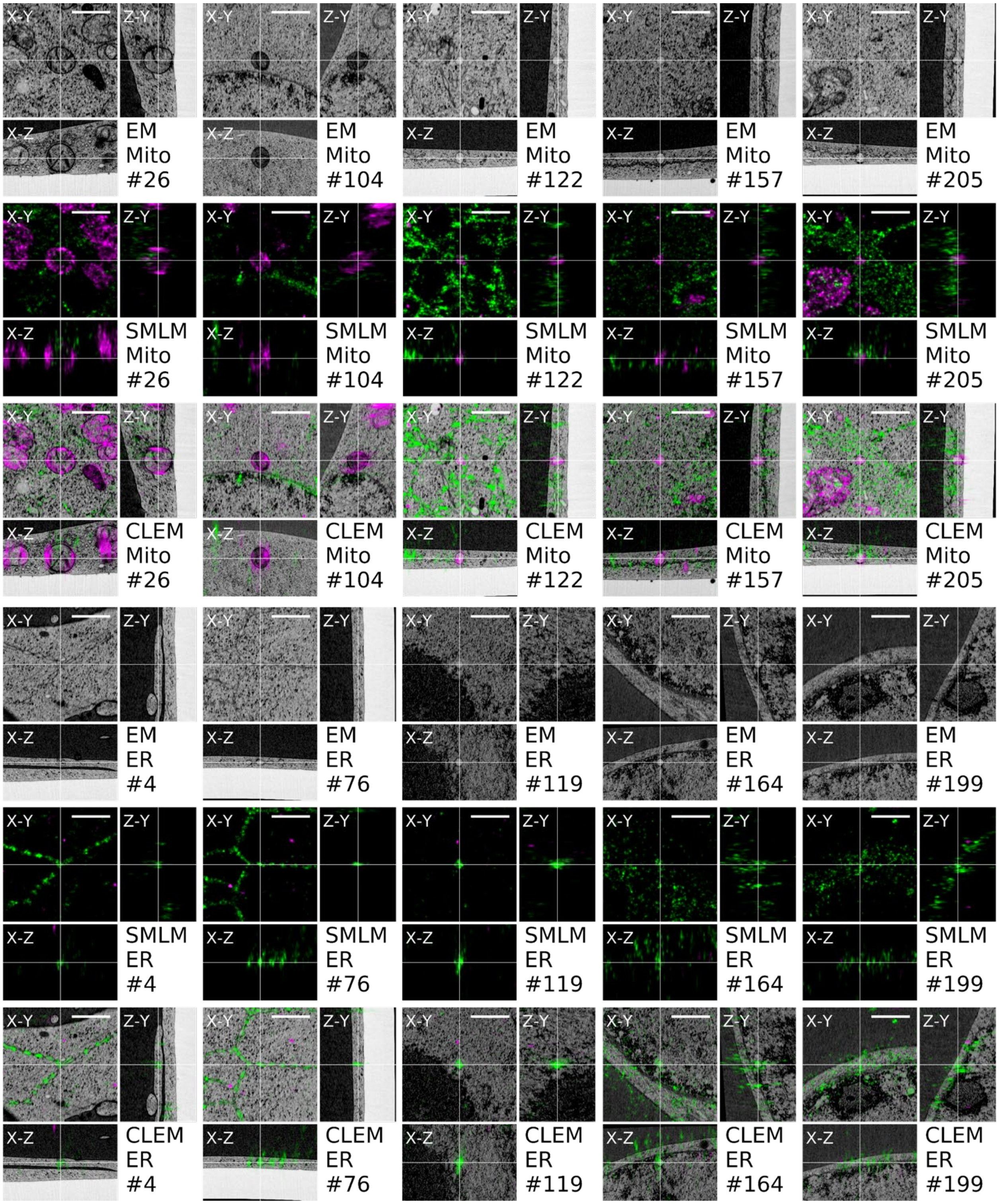


**Fig. S15.**

**High resolution CLEM images of various regions in the cell shown in Figure 3.** Rows 1 and 4: X-Y, Z-Y and X-Z cross-sections of EM data sets. Rows 2 and 5: X-Y, Z-Y and X-Z cross-sections of PALM data sets at the same coordinates. Rows 3 and 6: X-Y, Z-Y and X-Z cross-sections of overlaid EM and PALM data sets. Scale bars: 1 μm

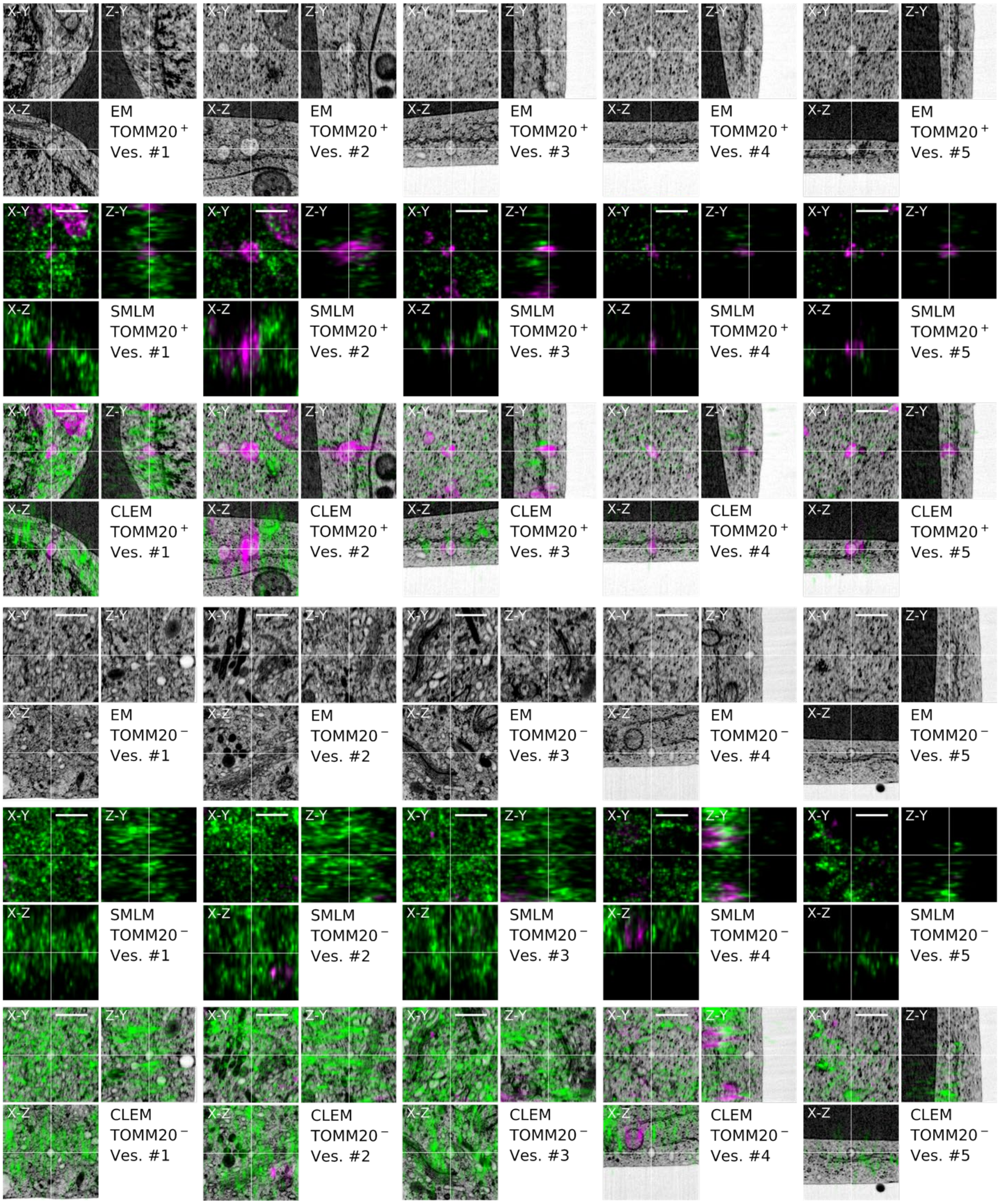
EM
TOMM20⁺
Ves. #1
EM
TOMM20⁺
Ves. #2
EM
TOMM20⁺
Ves. #3
EM
TOMM20⁺
Ves. #4
EM
TOMM20⁺
Ves. #5
SMLM
TOMM20⁺
Ves. #1
SMLM
TOMM20⁺
Ves. #2
SMLM
TOMM20⁺
Ves. #3
SMLM
TOMM20⁺
Ves. #4
SMLM
TOMM20⁺
Ves. #5
CLEM
TOMM20⁺
Ves. #1
CLEM
TOMM20⁺
Ves. #2
CLEM
TOMM20⁺
Ves. #3
CLEM
TOMM20⁺
Ves. #4
CLEM
TOMM20⁺
Ves. #5
EM
TOMM20⁻
Ves. #1
EM
TOMM20⁻
Ves. #2
EM
TOMM20⁻
Ves. #3
EM
TOMM20⁻
Ves. #4
EM
TOMM20⁻
Ves. #5
SMLM
TOMM20⁻
Ves. #1
SMLM
TOMM20⁻
Ves. #2
SMLM
TOMM20⁻
Ves. #3
SMLM
TOMM20⁻
Ves. #4
SMLM
TOMM20⁻
Ves. #5
CLEM
TOMM20⁻
Ves. #1
CLEM
TOMM20⁻
Ves. #2
CLEM
TOMM20⁻
Ves. #3
CLEM
TOMM20⁻
Ves. #4
CLEM
TOMM20⁻
Ves. #5
X-Y
Z-Y
X-Z

**Fig. S16.**

**Correlative cryo-SMLM/FIB-SEM orthoslices of TOMM20-positive and TOMM20-negative vesicles.** Rows 1-3: TOMM20-positive vesicles. Rows 4-6, TOMM20-negative vesicles (Fig. 3). Scale bars: 0.5µm

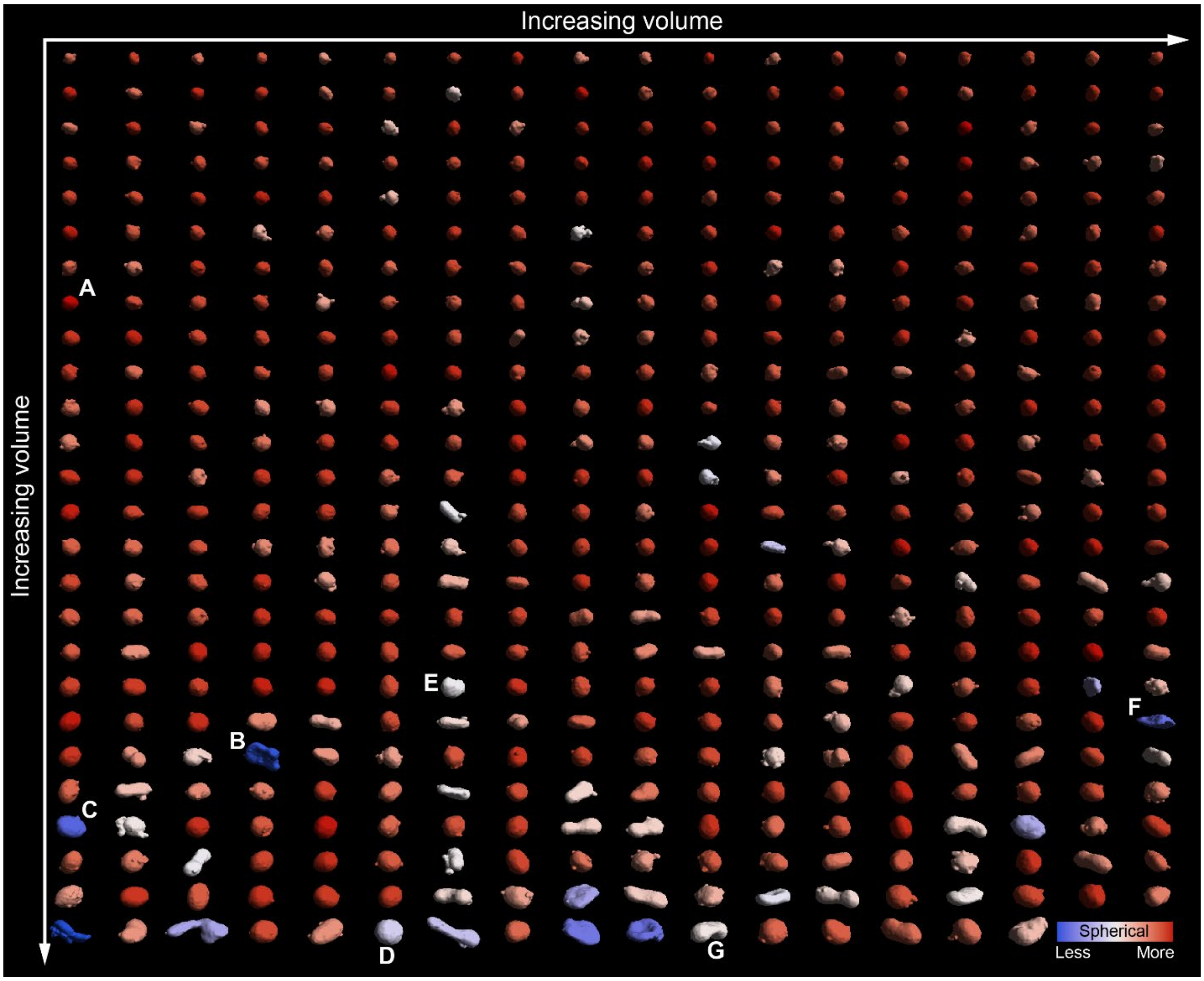


**Fig. S17.**

**FIB-SEM segmentation of peroxisomes identified by correlative cryo-SMLM.** 466 peroxisomal targeting signal (SKL) containing vesicles in two HeLa cells expressing Emerald-SKL ordered left to right and top to bottom by volume, and color coded according to their degree of deviation from spherical shape (supplementary note 8). Letters correspond to specific peroxisomes in Fig. 4.

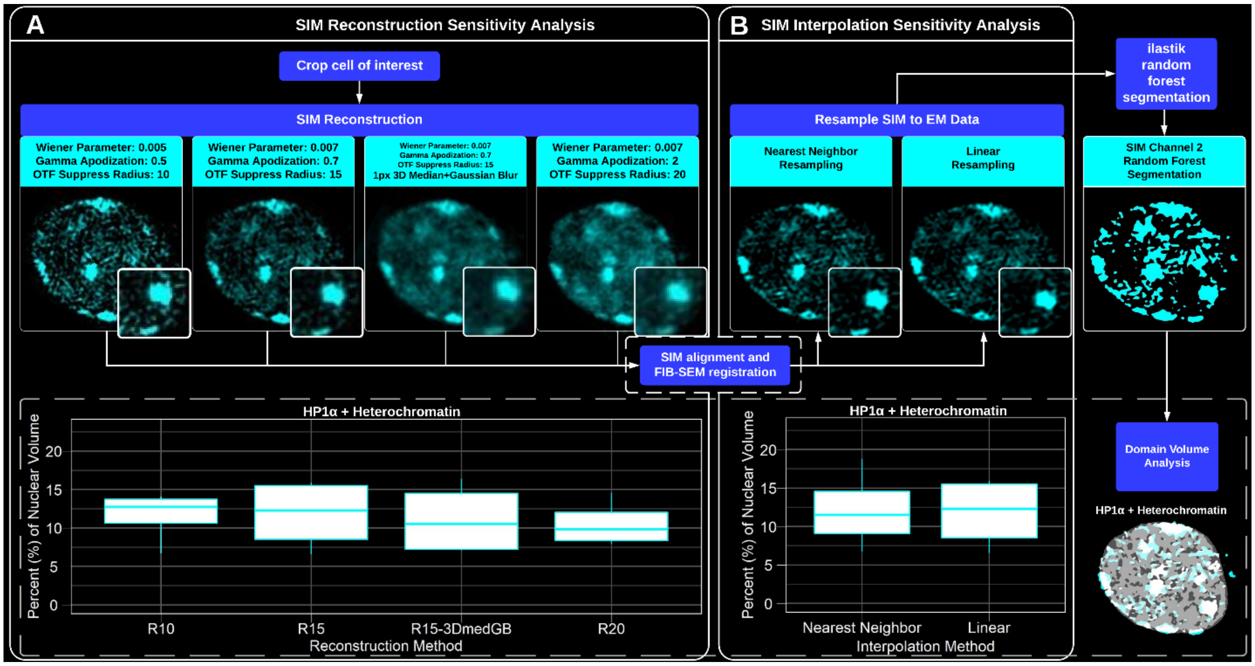


**Fig. S18.**

**Reconstruction and interpolation sensitivity analyses. (A)** SIM images of a representative CGN nucleus for different reconstruction parameters including: Wiener filter, gamma apodization, and the radius of the singularity suppression at the OTF origins. Lower dashed box contains HP1α-heterochromatin coincidence as a percentage of nuclear volume of all CGN nuclei for the parameter sets shown above. **(B)** SIM images using the second set of reconstruction parameters in (A), resampled to EM resolution using either nearest neighbor (left) or linear interpolation (right). Lower dashed box compares HP1α- heterochromatin coincidence as the percent volume for all CGN nuclei.

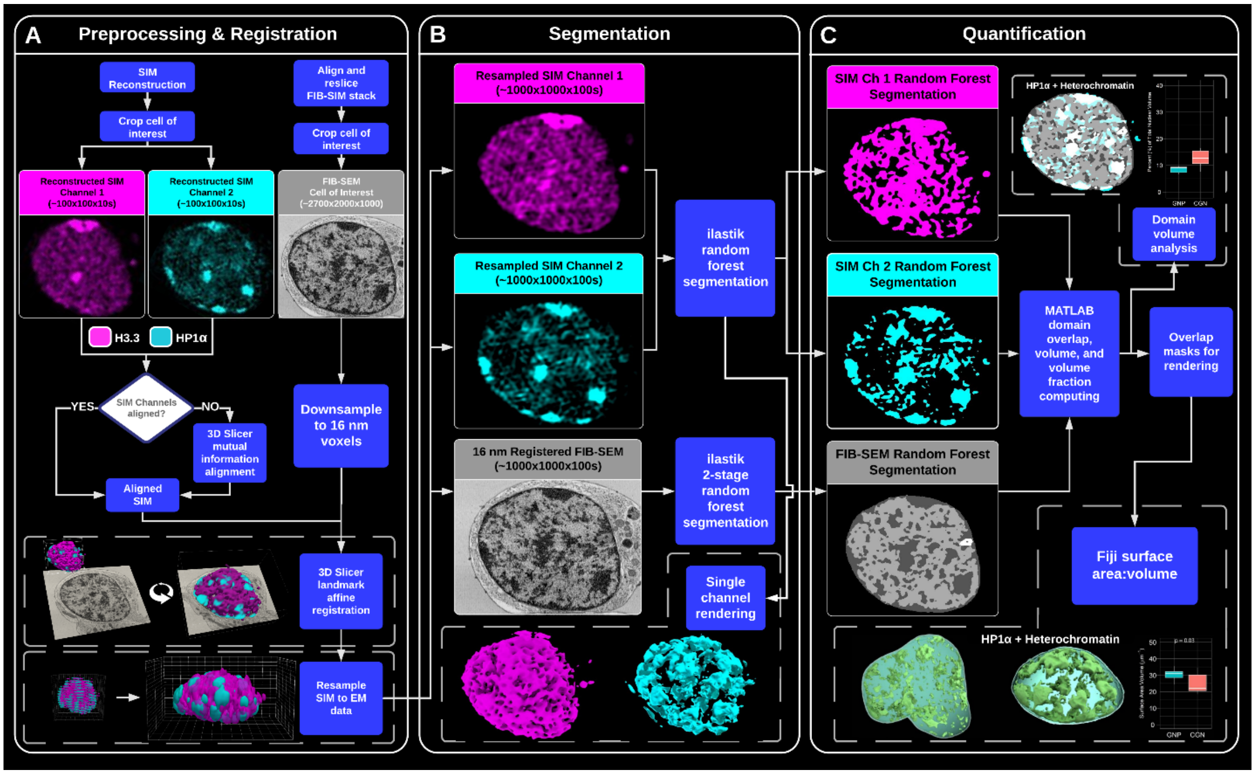


**Fig. S19.**

**Flowchart of Cryo-SIM/FIB-SEM data processing pipeline for granule neuron nuclei. (A)** Preprocessing and registration procedure for a representative granule neuron nucleus. Representative post-reconstructed SIM (H3.3 and HP1α, shown in magenta and cyan, respectively) and stack-aligned FIB-SEM (grayscale) data are shown at top. Upper dashed box presents representative renderings of the affine registration procedure (not to scale), and lower dashed box presents representative renderings of the resampling procedure (not to scale). **(B)** Orthoslices of the resampled SIM (magenta and cyan) and FIB-SEM (gray) data, along with 3D renderings of the SIM data after a 2-pixel, 32-nm Gaussian filter and binarization by a random forest algorithm (dashed box). **(C)** Orthoslices through the binarized SIM and segmented FIB-SEM data. Upper dashed box presents HP1α and heterochromatin coincidence in a representative nucleus, with box plot at right summarizing the same across all GNP and CGN nuclei. Lower dashed box contains HP1α- heterochromatin coincidence renderings for a representative GNP and CGN. The box plot at right summarizes the surface area to volume ratio measurements for this chromatin domain subtype across all CGN and GNP samples.

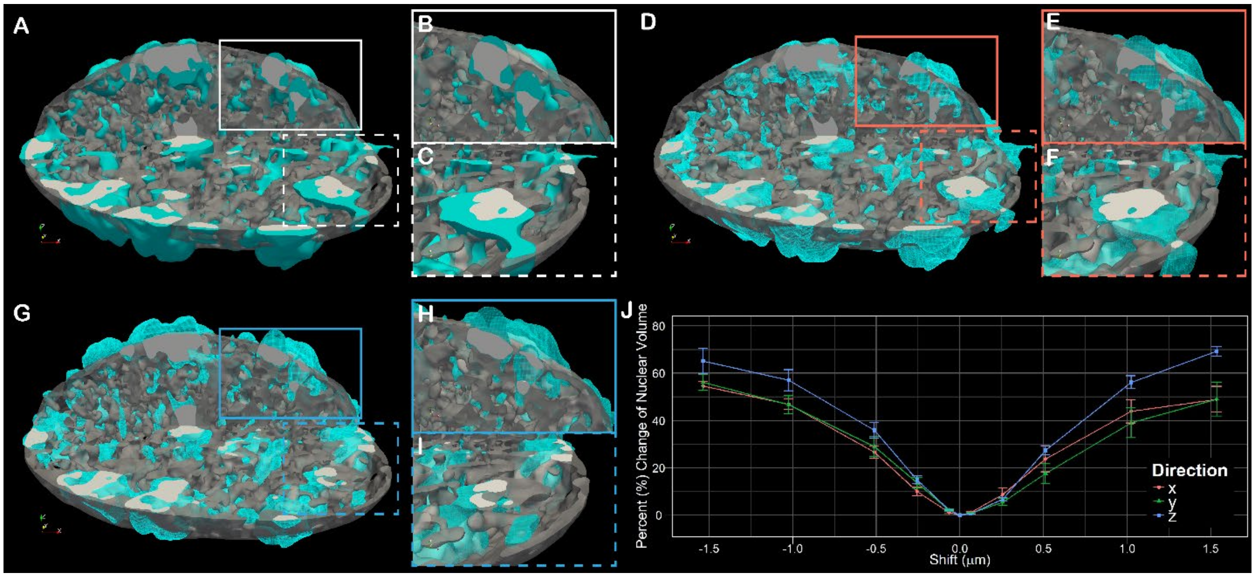


**Fig. S20.**

**Cryo-SIM/FIB-SEM registration sensitivity. (A)** Rendering of an exemplary registered CGN nucleus with random forest binarized HP1α SIM data (cyan) and the FIB-SEM segmented heterochromatin (gray). The coincidence of binary light and segmented EM data is shown in white. **(B)** Zoomed view of an XZ slice through a region of HP1α and heterochromatin overlap. **(C)** Same as (B) for an XY slice. **(D-E)** Same as (A-C) but after shifting the HP1α data 256 nm in the x direction. **(G-I)** Same as (A-C) but after shifting the HP1α data 256 nm in the z direction. **(J)** SIM registration sensitivity analysis comparing the change in the HP1α-heterochromatin percent volume averaged across two GNP and two CGN cells as the binary HP1α data are shifted in x, y, and z dimensions.

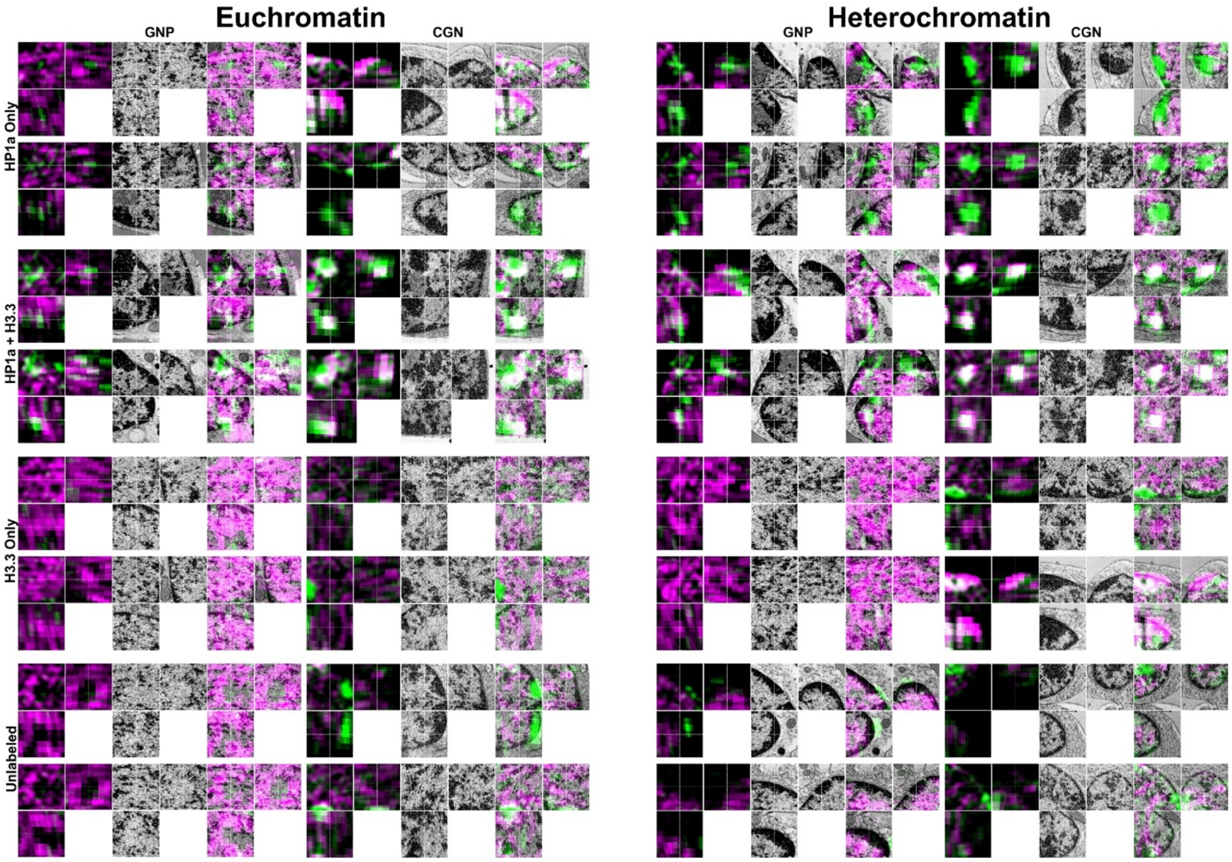


**Fig. S21.**

**Selected orthoslices of differently labeled chromatin domains.** Classical, EM-determined euchromatin and heterochromatin domains are shown on the left and right, respectively. GNP and CGN examples are shown on the left and right for each of the two domain types, with the presence or absence of nuclear domain reference proteins HP1α and H3.3 organized by column as indicated at left. Two 2x2x2 μm ROIs are shown for each labeled subdomain.

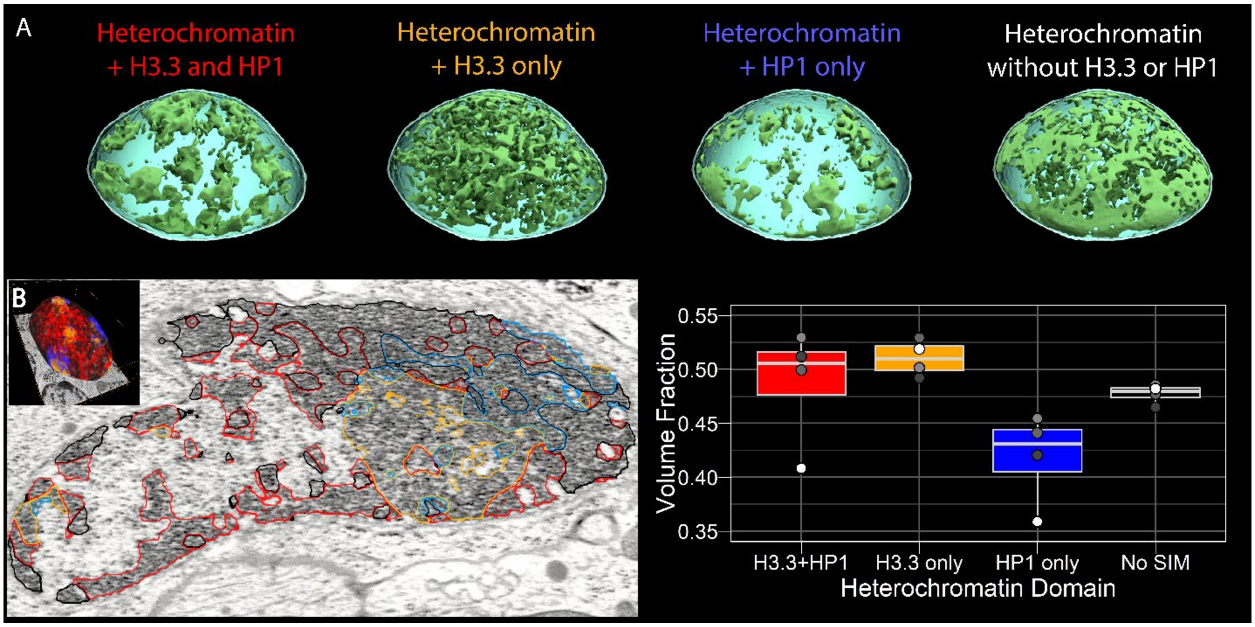


**Fig. S22.**

**Characterization of CryoCLEM heterochromatin subdomains. (A)** Isosurface renderings of distinct heterochromatin subdomains defined by either the presence or absence of H3.3 and/or HP1α. **(B)** Left: A representative FIB-SEM slice highlighting regions where either H3.3 or HP1α, or both, coincide. Border colors are defined in the box plot at right. Inset shows full volumetric rendering of one of the four cells from the analysis. Right: Box plots of the volume fraction-above-threshold for all combinations of FIB-SEM/SIM coincidence.

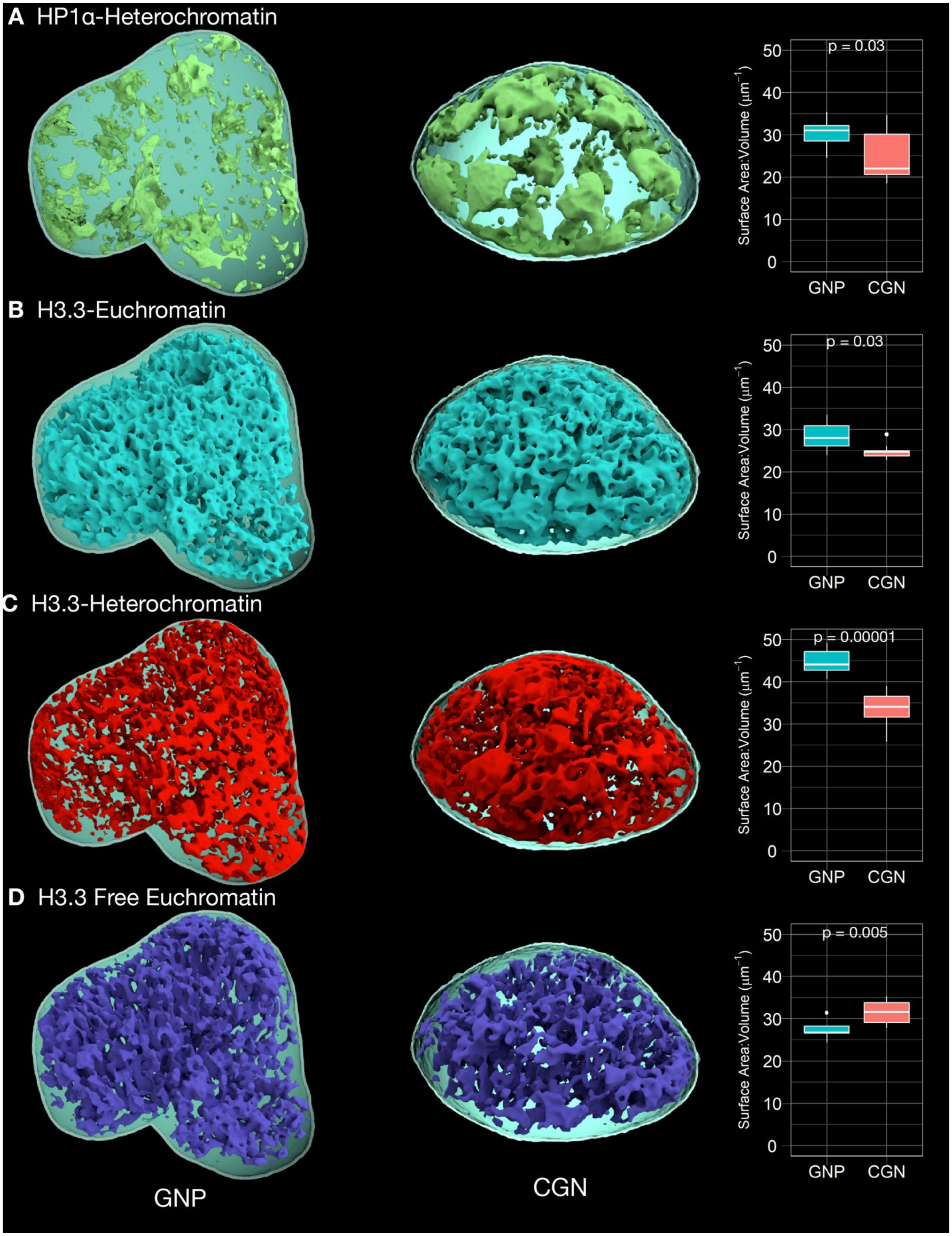
A HP1α-Heterochromatin
p = 0.03
Surface Area:Volume (μm⁻¹)
GNP
CGN
B H3.3-Euchromatin
p = 0.03
Surface Area:Volume (μm⁻¹)
GNP
CGN
C H3.3-Heterochromatin
p = 0.00001
Surface Area:Volume (μm⁻¹)
GNP
CGN
D H3.3 Free Euchromatin
p = 0.005
Surface Area:Volume (μm⁻¹)
GNP
CGN
GNP
CGN

**Fig. S23.**

**Differentiation stage-specific condensation of key chromatin subdomains identified by 3D CryoCLEM.** Representative isosurface renderings of the coincidence between the EM segmentation and the cryo-SIM signal along with box plots of the surface area to volume for all cells analyzed in Fig. 7. **(A)** HP1α and heterochromatin, **(B)** H3.3 and euchromatin, **(C)** H3.3 and heterochromatin, and **(D)** H3.3-free euchromatin.

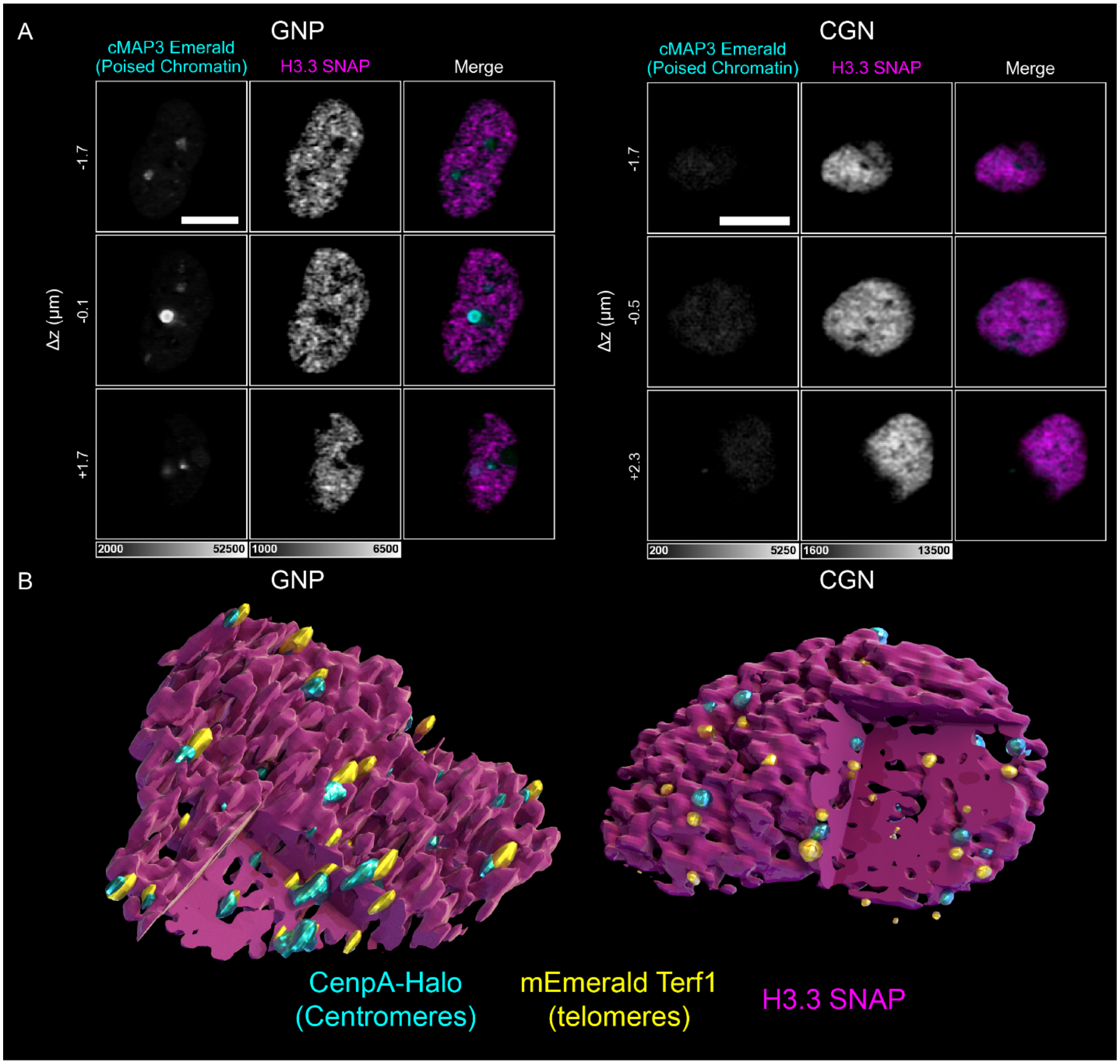


**Fig. S24.**

**Lattice light sheet SIM mode characterization of CryoCLEM chromatin domains. (A)** XY planes from LLSM SIM images volumes of a GNP and a CGN nucleus, both labeled with mEmerald-cMAP3 (cyan, poised chromatin) and SNAP-H3.3 conjugated to JF532, illustrating that H3.3 voids harbor poised chromatin (Movie S5). Scale bars: 5 μm. **(B)** Isosurface renderings of LLSM SIM data of a GNP and a CGN nucleus, both labeled with SNAP-H3.3 conjugated to JF646 (magenta, euchromatin marker), mEmerald-Terf1 (yellow, telomeres) and Halo-CENPA conjugated to JF532 (cyan, centromeres).

(A)

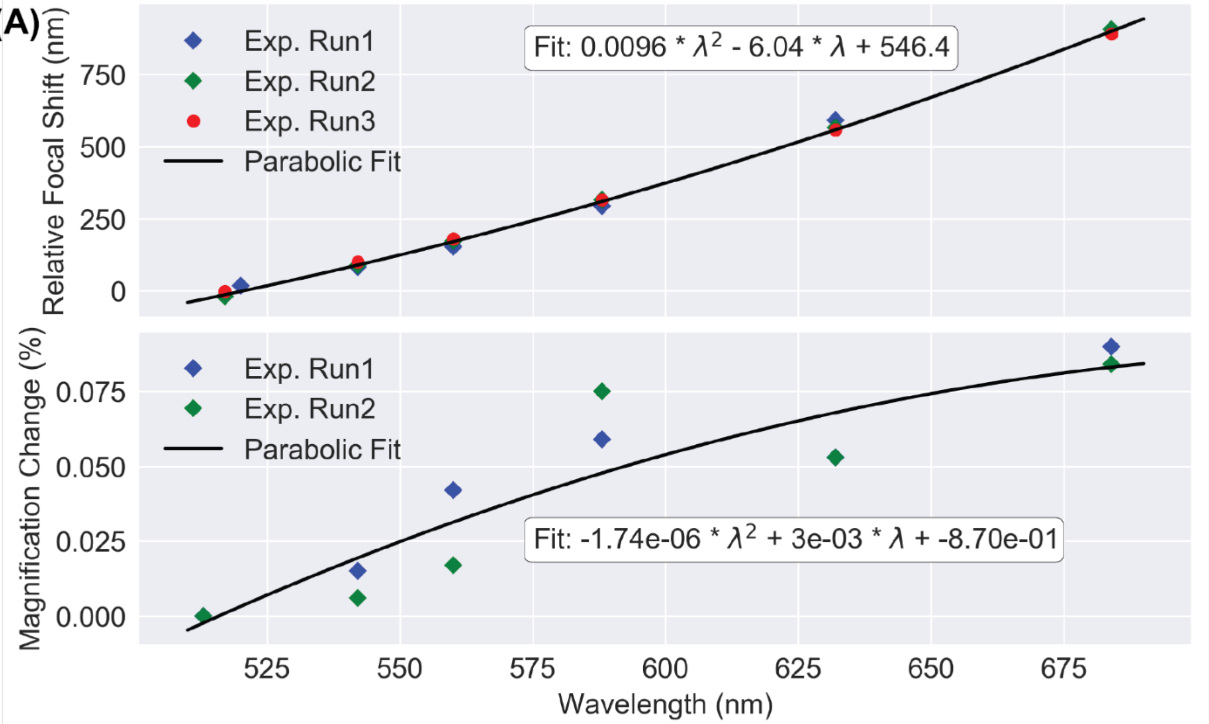


(B)

| | $\lambda_{exc}$ (nm) | Emission Filter | ΔZ (nm) |
|---|---|---|---|
| mEmerald | 488 | 488RU + 520/35 | -20.2 |
| JF525 | 532 | 532RU + 633SP | 176.3 |
| Green Bead | 488 | 488RU + 520/35 | -3.8 |
| Green Bead | 488 | 532RU + 633SP | 103.1 |
| Orange Bead | 488 | 532RU + 633SP | 197.3 |
| Orange Bead | 532 | 532RU + 633SP | 189.8 |

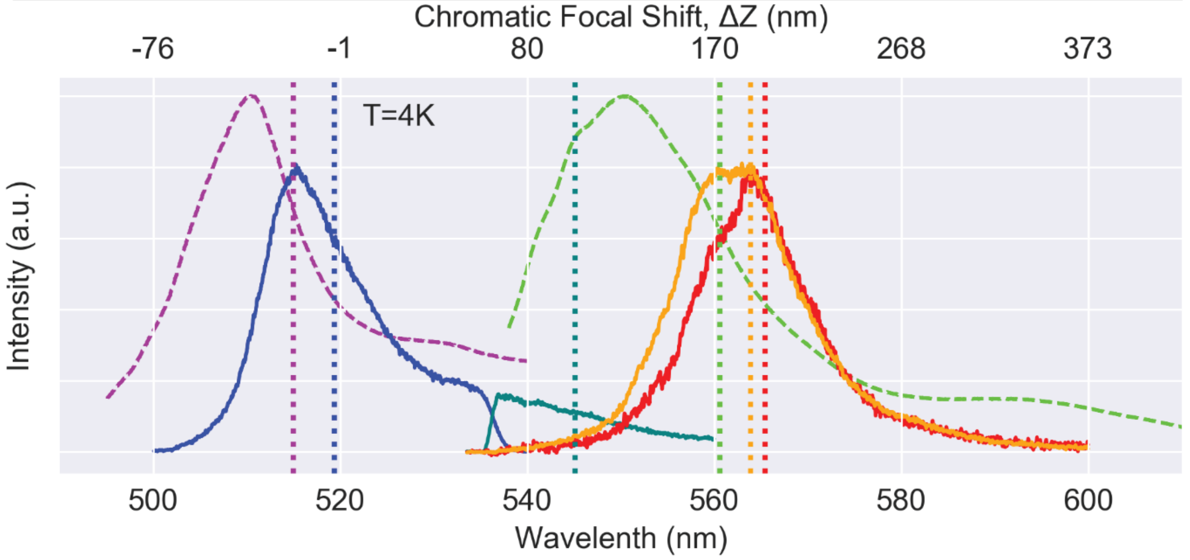

**Fig. S25.**

**Chromatic performance of the system with Nikon CFI L Plan EPI CRB 100x NA=0.85 air objective. (A)** Dependence of the relative focal plane position (top) and relative magnification (bottom) on wavelength. **(B)** Spectra of the fluorophores (dashed lines) and fluorescent beads (solid lines) at T = 4K with different excitation conditions and different emission filters. Vertical dotted lines of the same color indicate weight-averaged emission centers and focal shifts for each spectrum, the values of the focal shifts $\Delta Z$ also shown in the table.

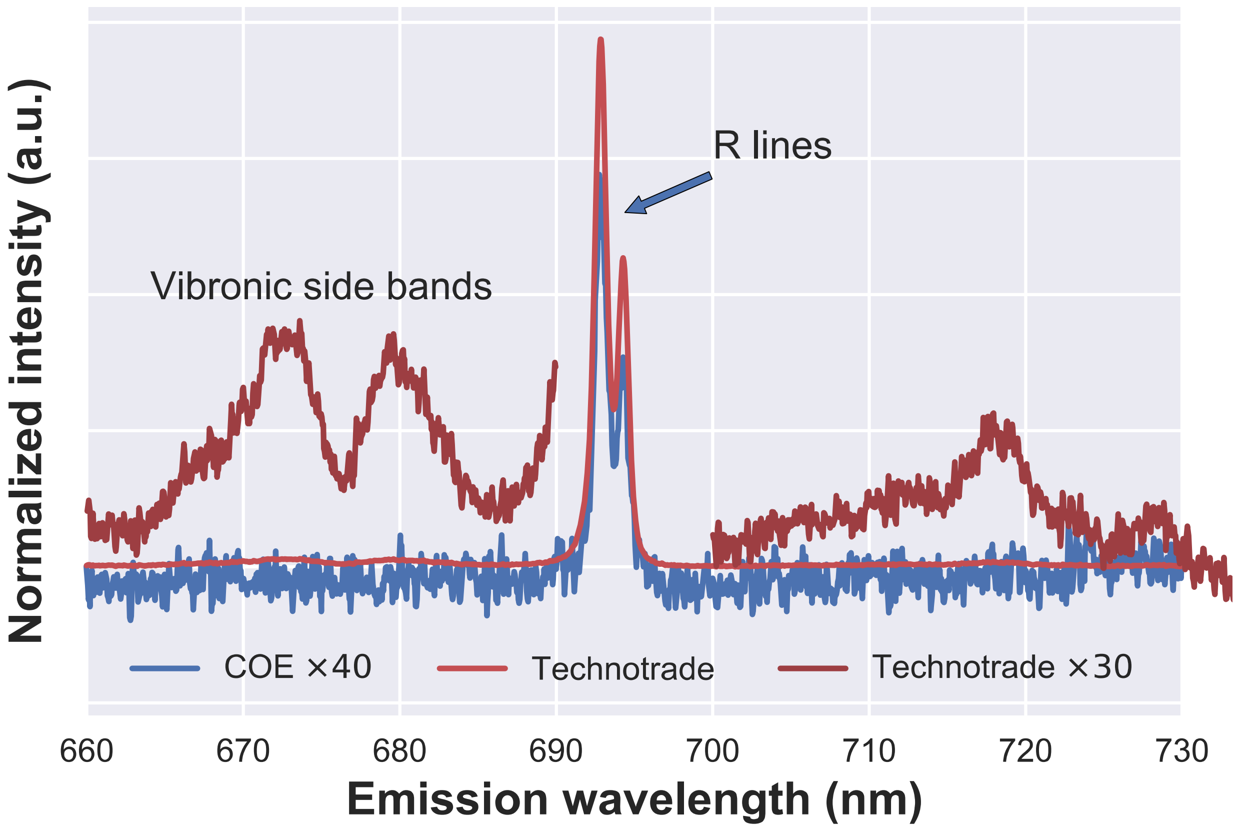


**Fig. S26.**

**Relative chromium (ruby) contamination of commercial sapphire coverslips.** Emission spectra of coverslips from two different manufacturers, under 561 nm excitation, normalized for camera exposure time, laser power and sample thickness, showing characteristic R-lines of atomic transitions in Cr. Light red line: TechnoTrade sapphire disk 500. Blue line: custom coverslip order, Nanjing Co-Energy Optical Crystal Co., Ltd, vertically magnified 40× for comparison. Dark red lines: TechnoTrade vertically magnified 30× to show vibronic side bands that can overlap with the emission of red fluorescent proteins and lead to excessive background for single molecule imaging.

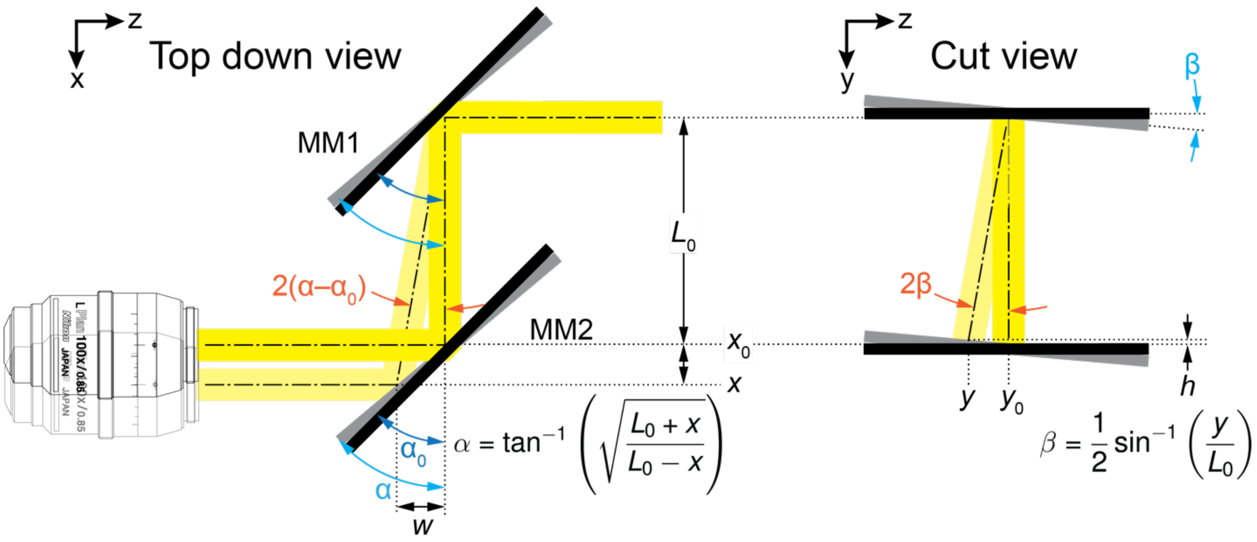


**Fig. S27.**

**Diagram illustrating how alignment of the optical path relative to the objective is maintained as the objective moves.** Top down view: when the objective moves in the direction shown from $x_0$ to $x$ in the plane of the coverslip, MM1 and MM2 must redirect the optical path (light yellow) so that it remains collinear with the axis of the objective at its new position. Angular alignment is maintained as long as MM1 and MM2 maintain the same relative angle between them. If $\alpha_0$ is the initial relative angle when the objective is at $x_0$, and $L_0$ is the distance between the pivot points of MM1 and MM2, then the relative angle α needed to keep the input beam centered when the objective moves to $x$ is given as shown above. Cut view: when the objective moves in the direction shown from $y_0$ to $y$ in the plane of the coverslip, the input beam will remain centered in the objective if the relative angle of both mirrors change by the amount β shown above. For derivations see supplementary note 3d.

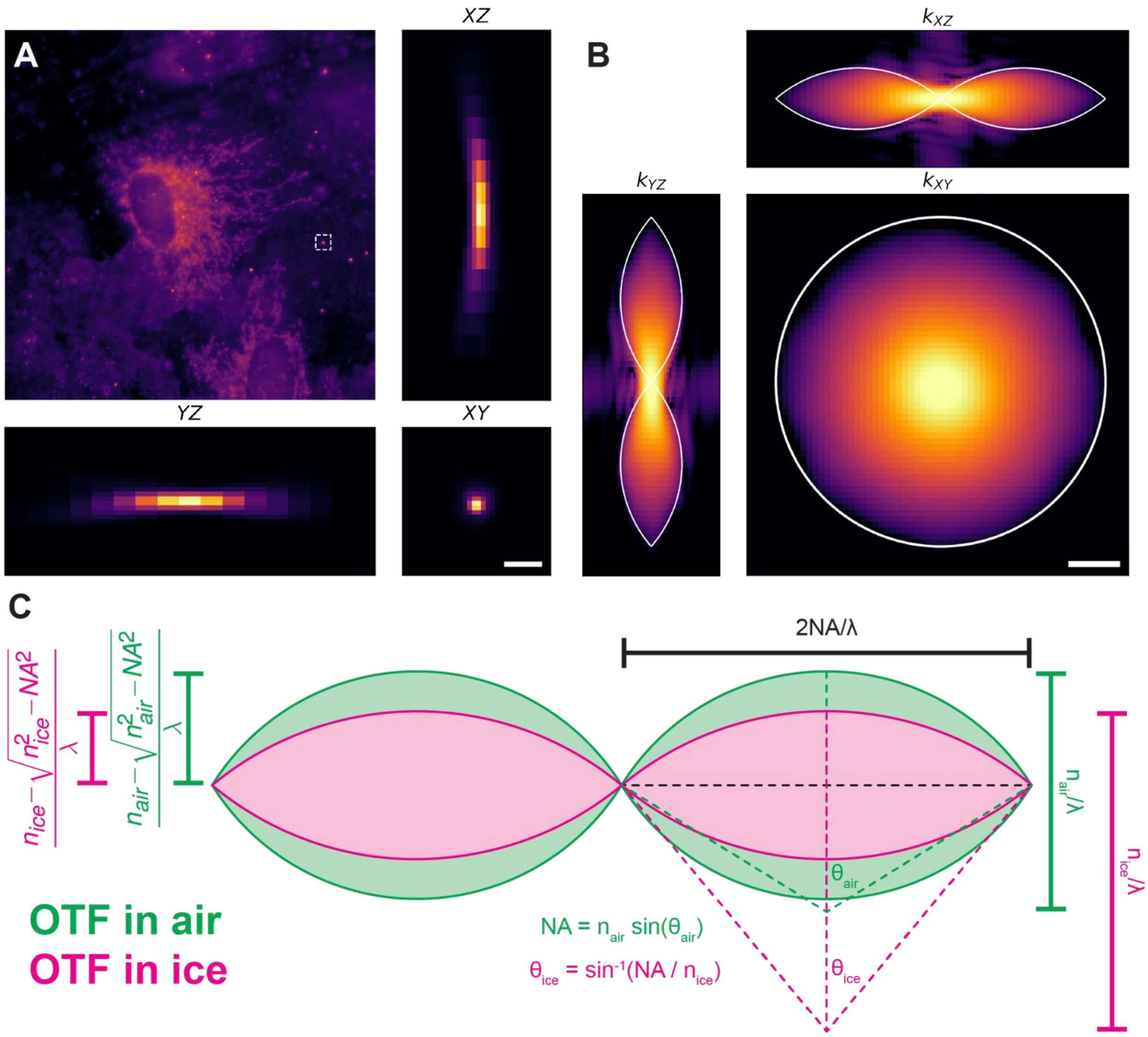


**Fig. S28.**

**Resolution characterization of the cryogenic microscope. (A)** A HeLa cell immobilized in vitreous ice expressing mEmerald-SKL (Fig. 4) surrounded by fluorescent bead fiducial markers. Max projections of the 3D PSF of the indicated bead (dashed square) along each of the three principal axes are shown. Scale bar, 1 µm. **(B)** 2D slices at principal planes of the corresponding 3D OTF. White lines indicate the theoretical maximal extent of the OTF. Scale bar, 1 $\mu m^{-1}$. **(C)** Comparison of a cut through the theoretical 3D OTF along the $k_{\rho z}$ plane for imaging in air (green) versus vitreous ice (pink). Lateral resolution is identical in the two cases, since the bending of the light rays at the ice interface due to Snell's law is exactly offset by the larger magnitude of the *k* vector in ice. Axial resolution, however, is higher in air than in ice, as shown by the scale bars and equations at left (equations can be derived from geometrical arguments based on the figure).

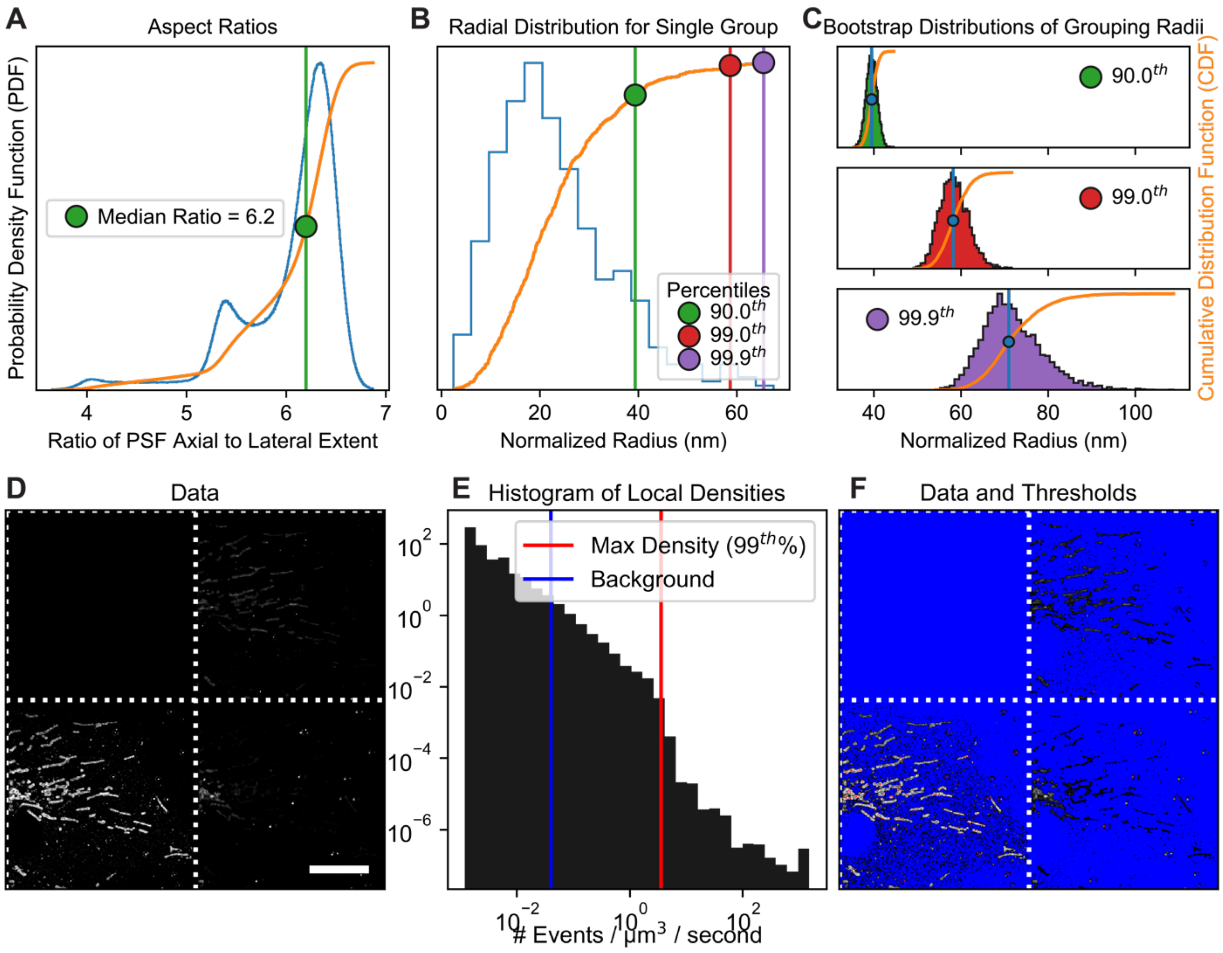


**Fig. S29.**

**Determining optimal grouping parameters from experimental data. (A)** Histogram (blue, left axis) and cumulative distribution (orange, right axis) of the PSF aspect ratios for all non-fiducial localizations in an example data set. The z-scaling parameter used in determining the axial grouping radius is given by the median of the distribution of aspect ratios (green line). **(B)** Histogram and cumulative distribution of normalized radii for synthetic groups (supplementary note 5b) with the $90^{th}$, $99^{th}$, and $99.9^{th}$ percentiles indicated. **(C)** Bootstrap distributions of the percentiles shown in (B) with median values indicated. **(D-F)** Space time event density is determined from the data (the 3D histogram renderings are shown as a mosaics) which, in combination with the normalized grouping radius, can be used to determine the grouping gap (supplementary note 5b and fig. S30). Scale bar: 10 µm.

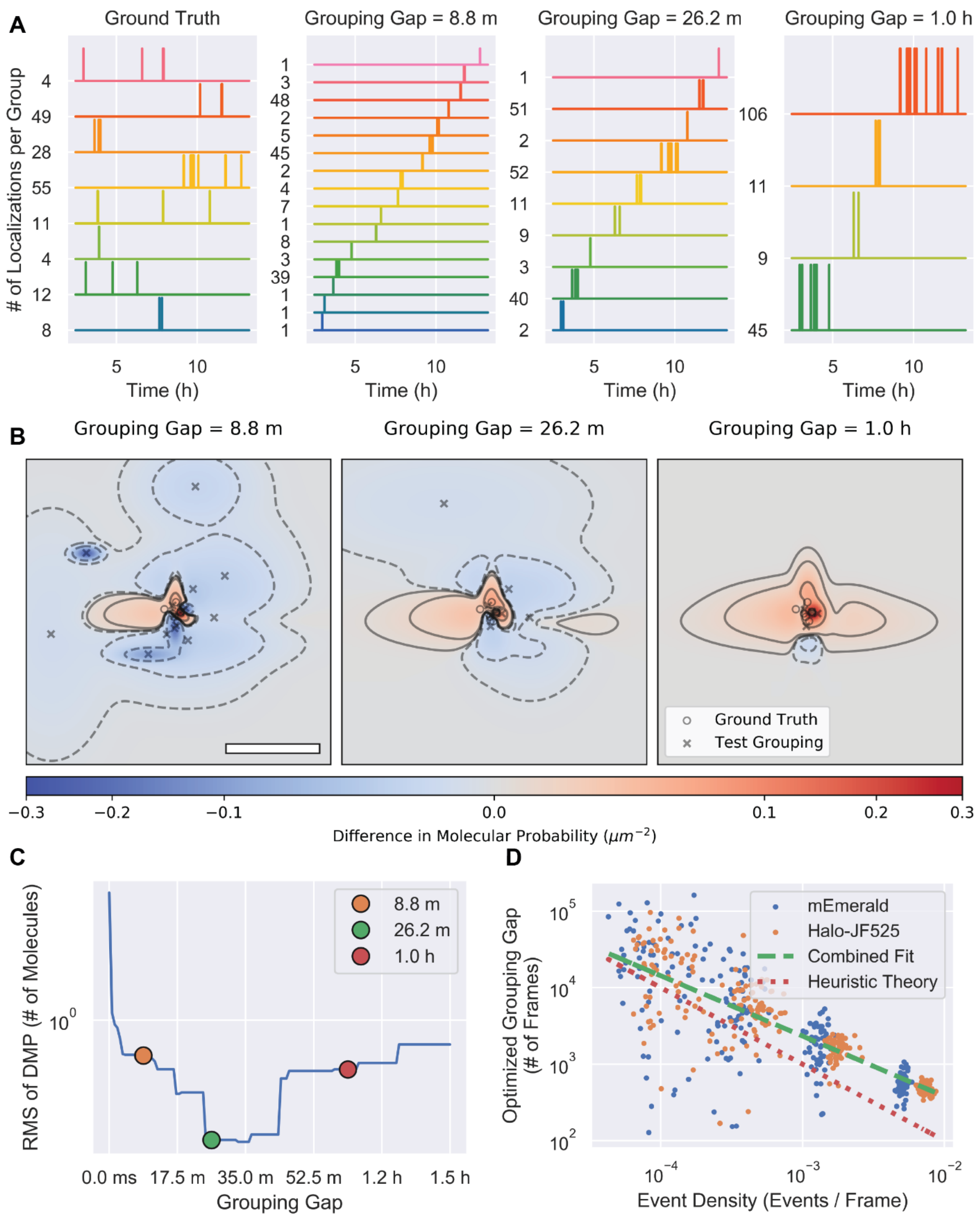
A
Ground Truth
Grouping Gap = 8.8 m
Grouping Gap = 26.2 m
Grouping Gap = 1.0 h
### of Localizations per Group
Time (h)
B
Grouping Gap = 8.8 m
Grouping Gap = 26.2 m
Grouping Gap = 1.0 h
Ground Truth
Test Grouping
Difference in Molecular Probability ($\mu m^{-2}$)
C
RMS of DMP (# of Molecules)
8.8 m
26.2 m
1.0 h
Grouping Gap
D
Optimized Grouping Gap (# of Frames)
mEmerald
Halo-JF525
Combined Fit
Heuristic Theory
Event Density (Events / Frame)

**Fig. S30.**

**Determining the optimal grouping gap from the data. (A**) Left column: a set of eight ground truth single molecule time traces (fig. S8 and supplementary note 4a) showing the number of localizations detected in each. Remaining columns: the results of grouping this set of single molecules with three arbitrary grouping gaps: 8.8 minutes, 26.2 minutes and 1 hour. **(B)** Differences in molecular probability (DMP) distributions between the exact grouping and grouping by the three arbitrary gaps in (A). Individual exact and arbitrary grouped localizations are indicated by circles and crosses, respectively. Scale bar, 50 nm. **(C)** Plot of the root mean squared DMP for different grouping gaps. **(D)** Scatter plot of optimized grouping gaps for mEmerald (blue dots) and Halo-JF525 (orange dots) as a function of event density. A line corresponding to the heuristic theory (supplementary note 5b) is shown as a dotted line and a power-law fit to both optimized data sets is shown as a green dashed line.

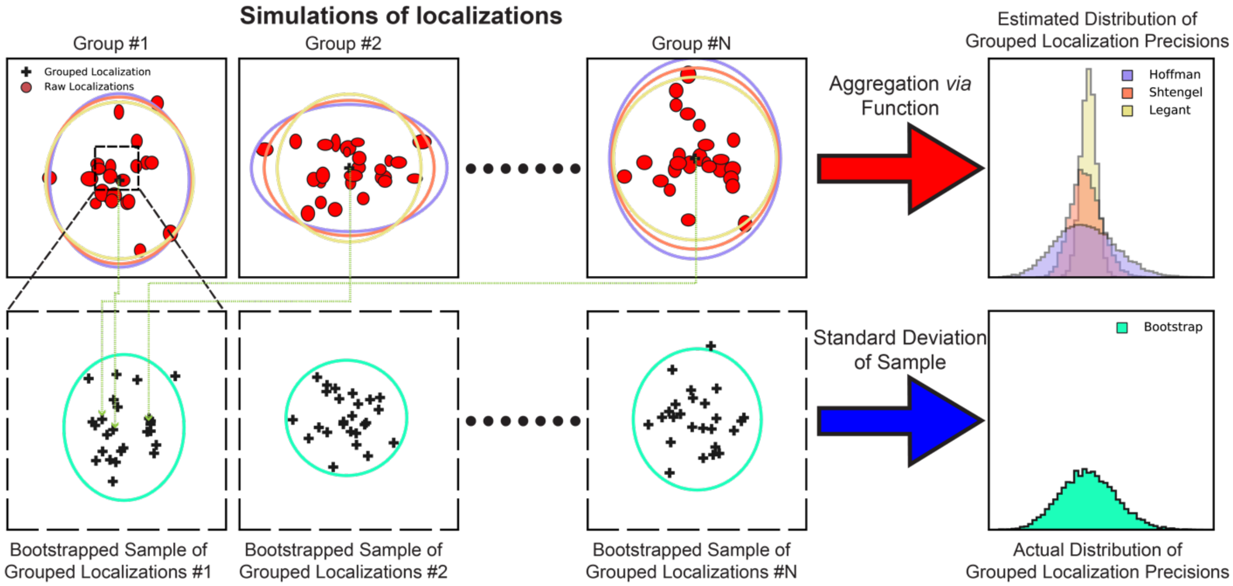


**Fig. S31.**

**Determining the localization precision of grouped emissive events.** Top row: simulated groups of localizations (red ellipses indicate position and precisions) along with the grouped position (black cross) and estimated grouped precision (magnified 60X) shown as large colored ellipses for three different grouping equations (supplementary note 5c). Bottom row: bootstrap samples of grouped positions along with the precision (standard deviation) of the sample (green ellipse, magnified 10X). Far right column, top: histograms of localization precisions in the horizontal direction for the three grouping functions. Far right column, bottom: ground truth distribution derived from bootstrap sampling.

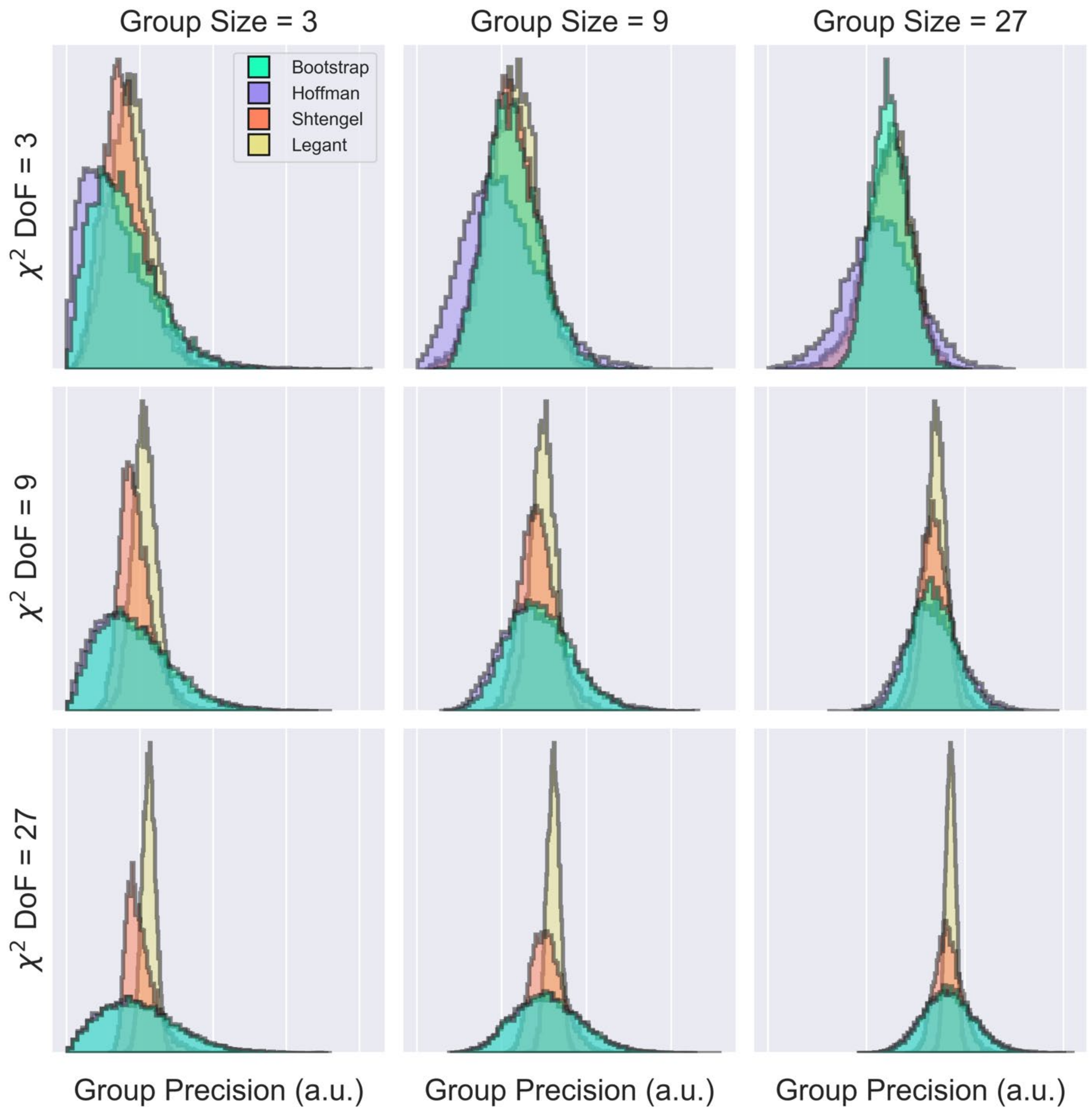


**Fig. S32A.**

**Comparing alternative methods of estimating localization precision for grouped localizations, synthetic data.** Distributions of grouped localization precisions as estimated by three different methods: Legant LLS PAINT (yellow, eq. 17, (*112*)); Shtengel iPALM (red, eq. 16, (*89*)); and Hoffman cryo-SMLM (purple, Eq. 15, supplementary note 5c). Green histograms show the bootstrap estimations (Fig. S31) of the distribution of precision. Each column and row present data from simulations with different numbers of localizations per group and different values for the degrees of freedom ($k$) of the chi-squared distribution ($\chi^2(k)$) from which the localization precisions are sampled. See also supplementary note 5. Note that the method developed here, Hoffman cryo-SMLM (purple), most closely resembles the result of ground truth simulations (green).

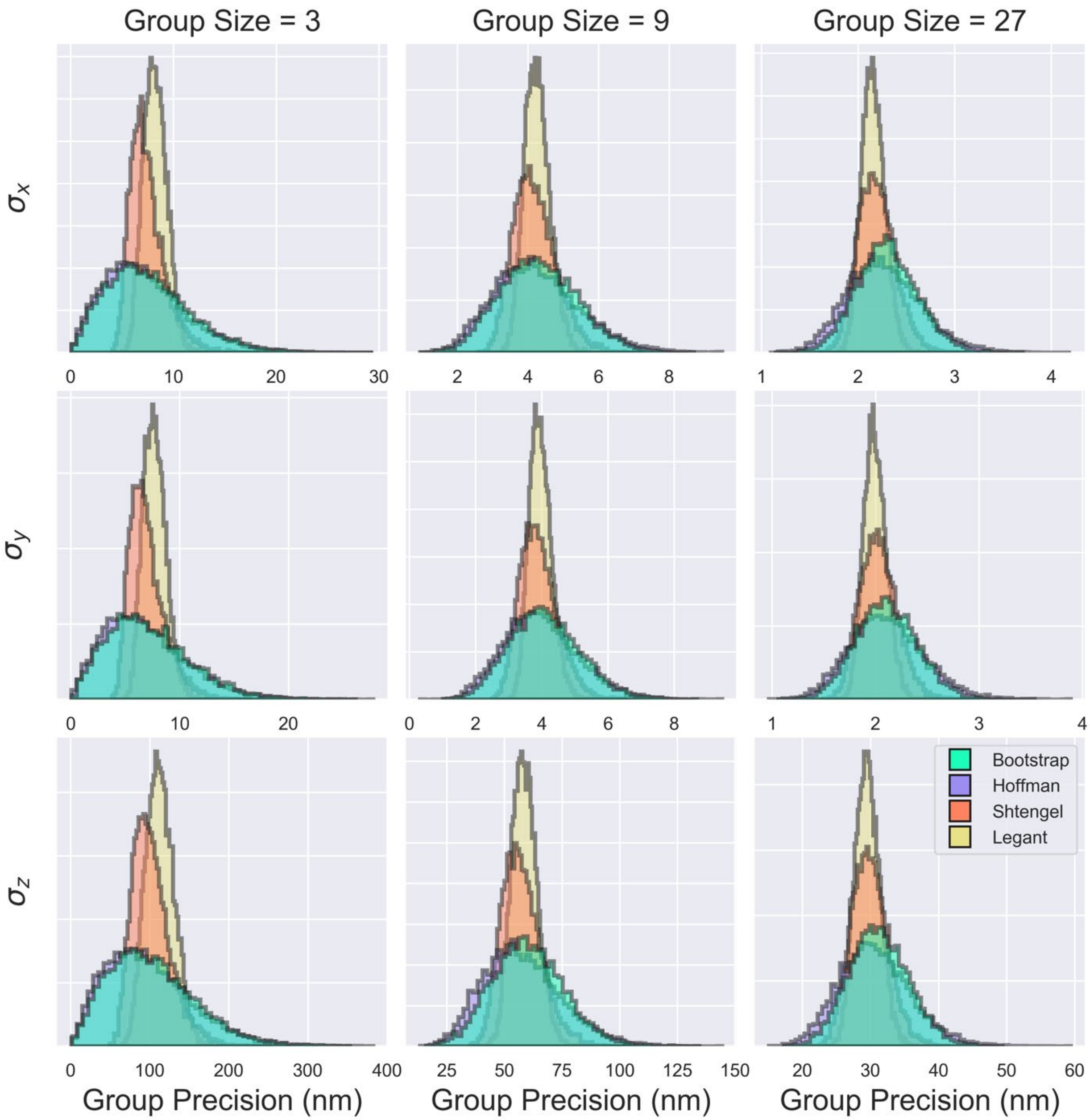


**Fig. S32B.**
**Comparing alternative methods of estimating localization precision for grouped localizations, Halo-Sec61β (JF525) data.**

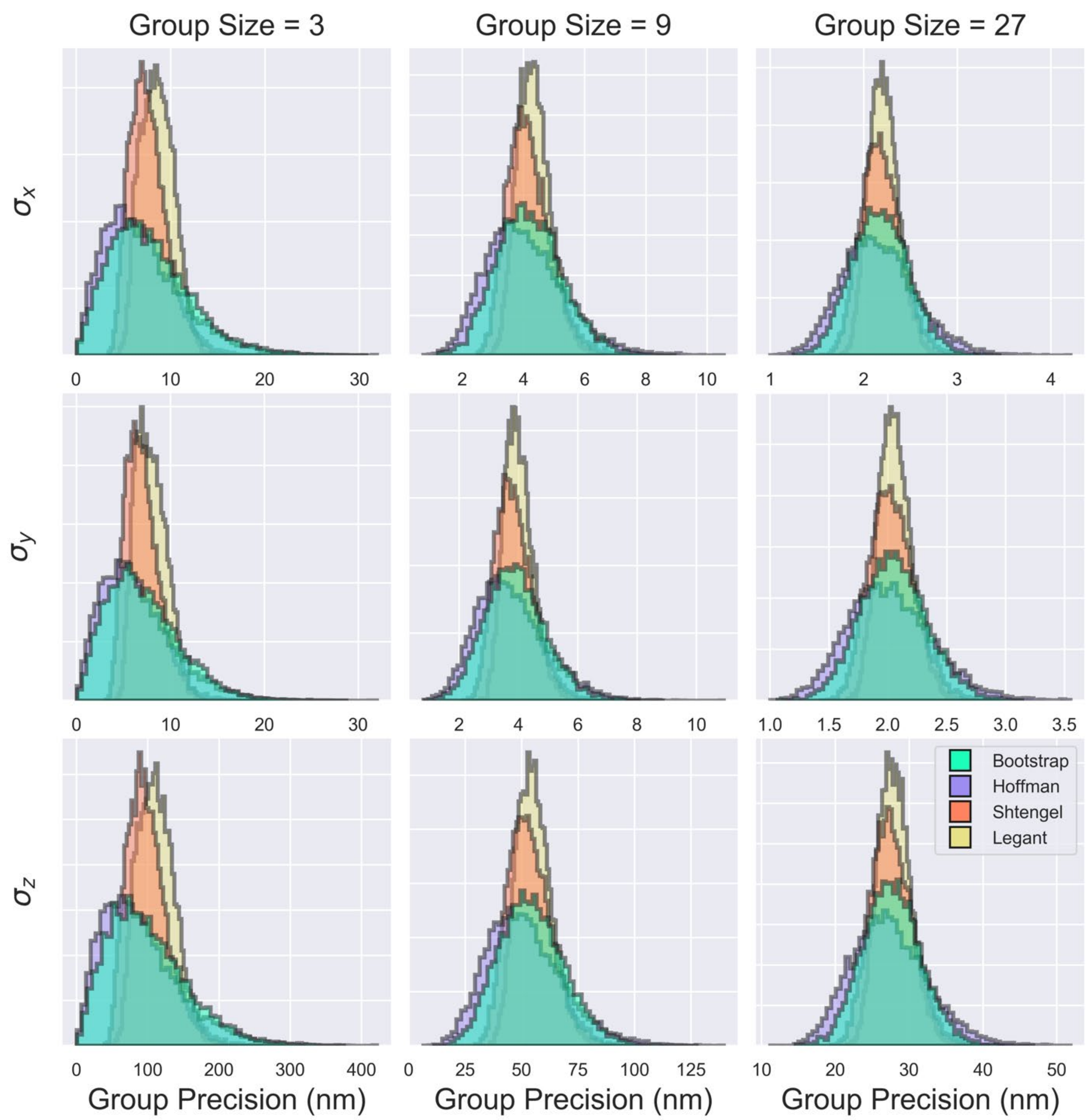


**Fig. S32C.**
**Comparing alternative methods of estimating localization precision for grouped localizations, mEmerald-Sec61β data.**

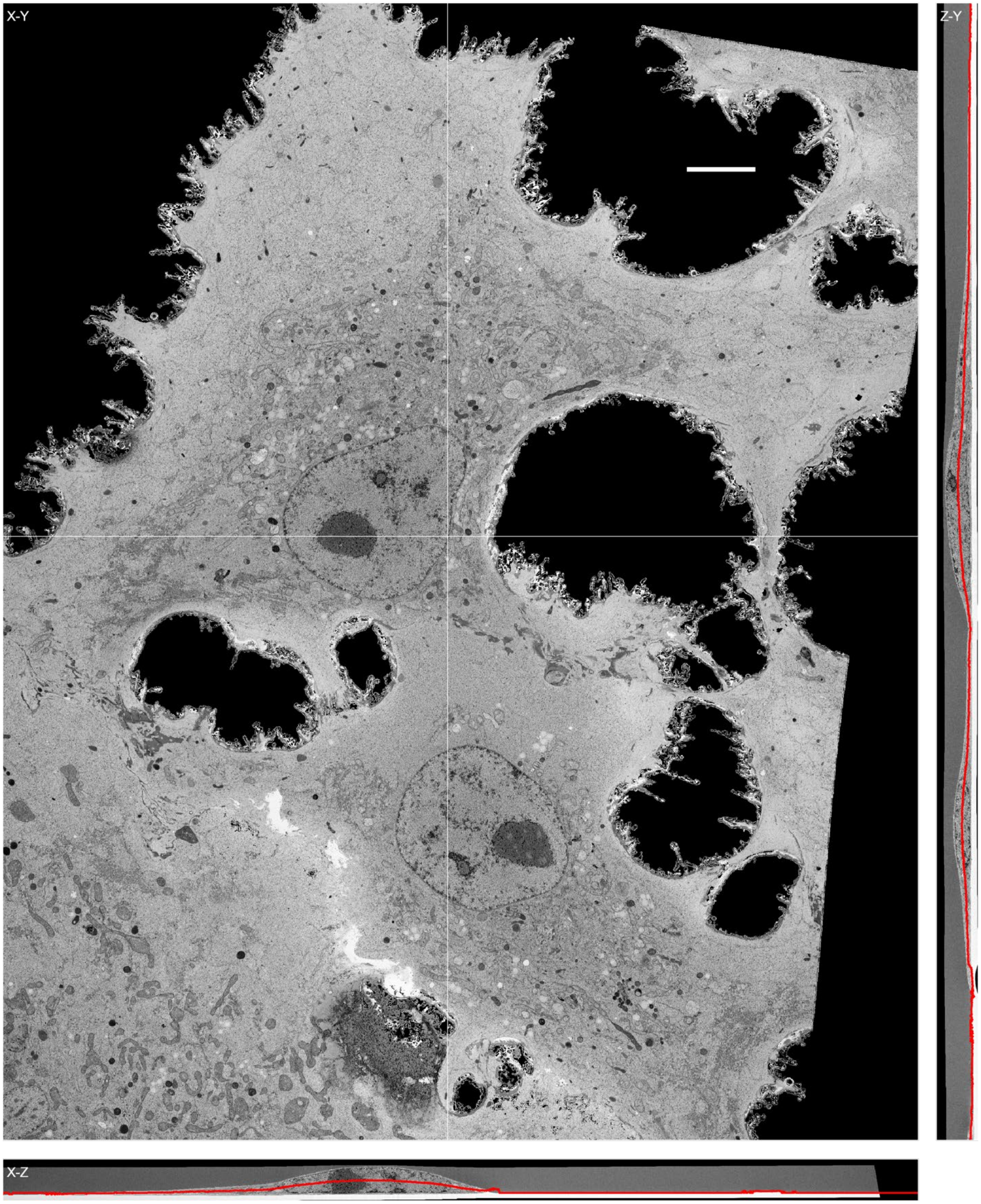
X-Y
Z-Y
X-Z

**Fig. S33.**

**Definition of mid-cell slice.** A representative "mid-cell" XY slice from the FIB-SEM volume of a cultured COS7 cell (Fig. 3). The 2D mid-cell pixel value at any (x, y) location is given by the value at the 3D volumetric pixel at the z-coordinate that corresponds to half the height of the cell at that location. ZY (right) and XZ (bottom) cross-sections are taken at positions indicated by white lines in the X-Y mid-cell slice. The red lines in ZY and XZ cross-sections indicate the z-coordinate of the mid-cell slice.

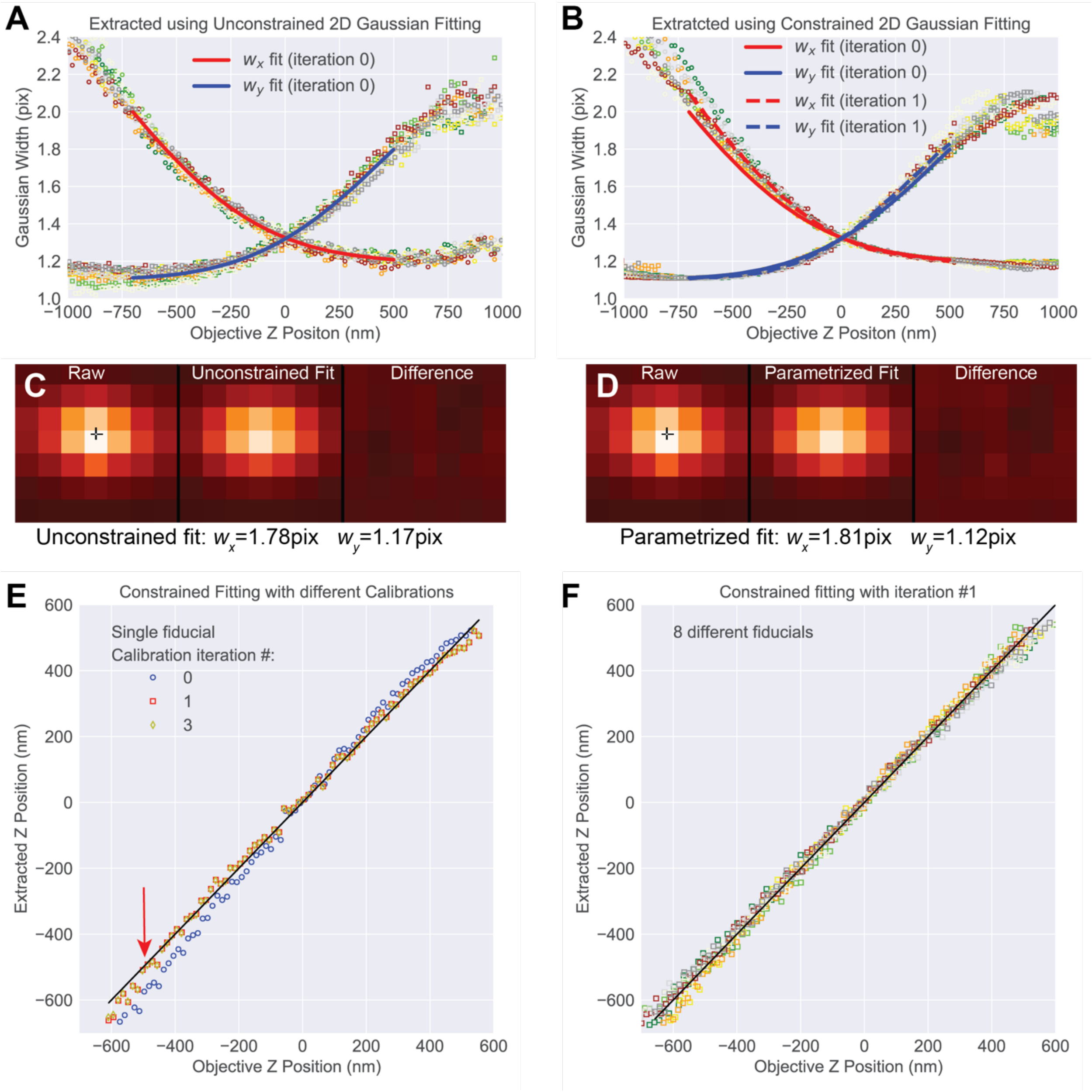


**Fig. S34.**

**Determining the Z coordinate in SMLM using astigmatic 2D Gaussian fitting. (A)** Gaussian widths from unconstrained fitting as functions of Z position for eight different fiducials (colored square markers). Fits of the x and y widths to 5$^{th}$ order polynomials are shown as solid lines. **(B)** Same as (A) but for constrained fitting, solid and dashed lines are polynomial fits from the initial step (A) and the 1$^{st}$ iteration, respectively. **(C)** Image of a selected fluorescent bead ~500nm below the focal plane (left), with unconstrained 2D Gaussian fit of the image (center), and the residuals (right). **(D)** Same as (C) for the 1$^{st}$ iteration of the constrained fit. **(E)** Z coordinate of the fiducial in (C) and (D) versus Z position for iteration 0 (blue circles), iteration 1 (red squares), and iteration 3 (yellow diamonds). Unity slope is indicated by a black line and the red arrow indicates the data in (C) and (D). **(F)** Same as (E) but for eight fiducials showing only the 1st iteration, as used for all the data in the paper.

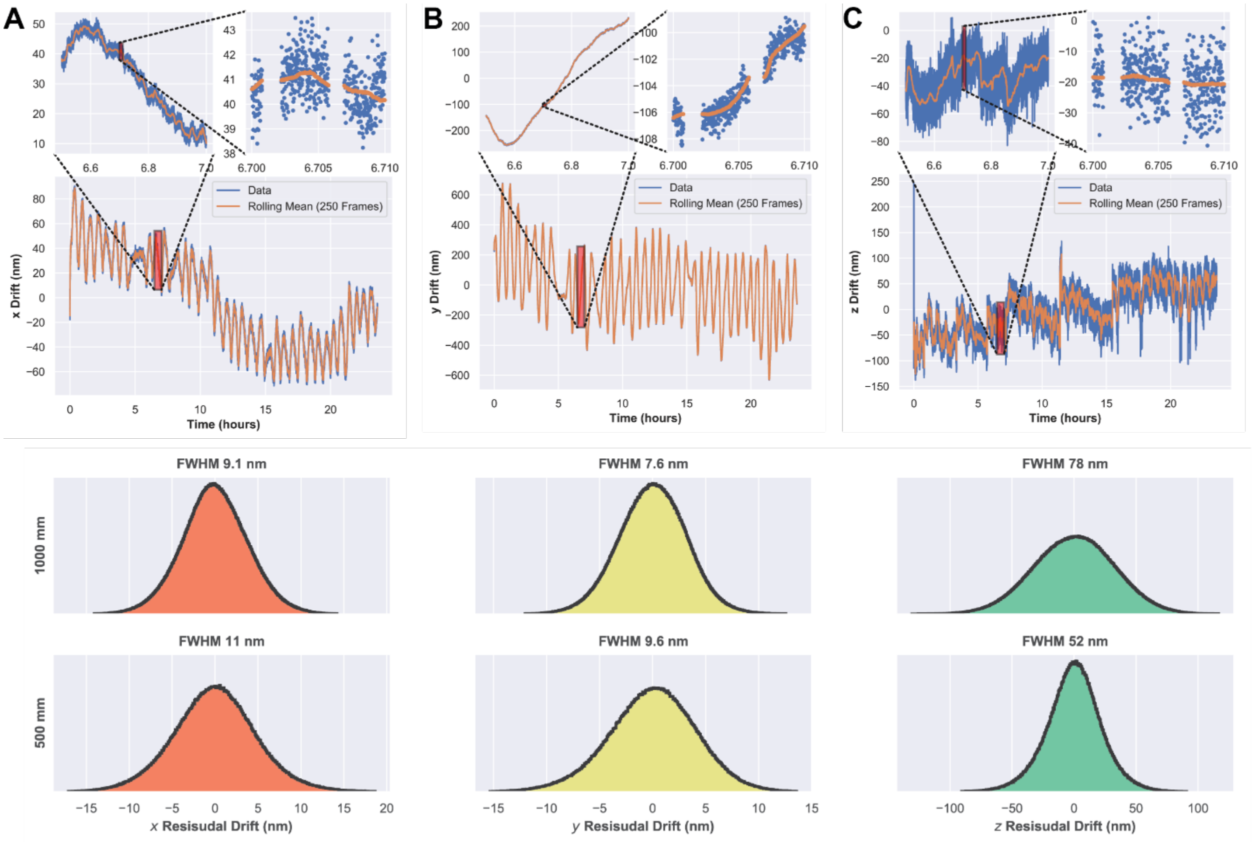


**Fig. S35.**

**Residual sample drift during cryo-SMLM acquisition, implications for resolution.** Top Row: Typical drift in the lateral **(A)** x, **(B)** y and axial **(C)** z directions, each at three different temporal magnifications. Bottom Row: Residual deviations from the mean for the five brightest fiducials in each data set in the x (orange), y (yellow) and z (green) directions for each of the astigmatism-generating cylindrical lenses (1000 mm or 500 mm focal length (FL)) used in this work. These residuals set lower bounds on the achievable localization accuracy in each direction. The 500 mm FL lens offers 33% better axial precision, but the 1000 mm lens yields 20% better lateral precision.

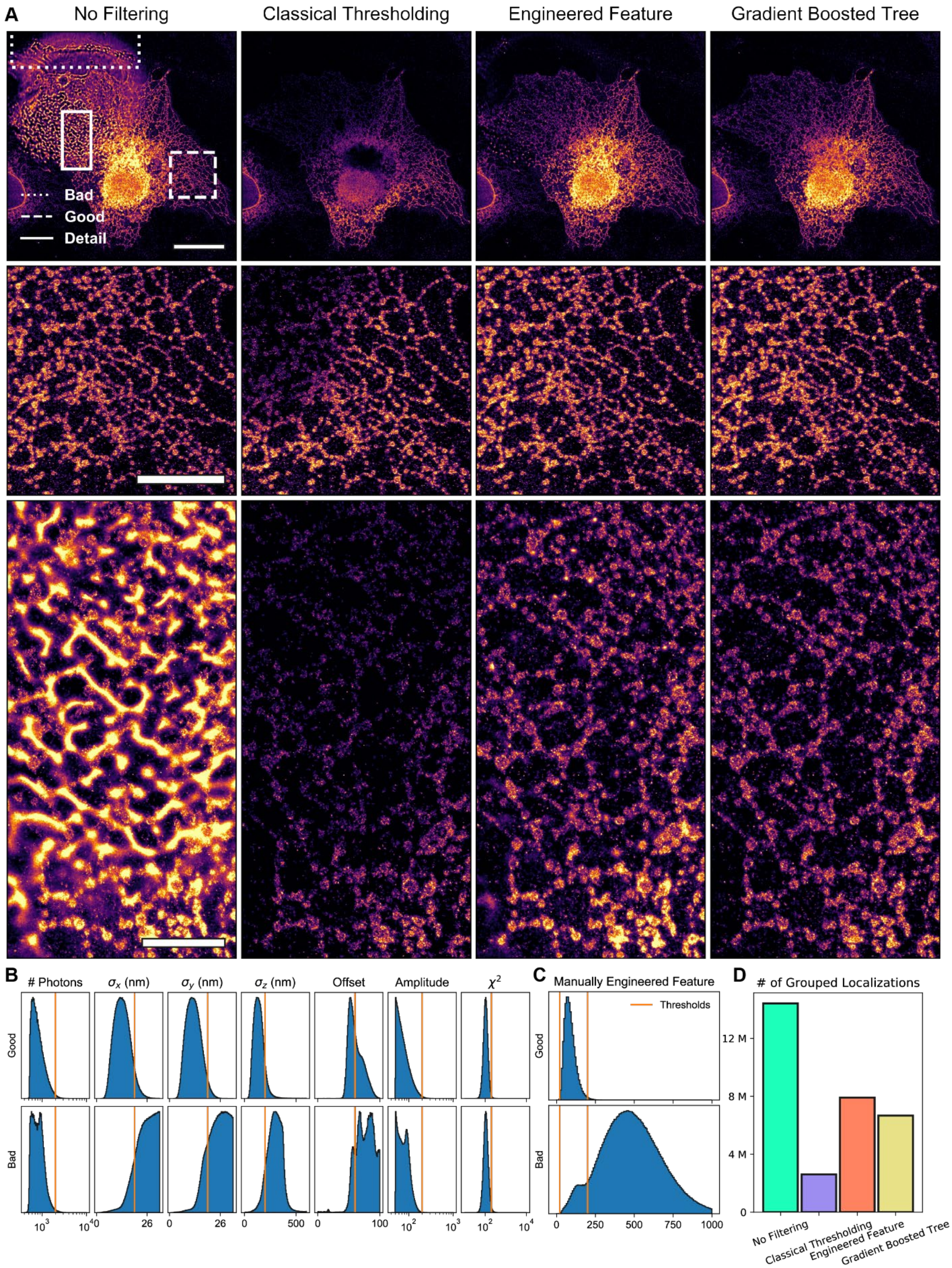
A
No Filtering
Classical Thresholding
Engineered Feature
Gradient Boosted Tree
Bad
Good
Detail
B
### Photons
$\sigma_x$ (nm)
$\sigma_y$ (nm)
$\sigma_z$ (nm)
Offset
Amplitude
$\chi^2$
Good
Bad
C
Manually Engineered Feature
Thresholds
D
### of Grouped Localizations
No Filtering
Classical Thresholding
Engineered Feature
Gradient Boosted Tree

**Fig. S36.**

**Comparison of methods for filtering poor localizations from SMLM data. (A)** An exemplary data set with contamination due to out of focus fluorescent detritus. First column: original data set, second column: data after filtering by setting thresholds on various localization parameters (cf. B), third column: data after thresholding on a manually engineered feature ($\chi^2 \times \mathrm{offset/amplitude}$), fourth column: data after filtering with a trained Gradient Boosted Tree classifier. Scale bars, 15, 5, and 3 μm. The second and third row show data from the "Good" and "Detail" insets indicated in the first image. **(B)** Classical thresholds were chosen by examining localization parameters in a "Good" area vs a "Bad" area (cf. A). **(C)** A similar strategy was used with the manually engineered feature. **(D)** The number of grouped localizations used to generate each image in the top row of **(A)**. We decided to use the gradient boosted tree method as it was the most robust and required the least human intervention, therefore avoiding human biases.

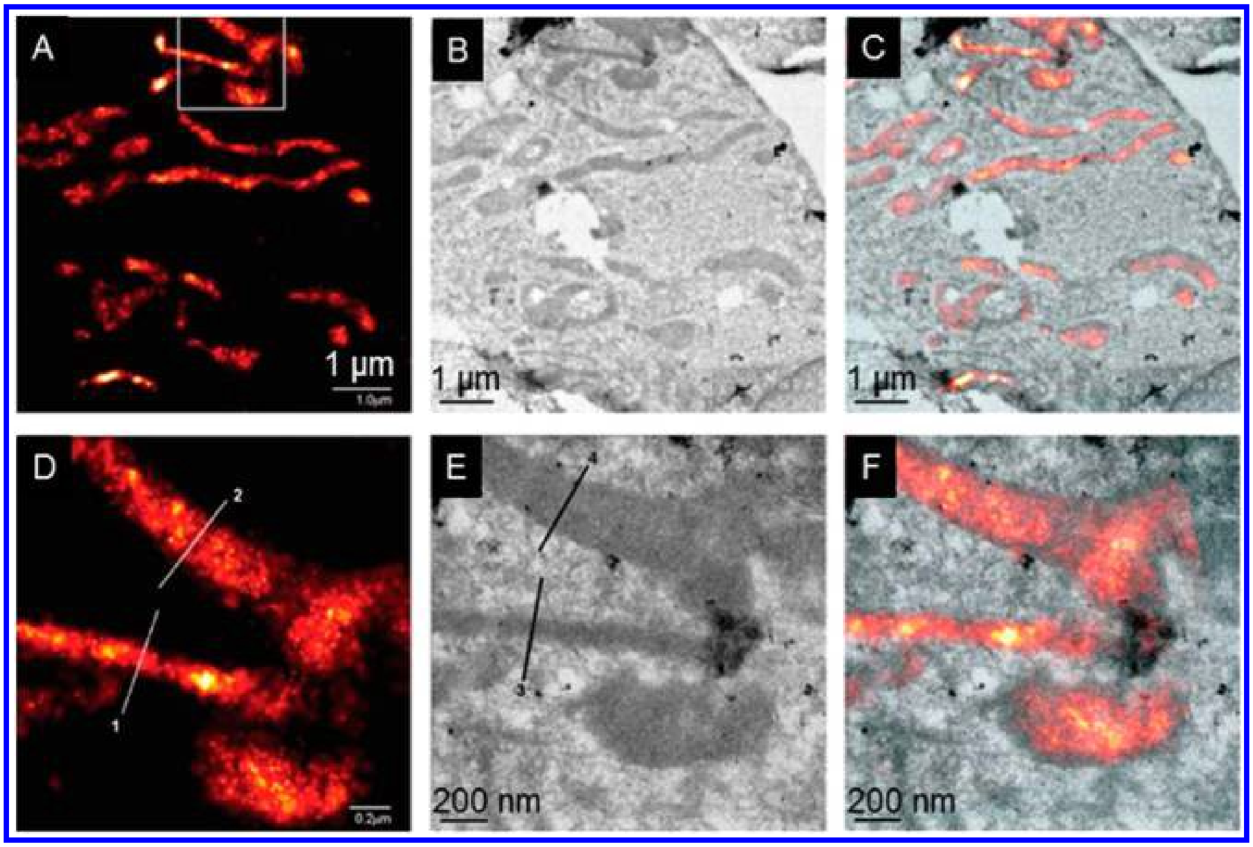


**Fig. S37.**

**Correlative PALM−TEM as an initial validation of PALM.** Correlative **(A)** PALM, **(B)** TEM, and **(C)** PALM/TEM overlay images of mitochondria in a cryoprepared thin section from a COS-7 cell expressing dEosFP-tagged cytochrome c oxidase import sequence. **(D−F)** Zoom-in of panels A−C corresponding to the box in panel A, showing that the matrix reporter molecules visualized by PALM extend up to the outer mitochondrial membrane visualized by TEM. Scale bars: 1.0 µm (A−C), 200 nm (D−F). Adapted from refs (*17*) and (*94*).

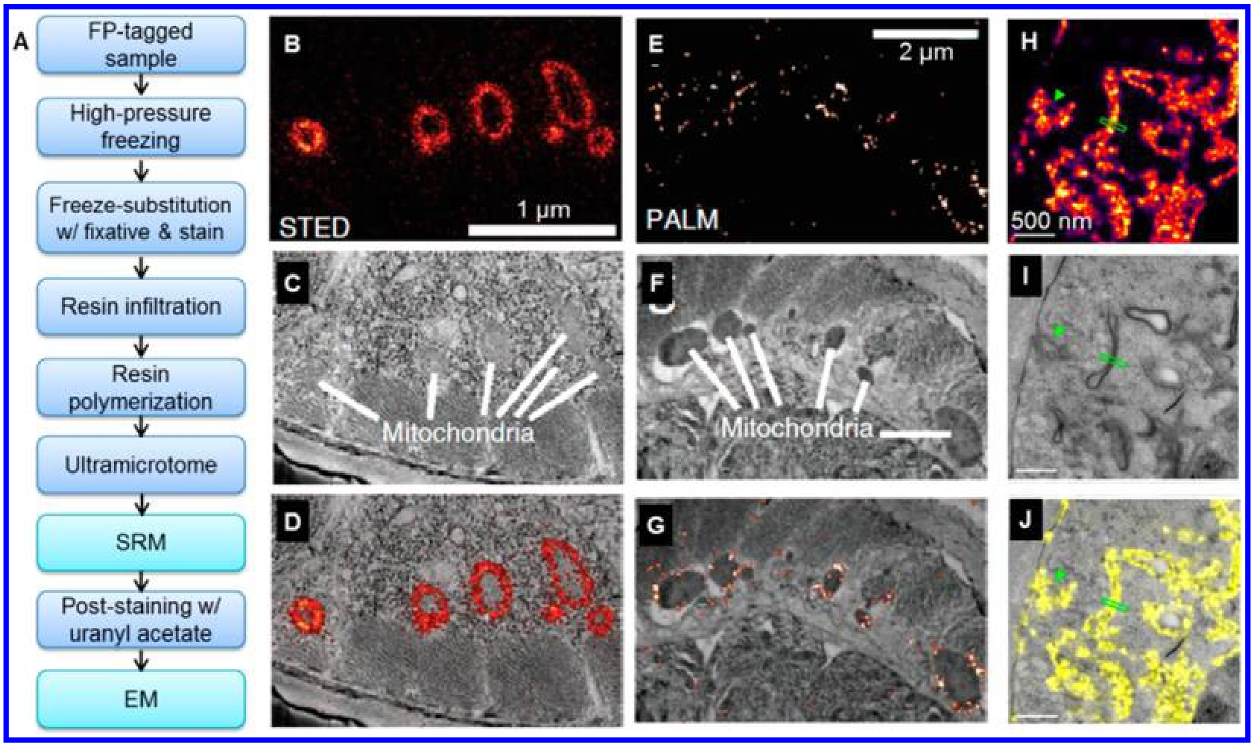


**Fig. S38.**

**Correlative SRM−EM of FP-tagged samples through resin embedding and sectioning. (A)** Flowchart of typical procedures. **(B−D)** Correlative (B) STED and (C) SEM images of a GMA-embedded ultrathin (120 nm) section of a nematode worm expressing TOM20−Citrine; (D) overlaid image. **(E−G)** Correlative (E) PALM and (F) SEM images of an ultrathin (70 nm) LR White section of another worm expressing TOM20−tdEos; (G) overlaid image. (B−G) Adapted from refs (*17*) and (*98*). **(H−J)** Correlative (H) SMLM and (I) TEM images of an ultrathin (100−150 nm) section of Lowicryl HM20-embedded HEK-293T cells tagged by EphA2-mVenus; (J) overlaid image. (H−J) Adapted from refs (*17*) and (*42*).

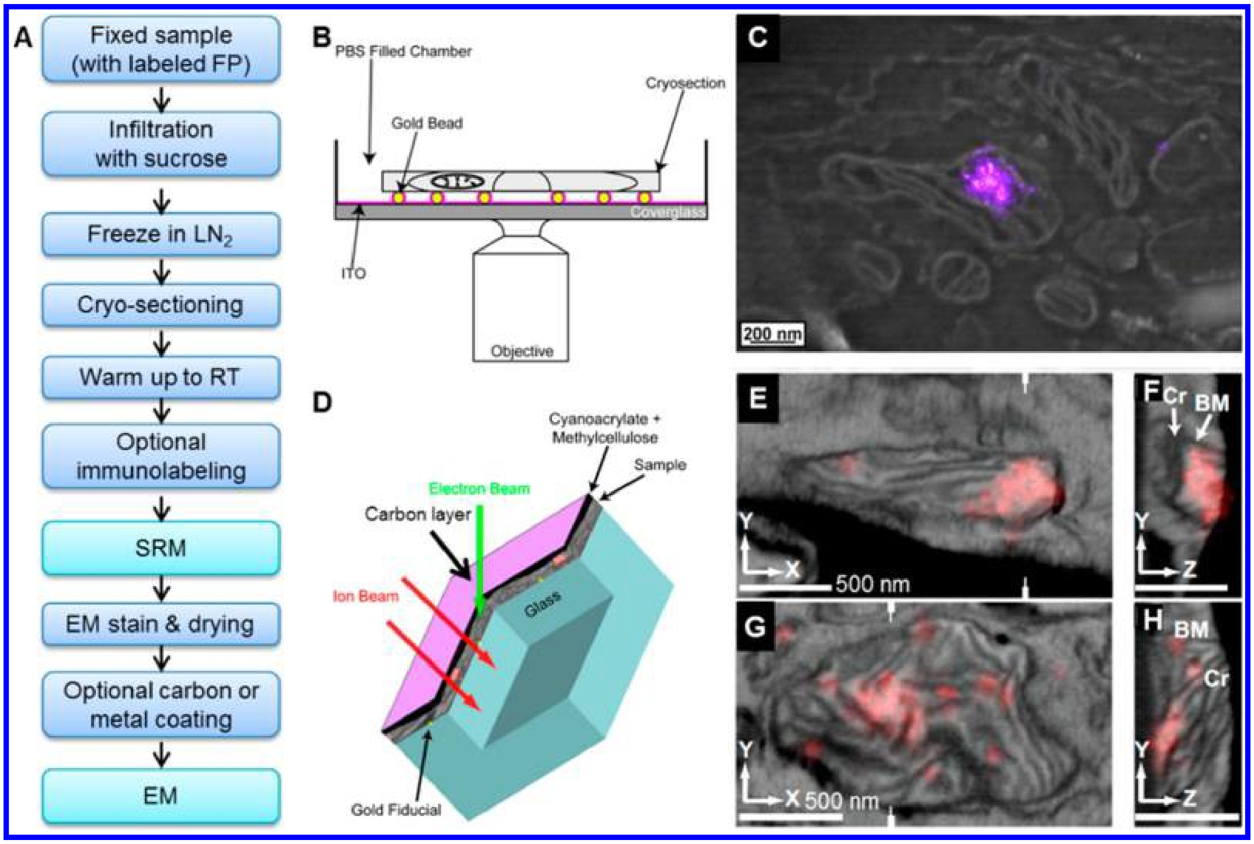


**Fig. S39.**

**Correlative SRM−EM of cryosectioned samples. (A)** Flowchart of typical procedures. **(B)** Schematic of sample geometry for correlative SRM−SEM of a cryosection on an ITO-coated coverslip. **(C)** Thus-obtained correlated SEM (grayscale)−PALM (magenta) data of a ~100-nm-thick cryosection of mouse fibroblast 3T3 cells tagged with TFAM-mEos2, showing that TFAM-mEos2 resides in the mitochondrial matrix surrounded by boundary and cristae membranes. (B, C) Adapted from refs (*17*) and (*96*). **(D)** Schematic of sample geometry for FIB−SEM of a cryosection on coverslip after iPALM SRM. Cryosection sample (gray) is coated with cyanoacrylate and methylcellulose (black), to smooth the topography of the cells and ensure more uniform ion milling, and covered with a thin layer of carbon (purple) for electrical conductivity. SEM images are taken at different depths as the ion beam mills through the sample to construct a 3D image. **(E−H)** Correlated and overlaid images of 3D FIB-SEM (grayscale) and 3D iPALM (red) data of a ~500-nm-thick cryosection of 3T3 cells tagged with TFAM-mEos2. (E, G) Slices in the x−y plane; (F, H) slices in the y−z plane along the hatch marks in panels E and G. Labels denote the mitochondrial boundary membrane (BM) and cristae (Cr). (D−H) Adapted from refs (*17*) and (*95*).

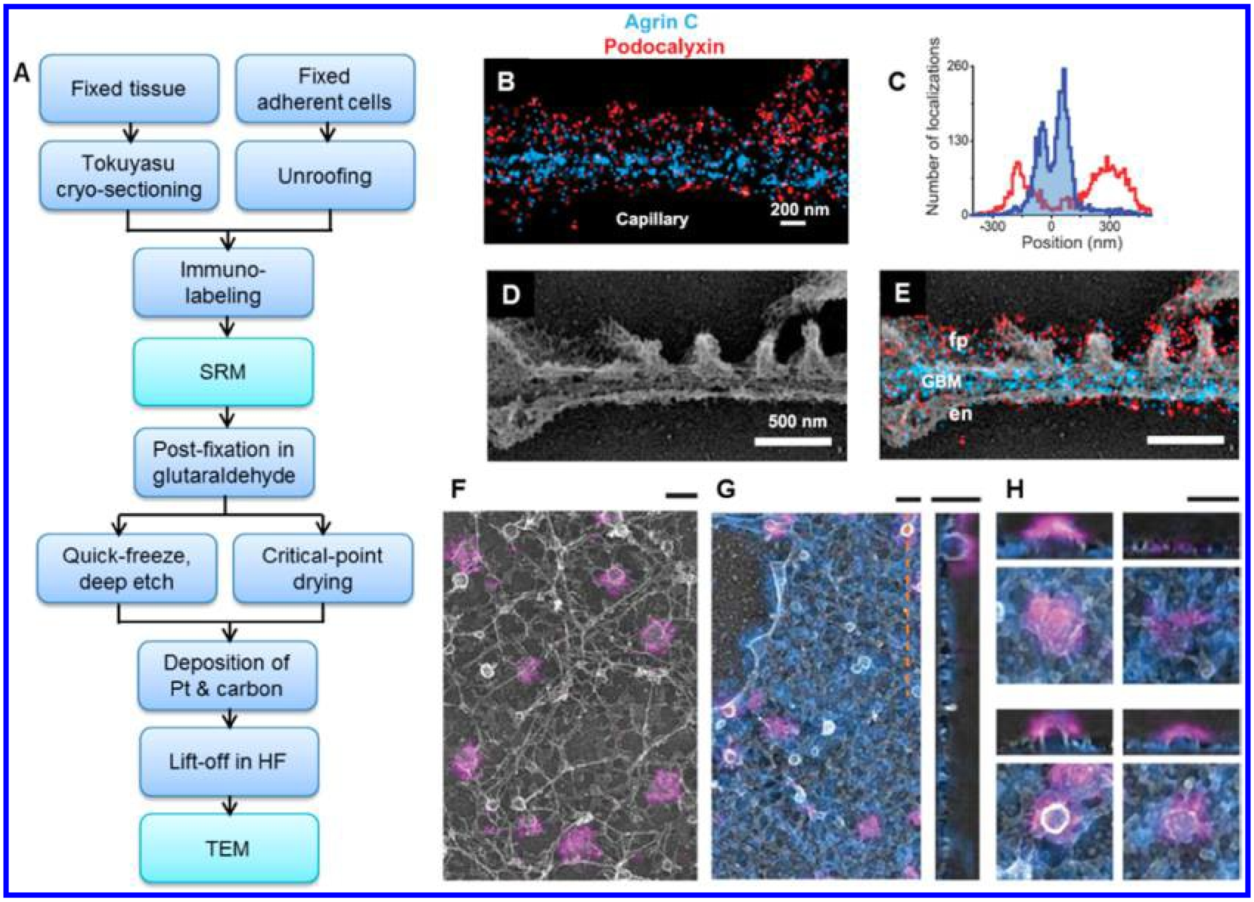


**Fig. S40.**

**Correlative SRM and metal-replica TEM. (A)** Flowchart of typical procedures. **(B)** Two-color STORM and **(C)** cross-sectional quantification of immunolabeled agrinC (blue) and podocalyxin (red) along a capillary region of a cryosection of mouse kidney tissue. **(D)** TEM of platinum deep-etch replica prepared from the same section. **(E)** Overlay of STORM and EM images shows ultrastructural features such as podocyte foot processes (fp), endothelial cells (en), and glomerular basement membrane (GBM). (B−E) Adapted from refs (*17*) and (*136*). **(F)** iPALM image of Alexa Fluor 647-labeled clathrin (magenta) at the inner bottom surface of an unroofed cell, correlated and overlaid with the metal-replica TEM image of the same surface (grayscale). **(G)** (Left) Two-color iPALM results of membrane- targeted myristoylated psCFP2 (blue) and clathrin−Alexa Fluor 647 (magenta) mapped onto the z projection of a TEM tomogram (grayscale). (Right) Magnified z slice along the orange dashed line. **(H)** Individual clathrin structures are shown in xy (z projection) and yz dimensions (tomogram slice). Scale bars: 200 nm (F−H). (F−H) Adapted from refs (*17*) and (*99*).

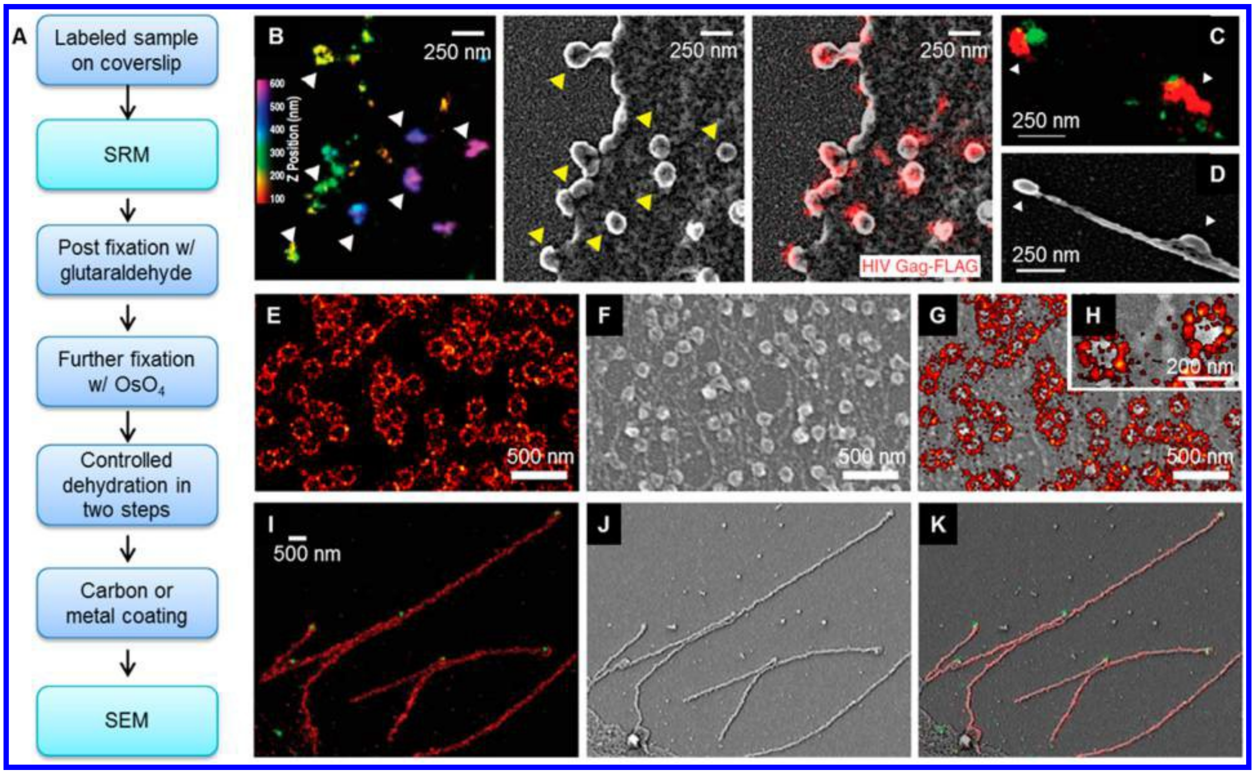


**Fig. S41.**

**Correlative SRM−SEM of unsectioned samples. (A)** Flowchart of typical procedures. **(B)** Correlative iPALM (left; colored for z) of COS- 7 cells expressing HIV Gag−FLAG immunolabeled with Alexa Fluor 647 and SEM (middle) of the same area. Overlay data are shown on the right. **(C, D)** Correlative (C) two-color iPALM and (D) SEM of two virus-like particles (white arrowheads) emanating from a cell expressing Gag−FLAG (red) and PSCFP2−CHMP2A (green). (B−D) Adapted from refs (*17*) and (*101*). (E−H) Correlative dSTORM-SEM of nuclear pore complexes. **(E)** dSTORM image of the integral membrane protein gp210 immunolabeled by Alexa Fluor 647. **(F)** Corresponding SEM image of the nucleoplasmic side of the nuclear envelope, visualizing nuclear baskets. **(G)** Overlaid image. **(H)** Zoom-in of panel G. Adapted from refs (*17*) and (*100*). (E−H). **(I−K)** Correlative two-color STORM (I) and SEM (J) images of budding Udorn virus filaments immunolabeled for M1 (labeled by Alexa Fluor 647; red) and vRNP (labeled by Alexa Fluor 568; green). (K) Overlaid image. (I−K) Adapted from refs (*17*) and (*39*).

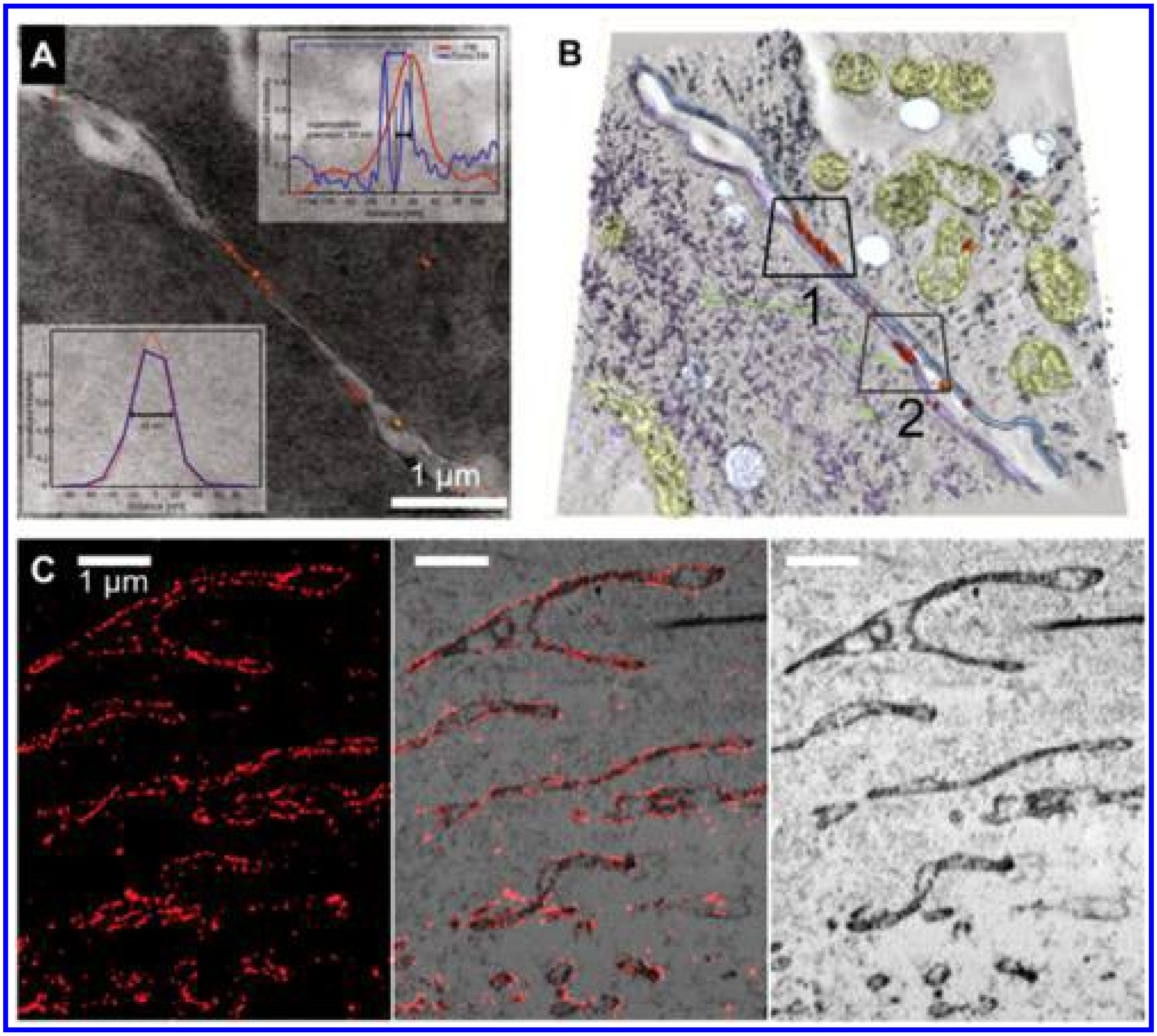


**Fig. S42.**

**Correlative SRM−EM of dye-labeled samples through resin embedding and sectioning. (A)** Superimposed images of correlated dSTORM of SNAPf−N-cadherin labeled with Alexa Fluor 647 (orange) and TEM micrograph for a 300-nm-thick Lowicryl HM20- embedded section of cultured L cells. **(B)** Overlay of dSTORM result (orange) with ET of the sample, with volume representation of intracellular organelles: mitochondria (yellow), vesicles (light blue), microtubules (green), and cytoplasmic compounds (violet). (A, B) Adapted from refs (*17*) and (*97*). **(C)** Correlative STORM and EM images of a 70-nm-thick UltraBed-embedded section of cultured BS-C-1 cells immunostained with Alexa Fluor 647 for TOM20. (Left) STORM image; (right) SEM image; (middle) overlaid image. (C) Adapted from refs (*17*) and (*39*).

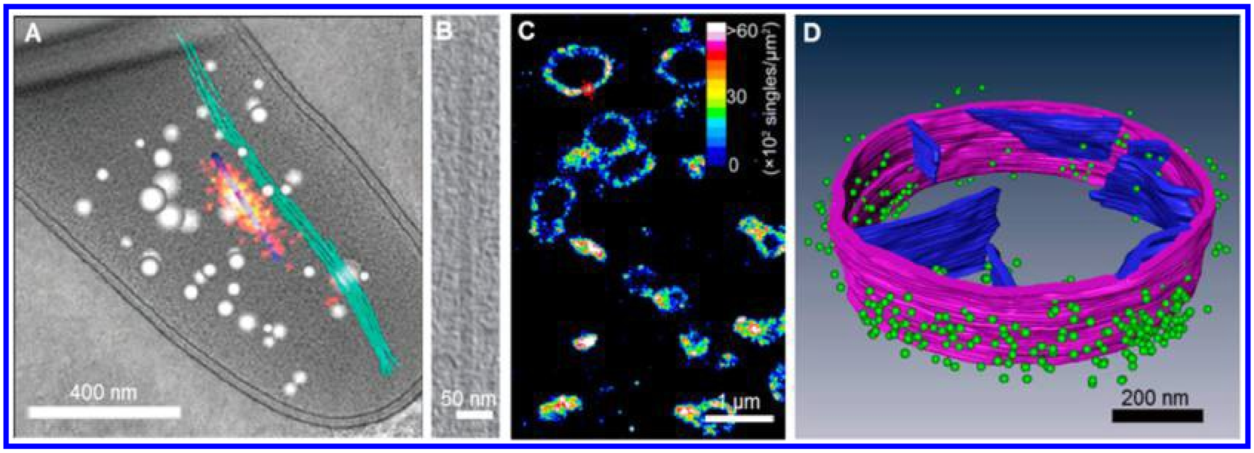


**Fig. S43.**

**Correlated cryo-PALM and cryo-ET. (A)** Virtual slice from a high-resolution 3D cryotomogram of plunge-frozen M. xanthus (grayscale) correlated with a cryo-PALM image of VipA−PA-GFP (red and yellow). Cellular substructures in the cryotomogram were categorized into different components: tubular structure (blue), filamentous bundles (green), and spherical granules (white). **(B)** Zoom-in of the cryo-ET result of the T6SS tubular structure as identified through the correlated cryo-PALM image. (A, B) Adapted from refs (*17*) and (*30*). Copyright 2014 Macmillan Publishers Limited. **(C)** Cryo-PALM image of a 200 nm vitreous slice of HEK293 cells labeled by Dronpa-tagged TOM20, a marker for the mitochondrial outer membrane. **(D)** Three-dimensional reconstruction of correlated cryo-SRM and cryo-ET for a vitreous section of a HEK293 cell expressing TOM20-Dronpa. Mitochondrial outer membrane (purple) and cristae (blue) are identified from cryo-ET data; Dronpa tag molecules (green) are from the brightest (top ~10%) single molecules in the cryo-PALM data. (C, D) Adapted from refs (*17*) and (*7*).

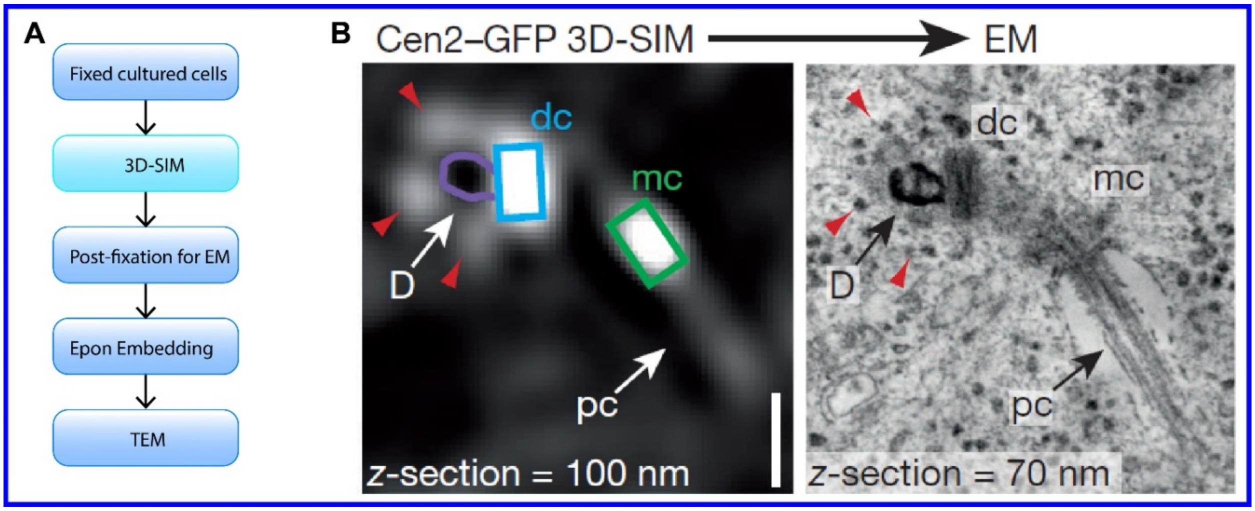


**Fig. S44.**

**Correlative 3D-SIM-TEM of cultured cells. (A)** Flowchart of the procedure. **(B)** Correlative 3D-SIM z-plane (left) and TEM (right) of a centrosome showing an incomplete procentriole loaded deuterosome connected to the daughter centriole. Purple line delineates TEM deuterosome. D, deuterosome, dc, daughter centriole; mc, mother centriole; pc: primary cilium. Adapted from ref (*137*).

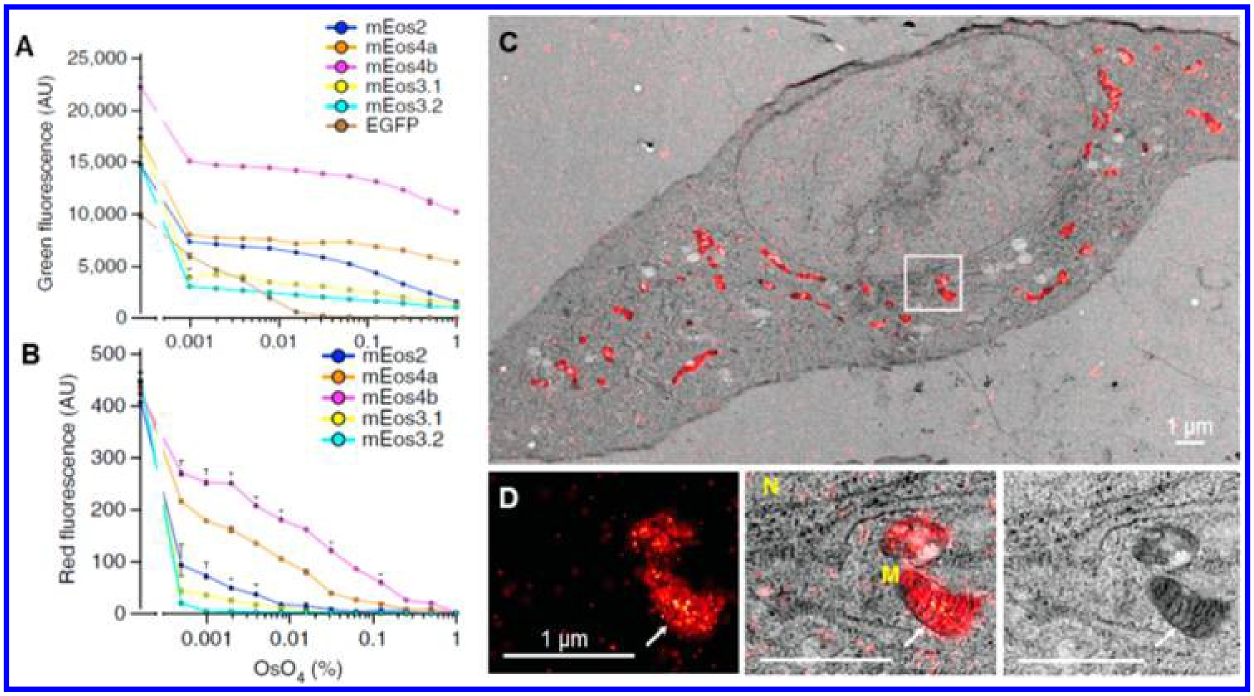


**Fig. S45.**

**Correlative SRM−EM of resin-embedded and sectioned samples by use of fixation-resistant FPs. (A, B)** Fluorescence retention capability of different FPs in vitro at different concentrations of osmium tetroxide, for green (prephotoconversion, A) and red (after-photoconversion, B) fluorescence signals, respectively. **(C)** Correlative PALM (red) and TEM (grayscale) images of a 60 nm GMA section of 3T3 cells expressing mitochondrially targeted mEos4a; sample was fixed by 0.5% osmium tetroxide. **(D)** PALM, overlaid, and TEM images of boxed area in panel C. Arrows indicate a mitochondrial cristae fold. N, nucleus; M, mitochondrion. Adapted from refs (*17*) and (*41*).

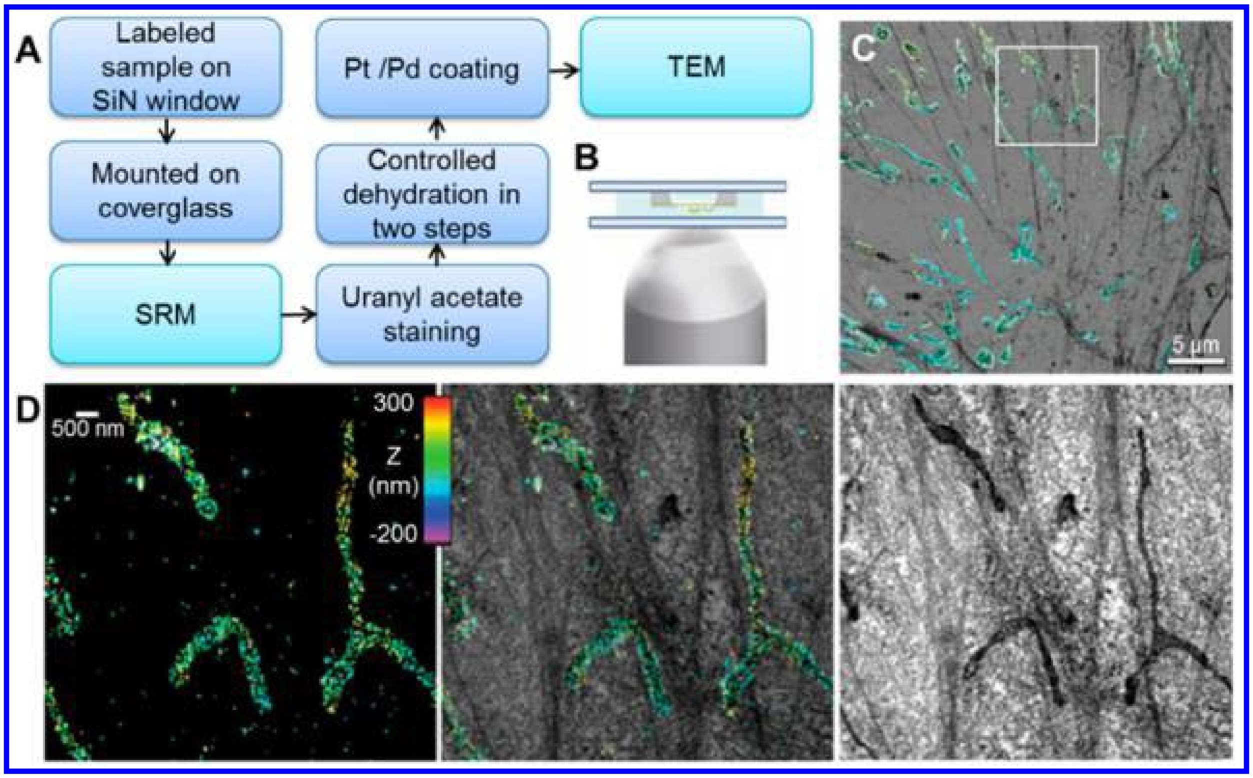


**Fig. S46.**

**Correlative SRM−TEM of unsectioned samples. (A)** Flowchart of typical procedures. **(B)** Scheme of sample mounting geometry for SRM of cells cultured on a SiN window. **(C)** Overlaid correlative 3D STORM (colored for z) and TEM (grayscale) images of mitochondria in a BS-C-1 cell immunostained for TOM20. **(D)** Zoom-in of the boxed regions in panel C, shown as both overlaid and separated STORM and TEM images. Adapted from refs (*17*) and (*39*).

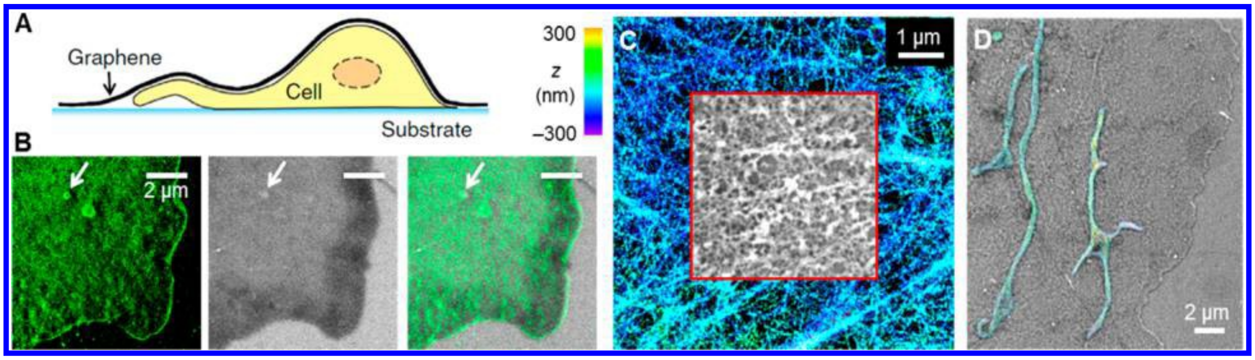


**Fig. S47.**
**Correlative SRM−SEM of wet cell samples via graphene encapsulation. (A)** Schematic: wet cell sample on a substrate (e.g., a coverslip) is covered by graphene to allow direct SEM in a conventional setup without dehydration. **(B)** Correlative STORM and graphene-based SEM results on an unstained wet cell. COS-7 cells on coverglass were membrane-labeled with a lipophilic stain, DiI, and fixed by 4% paraformaldehyde. After STORM of DiI (left), sample was encapsulated by graphene for SEM (center) without metal staining or dehydration. Arrow points to a vesicle that is visualized in both modalities due to local internalization of cell membrane. **(C)** Correlated 3D-STORM (colored for z) and graphene-based EM (black-and-white; center) of the phalloidin-labeled actin cytoskeleton in a wet, membrane-extracted COS-7 cell stained with tannic acid and uranyl acetate. **(D)** Correlated and overlaid 3D-STORM and graphene SEM images of a wet COS-7 cell. For STORM, the sample was immunolabeled for TOM20, a mitochondrial outer-membrane marker. For graphene SEM, the sample was stained with uranyl acetate for membrane contrast. Adapted from refs (*17*) and (*138*).

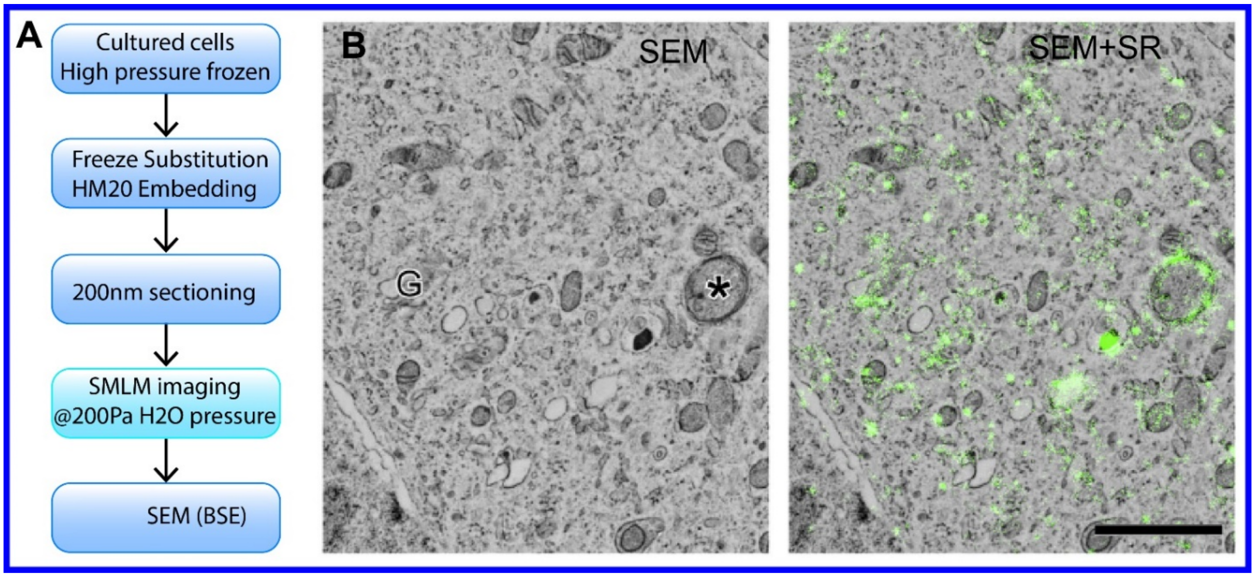


**Fig. S48.**

**Correlative SMLM-SEM of resin-embedded cultured cells in integrated LM-EM microscope. (A)** Flowchart of the procedure. **(B)** SEM image (left) and overlay of SMLM and SEM showing localization of the GFP-C1 construct. G: Golgi, Asterisk: putative autophagosome. Scale bar 2 mm. Adapted from ref (*40*).

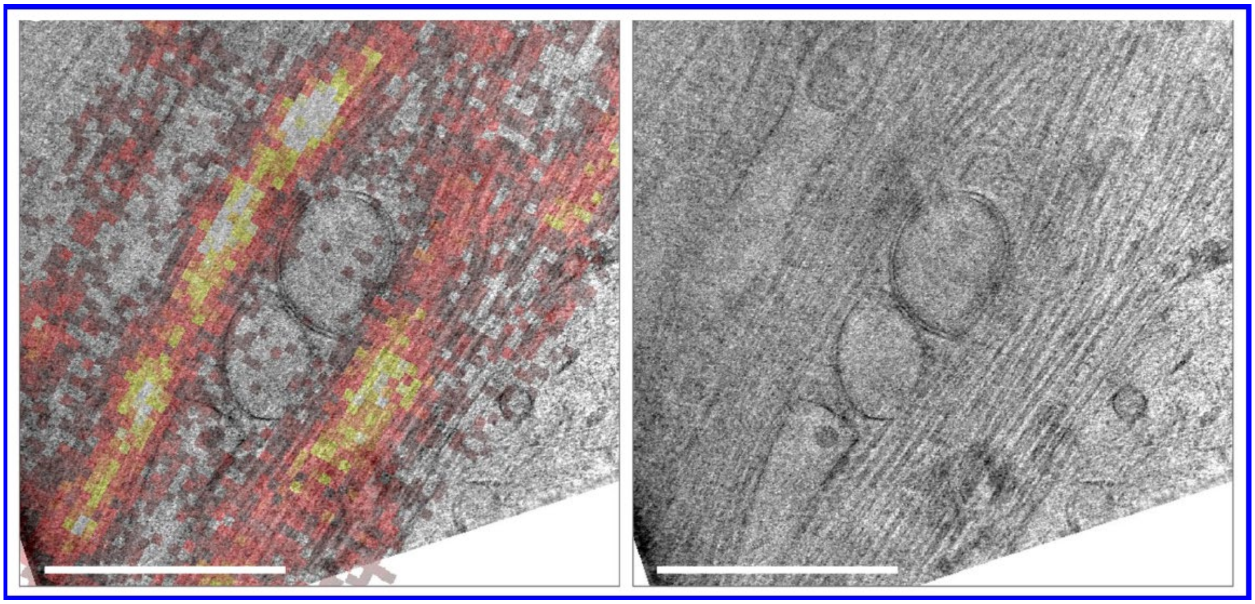

**Fig. S49.**
**Correlated cryo-PALM and cryo-ET on U2OS cells, transfected with rsEGFP2-MAP2.** Scale bars 1um. Adapted from ref (*29*).

#### Table S1.

| *EM method* | *SRM method* | *Fixation* | *Sample Processing* | *SRM label* | *Nature of sample* | *EM quality* | *SRM quality* | *Reference* |
|---|---|---|---|---|---|---|---|---|
| *TEM* | PALM | 4% PFA + 0.5% GA | Tokuyasu Cryo-sectioning | dEosFP | Monkey cell line (COS-7) | Poor ultrustructure preservation | Good | Betzig et al. 2006 (*94*) Fig. S37 |
| *SEM (BSE)* | STED, PALM | Cryo (HPF) | FS, Acrilic RE, sectioning | Citrine (STED), Dendra and tdEos (PALM) | C. elegans worms | Poor ultrustructure preservation | Limited fluorophore survival | Watanabe et al. 2011 (*98*) Fig. S38 |
| *FIB-SEM (BSE)* | iPALM | 4% PFA + 2% GA | Tokuyasu Cryo-sectioning, FIB | mEos2 | Mouse cell line (3T3sw) | Poor ultrustructure preservation | Good | Kopek et al. 2012 (*95*) Fig. S39 |
| *SEM (BSE)* | PALM | 4% PFA + 2% GA | Tokuyasu Cryo-sectioning, metal coating | mEos2, PS-CFP2, caged dye | Mouse cell line (3T3sw) | Poor ultrustructure preservation | Good | Kopek et al. 2013 (*96*) Fig. S39 |
| *Pt-replica, TEM* | STORM | 4% PFA, 3% PFA +0.05% GA | Tokuyasu Cryo-sectioning, quick freeze–deep etch, Pt- replica | Alexa 647 | Mouse and human kidney tissue | Limited to membranous capillary structures | Good | Suleiman et al. 2013 (*136*) Fig. S40 |
| *Pt-replica, TEM, ET* | iPALM | Unroofed by sonication, then 2% PFA + 0.04% GA | unroofing, CPD, Pt-replica | psCFP2, Alexa 647 | Rat cell line (PC12-GR5) | Limited to membrane bound structures | Good | Sochacki et al. 2014 (*99*) Fig. S40 |
| *Pt-replica, TEM,* | dSTORM | Unroofed by sonication, then 4% PFA, or 2% PFA+2% GA | unroofing, HMDS drying, Pt-replica | Alexa 488, Alexa 647 | Primary rat hippocampal neurons (cultured) | Limited to structures at the exposed surface | Good | Vassilopoulos et al. 2019 (*102*) |
| *SEM (SE)* | iPALM | 4% PFA + 0.2% GA + 0.2% Triton | Post staining, CPD, metal coating | psCFP2, Alexa 647 | Monkey cell line (COS-7) | Limited to surface morphology | Good | Van Engelenburg et al. 2014 (*101*) Fig. S41 |
| *TEM, ET* | dSTORM | Cryo (HPF) | FS, Acrilic RE, sectioning | Alexa 647, SiR | Mammalian cell lines (L cells, HeLa) | Poor ultrustructure preservation | Good | Perkovic et al. 2014 (*97*) Fig. S42 |
| *Cryo-ET* | Cryo-PALM | Cryo (plunge-freezing) | | PA-GFP | Bacteria (*M. xanthus*) | Good | Limited labeling density | Chang et al. 2014 (*30*) Fig. S43 |
| *SEM (SE)* | dSTORM | 2% PFA | CPD, carbon coating | Alexa 647 | X. laevis oocytes | Limited to membrane bound structures | Good | Löschberger et al. 2014 (*100*) Fig. S41 |
| *TEM* | 3D-SIM | Methanol 0.5% Triton | Post fixation + staining; Epon RE, sectioning | GFP, DAPI | Primary mouse ependymal progenitors | good (SIM only - fast) | Unclear preservation other than centrosomes | Al Jord et al. 2014 (*137*) Fig. S44 |
| *TEM, SEM (BSE)* | PALM | 4% PFA + 0.2% GA + HPF) followed by 1% UA + 0.5% $OsO_4$ | FS, Acrilic RE, sectioning | mEos4a, mEos4b | Mouse cell line (3T3) | good | Limited fluorophore survival | Paez-Segala et al. 2015 (*41*) Fig. S45 |
| *TEM* | SMLM | Cryo (HPF), followed by 0.2% UA, 0-0.1% TA during FS | FS, Acrilic RE, sectioning | mGFP, mVenus, mRuby2 | Human cell line (HEK293T) | poor ultrustructure preservation | Good | Johnson et al. 2015 (*42*) Fig. S38 |

| | | | | | | | |
|---|---|---|---|---|---|---|---|
| *TEM, SEM (BSE, SE)* | 3D and 2D STORM | 4% PFA + 0.1% GA + 0.2% Triton | CPD, metal coating; RE, sectioning | Alexa 647, Alexa 568 | Mammalian cell lines (BS-C-1, A549), influenza virus | poor ultrustructure preservation | Good | Kim et al. 2015 (*39*) Fig. S41 and S46 |
| *SEM (SE)* | 3D and 2D STORM | 4% PFA or 0.3% GA + 0.25% Triton | Graphene encapsulation | Alexa 647, Cy3B, CM-DiI | Monkey cell line (COS-7) | Limited to surface morphology | Good | Wojcik et al. 2015 (*138*) Fig. S47 |
| *Cryo-ET* | Cryo-PALM | Cryo (HPF) | Vitreous sectioning | Dronpa | Human cell line (HEK293) | good | Limited labeling density | Liu et al. 2015 (*7*) Fig. S43 |
| *SEM (BSE)* | SMLM @200 Pa of $H_20$ | 4% PFA + Cryo | FS, Acrylic RE, sectioning, integrated LM-SEM | YFP, GFP | Human cell line (HeLa) | poor ultrustructure preservation | Good | Peddie et al. 2017 (*40*) Fig. S48 |
| *Cryo-ET* | Cryo-PALM | Cryo (plunge-freezing) | | rsEGFP2 | Human cell line (U2OS) | Good | Good | Tuijtel et al. 2019 (*29*) Fig. S49 |

SR-CLEM techniques, adapted from 1, modified and expanded. Cryo SR-CLEM techniques are shaded blue.

BSE – back-scattered electron (detector)

HPF – high-pressure freezing

FS – freeze-substitution (usually along with chemical fixation)

CPD – critical point drying

RE – resin embedding

**Table S2.**

| *Fluorophore* | Excitation Max (nm) | Emission Max (nm) | Temperature at which suitable for Cryo-SMLM | Notes |
|---|---|---|---|---|
| *EGFP* | 488 | 507 | 77 | c.f. Fig. 1 |
| *pHluorin* | 475 | 509 | 77 | Similar results to eGFP and mEmerald. Retains pH sensitivity. |
| *mEmerald* | 487 | 509 | 77 | c.f. Fig. 1 |
| *mEos3.2 Green* | 507 | 516 | 77 | No switching between green and red states observed. Green state blinks |
| *Dronpa* | 503 | 518 | 77 | |
| *TagYFP* | 508 | 524 | 77 | c.f. Fig. 1 |
| *mVenus* | 515 | 527 | 77 | Two emissive states: one with 488 nm and another with 532 nm excitation. We observed that the two states can convert to one another under illumination. The kinetics of this conversion is on the order of minutes with ~kW/cm^2 intensities. |
| *JF525* | 525 | 549 | 4 | c.f. Fig. 1 |
| *mOrange2* | 549 | 565 | Not tested | Two emission bands, one at the normal RT wavelengths and one overlapping with GFP |
| *JF549* | 549 | 571 | 4* | Almost 4 days of constant illumination required before acceptable blinking is achieved. c.f. Fig. 1 |
| *mEos3.2 Red* | 572 | 580 | N/A | No switching between green and red states observed. Red state does not blink |
| *mRuby3* | 558 | 592 | N/A | No photoswitching observed |
| *mApple* | 568 | 592 | N/A | No photoswitching observed |
| *JF585* | 585 | 609 | N/A | No photoswitching observed |
| *mCherry* | 587 | 610 | 4* | Poor results at 4K (c.f. Fig. 1). No photoswitching observed at 77K |

| | | | | |
|---|---|---|---|---|
| *mKeima* | 440 | 620 | N/A | Intense emission in the GFP channel (488 nm excitation and 520/35 emission) along with dark state shelving and blinking. No blinking was observed in the room temperature emission channel |
| *PA-mKate* | 586 | 628 | N/A | Strong blue channel (488 nm excitation) emission. Unsuitable for two color imaging. Blinking in the red channel (561 nm excitation) poor but not quantified. |
| *TagRFP657* | 611 | 657 | N/A | No photoswitching observed |
| *mCardinal* | 604 | 659 | N/A | No photoswitching observed |
| *JF646* | 646 | 664 | N/A | No photoswitching observed |
| *Alexa Fluor 647* | 650 | 665 | N/A | No photoswitching observed |
| *DiSC3(5)* | 651 | 675 | N/A | Tested at 25% of the power reported in (*34*). Very little reduction in emission over a four-hour period. |

Excitation and emission wavelengths were collected from fpbase.org (*139*).

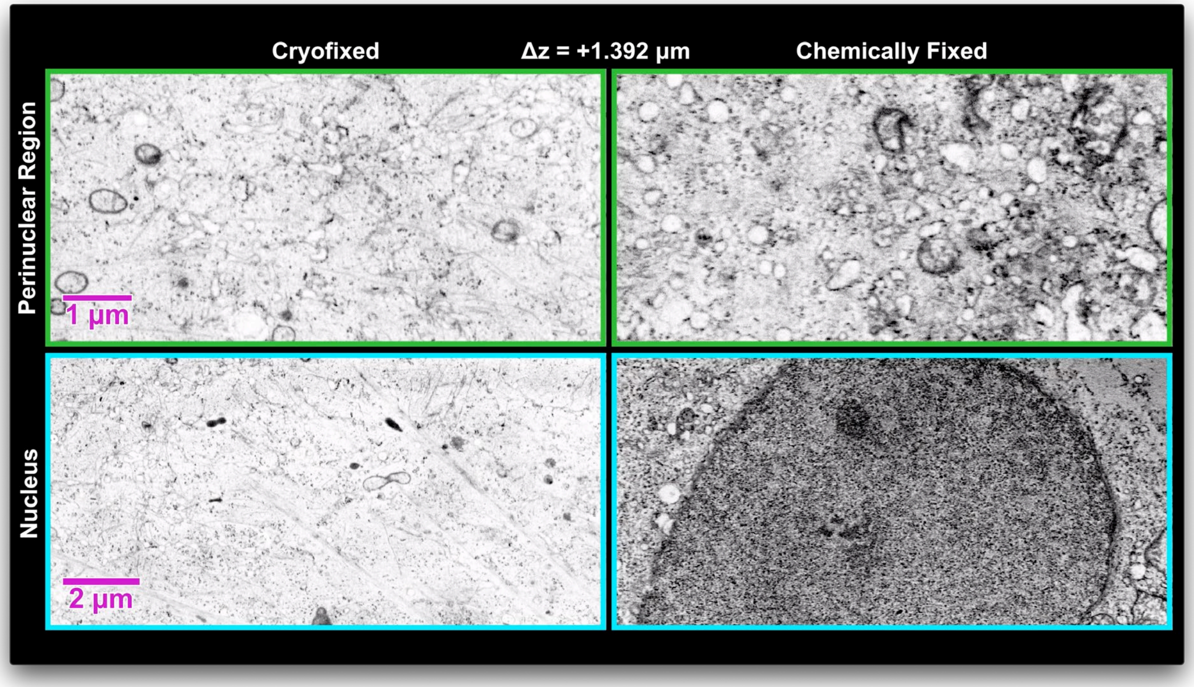


**Movie S1.**

**Comparison of EM ultrastructure in cryofixed versus chemically fixed adherent cells.** XY orthoslices through FIB-SEM volumes of a high-pressure frozen cell (left) and a cell fixed with 4% PFA and 0.1% glutaraldehyde in the perinuclear (top) and nuclear (bottom) regions. See also supplementary note 2b. Distance from the static views shown in fig. S3 is shown at top (Fig. 1, E, F, K, L).

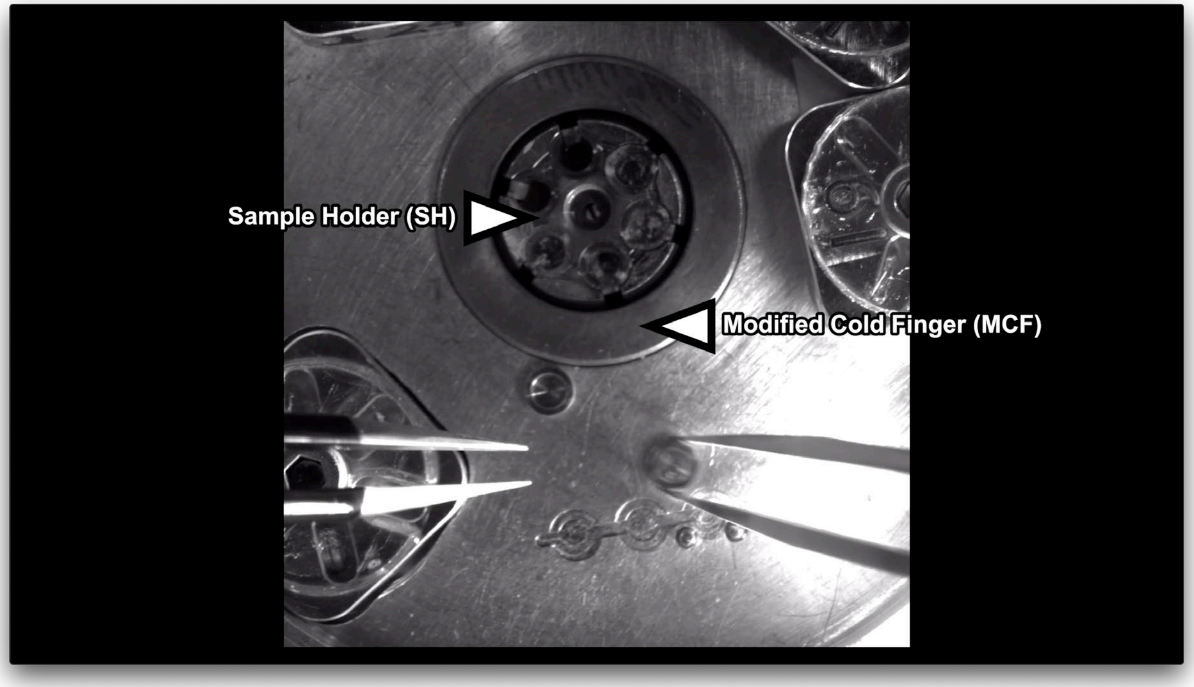


**Movie S2.**
**Coverslip cleaning and loading procedure.** Video recorded through a Nikon stereoscope of planchette removal, coverslip cleaning and coverslip loading as described in supplementary note 2c.

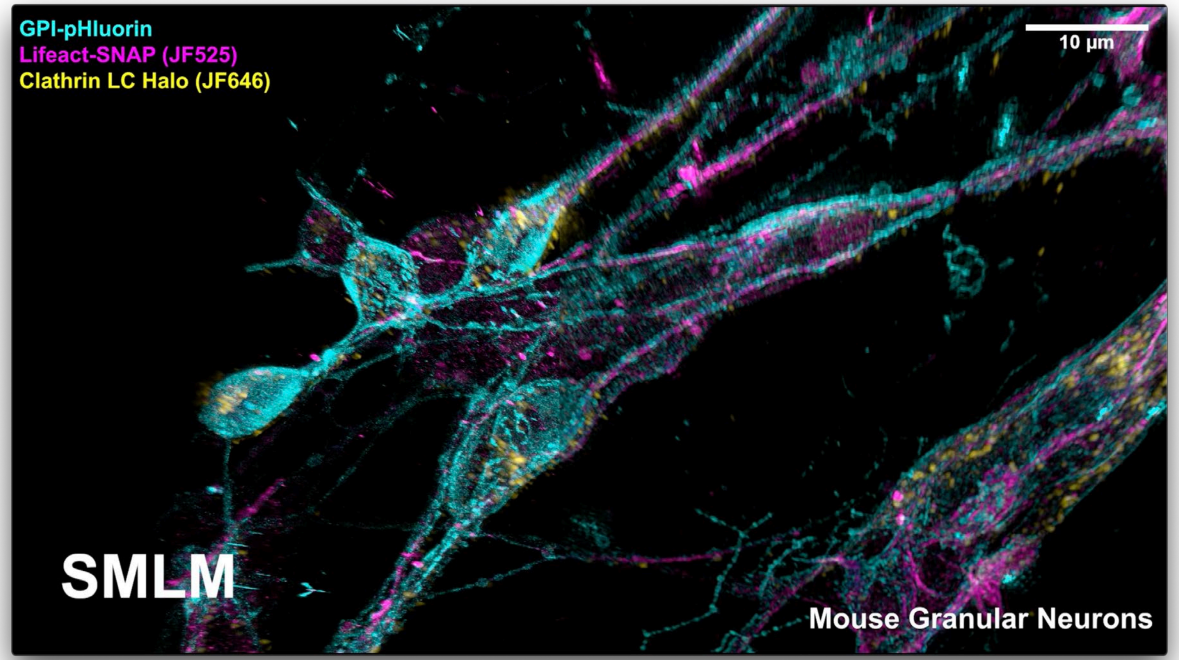


**Movie S3.**
Three color Cryo-SIM correlated with two color SMLM of high pressure frozen mouse granular neurons expressing GPI-pHluorin, Lifeact-SNAP conjugated to JF525, and Clathrin LC Halo conjugated to JF646.

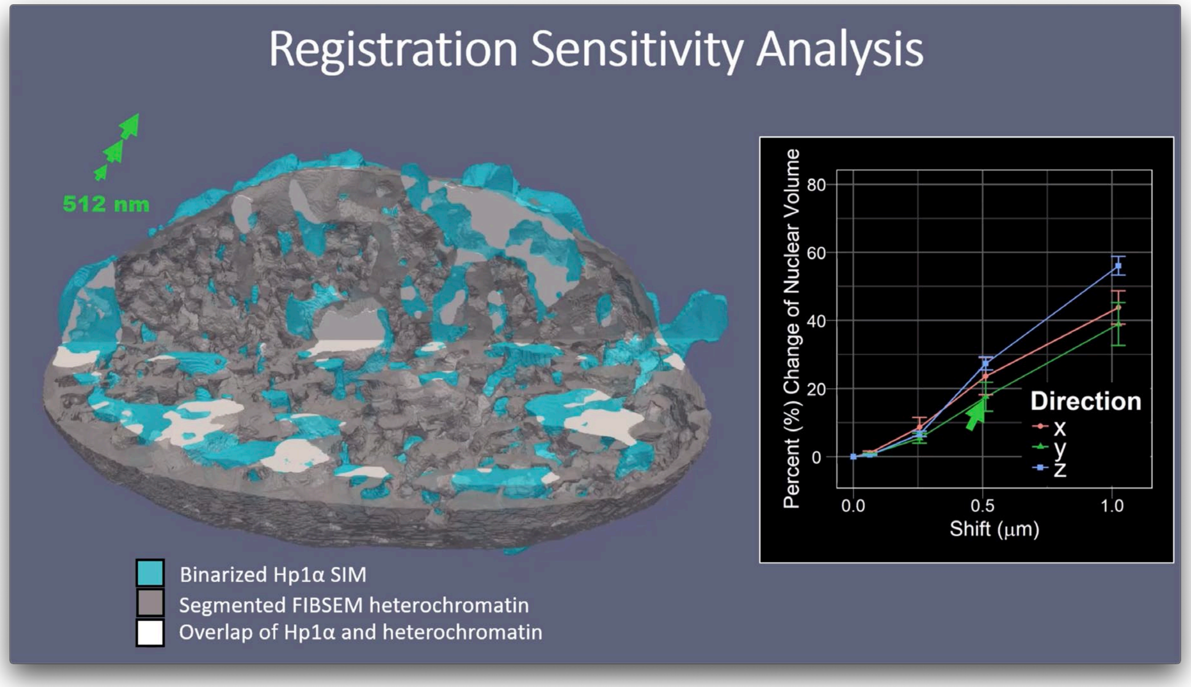


**Movie S4.**
Animated sensitivity analysis (c.f. supplementary note 12).

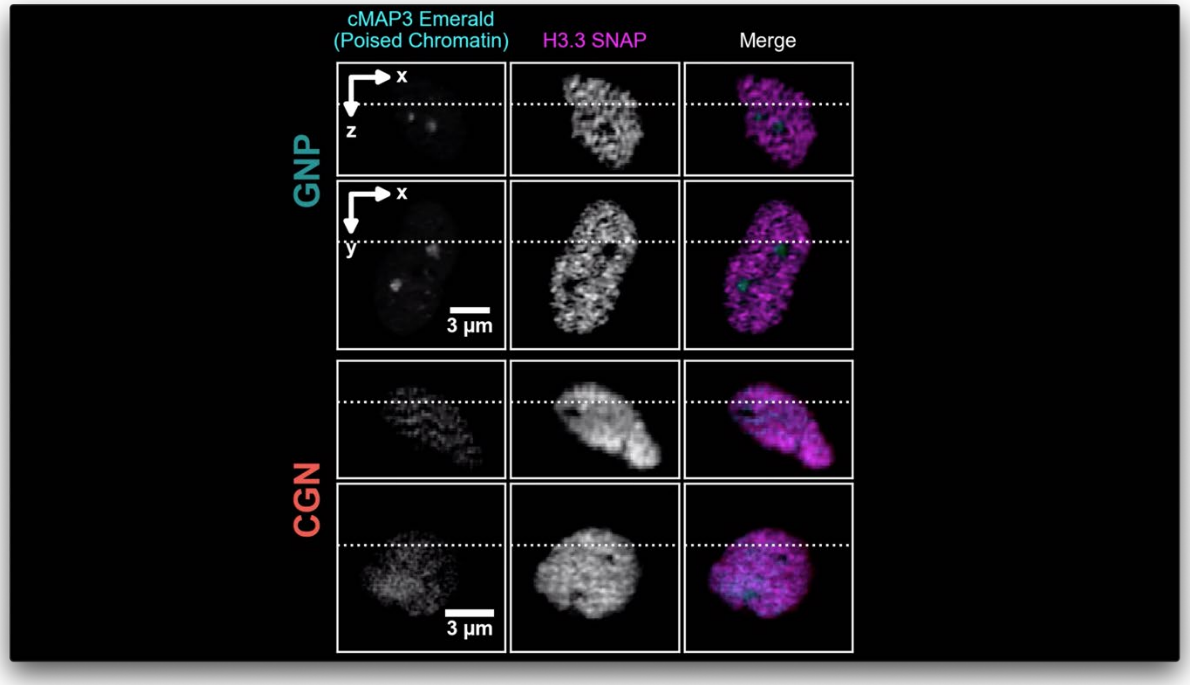


**Movie S5.**
Animated orthoslices for Fig. S24.

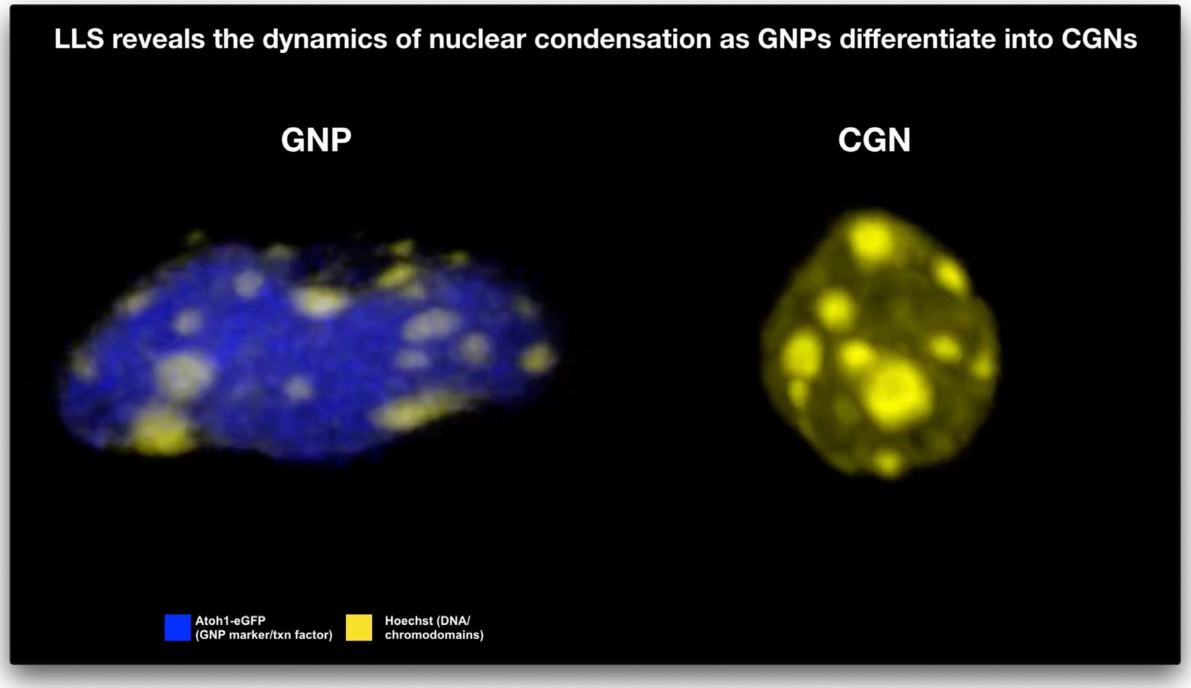


**Movie S6.**
Animated videos of GNP and CGN nuclei for Fig. 7A.

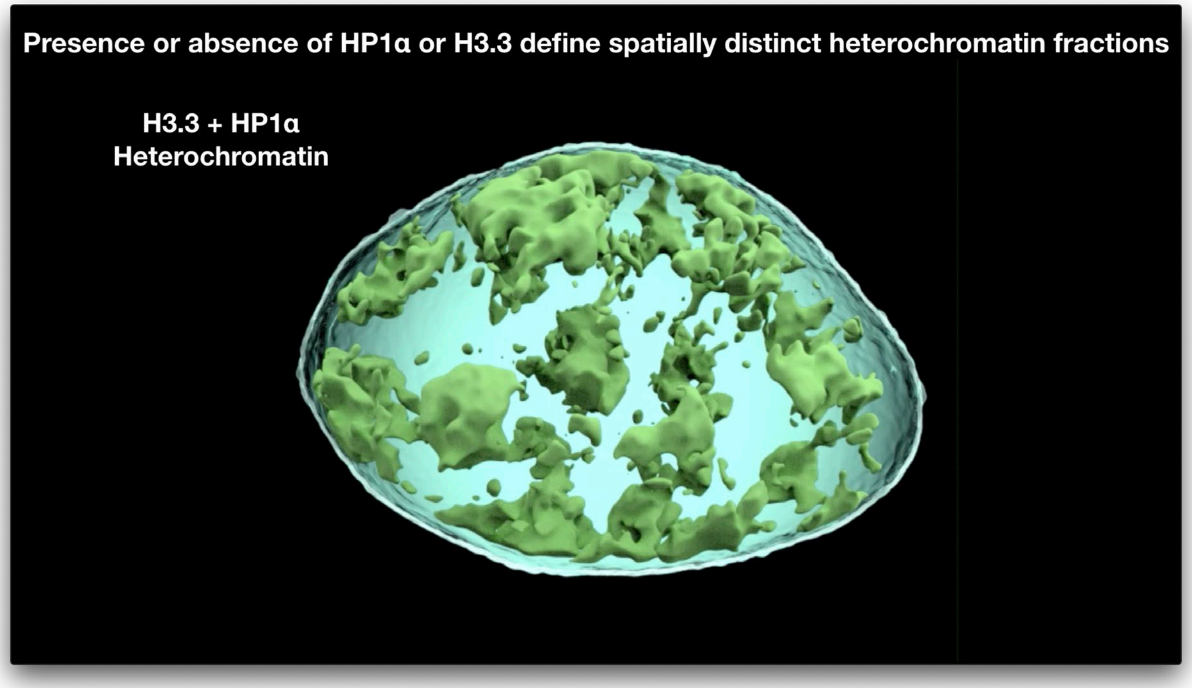


**Movie S7.**
3D surface renderings of CLEM defined heterochromatin subdomains for the CGN nuclei in Fig. S22A.
